# *corto*: a lightweight R package for Gene Network Inference and Master Regulator Analysis

**DOI:** 10.1101/2020.02.10.942623

**Authors:** Daniele Mercatelli, Gonzalo Lopez-Garcia, Federico M. Giorgi

## Abstract

**Motivation:** Gene Network Inference and Master Regulator Analysis (MRA) have been widely adopted to define specific transcriptional perturbations from gene expression signatures. Several tools exist to perform such analyses, but most require a computer cluster or large amounts of RAM to be executed.

**Results:** We developed corto, a fast and lightweight R package to infer gene networks and perform MRA from gene expression data, with optional corrections for Copy Number Variations (CNVs) and able to run on signatures generated from RNA-Seq or ATAC-Seq data. We extensively benchmarked it to infer context-specific gene networks in 39 human tumor and 27 normal tissue datasets.

**Availability:** Cross-platform and multi-threaded R package on CRAN (stable version) https://cran.rproject.org/package=corto and Github (development release) https://github.com/federicogiorgi/corto.

## Introduction

The advent of high throughput methods to quantify transcript abundances has offered the possibility to measure gene co-expression across hundreds of samples and thousands of genes. The principle of co-expression has fueled the generation of several gene regulatory network representations over the past decade, and still constitutes the main source of genome-wide network inference pipelines (Mercatelli et al., 2019a). Recently, scientists have developed tools to further leverage gene network models and interrogate them via MRA to identify regulatory subnetworks active in specific experimental conditions, tumor subtypes or even individual patients (Ding et al., 2018), or to improve the readout in noisy or single-cell datasets by aggregating genes (Mercatelli et al., 2019b).

The majority of current tools for gene network inference require either a computing cluster to be executed and/or a high amount of RAM. Therefore, we developed a lightweight R package dubbed *corto* (“correlation tool”) to infer significant, direct edges between a user-provided list of source genes (“centroids”), such as Transcription Factors (TFs). The algorithm underneath *corto* infers direct TF-target relationships by applying DPI on correlation triplets, as proposed in (Reverter and Chan, 2008). Our tool provides networks as Bioconductor regulon objects that can be immediately used in MRA by corto itself or by other tools, e.g. the VIPER pipeline (Alvarez et al., 2016). We benchmarked *corto* across dozens of transcritpomics datasets, with respect to other analogous tools.

## Functionalities

The *corto* algorithm expands the well-established pipeline of the public Java tool ARACNe-AP (Lachmann et al., 2016), and allows the user to perform downstream MRA and visualization of Master Regulators. The R implementation of its algorithms are fully multithreaded, with a user-friendly progress bar indicating the estimated time to completion. The functionalities of the package can be categorized as follows:

1. Gene Network Inference. This is based on optimized pairwise correlation, DPI and bootstrapping to evaluate the significant edges. Further details on the inference algorithm are available on the package vignettes and on the Github page.
2. CNV correction. As the presence of CNVs can influence and bias the generation of gene networks (Schubert et al., 2019), *corto* gives the optional possibility to use CNV data to correct target expression profiles via linear regression. An example of *corto* CNV-corrected network analysis in the TCGA Glioblastoma (GBM) dataset is provided in the package vignette and in **Supp Figure S1**.
3. Master Regulator Analysis. The mra function within our package calculates the enrichment of each TF-centered network on a user-selected signature, provided as two gene expression matrices (e.g. treatment vs control). This kind of analysis is exemplified in detail in **Supp Vignette 1**.
4. Visualization of Network Enrichment. *corto* can generate MRA plots as visualized in **Figure 1**. For each most significant or user-specified centroid, the function visualizes the distribution of its targets across a signature, as red (positively correlated targets) or blue (negatively correlated) vertical bars. The Normalized Enrichment Score (NES) and the corresponding p-value (based on permutation tests and signature sample shuffling) are shown. Also, the most correlated targets are shown as a mini network, connected to the centroid with pointed arrows (positively correlated) or blunted arrows (negatively correlated), and shown as red or blue if activated or repressed in the provided contrast.

Classic MRA is based on a provided gene network and on a gene expression differential signature, based on Microarrays or RNA-Seq. To run *corto* on these data types, we suggest to use RMA and VST normalization, which have been shown to be optimal for coexpression analyses (Giorgi et al., 2013). On top of these data types, corto can operate also on ATAC-Seq-derived signatures, which measure the differential chromatin status between two conditions, as shown in **Supp Vignette 2**.

**Figure 1.**
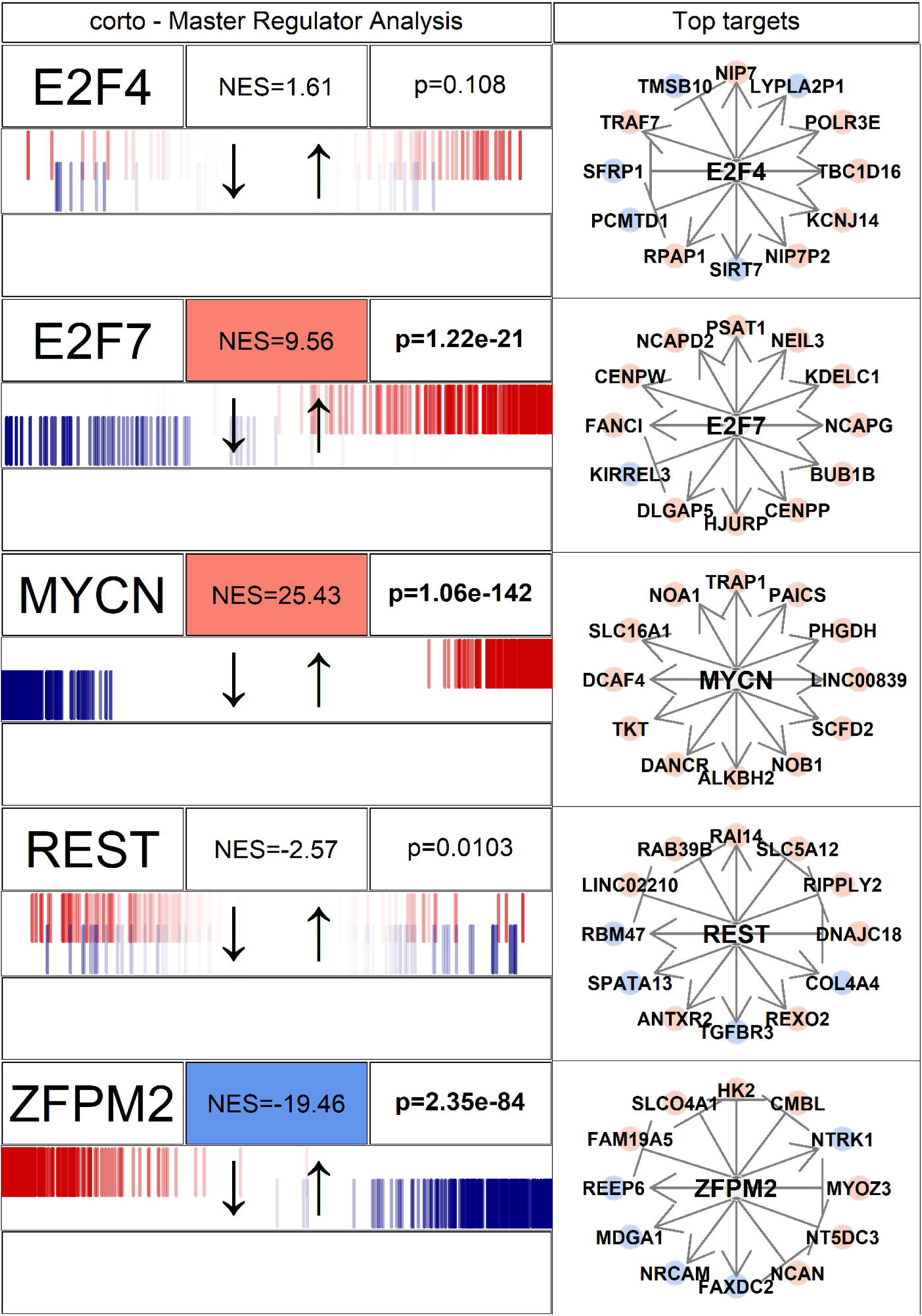
Example of Master Regulator Analysis performed by corto.

## Benchmarking

We tested *corto* extensively against the Java tool ARACNe-AP (Lachmann et al., 2016) and other R-based tools for gene network inference, such as minet (Meyer et al., 2008) and RTN (Castro et al., 2016). Our analysis shows that *corto* is consistently at least 10x faster than its competitors even in single-thread mode, obtaining significantly similar networks (**Supp Vignette 3**). Using the ENCODE ChIP-Seq dataset, we estimated the accuracy of corto, which is consistently above 75% in all tested cell lines (**Supp Benchmark**). We also detected a high similarity in MRA analyses executed with networks generated by *corto* and the other tool generating regulon objects, ARACNe-AP (**Supp Vignette 4**). More examples and details on *corto* and MRA are available in **Supp File S1**. In **Figure 1**, we tested a *corto*-generated Neuroblastoma network (Kocak et al., 2013) on a MYCN amplified vs. not amplified signature and visualize the results for selected TF networks. Unsurprisingly (more details in **Supp Vignette 1**), *corto* shows MYCN network amongst the most upregulated ones, together with the pro-proliferative E2F7, while E2F4 and REST are not affected within significance. On the other hand, the pro-differentiation TF ZFPM1 seems to be downregulated by the MYCN amplification. Top targets for each TF are shown.

## Conclusions

We propose *corto* as a novel, lightweight and robust tool for rapid gene network inference and MRA in large-scale transcriptomics datasets. While benchmarked here in human RNA-Seq and ATAC-Seq data, there is no intrinsic limit to corto applications, which can be naturally extended to other organisms and quantitative biological data, such as proteomics datasets.

## Supporting information

Supp Benchmark

Supp Vignette

## References

Alvarez, M. J., Shen, Y., Giorgi, F. M., Lachmann, A., Ding, B. B., Ye, B. H., and Califano, A. (2016). Functional characterization of somatic mutations in cancer using network-based inference of protein activity. Nature genetics, 48(8), 838.

Castro, M. A., de Santiago, I., Campbell, T. M., Vaughn, C., Hickey, T. E., Ross, E., Tilley, W. D., Markowetz, F., Ponder, B. A., and Meyer, K. B. (2016). Regulators of genetic risk of breast cancer identified by integrative network analysis. Nature genetics, 48(1), 12.

Ding, H., Douglass, E. F., Sonabend, A. M., Mela, A., Bose, S., Gonzalez, C., Canoll, P. D., Sims, P. A., Alvarez, M. J., and Califano, A. (2018). Quantitative assessment of protein activity in orphan tissues and single cells using the metaviper algorithm. Nature communications, 9(1), 1471.

Giorgi, F. M., Del Fabbro, C., and Licausi, F. (2013). Comparative study of rna-seq-and microarray-derived coexpression networks in arabidopsis thaliana. Bioinformatics, 29(6), 717–724.

Kocak, H., Ackermann, S., Hero, B., Kahlert, Y., Oberthuer, A., Juraeva, D., Roels, F., Theissen, J., Westermann, F., Deubzer, H., et al. (2013). Hox-c9 activates the intrinsic pathway of apoptosis and is associated with spontaneous regression in neuroblastoma. Cell death & disease, 4(4), e586–e586.

Lachmann, A., Giorgi, F. M., Lopez, G., and Califano, A. (2016). ARACNe-AP: gene network reverse engineering through adaptive partitioning inference of mutual information. Bioinformatics, 32(14), 2233–2235.

Mercatelli, D., Scalambra, L., Triboli, L., Ray, F., and Giorgi, F. M. (2019a). Gene regulatory network inference resources: A practical overview. Biochimica et Biophysica Acta (BBA)-Gene Regulatory Mechanisms, page 194430.

Mercatelli, D., Ray, F., and Giorgi, F. M. (2019b). Pan-cancer and single-cell modelling of genomic alterations through gene expression. Frontiers in genetics, 10, 671.

Meyer, P. E., Lafitte, F., and Bontempi, G. (2008). minet: Ar/bioconductor package for inferring large transcriptional networks using mutual information. BMC bioinformatics, 9(1), 461.

Nicolle, R., Radvanyi, F., and Elati, M. (2015). Coregnet: reconstruction and integrated analysis of co-regulatory networks. Bioinformatics, 31(18), 3066–3068.

Reverter, A. and Chan, E. K. (2008). Combining partial correlation and an information theory approach to the reversed engineering of gene co-expression networks. Bioinformatics, 24(21), 2491–2497.

Schubert, M., Colomé-Tatché, M., and Foijer, F. (2019). Gene networks in cancer are biased by aneuploidies and sample impurities. Biochim Biophys Acta Gene Regul Mech, page 194444.

