## Supplementary material for "*corto*: a lightweight R package for Gene Network Inference and Master Regulator Analysis": Supp Benchmark

We tested *corto* on 39 tumor datasets, including the entire collection of TCGA (Weinstein et al., 2013), and on 27 normal tissue datasets from GTEx (Ardlie and Dermitzakis, 2015), to generate context-specific inferences of gene regulatory networks (**Supp Table S1**). We compared *corto* with ARACNe-AP (Lachmann et al., 2016), the other tool able to generate regulon objects for MRA. *corto*-generated networks are significantly similar to ARACNe-AP networks across all datasets (Supp Table S1), with edge intersections around 0.1 Jaccard Index (JI) All networks generated by ARACNe-AP and corto are provided on our website at https://giorgilab.org/corto-the-correlationtool/. Computationally, corto can generate networks significantly faster than ARACNe-AP, and using less RAM (**Supp Table S2**). In general, on tests performed with 100 and 1000 bootstraps corto is roughly 9 times faster and uses half the RAM of ARACNe-AP. *corto* can be efficiently run on an average desktop computer with context-specific datasets of hundreds of samples. In order to test if corto can infer realistic gene regulatory networks, we downloaded the entire ENCODE ChIP-Seq matrix, containing data for 187 transcription factors across 118 cell lines (Auerbach et al., 2013). For each ENCODE experiment we generated putative target genes, defined as genes whose Transcription Starting Site(s) fall within 250 nucleotides of the TF binding regions. For each ENCODE cell line, we histologically matched the closest gene expression dataset from TCGA or GTEx. We then calculated the accuracy of the overlap between gene regulatory networks inferred for selected TFs by ChIP-Seq, and the corresponding TF-centered networks from ARACNe-AP and *corto*. We measured that the intersection between ChIP-Seq and *corto*-inferred networks was significantly higher than expected and that *corto* is consistently more accurate than ARACNe-AP in all contexts, with accuracy consistently above 75% (**Supp Table S3**). Finally, we tested *corto* in MRA scenarios (see Figure below). In the example Figure, we stratified cancer samples according to the gene expression of the Epidermal Growth Factor Receptor (EGFR) in the TCGA Glioblastoma dataset. We then performed MRA between the samples with the highest level of EGFR (4th quartile) and the samples with the lowest level (1st quartile), using VIPER (Alvarez et al., 2016) with *corto* and ARACNe-AP networks. The results obtained with *corto* networks are almost identical to those obtained with ARACNe-AP, despite the JI of 0.1. Our example shows the activation of the TF SOX9 downstream of a EGFR overexpression in Glioblastoma, as previously validated (Liu et al., Molecular Cell 2015, PMID 26455392).

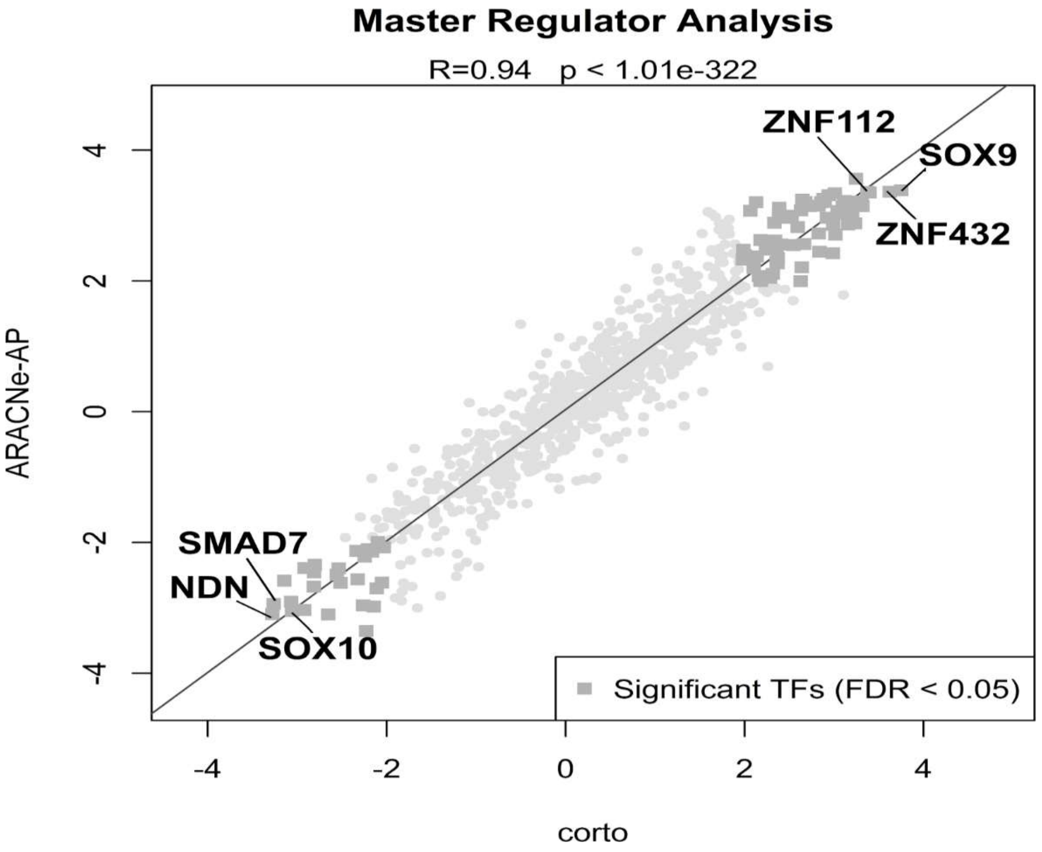


Correlation between the VIPER TF activities based on corto and ARACNe-AP networks. Significant TFs using both network inference tools are shown as darker gray squares. The 6 most significant up- and down-regulated TFs are shown as labels.

**References**

Alvarez, M. J., Shen, Y., Giorgi, F. M., Lachmann, A., Ding, B. B., Ye, B. H., and Califano, A. (2016). Functional characterization of somatic mutations in cancer using network-based inference of protein activity. Nature genetics, 48(8), 838.

Ardlie, K. G. and Dermitzakis, E. T. (2015). The genotype-tissue expression (gtex) pilot analysis: multitissue gene regulation in humans. Science, 348(6235), 648–660.

Auerbach, R. K., Chen, B., and Butte, A. J. (2013). Relating genes to function: identifying enriched transcription factors using the encode chip-seq significance tool. Bioinformatics, 29(15), 1922–1924.

Lachmann, A., Giorgi, F. M., Lopez, G., and Califano, A. (2016). Aracne-ap: gene network reverse engineering through adaptive partitioning inference of mutual information. Bioinformatics, 32(14), 2233–2235.

Liu, F., Hon, G. C., Villa, G. R., Turner, K. M., Ikegami, S., Li, B., Kuan, S., Lee, A. Y., et al. (2015). Egfr mutation promotes glioblastoma through epigenome and transcription factor network remodeling. Molecular cell, 60(2), 307–318.

Weinstein, J. N., Collisson, E. A., Mills, G. B., Shaw, K. R. M., Ozenberger, B. A., Ellrott, K., Shmulevich, I., Sander, C., Stuart, J. M., Network, C. G. A. R., et al. (2013). The cancer genome atlas pan-cancer analysis project. Nature genetics, 45(10), 1113.

***corto* : a lightweight R package to infer gene regulatory networks** by Mercatelli, Lopez-Garcia and Giorgi

**Supplementary Table S1**

Comparison between *corto* and ARACNe-AP networks

FET: Fisher's Exact Score. Calculated by R 3.6.1 (minimum measurable p-value: $10^{-302}$)

| Dataset | Nr.Samples | Nr.Nodes.Aracne | Nr.Nodes.Corto | Nr.Edges.Aracne | Nr.Edges.Corto | Nr.Edges.Intersection | Jaccard.Index | pvalue.FET | Type | Source |
|---|---|---|---|---|---|---|---|---|---|---|
| gtex_Adipose_Tissue | 797 | 21066 | 20650 | 1016897 | 170833 | 131061 | 0.124032218 | $<10^{-302}$ | RNA-Seq, VST-normalized (DESeq) | *https://gtexportal.org/home/* |
| gtex_Adrenal_Gland | 190 | 19969 | 16846 | 1681529 | 263912 | 204329 | 0.11735546 | $<10^{-302}$ | RNA-Seq, VST-normalized (DESeq) | *https://gtexportal.org/home/* |
| gtex_Blood | 537 | 20735 | 19623 | 635148 | 229991 | 91282 | 0.117957194 | $<10^{-302}$ | RNA-Seq, VST-normalized (DESeq) | *https://gtexportal.org/home/* |
| gtex_Blood_Vessel | 913 | 20972 | 20494 | 948392 | 157214 | 118906 | 0.120508767 | $<10^{-302}$ | RNA-Seq, VST-normalized (DESeq) | *https://gtexportal.org/home/* |
| gtex_Brain | 1671 | 21808 | 21965 | 493147 | 127859 | 66673 | 0.12027608 | $<10^{-302}$ | RNA-Seq, VST-normalized (DESeq) | *https://gtexportal.org/home/* |
| gtex_Breast | 290 | 20826 | 19841 | 1312433 | 275068 | 199046 | 0.143357905 | $<10^{-302}$ | RNA-Seq, VST-normalized (DESeq) | *https://gtexportal.org/home/* |
| gtex_Colon | 507 | 21056 | 20173 | 766051 | 229830 | 129206 | 0.149082413 | $<10^{-302}$ | RNA-Seq, VST-normalized (DESeq) | *https://gtexportal.org/home/* |
| gtex_Esophagus | 1021 | 21380 | 21136 | 645133 | 177173 | 90660 | 0.123912384 | $<10^{-302}$ | RNA-Seq, VST-normalized (DESeq) | *https://gtexportal.org/home/* |
| gtex_Heart | 600 | 20605 | 19670 | 1012172 | 181549 | 132503 | 0.12485936 | $<10^{-302}$ | RNA-Seq, VST-normalized (DESeq) | *https://gtexportal.org/home/* |
| gtex_Kidney | 45 | 19282 | 8843 | 990727 | 45734 | 44491 | 0.044851155 | $<10^{-302}$ | RNA-Seq, VST-normalized (DESeq) | *https://gtexportal.org/home/* |
| gtex_Liver | 175 | 19654 | 17202 | 1621251 | 255744 | 198735 | 0.118417289 | $<10^{-302}$ | RNA-Seq, VST-normalized (DESeq) | *https://gtexportal.org/home/* |
| gtex_Lung | 427 | 20870 | 19348 | 1179775 | 208211 | 162637 | 0.132727084 | $<10^{-302}$ | RNA-Seq, VST-normalized (DESeq) | *https://gtexportal.org/home/* |
| gtex_Muscle | 564 | 20441 | 20554 | 1321222 | 209506 | 156752 | 0.114086418 | $<10^{-302}$ | RNA-Seq, VST-normalized (DESeq) | *https://gtexportal.org/home/* |
| gtex_Nerve | 414 | 20822 | 19872 | 1351331 | 222306 | 174887 | 0.12503092 | $<10^{-302}$ | RNA-Seq, VST-normalized (DESeq) | *https://gtexportal.org/home/* |
| gtex_Ovary | 133 | 19852 | 16442 | 1709431 | 240587 | 201821 | 0.115445227 | $<10^{-302}$ | RNA-Seq, VST-normalized (DESeq) | *https://gtexportal.org/home/* |
| gtex_Pancreas | 248 | 19932 | 16755 | 1778586 | 225516 | 194610 | 0.107549522 | $<10^{-302}$ | RNA-Seq, VST-normalized (DESeq) | *https://gtexportal.org/home/* |
| gtex_Pituitary | 183 | 20584 | 18104 | 1605400 | 293898 | 220364 | 0.13125233 | $<10^{-302}$ | RNA-Seq, VST-normalized (DESeq) | *https://gtexportal.org/home/* |
| gtex_Prostate | 152 | 20370 | 17776 | 1682421 | 282550 | 230446 | 0.132858275 | $<10^{-302}$ | RNA-Seq, VST-normalized (DESeq) | *https://gtexportal.org/home/* |
| gtex_Salivary_Gland | 97 | 19985 | 15846 | 1661204 | 242401 | 183161 | 0.106461472 | $<10^{-302}$ | RNA-Seq, VST-normalized (DESeq) | *https://gtexportal.org/home/* |
| gtex_Skin | 1203 | 21604 | 21822 | 605408 | 166183 | 73714 | 0.105626063 | $<10^{-302}$ | RNA-Seq, VST-normalized (DESeq) | *https://gtexportal.org/home/* |
| gtex_Small_Intestine | 137 | 20536 | 18120 | 1262939 | 342451 | 214552 | 0.154260956 | $<10^{-302}$ | RNA-Seq, VST-normalized (DESeq) | *https://gtexportal.org/home/* |
| gtex_Spleen | 162 | 20084 | 16255 | 1775758 | 217518 | 188377 | 0.104369829 | $<10^{-302}$ | RNA-Seq, VST-normalized (DESeq) | *https://gtexportal.org/home/* |
| gtex_Stomach | 262 | 20514 | 18643 | 1118816 | 308233 | 185117 | 0.149055665 | $<10^{-302}$ | RNA-Seq, VST-normalized (DESeq) | *https://gtexportal.org/home/* |
| gtex_Testis | 259 | 22367 | 20476 | 1156684 | 328515 | 187059 | 0.144097709 | $<10^{-302}$ | RNA-Seq, VST-normalized (DESeq) | *https://gtexportal.org/home/* |
| gtex_Thyroid | 446 | 20850 | 19758 | 1435642 | 217928 | 181870 | 0.123578175 | $<10^{-302}$ | RNA-Seq, VST-normalized (DESeq) | *https://gtexportal.org/home/* |
| gtex_Uterus | 111 | 19737 | 15318 | 1764353 | 216298 | 183945 | 0.10237902 | $<10^{-302}$ | RNA-Seq, VST-normalized (DESeq) | *https://gtexportal.org/home/* |
| gtex_Vagina | 115 | 20031 | 16765 | 1507841 | 313854 | 215914 | 0.134460428 | $<10^{-302}$ | RNA-Seq, VST-normalized (DESeq) | *https://gtexportal.org/home/* |
| kocak_NBL | 649 | 18849 | 18759 | 1047641 | 182259 | 146554 | 0.135279034 | $<10^{-302}$ | Microarrays, RMA-normalized | *https://www.ncbi.nlm.nih.gov/geo/query/acc.cgi?acc=GSE45547* |
| metabric_BRCA | 996 | 17589 | 12901 | 1536258 | 99408 | 84469 | 0.054454076 | $<10^{-302}$ | RNA-Seq, VST-normalized (DESeq) | *http://www.cbioportal.org/datasets (Discovery dataset)* |
| nrc_NBL | 283 | 17324 | 16618 | 1062930 | 273568 | 187798 | 0.163487421 | $<10^{-302}$ | Microarrays, RMA-normalized | *https://www.ncbi.nlm.nih.gov/geo/query/acc.cgi?acc=GSE85047* |
| target_NBL | 417 | 20588 | 20586 | 1010004 | 261634 | 142263 | 0.125966132 | $<10^{-302}$ | RNA-Seq + microarrays, TDM-normalized | *https://ocg.cancer.gov/programs/target/projects/neuroblastoma* |
| target_NBLrnaseq | 168 | 24384 | 21617 | 1999921 | 334680 | 249267 | 0.11953337 | $<10^{-302}$ | RNA-Seq, VST-normalized (DESeq) | *https://ocg.cancer.gov/programs/target/projects/neuroblastoma* |
| tcga_ACC | 79 | 17588 | 8820 | 1462660 | 32367 | 27505 | 0.018742479 | $<10^{-302}$ | RNA-Seq, VST-normalized (DESeq) | *https://gdac.broadinstitute.org/* |
| tcga_BLCA | 427 | 18912 | 19049 | 1404927 | 247732 | 176970 | 0.119923642 | $<10^{-302}$ | RNA-Seq, VST-normalized (DESeq) | *https://gdac.broadinstitute.org/* |
| tcga_BRCA | 1212 | 19243 | 19714 | 798603 | 162262 | 107453 | 0.125909877 | $<10^{-302}$ | RNA-Seq, VST-normalized (DESeq) | *https://gdac.broadinstitute.org/* |
| tcga_CESC | 309 | 18716 | 18277 | 1596515 | 211296 | 157233 | 0.095259358 | $<10^{-302}$ | RNA-Seq, VST-normalized (DESeq) | *https://gdac.broadinstitute.org/* |
| tcga_CHOL | 45 | 17530 | 7361 | 1025393 | 43668 | 38240 | 0.037096644 | $<10^{-302}$ | RNA-Seq, VST-normalized (DESeq) | *https://gdac.broadinstitute.org/* |
| tcga_COAD | 328 | 18585 | 18358 | 1342130 | 242348 | 174537 | 0.123790286 | $<10^{-302}$ | RNA-Seq, VST-normalized (DESeq) | *https://gdac.broadinstitute.org/* |
| tcga_DLBC | 48 | 17009 | 7385 | 875156 | 81823 | 71597 | 0.08086566 | $<10^{-302}$ | RNA-Seq, VST-normalized (DESeq) | *https://gdac.broadinstitute.org/* |
| tcga_ESCA | 196 | 19289 | 18022 | 1586364 | 212379 | 150967 | 0.091618642 | $<10^{-302}$ | RNA-Seq, VST-normalized (DESeq) | *https://gdac.broadinstitute.org/* |
| tcga_GBM | 171 | 18404 | 17027 | 1658235 | 236792 | 192030 | 0.112760034 | $<10^{-302}$ | RNA-Seq, VST-normalized (DESeq) | *https://gdac.broadinstitute.org/* |
| tcga_HNSC | 566 | 19244 | 19555 | 1288606 | 219272 | 165886 | 0.123611765 | $<10^{-302}$ | RNA-Seq, VST-normalized (DESeq) | *https://gdac.broadinstitute.org/* |
| tcga_KICH | 91 | 18260 | 15559 | 1580958 | 248630 | 181975 | 0.11044766 | $<10^{-302}$ | RNA-Seq, VST-normalized (DESeq) | *https://gdac.broadinstitute.org/* |
| tcga_KIRC | 606 | 19255 | 19741 | 1057848 | 227957 | 156918 | 0.139002398 | $<10^{-302}$ | RNA-Seq, VST-normalized (DESeq) | *https://gdac.broadinstitute.org/* |
| tcga_KIRP | 323 | 18760 | 19241 | 1379865 | 284284 | 200480 | 0.136970859 | $<10^{-302}$ | RNA-Seq, VST-normalized (DESeq) | *https://gdac.broadinstitute.org/* |
| tcga_LAML | 173 | 17773 | 17038 | 1569326 | 231900 | 182259 | 0.112577341 | $<10^{-302}$ | RNA-Seq, VST-normalized (DESeq) | *https://gdac.broadinstitute.org/* |
| tcga_LGG | 530 | 18987 | 18890 | 1113177 | 211954 | 157796 | 0.135176278 | $<10^{-302}$ | RNA-Seq, VST-normalized (DESeq) | *https://gdac.broadinstitute.org/* |

| | | | | | | | | | | |
|---|---|---|---|---|---|---|---|---|---|---|
| tcga_LIHC | 423 | 18638 | 18978 | 1340322 | 254331 | 180016 | 0.127252433 | $<10^{-302}$ | RNA-Seq, VST-normalized (DESeq) | *https://gdac.broadinstitute.org/* |
| tcga_LUAD | 576 | 19011 | 19212 | 1275070 | 218441 | 161570 | 0.121304172 | $<10^{-302}$ | RNA-Seq, VST-normalized (DESeq) | *https://gdac.broadinstitute.org/* |
| tcga_LUSC | 552 | 19215 | 19640 | 1177957 | 225438 | 155412 | 0.124530542 | $<10^{-302}$ | RNA-Seq, VST-normalized (DESeq) | *https://gdac.broadinstitute.org/* |
| tcga_MESO | 87 | 18060 | 11054 | 1498092 | 71690 | 57275 | 0.037867593 | $<10^{-302}$ | RNA-Seq, VST-normalized (DESeq) | *https://gdac.broadinstitute.org/* |
| tcga_OV | 307 | 18967 | 17609 | 1699890 | 151024 | 123273 | 0.071353366 | $<10^{-302}$ | RNA-Seq, VST-normalized (DESeq) | *https://gdac.broadinstitute.org/* |
| tcga_PAAD | 183 | 18612 | 18028 | 1534310 | 298000 | 207845 | 0.127946739 | $<10^{-302}$ | RNA-Seq, VST-normalized (DESeq) | *https://gdac.broadinstitute.org/* |
| tcga_PCPG | 187 | 18268 | 17285 | 1572415 | 234337 | 171005 | 0.104542451 | $<10^{-302}$ | RNA-Seq, VST-normalized (DESeq) | *https://gdac.broadinstitute.org/* |
| tcga_PRAD | 550 | 18982 | 19005 | 975298 | 218124 | 153319 | 0.147407516 | $<10^{-302}$ | RNA-Seq, VST-normalized (DESeq) | *https://gdac.broadinstitute.org/* |
| tcga_READ | 105 | 18125 | 13812 | 1612057 | 128961 | 100152 | 0.061036063 | $<10^{-302}$ | RNA-Seq, VST-normalized (DESeq) | *https://gdac.broadinstitute.org/* |
| tcga_SARC | 265 | 18711 | 18099 | 1517489 | 238774 | 167355 | 0.105327055 | $<10^{-302}$ | RNA-Seq, VST-normalized (DESeq) | *https://gdac.broadinstitute.org/* |
| tcga_SKCM | 473 | 18965 | 19042 | 1292735 | 219030 | 156295 | 0.115306868 | $<10^{-302}$ | RNA-Seq, VST-normalized (DESeq) | *https://gdac.broadinstitute.org/* |
| tcga_STAD | 450 | 19844 | 19898 | 1328546 | 272982 | 194215 | 0.138004126 | $<10^{-302}$ | RNA-Seq, VST-normalized (DESeq) | *https://gdac.broadinstitute.org/* |
| tcga_TGCT | 156 | 19300 | 18839 | 1265423 | 362158 | 233862 | 0.167797095 | $<10^{-302}$ | RNA-Seq, VST-normalized (DESeq) | *https://gdac.broadinstitute.org/* |
| tcga_THCA | 568 | 18797 | 19114 | 951870 | 203952 | 150385 | 0.149571778 | $<10^{-302}$ | RNA-Seq, VST-normalized (DESeq) | *https://gdac.broadinstitute.org/* |
| tcga_THYM | 122 | 18581 | 17811 | 1287751 | 353430 | 231165 | 0.163944948 | $<10^{-302}$ | RNA-Seq, VST-normalized (DESeq) | *https://gdac.broadinstitute.org/* |
| tcga_UCEC | 201 | 18726 | 17333 | 1524483 | 249891 | 171064 | 0.106694276 | $<10^{-302}$ | RNA-Seq, VST-normalized (DESeq) | *https://gdac.broadinstitute.org/* |
| tcga_UCS | 57 | 18320 | 4499 | 942251 | 12304 | 11422 | 0.012110699 | $<10^{-302}$ | RNA-Seq, VST-normalized (DESeq) | *https://gdac.broadinstitute.org/* |
| tcga_UVM | 80 | 17353 | 14306 | 1324565 | 265504 | 183218 | 0.130232697 | $<10^{-302}$ | RNA-Seq, VST-normalized (DESeq) | *https://gdac.broadinstitute.org/* |
| versteeg_NBL | 118 | 20454 | 12118 | 1923812 | 108650 | 84719 | 0.043495985 | $<10^{-302}$ | Microarrays, RMA-normalized | *https://www.ncbi.nlm.nih.gov/geo/query/acc.cgi?acc=GSE16476* |

***corto*** **: a lightweight R package to infer gene regulatory networks** by Mercatelli, Lopez-Garcia and Giorgi

**Supplementary Table S2A**

Time and memory benchmarks for network generation

| ***corto*** benchmark | | **nbootstraps = 100, nthreads = 4, p-value = 1e-8** | | **nbootstraps = 1000, nthreads = 4, p-value = 1e-8** | |
|---|---|---|---|---|---|
| **TCGA dataset** | **Nr. Samples** | **Elapsed Time (minutes)** | **Peak RAM Used (MiB)** | **Elapsed Time (minutes)** | **Peak RAM Used (MiB)** |
| ACC | 79 | 1.6 | 464.8 | 14.91 | 464.9 |
| BLCA | 427 | 23.61 | 1026 | 222.17 | 1025.9 |
| BRCA | 1212 | 77.96 | 2342.1 | 735.23 | 1912.2 |
| CESC | 309 | 12.72 | 1183.3 | 117.37 | 960.8 |
| CHOL | 45 | 1.36 | 575.5 | 12.59 | 449.8 |
| COAD | 328 | 25.16 | 802.9 | 235.57 | 995.9 |
| DLBC | 48 | 1.9 | 567.8 | 17.56 | 478.2 |
| ESCA | 196 | 9.24 | 565.9 | 87.08 | 706.5 |
| GBM | 171 | 10.22 | 537.1 | 98.1 | 671.9 |
| HNSC | 566 | 37.5 | 1198.6 | 350.57 | 1202.1 |
| KICH | 91 | 9.23 | 596.7 | 84 | 596.1 |
| KIRC | 606 | 57.81 | 1469.7 | 521.18 | 1553.2 |
| KIRP | 323 | 29.03 | 915.8 | 272.57 | 925 |
| LAML | 173 | 10.25 | 634.5 | 94.58 | 787.9 |
| **LGG** | **530** | **52.47** | **1453.4** | 480.37 | 1453.4 |
| LIHC | 423 | 27 | 860.5 | 253.06 | 1040.9 |
| LUAD | 576 | 32.59 | 973.6 | 297.6 | 973.7 |
| LUSC | 552 | 35.16 | 1188.4 | 325.4 | 1188.4 |
| OV | 307 | 8.12 | 729 | 76.49 | 729.1 |
| PAAD | 183 | 15.16 | 682.7 | 144.64 | 682.8 |
| PCPG | 187 | 11.06 | 547.8 | 104.38 | 684.4 |
| PRAD | 550 | 55.17 | 1378.8 | 518.73 | 1608.6 |
| READ | 105 | 4.53 | 683.2 | 42.6 | 623.9 |
| SARC | 265 | 14.67 | 747.6 | 137.52 | 747.5 |
| SKCM | 473 | 31.13 | 1115.8 | 290 | 1115.7 |
| STAD | 450 | 30.77 | 1097.8 | 289.58 | 1097.8 |
| TGCT | 156 | 29.09 | 729.3 | 281.12 | 901.2 |
| THCA | 568 | 60.76 | 1489.3 | 568.87 | 1477.2 |

| | | | | | |
|---|---|---|---|---|---|
| THYM | 122 | 24.27 | 738.5 | 229.09 | 732.4 |
| UCEC | 201 | 13.38 | 698.4 | 130.78 | 865.6 |
| UCS | 57 | 0.8 | 470.2 | 7.37 | 568.7 |
| UVM | 80 | 11.91 | 471.6 | 114.33 | 589.5 |

peakRAM package 1.0.2
R version 3.6.1
running under Microsoft Windows 10 Pro (build 18362)
Intel(R) Core(TM) i5-8400 CPU @ 2.80GHz, 2808 Mhz, 6 cores
RAM 32 GB

***corto* : a lightweight R package to infer gene regulatory networks** by Mercatelli, Lopez-Garcia and Giorgi

**Supplementary Table S3**

Comparison between *corto* and ARACNe-AP networks with ENCODE ChIP-Seq inferred regulatory networks

| Cell Line | RNASeq dataset | TFs in both | ChIPSeq edges | Aracne Edges | Corto Edges | TP Aracne | TP Corto | FP Aracne | FP Corto | TN Aracne | TN Corto | FN Aracne | FN Corto | Aracne Accuracy | Corto Accuracy |
|---|---|---|---|---|---|---|---|---|---|---|---|---|---|---|---|
| A549 | tcga_LUAD | 20 | 67177 | 25013 | 5260 | 6030 | 1304 | 18983 | 3956 | 386153 | 401180 | 60154 | 64880 | 83.21% | 85.40% |
| BE2_C | kocak_NBL | 1 | 3348 | 939 | 145 | 193 | 35 | 746 | 110 | 19956 | 20592 | 3102 | 3260 | 83.97% | 85.96% |
| Caco-2 | tcga_COAD | 1 | 3762 | 1312 | 79 | 296 | 25 | 1016 | 54 | 18922 | 19884 | 3351 | 3622 | 81.48% | 84.41% |
| ECC-1 | tcga_UCEC | 4 | 3470 | 6266 | 1074 | 300 | 52 | 5966 | 1022 | 84922 | 89866 | 3140 | 3388 | 90.35% | 95.33% |
| Gliobla | tcga_GBM | 1 | 4666 | 1171 | 413 | 313 | 113 | 858 | 300 | 18238 | 18796 | 4176 | 4376 | 78.66% | 80.17% |
| HCT-116 | tcga_COAD | 3 | 14889 | 4834 | 692 | 1530 | 256 | 3304 | 436 | 52785 | 55653 | 13130 | 14404 | 76.77% | 79.02% |
| HEK293 | tcga_KIRP | 4 | 10214 | 3438 | 635 | 552 | 79 | 2886 | 556 | 81331 | 83661 | 9559 | 10032 | 86.81% | 88.78% |
| HEK293-T-REx | tcga_KIRP | 1 | 5208 | 1350 | 134 | 350 | 30 | 1000 | 104 | 17421 | 18317 | 4814 | 5134 | 75.35% | 77.79% |
| HeLa-S3 | tcga_UCEC | 35 | 107985 | 52303 | 13246 | 10429 | 3204 | 41874 | 10042 | 675941 | 707773 | 96041 | 103266 | 83.27% | 86.25% |
| HepG2 | tcga_LIHC | 45 | 130484 | 60239 | 17337 | 11759 | 4164 | 48480 | 13173 | 882694 | 918001 | 116412 | 124007 | 84.44% | 87.05% |
| MCF-7 | tcga_BRCA | 6 | 26742 | 7177 | 2129 | 1593 | 377 | 5584 | 1752 | 110244 | 114076 | 24059 | 25275 | 79.05% | 80.90% |
| NB4 | tcga_LAML | 3 | 17668 | 5210 | 385 | 1775 | 158 | 3435 | 227 | 50188 | 53396 | 15351 | 16968 | 73.45% | 75.70% |
| PANC-1 | tcga_PAAD | 2 | 4004 | 4297 | 2279 | 443 | 217 | 3854 | 2062 | 39330 | 41122 | 3541 | 3767 | 84.32% | 87.64% |
| T-47D | tcga_BRCA | 5 | 8994 | 9796 | 3602 | 982 | 273 | 8814 | 3329 | 100215 | 105700 | 7894 | 8603 | 85.83% | 89.88% |
| U87 | tcga_GBM | 1 | 4777 | 1277 | 421 | 390 | 149 | 887 | 272 | 17976 | 18591 | 4332 | 4573 | 77.87% | 79.46% |
