## Supplementary material for "*corto*: a lightweight R package for Gene Network Inference and Master Regulator Analysis": Supp File S1

**corto: a lightweight R package to infer gene regulatory networks** by Mercatelli, Lopez-Garcia and Giorgi

**Supplementary File S1**

This document contains four graphs for each TCGA dataset type, indicated as their official TCGA acronym (https://gdc.cancer.gov/resources-tcga-users/tcga-code-tables/tcga-study-abbreviations). Briefly, in the Master Regulator Analysis, the correlation between activity scores (expressed as Normalized Enrichment Scores of all TFs on a EGFR high vs. low signature) is shown. Top 3 most significant up- and down-regulated TFs are highlighted. A second graph, showing how patients were subset for high vs. low EGFR expression, is included. Vertical dotted grey lines corresponding to 1$^{st}$ and 3$^{rd}$ quartile are depicted. Horizontal dashed lines correspond to the 1$^{st}$ and 3$^{rd}$ EGFR expression quartile. Patients below the 1$^{st}$ quartile are taken as low EGFR expressing patients, while patients above the 3$^{rd}$ quartile are taken as high EGFR expressing patients for Master Regulator Analysis. The last two graphs represent the msVIPER tables obtained by using the ARACNe-AP or the corto regulon, respectively. Induced and repressed target genes, according to the regulatory model for the corresponding regulator (Set column), are depicted by blue and red vertical lines, respectively. General activity (i.e. repression or activation) is shown by a blue or red box in the “Act” column, while the “Exp” column reflects the differential expression of the corresponding regulator in the EGFR high vs. low comparison.

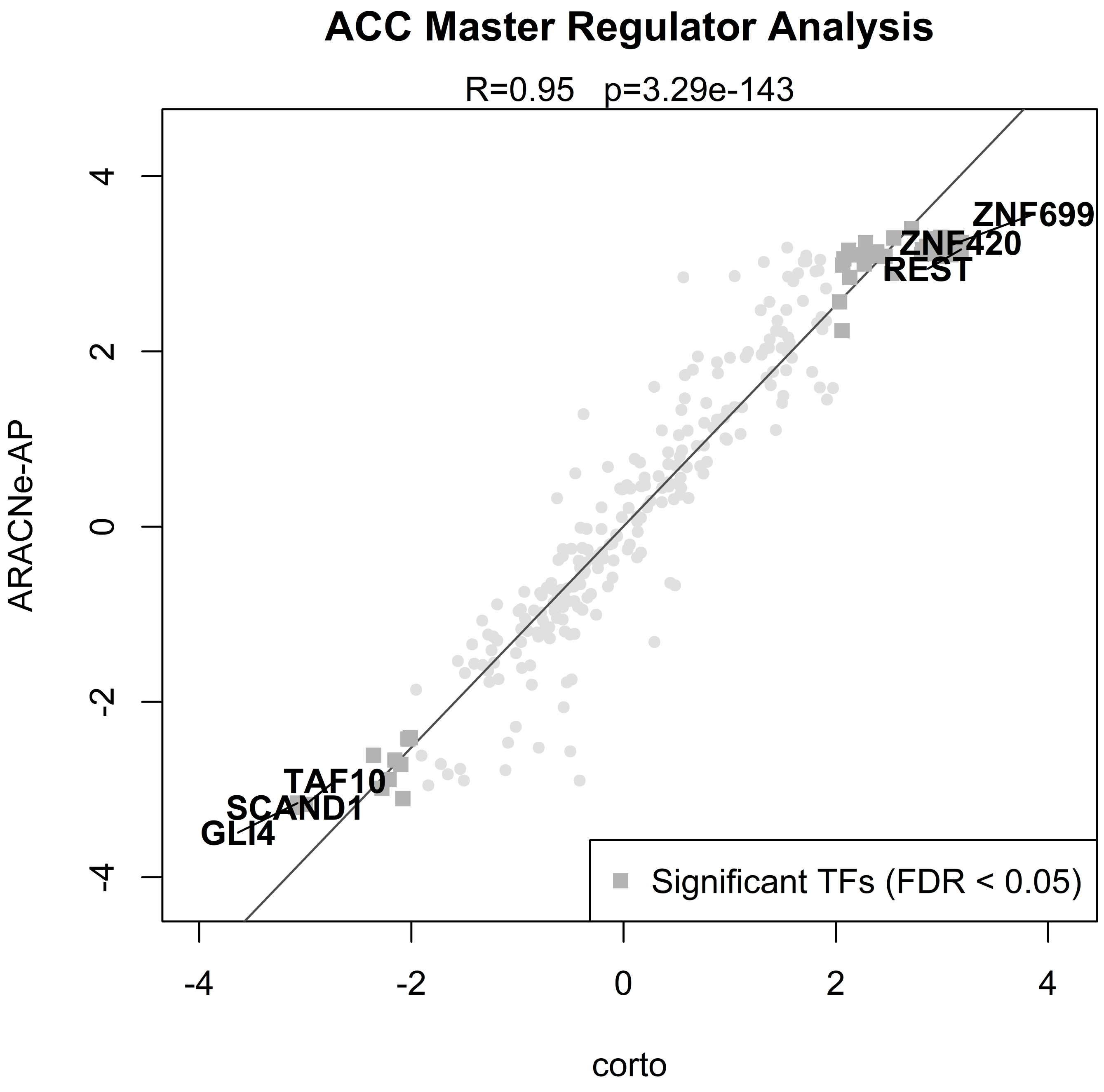

ACC Master Regulator Analysis
R=0.95 p=3.29e-143
ARACNe-AP
corto
-4
-2
0
2
4
ZNF699
ZNF420
REST
TAF10
SCAND1
GLI4
Significant TFs (FDR < 0.05)

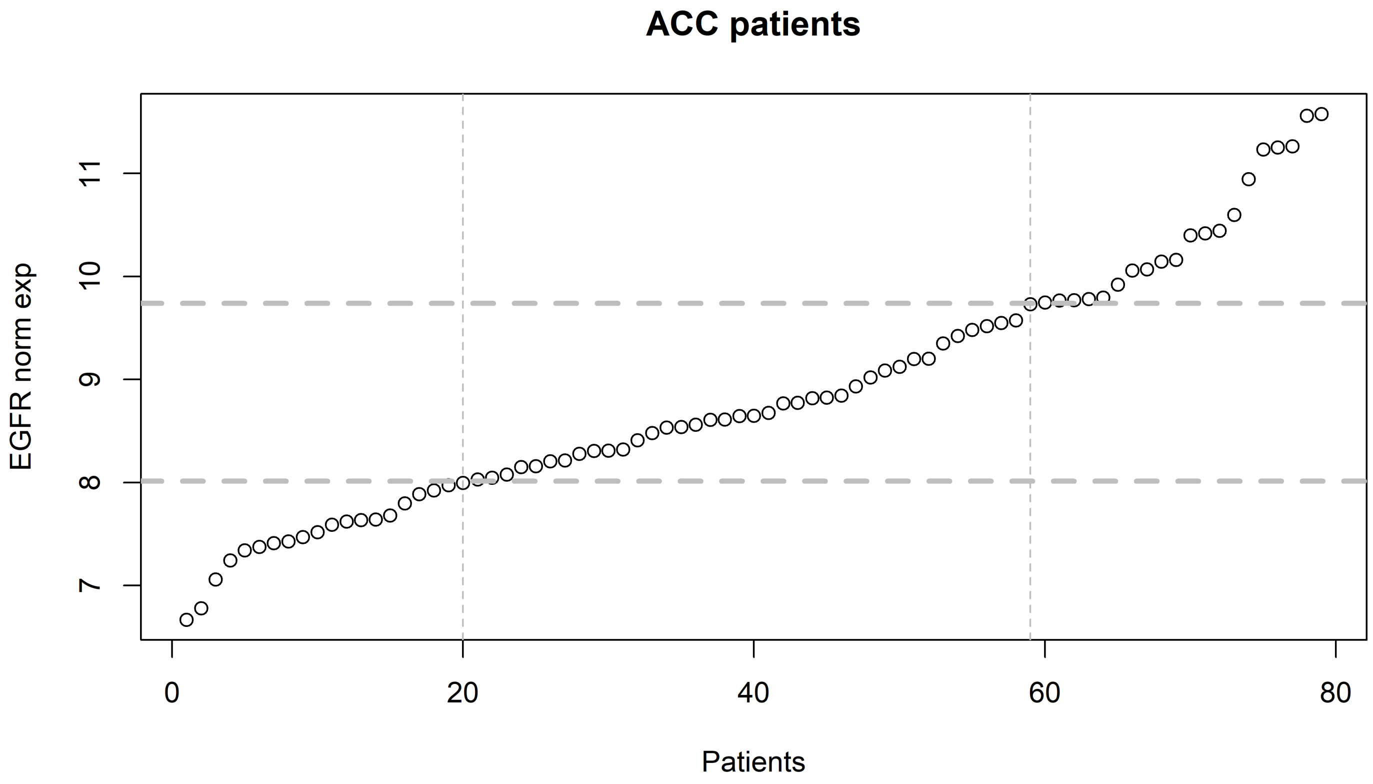
ACC patients
EGFR norm exp
7
8
9
10
11
0
20
40
60
80
Patients

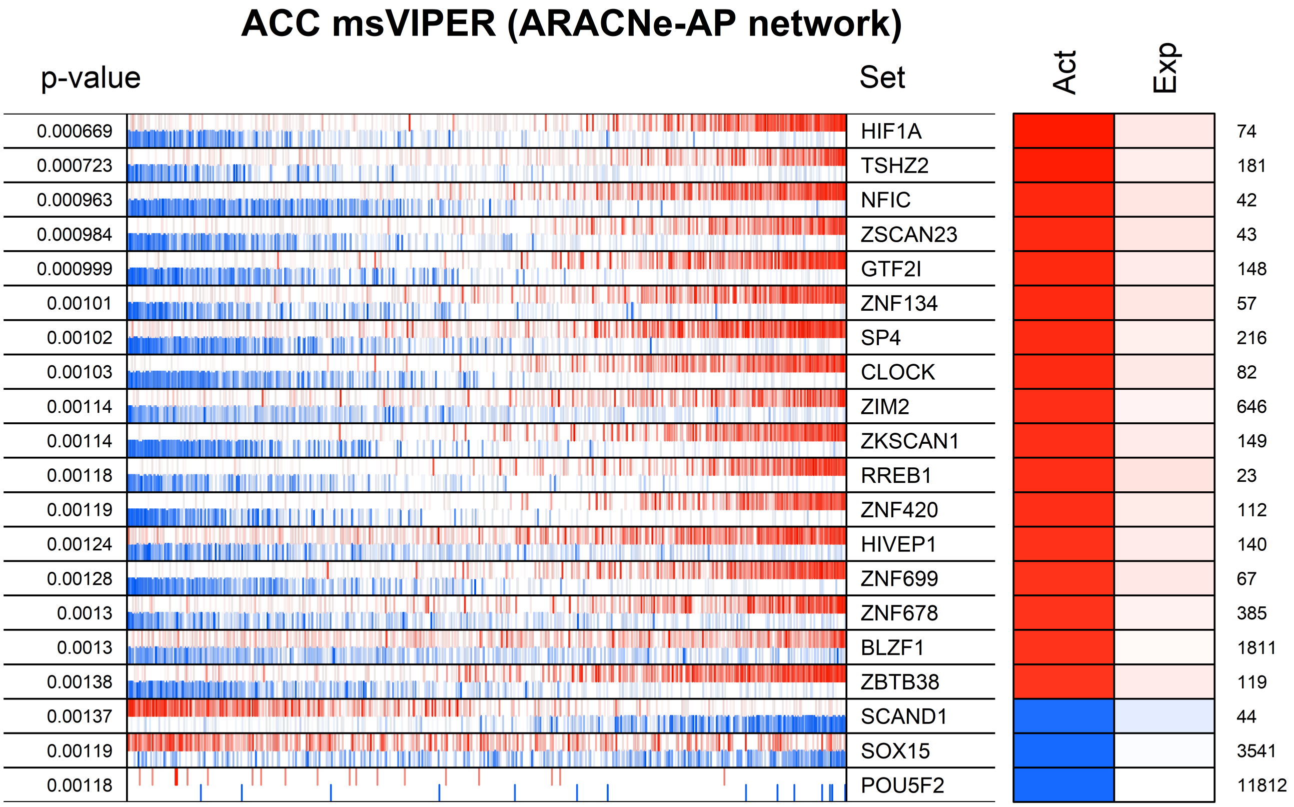
ACC msVIPER (ARACNe-AP network)
p-value
Set
Act
Exp
0.000669 HIF1A 74
0.000723 TSHZ2 181
0.000963 NFIC 42
0.000984 ZSCAN23 43
0.000999 GTF2I 148
0.00101 ZNF134 57
0.00102 SP4 216
0.00103 CLOCK 82
0.00114 ZIM2 646
0.00114 ZKSCAN1 149
0.00118 RREB1 23
0.00119 ZNF420 112
0.00124 HIVEP1 140
0.00128 ZNF699 67
0.0013 ZNF678 385
0.0013 BLZF1 1811
0.00138 ZBTB38 119
0.00137 SCAND1 44
0.00119 SOX15 3541
0.00118 POU5F2 11812

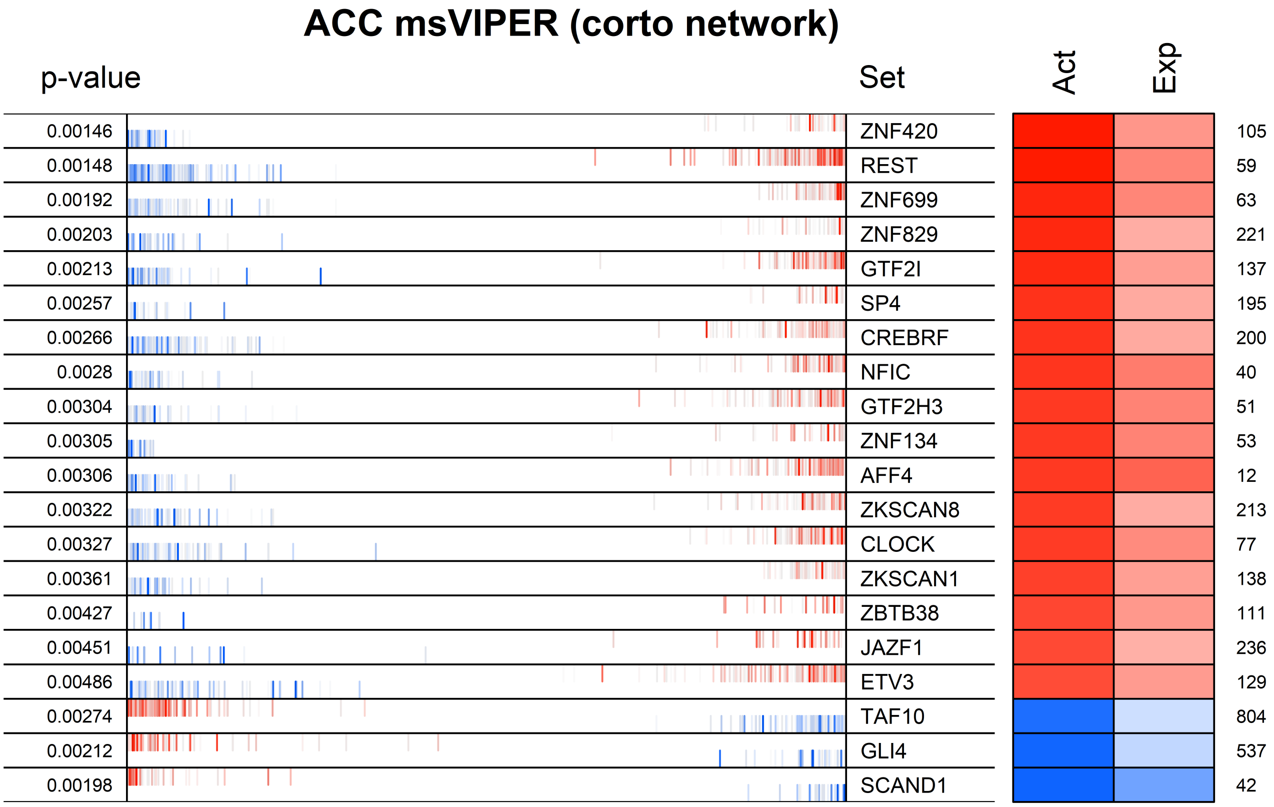
ACC msVIPER (corto network)
p-value
Set
Act
Exp
0.00146
ZNF420
105
0.00148
REST
59
0.00192
ZNF699
63
0.00203
ZNF829
221
0.00213
GTF2I
137
0.00257
SP4
195
0.00266
CREBRF
200
0.0028
NFIC
40
0.00304
GTF2H3
51
0.00305
ZNF134
53
0.00306
AFF4
12
0.00322
ZKSCAN8
213
0.00327
CLOCK
77
0.00361
ZKSCAN1
138
0.00427
ZBTB38
111
0.00451
JAZF1
236
0.00486
ETV3
129
0.00274
TAF10
804
0.00212
GLI4
537
0.00198
SCAND1
42

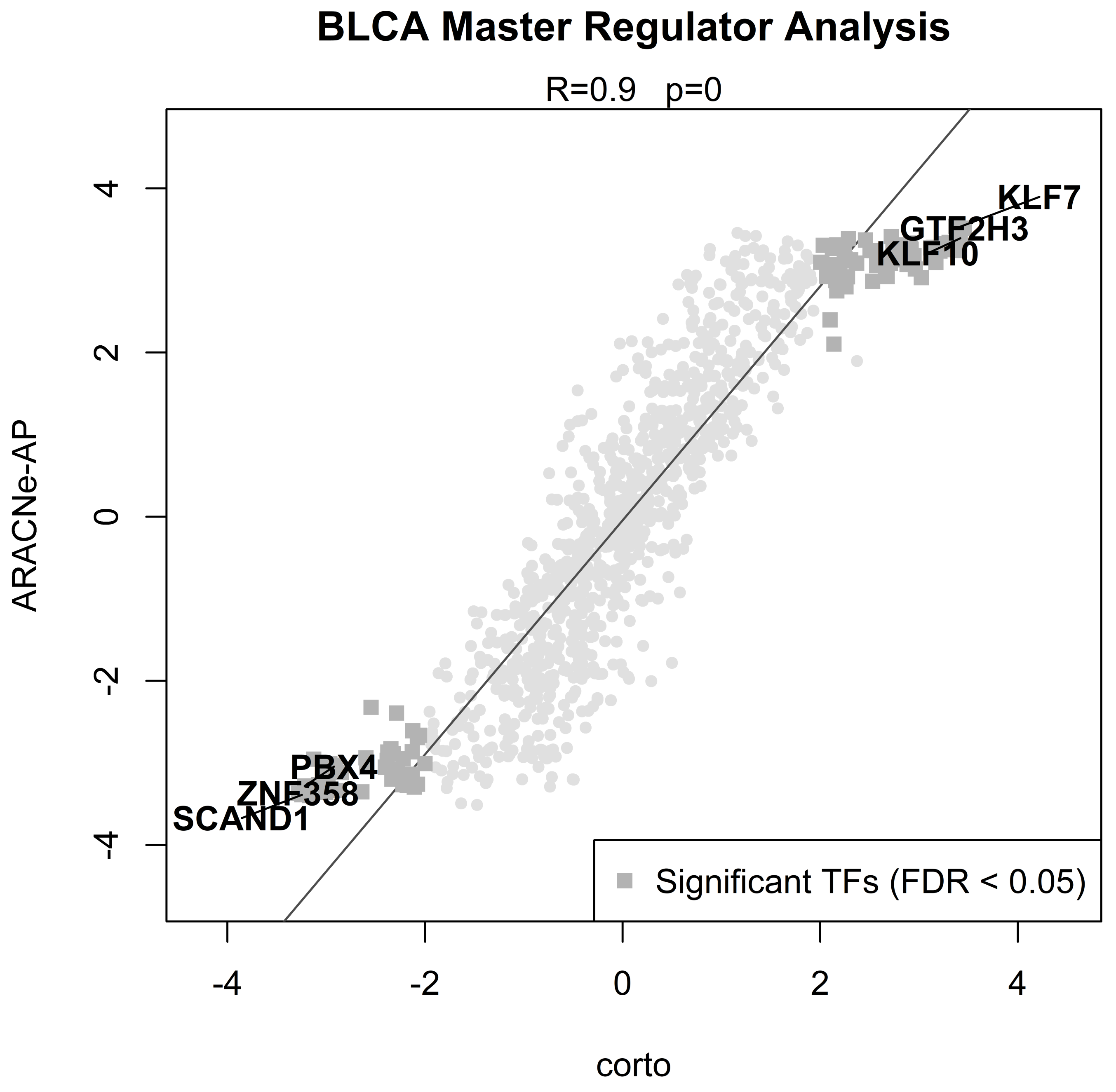
BLCA Master Regulator Analysis
R=0.9 p=0
KLF7
GTF2H3
KLF10
PBX4
ZNF358
SCAND1
Significant TFs (FDR < 0.05)
ARACNe-AP
corto
4
2
0
-2
-4
-4
-2
0
2
4

### BLCA patients

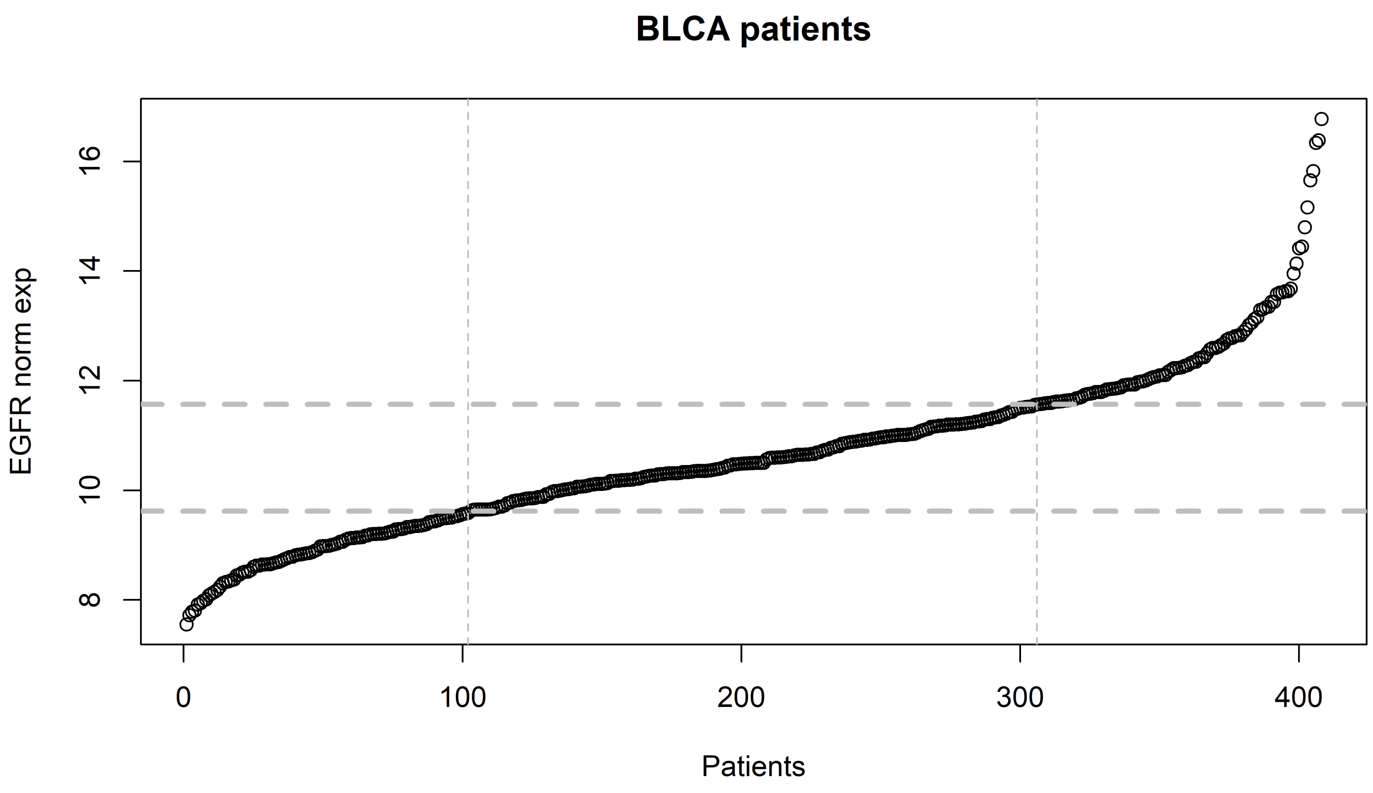

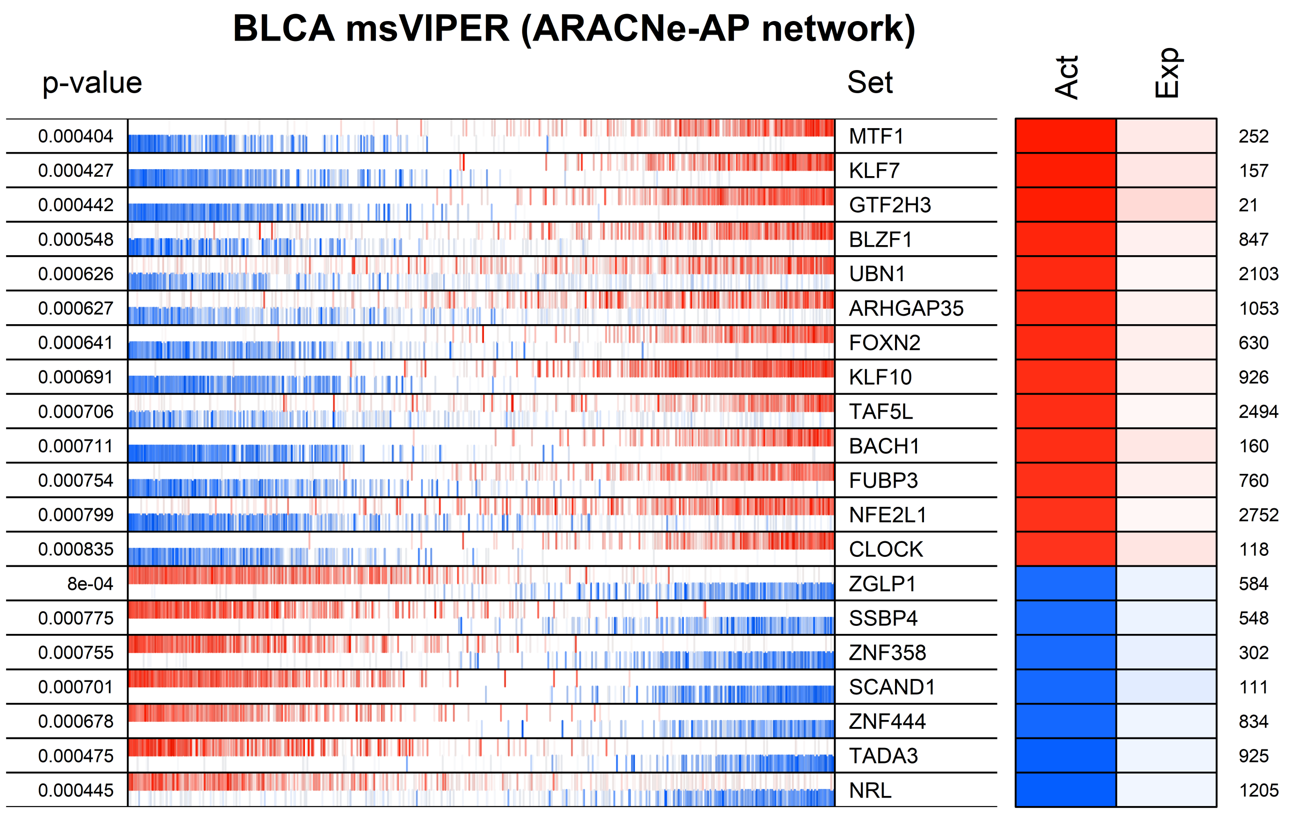
BLCA msVIPER (ARACNe-AP network)
p-value
Set
Act
Exp
0.000404
0.000427
0.000442
0.000548
0.000626
0.000627
0.000641
0.000691
0.000706
0.000711
0.000754
0.000799
0.000835
8e-04
0.000775
0.000755
0.000701
0.000678
0.000475
0.000445
MTF1
KLF7
GTF2H3
BLZF1
UBN1
ARHGAP35
FOXN2
KLF10
TAF5L
BACH1
FUBP3
NFE2L1
CLOCK
ZGLP1
SSBP4
ZNF358
SCAND1
ZNF444
TADA3
NRL
252
157
21
847
2103
1053
630
926
2494
160
760
2752
118
584
548
302
111
834
925
1205

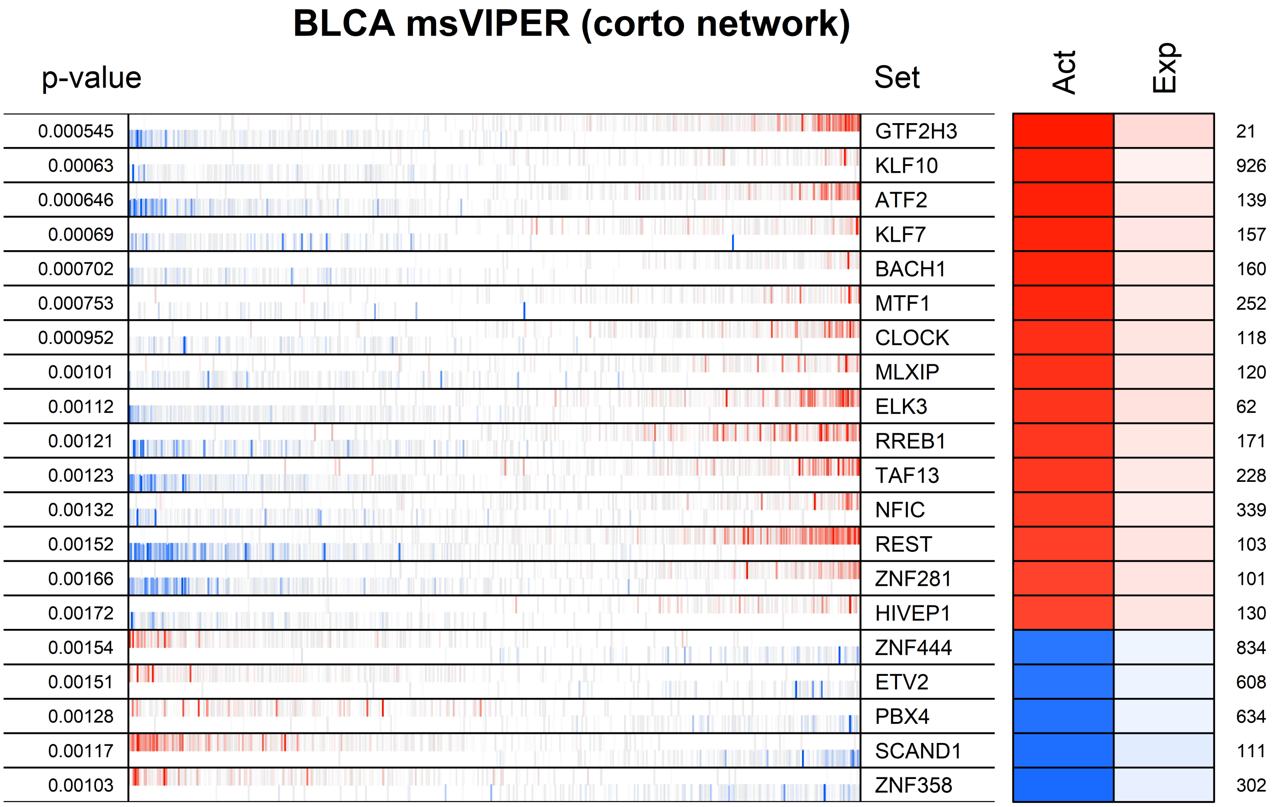
BLCA msVIPER (corto network)
p-value
Set
Act
Exp
0.000545 GTF2H3 21
0.00063 KLF10 926
0.000646 ATF2 139
0.00069 KLF7 157
0.000702 BACH1 160
0.000753 MTF1 252
0.000952 CLOCK 118
0.00101 MLXIP 120
0.00112 ELK3 62
0.00121 RREB1 171
0.00123 TAF13 228
0.00132 NFIC 339
0.00152 REST 103
0.00166 ZNF281 101
0.00172 HIVEP1 130
0.00154 ZNF444 834
0.00151 ETV2 608
0.00128 PBX4 634
0.00117 SCAND1 111
0.00103 ZNF358 302

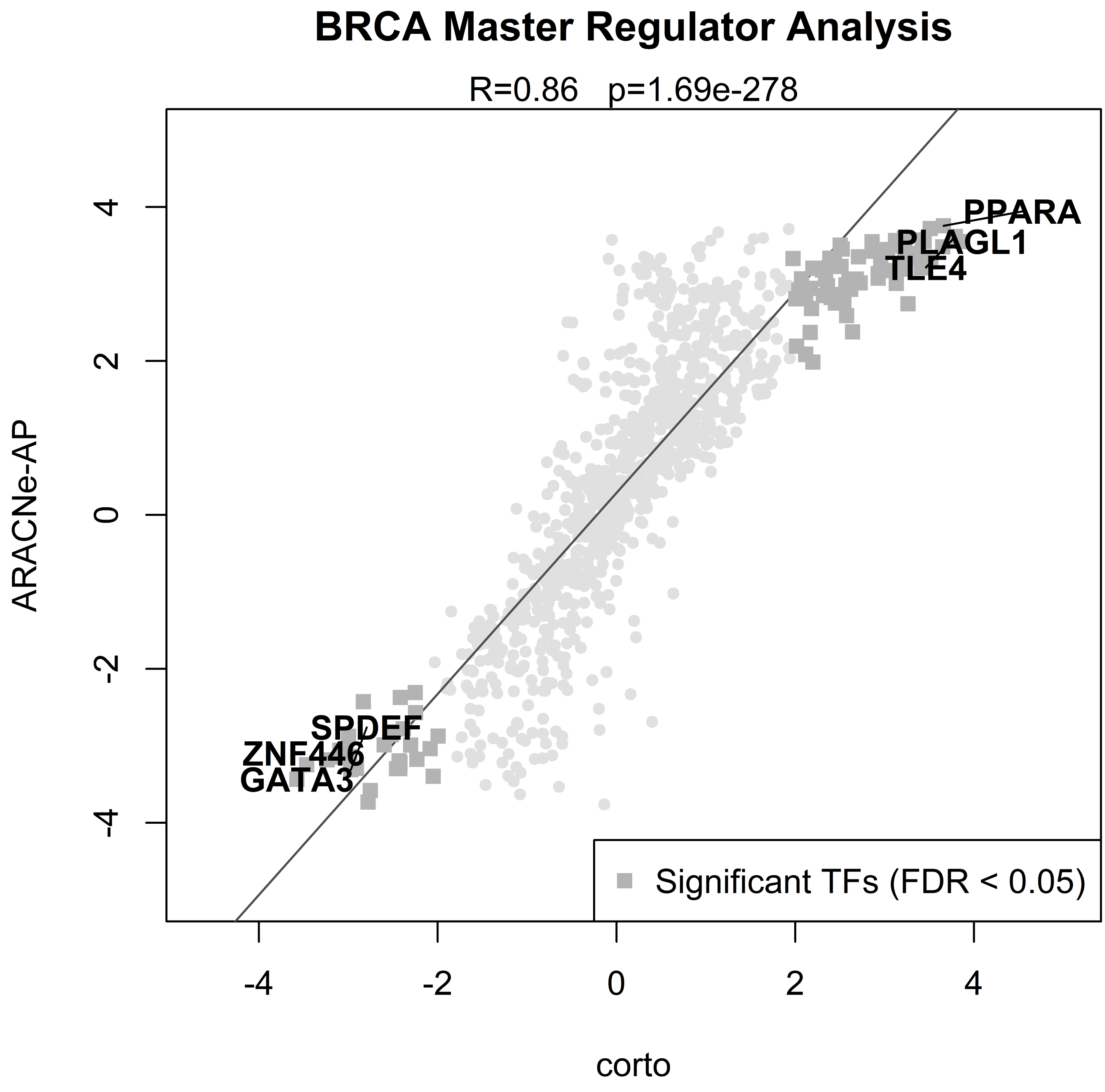
BRCA Master Regulator Analysis
R=0.86 p=1.69e-278
ARACNe-AP
corto
PPARA
PLAGL1
TLE4
SPDEF
ZNF446
GATA3
Significant TFs (FDR < 0.05)
4
2
0
-2
-4
-4
-2
0
2
4

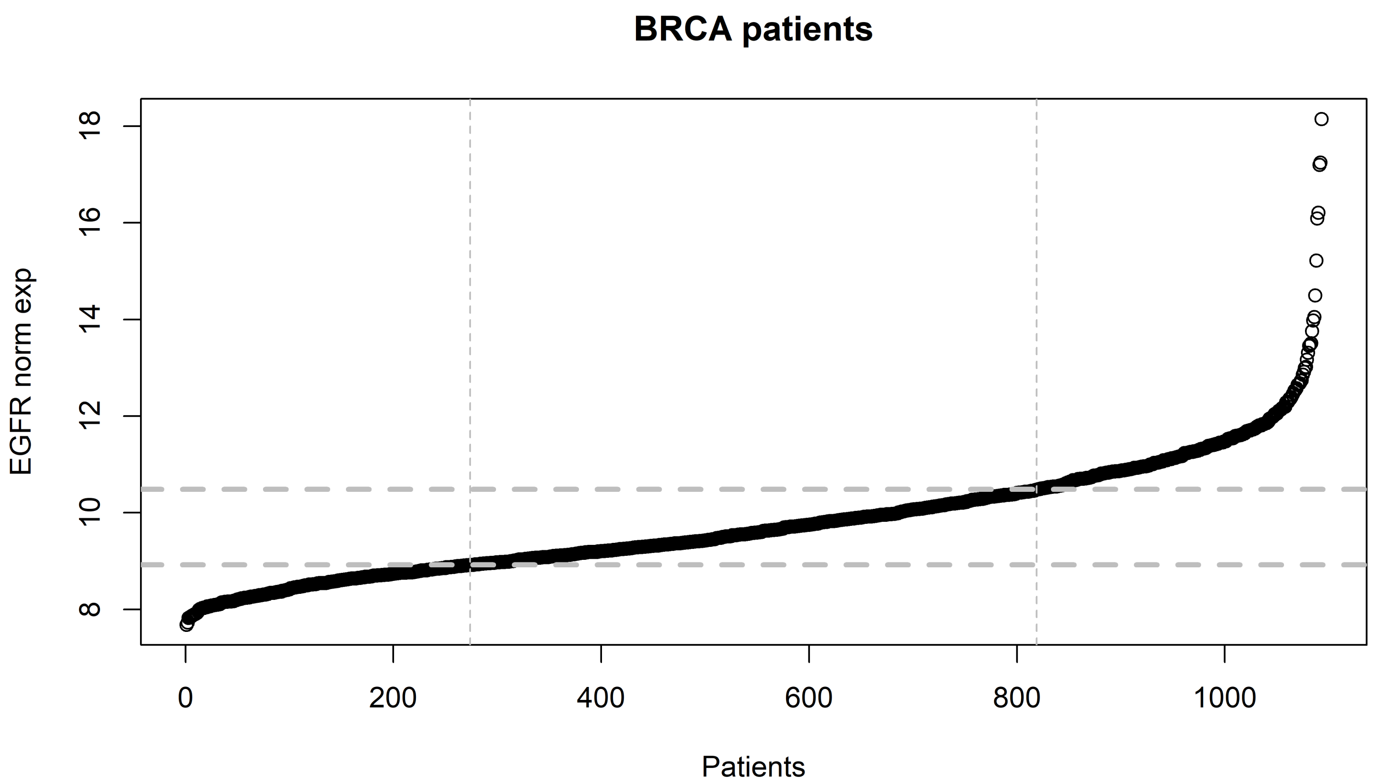
BRCA patients
EGFR norm exp
8
10
12
14
16
18
0
200
400
600
800
1000
Patients

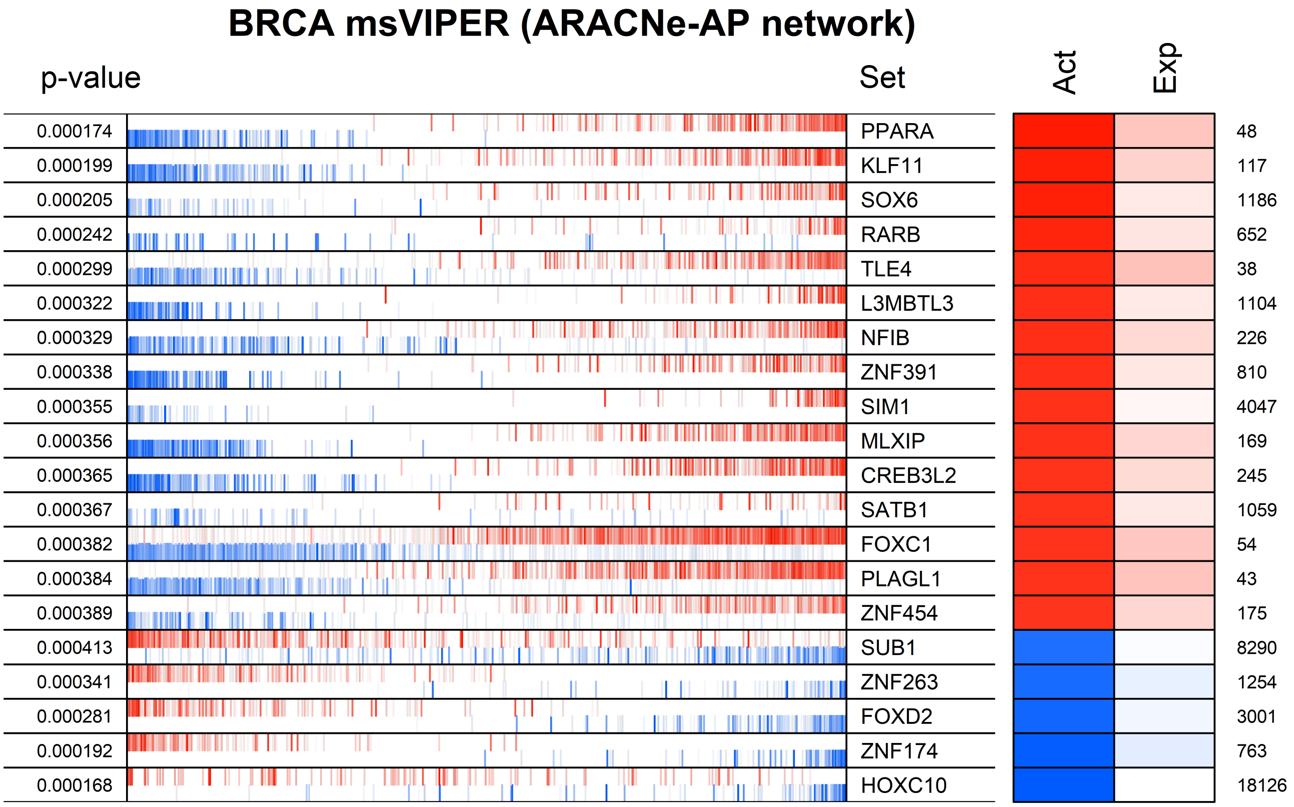
BRCA msVIPER (ARACNe-AP network)
p-value
Set
Act
Exp
0.000174 PPARA 48
0.000199 KLF11 117
0.000205 SOX6 1186
0.000242 RARB 652
0.000299 TLE4 38
0.000322 L3MBTL3 1104
0.000329 NFIB 226
0.000338 ZNF391 810
0.000355 SIM1 4047
0.000356 MLXIP 169
0.000365 CREB3L2 245
0.000367 SATB1 1059
0.000382 FOXC1 54
0.000384 PLAGL1 43
0.000389 ZNF454 175
0.000413 SUB1 8290
0.000341 ZNF263 1254
0.000281 FOXD2 3001
0.000192 ZNF174 763
0.000168 HOXC10 18126

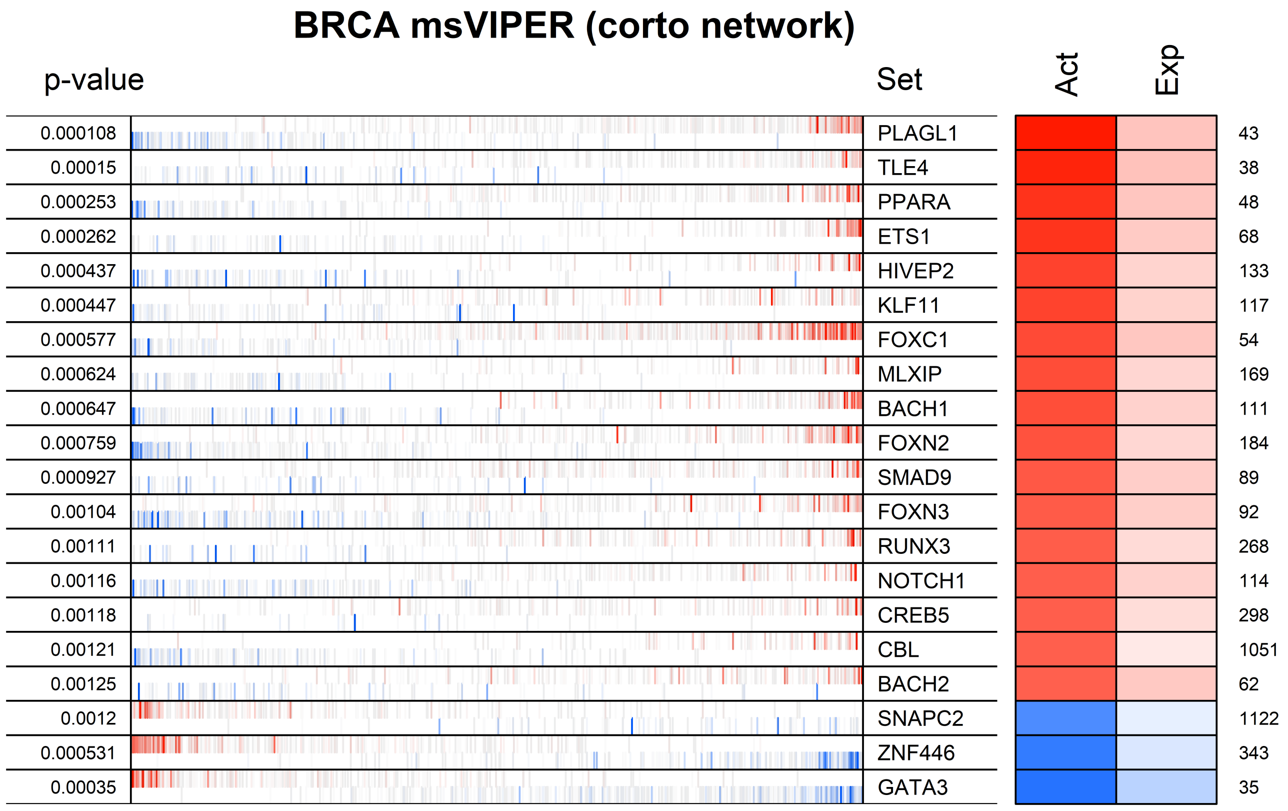
BRCA msVIPER (corto network)
p-value
Set
Act
Exp
0.000108
PLAGL1
43
0.00015
TLE4
38
0.000253
PPARA
48
0.000262
ETS1
68
0.000437
HIVEP2
133
0.000447
KLF11
117
0.000577
FOXC1
54
0.000624
MLXIP
169
0.000647
BACH1
111
0.000759
FOXN2
184
0.000927
SMAD9
89
0.00104
FOXN3
92
0.00111
RUNX3
268
0.00116
NOTCH1
114
0.00118
CREB5
298
0.00121
CBL
1051
0.00125
BACH2
62
0.0012
SNAPC2
1122
0.000531
ZNF446
343
0.00035
GATA3
35

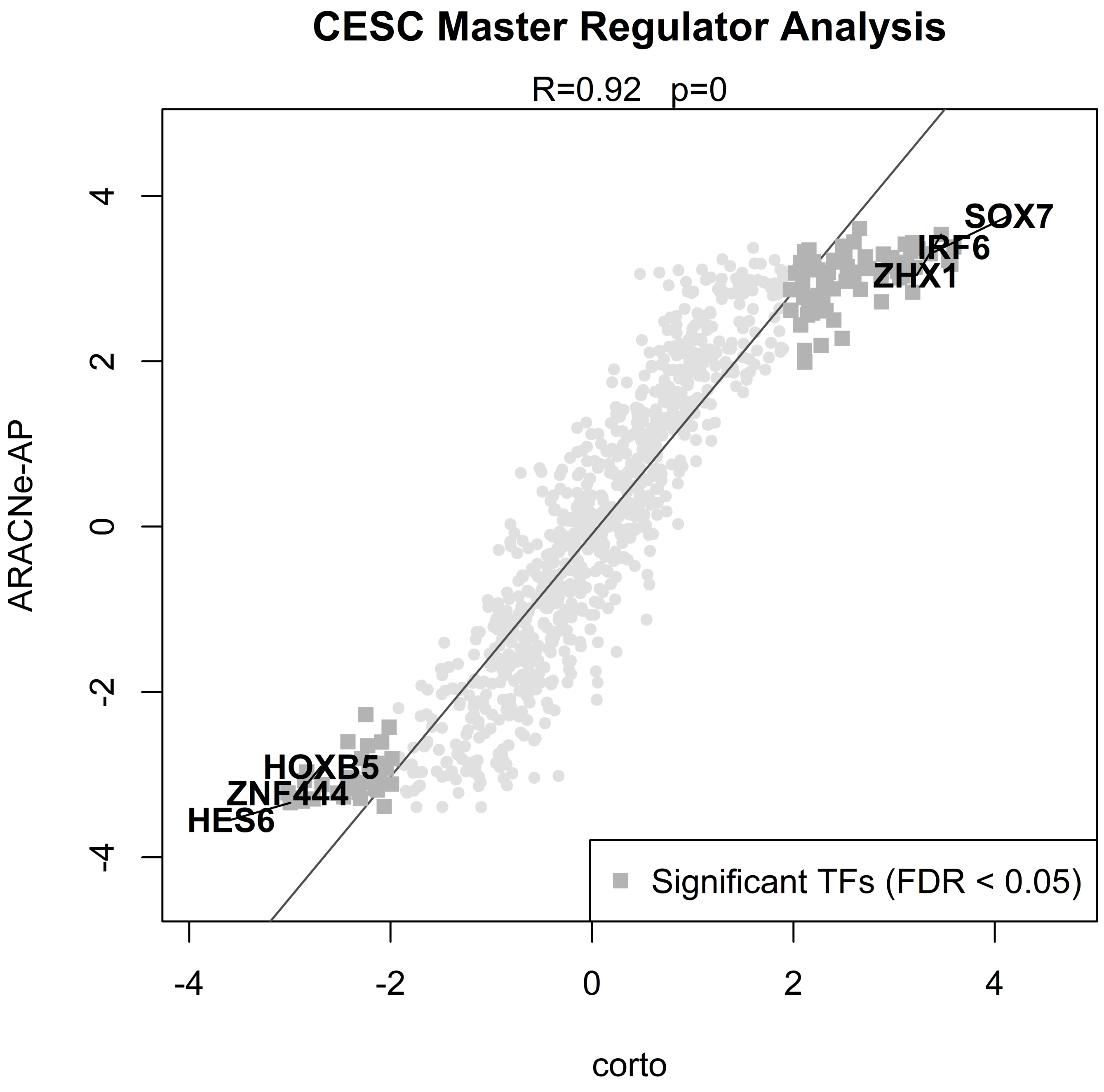

CESC Master Regulator Analysis
R=0.92 p=0
ARACNe-AP
corto
SOX7
IRF6
ZHX1
HOXB5
ZNF444
HES6
Significant TFs (FDR < 0.05)
-4
-2
0
2
4

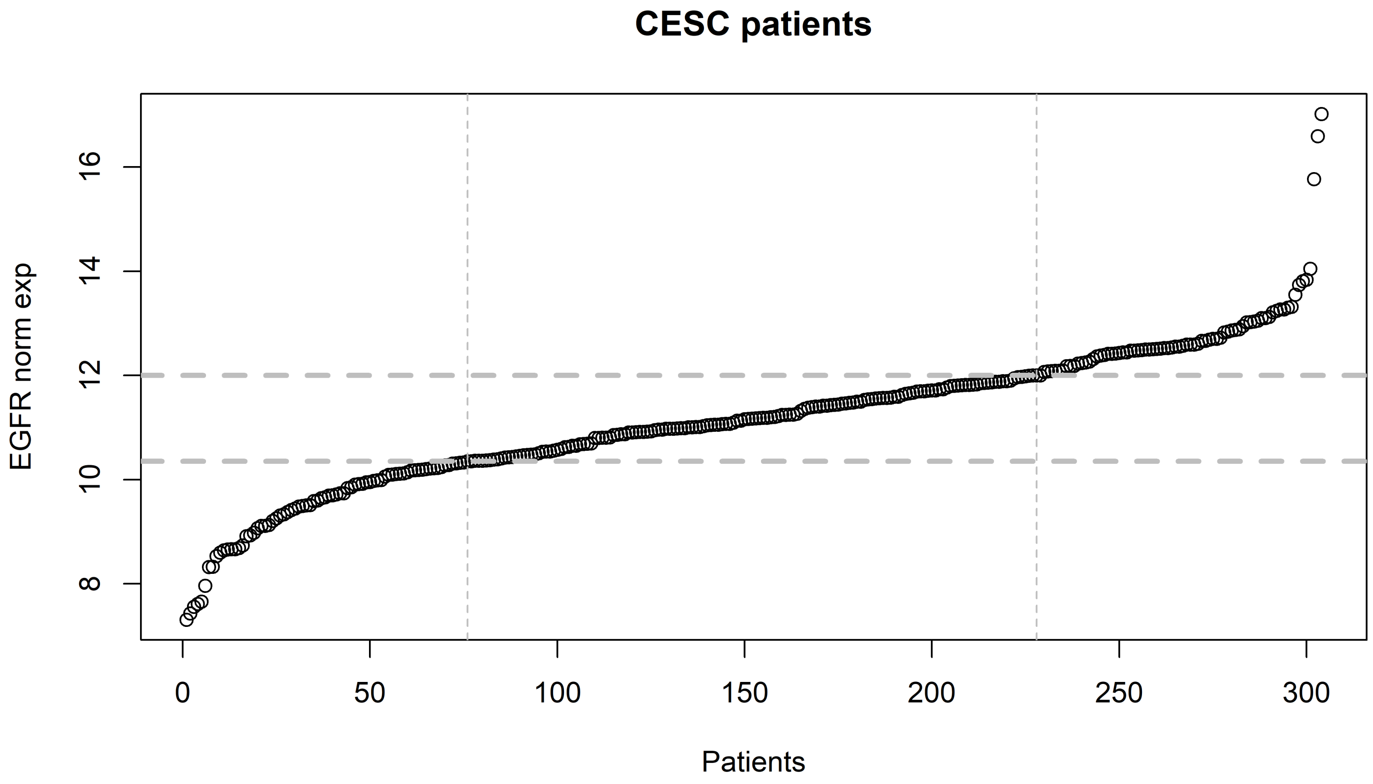

CESC patients
EGFR norm exp
Patients
8
10
12
14
16
0
50
100
150
200
250
300

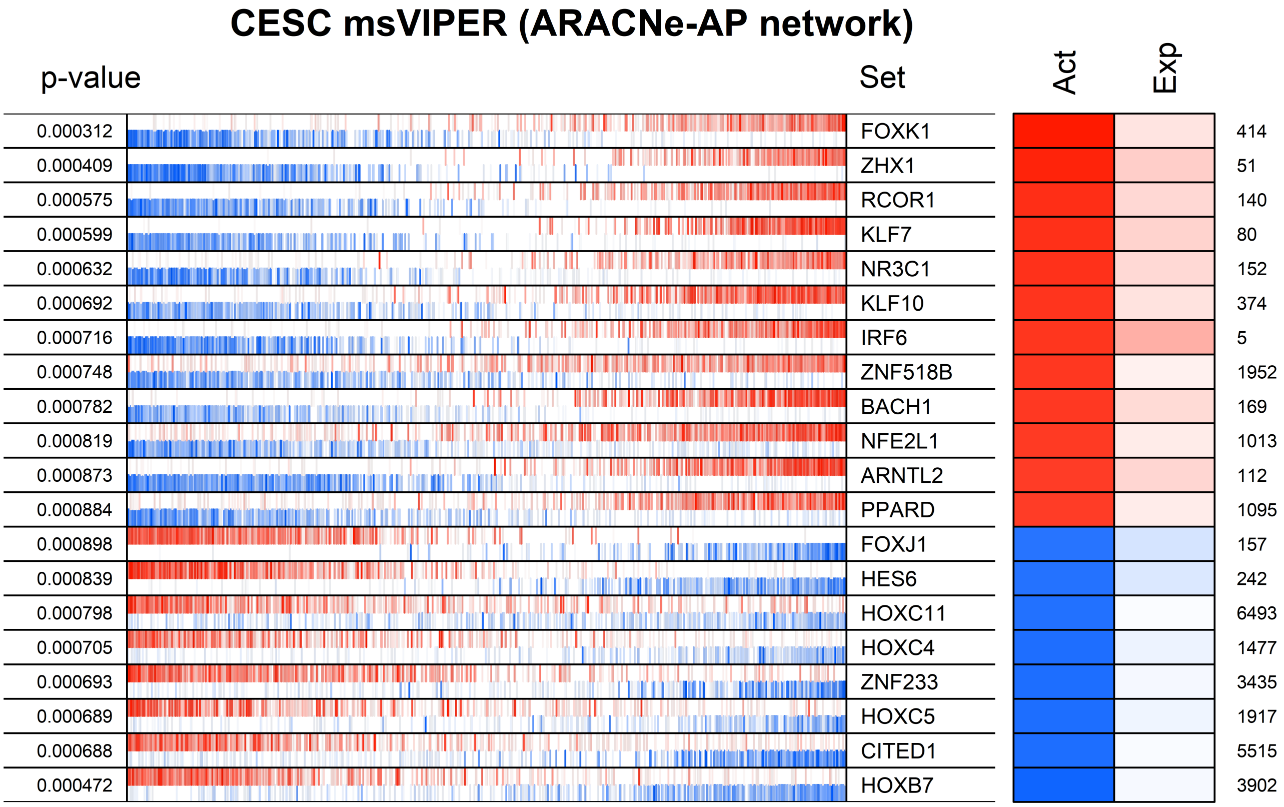
CESC msVIPER (ARACNe-AP network)
p-value
Set
Act
Exp
0.000312 FOXK1 414
0.000409 ZHX1 51
0.000575 RCOR1 140
0.000599 KLF7 80
0.000632 NR3C1 152
0.000692 KLF10 374
0.000716 IRF6 5
0.000748 ZNF518B 1952
0.000782 BACH1 169
0.000819 NFE2L1 1013
0.000873 ARNTL2 112
0.000884 PPARD 1095
0.000898 FOXJ1 157
0.000839 HES6 242
0.000798 HOXC11 6493
0.000705 HOXC4 1477
0.000693 ZNF233 3435
0.000689 HOXC5 1917
0.000688 CITED1 5515
0.000472 HOXB7 3902

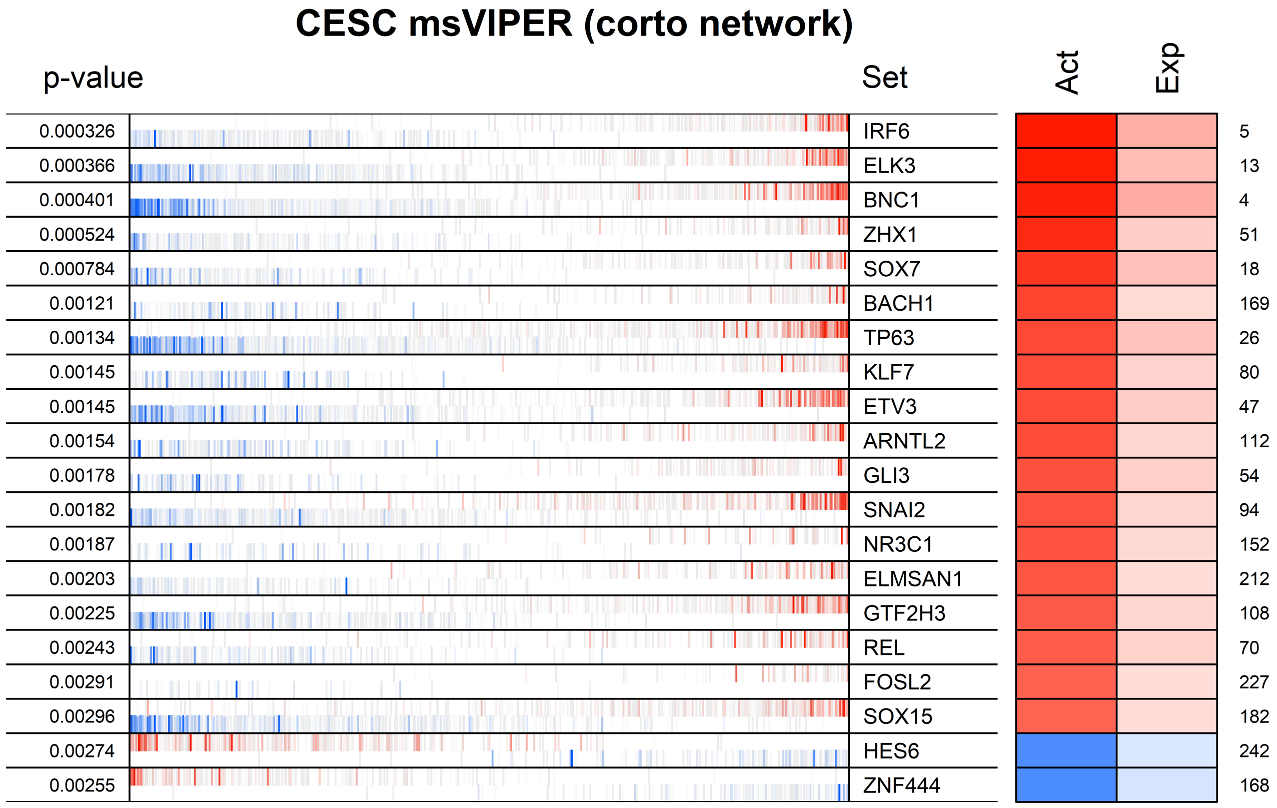
CESC msVIPER (corto network)
p-value
Set
Act
Exp
0.000326 IRF6 5
0.000366 ELK3 13
0.000401 BNC1 4
0.000524 ZHX1 51
0.000784 SOX7 18
0.00121 BACH1 169
0.00134 TP63 26
0.00145 KLF7 80
0.00145 ETV3 47
0.00154 ARNTL2 112
0.00178 GLI3 54
0.00182 SNAI2 94
0.00187 NR3C1 152
0.00203 ELMSAN1 212
0.00225 GTF2H3 108
0.00243 REL 70
0.00291 FOSL2 227
0.00296 SOX15 182
0.00274 HES6 242
0.00255 ZNF444 168

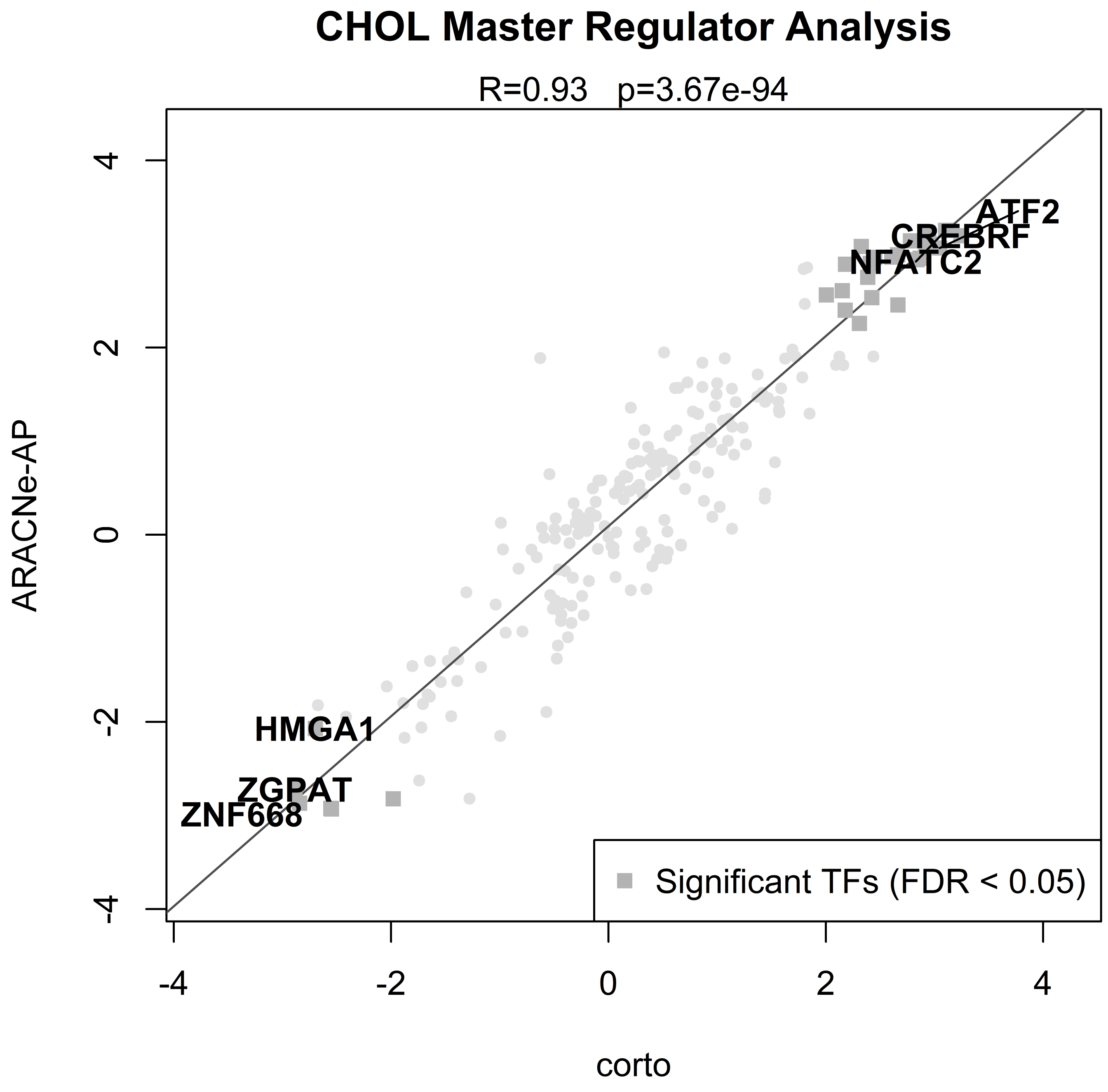
CHOL Master Regulator Analysis
R=0.93 p=3.67e-94
ARACNe-AP
corto
4
2
0
-2
-4
-4
-2
0
2
4
ATF2
CREBRF
NFATC2
HMGA1
ZGPAT
ZNF668
Significant TFs (FDR < 0.05)

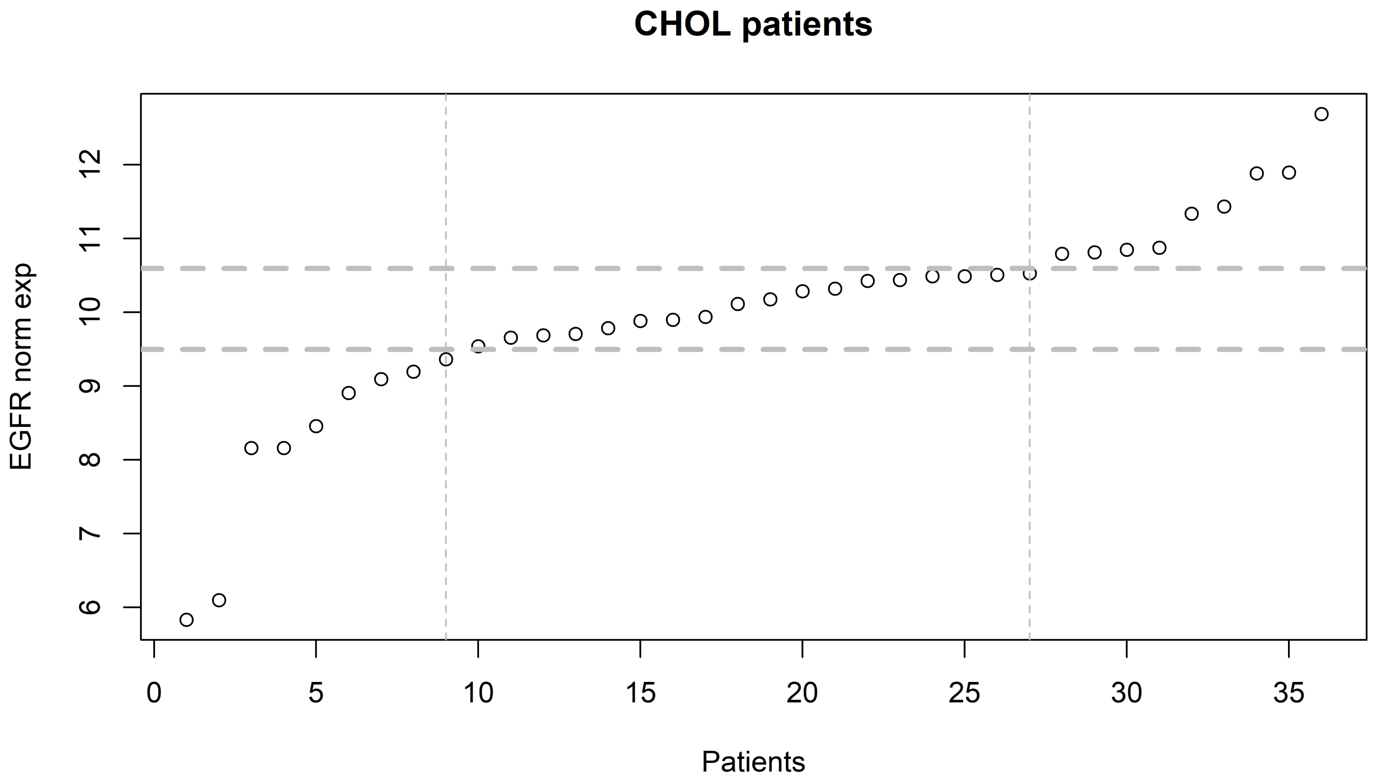

CHOL patients
EGFR norm exp
Patients
6
7
8
9
10
11
12
0
5
10
15
20
25
30
35

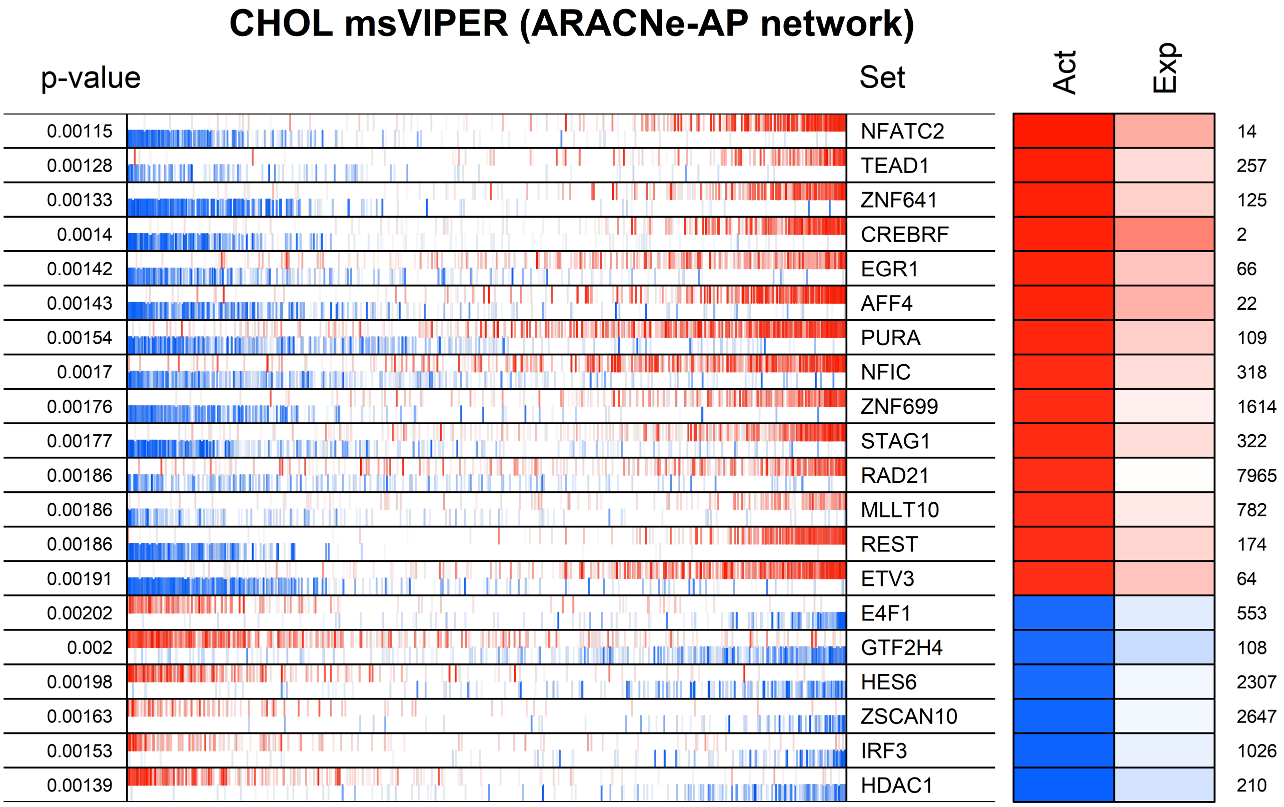
CHOL msVIPER (ARACNe-AP network)
p-value
Set
Act
Exp
0.00115
NFATC2
14
0.00128
TEAD1
257
0.00133
ZNF641
125
0.0014
CREBRF
2
0.00142
EGR1
66
0.00143
AFF4
22
0.00154
PURA
109
0.0017
NFIC
318
0.00176
ZNF699
1614
0.00177
STAG1
322
0.00186
RAD21
7965
0.00186
MLLT10
782
0.00186
REST
174
0.00191
ETV3
64
0.00202
E4F1
553
0.002
GTF2H4
108
0.00198
HES6
2307
0.00163
ZSCAN10
2647
0.00153
IRF3
1026
0.00139
HDAC1
210

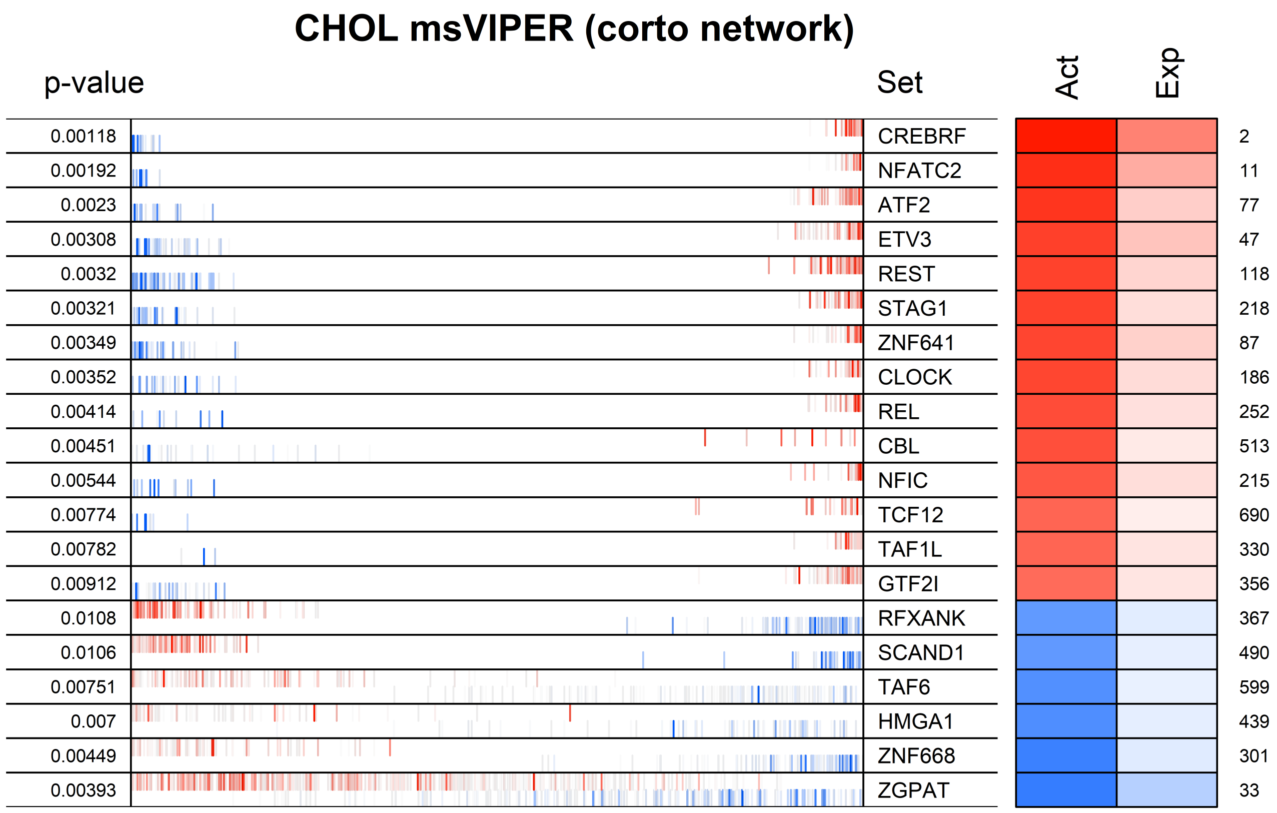

CHOL msVIPER (corto network)
p-value
Set
Act
Exp
0.00118
CREBRF
2
0.00192
NFATC2
11
0.0023
ATF2
77
0.00308
ETV3
47
0.0032
REST
118
0.00321
STAG1
218
0.00349
ZNF641
87
0.00352
CLOCK
186
0.00414
REL
252
0.00451
CBL
513
0.00544
NFIC
215
0.00774
TCF12
690
0.00782
TAF1L
330
0.00912
GTF2I
356
0.0108
RFXANK
367
0.0106
SCAND1
490
0.00751
TAF6
599
0.007
HMGA1
439
0.00449
ZNF668
301
0.00393
ZGPAT
33

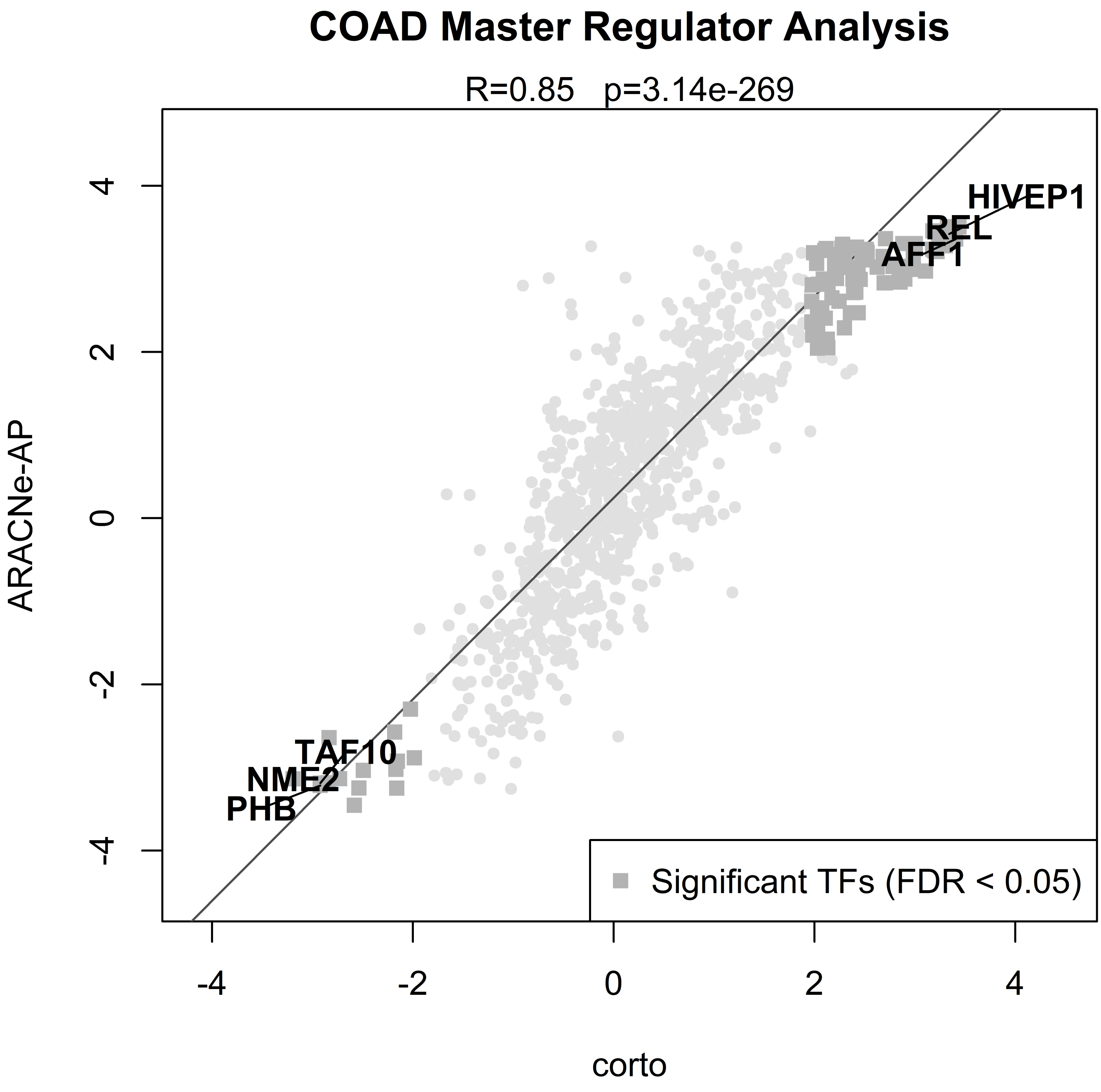

COAD Master Regulator Analysis
R=0.85 p=3.14e-269
ARACNe-AP
corto
HIVEP1
REL
AFF1
TAF10
NME2
PHB
Significant TFs (FDR < 0.05)
-4
-2
0
2
4

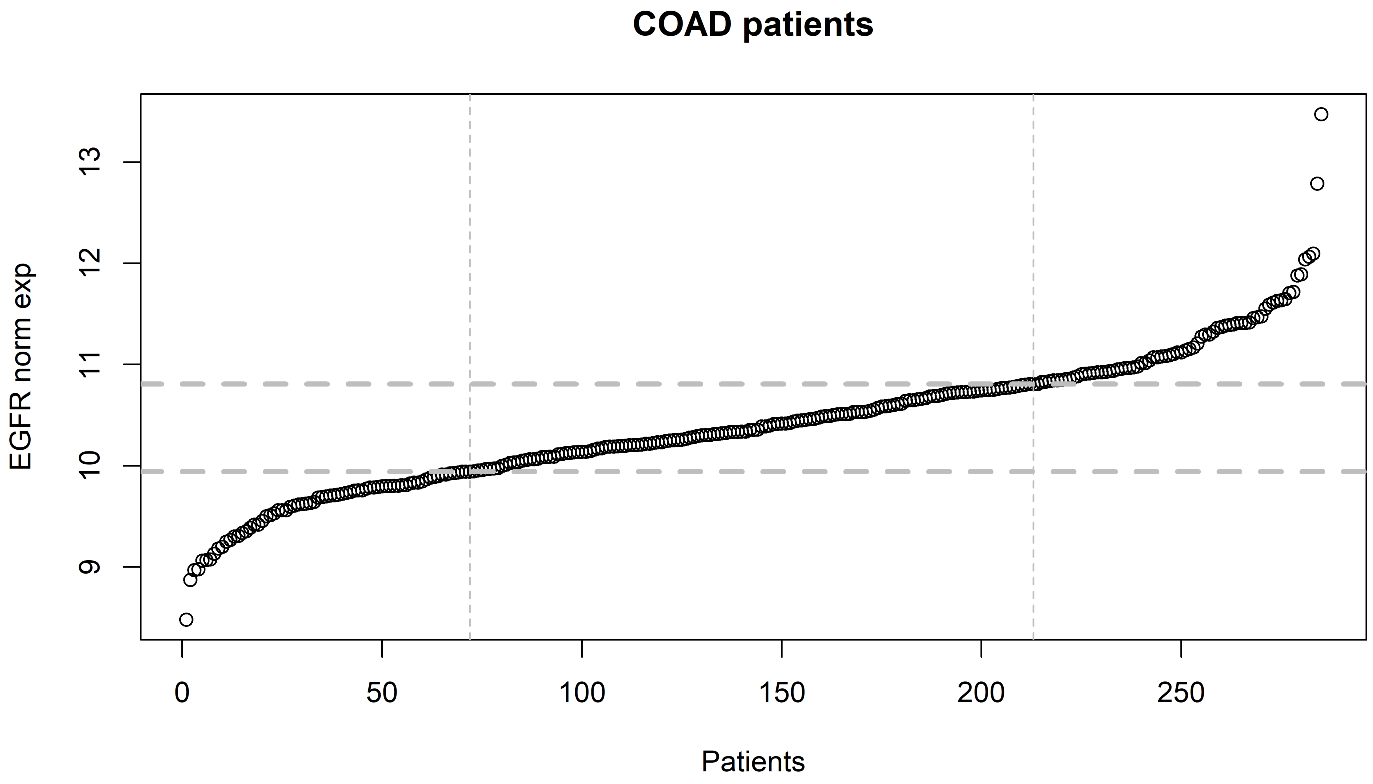
COAD patients
EGFR norm exp
9
10
11
12
13
0
50
100
150
200
250
Patients

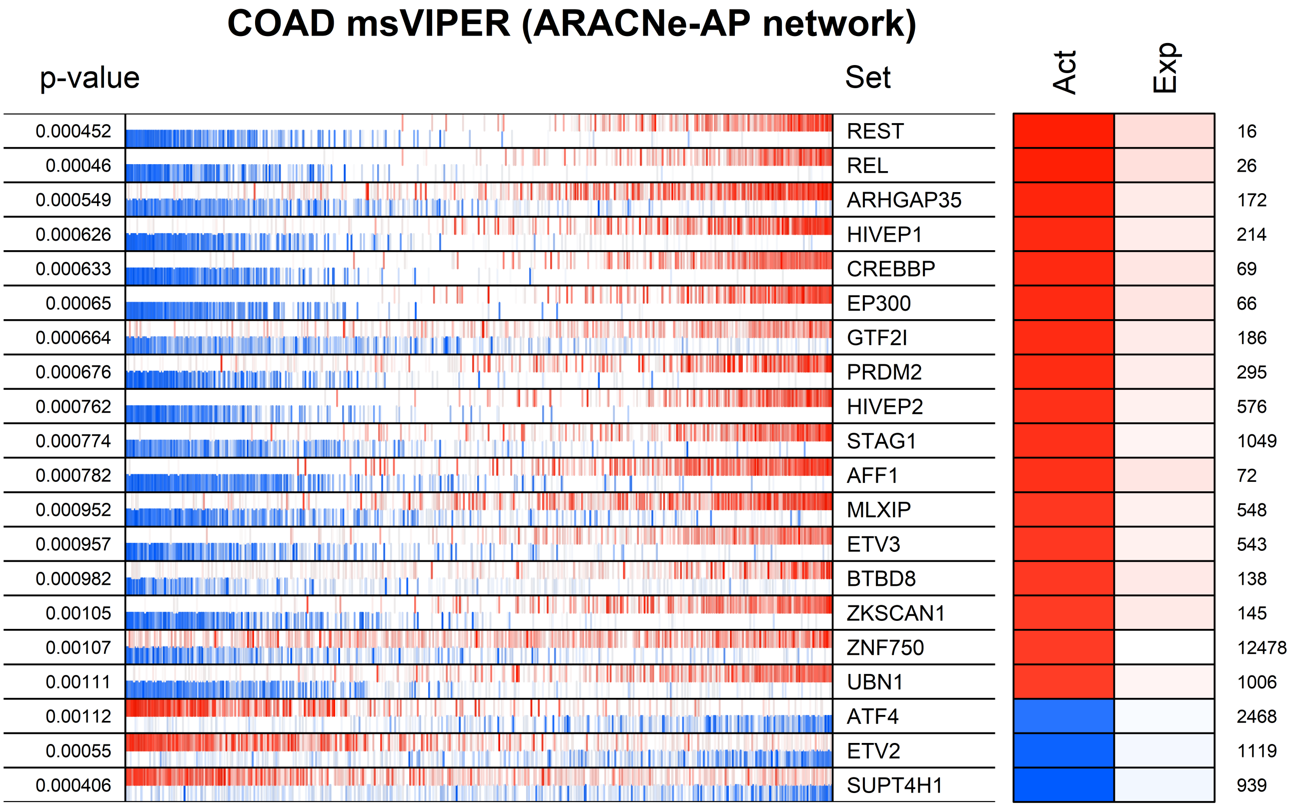
COAD msVIPER (ARACNe-AP network)
p-value
Set
Act
Exp
0.000452 REST 16
0.00046 REL 26
0.000549 ARHGAP35 172
0.000626 HIVEP1 214
0.000633 CREBBP 69
0.00065 EP300 66
0.000664 GTF2I 186
0.000676 PRDM2 295
0.000762 HIVEP2 576
0.000774 STAG1 1049
0.000782 AFF1 72
0.000952 MLXIP 548
0.000957 ETV3 543
0.000982 BTBD8 138
0.00105 ZKSCAN1 145
0.00107 ZNF750 12478
0.00111 UBN1 1006
0.00112 ATF4 2468
0.00055 ETV2 1119
0.000406 SUPT4H1 939

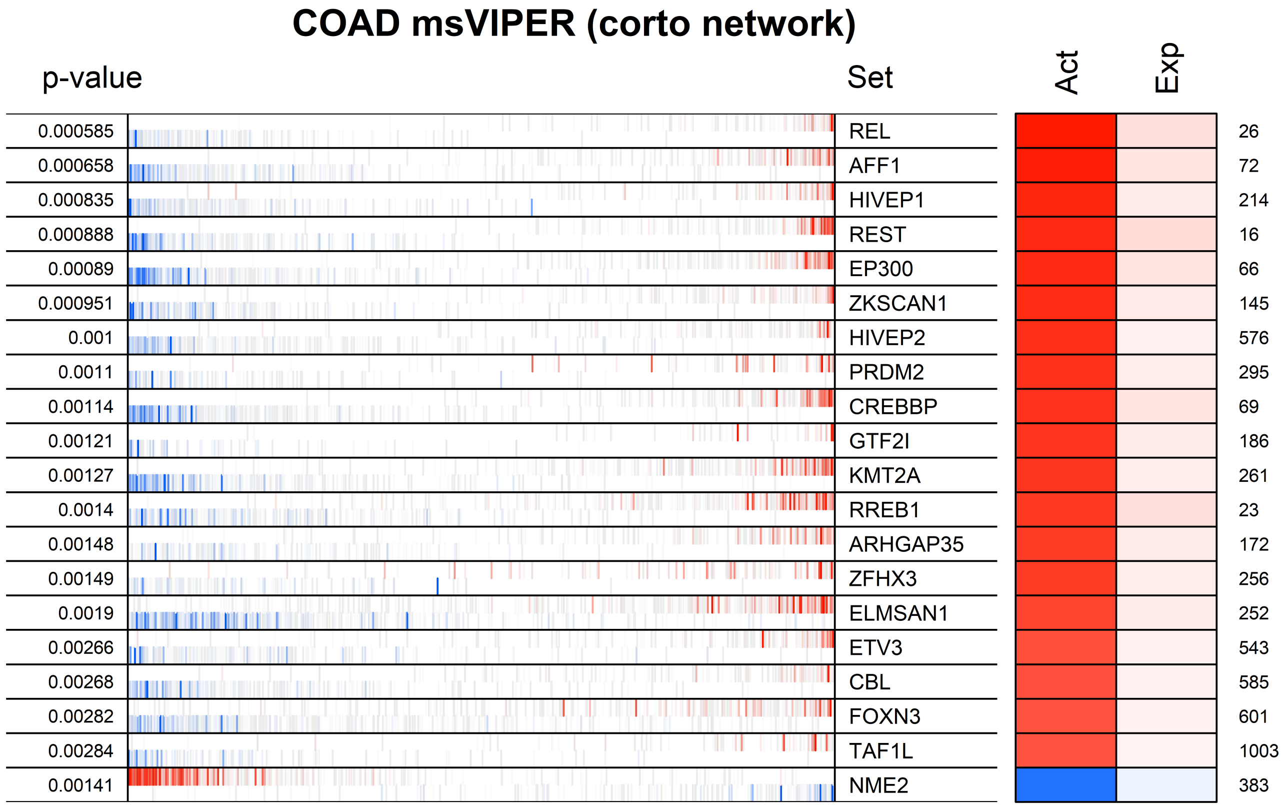

COAD msVIPER (corto network)
p-value
Set
Act
Exp
0.000585
0.000658
0.000835
0.000888
0.00089
0.000951
0.001
0.0011
0.00114
0.00121
0.00127
0.0014
0.00148
0.00149
0.0019
0.00266
0.00268
0.00282
0.00284
0.00141
REL
AFF1
HIVEP1
REST
EP300
ZKSCAN1
HIVEP2
PRDM2
CREBBP
GTF2I
KMT2A
RREB1
ARHGAP35
ZFHX3
ELMSAN1
ETV3
CBL
FOXN3
TAF1L
NME2
26
72
214
16
66
145
576
295
69
186
261
23
172
256
252
543
585
601
1003
383

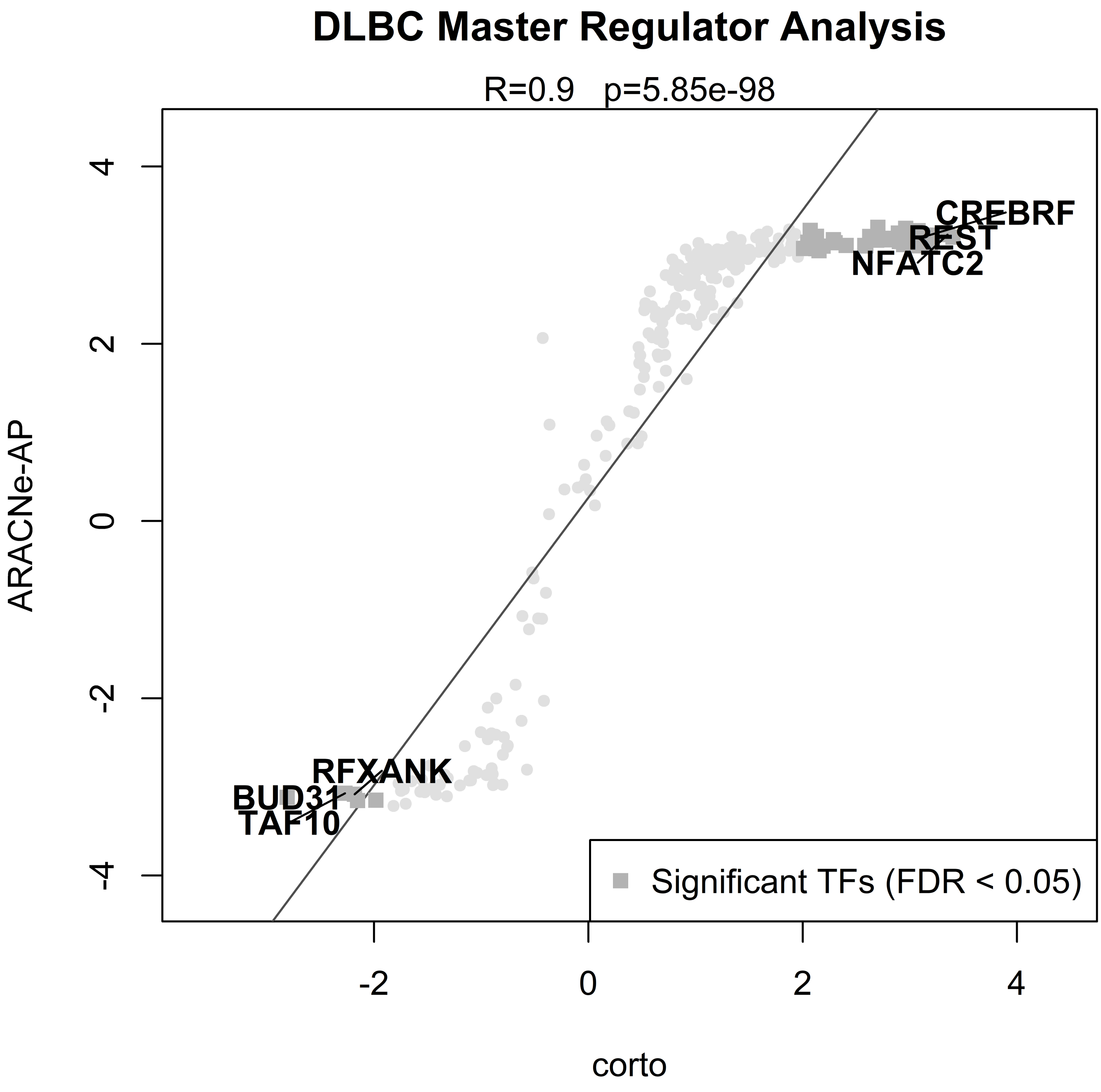

DLBC Master Regulator Analysis
R=0.9 p=5.85e-98
ARACNe-AP
corto
CREBRF
REST
NFATC2
RFXANK
BUD31
TAF10
Significant TFs (FDR < 0.05)
4
2
0
-2
-4
-2
0
2
4

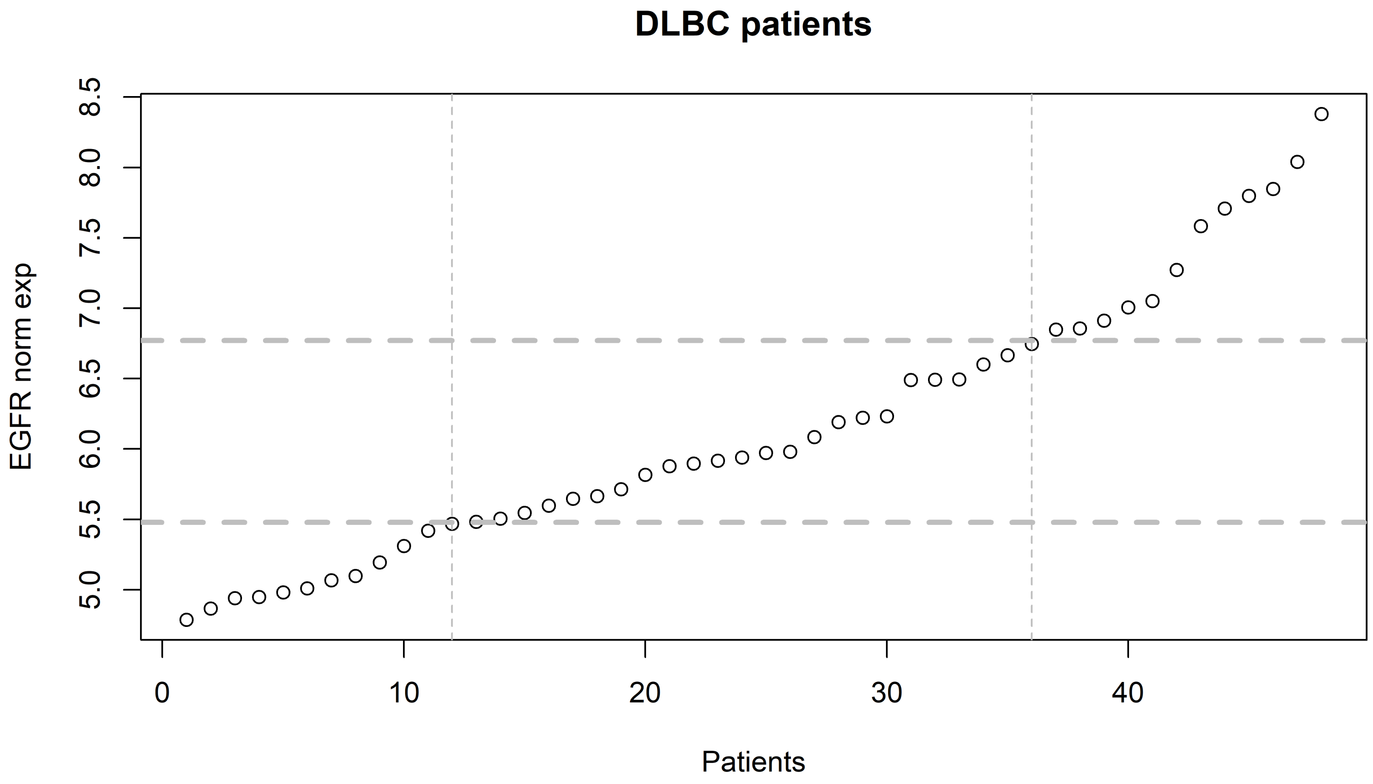
DLBC patients
EGFR norm exp
Patients
5.0
5.5
6.0
6.5
7.0
7.5
8.0
8.5
0
10
20
30
40

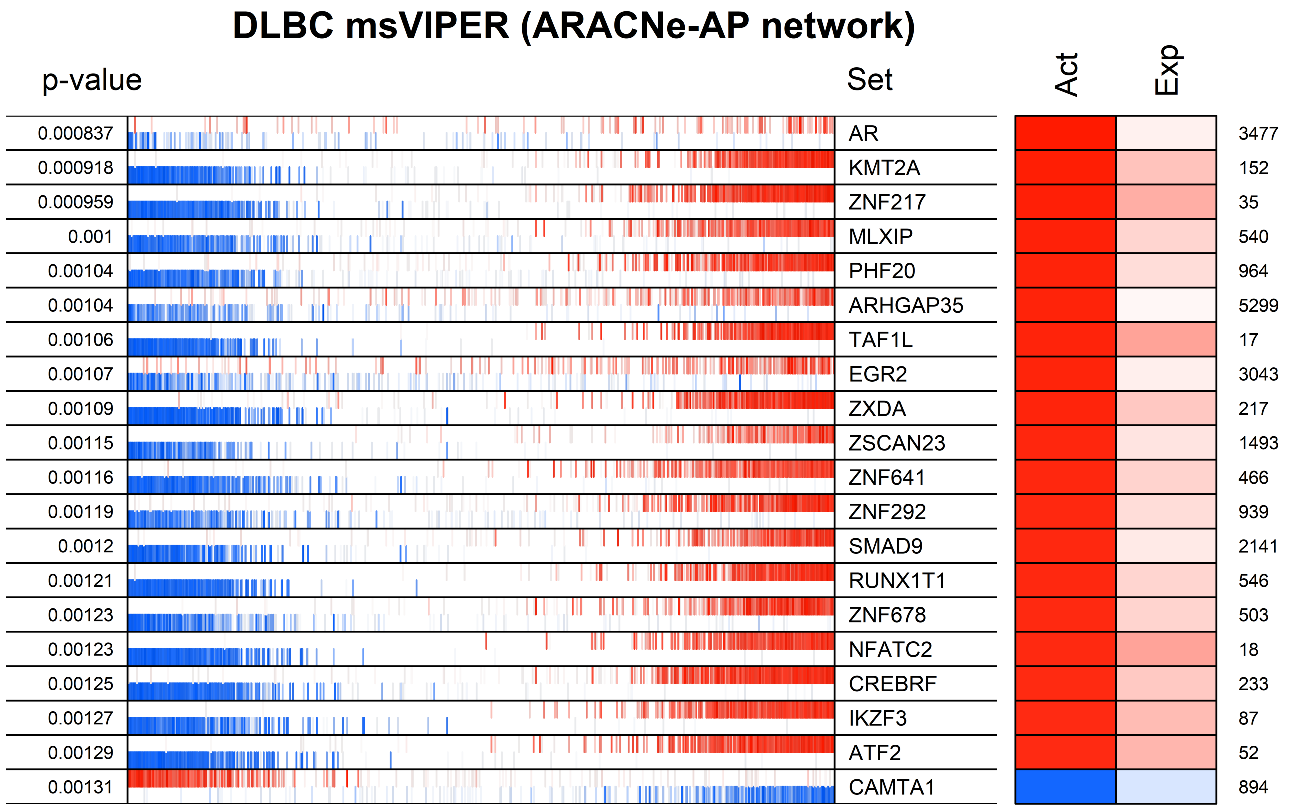

DLBC msVIPER (ARACNe-AP network)
p-value
Set
Act
Exp
0.000837
AR
3477
0.000918
KMT2A
152
0.000959
ZNF217
35
0.001
MLXIP
540
0.00104
PHF20
964
0.00104
ARHGAP35
5299
0.00106
TAF1L
17
0.00107
EGR2
3043
0.00109
ZXDA
217
0.00115
ZSCAN23
1493
0.00116
ZNF641
466
0.00119
ZNF292
939
0.0012
SMAD9
2141
0.00121
RUNX1T1
546
0.00123
ZNF678
503
0.00123
NFATC2
18
0.00125
CREBRF
233
0.00127
IKZF3
87
0.00129
ATF2
52
0.00131
CAMTA1
894

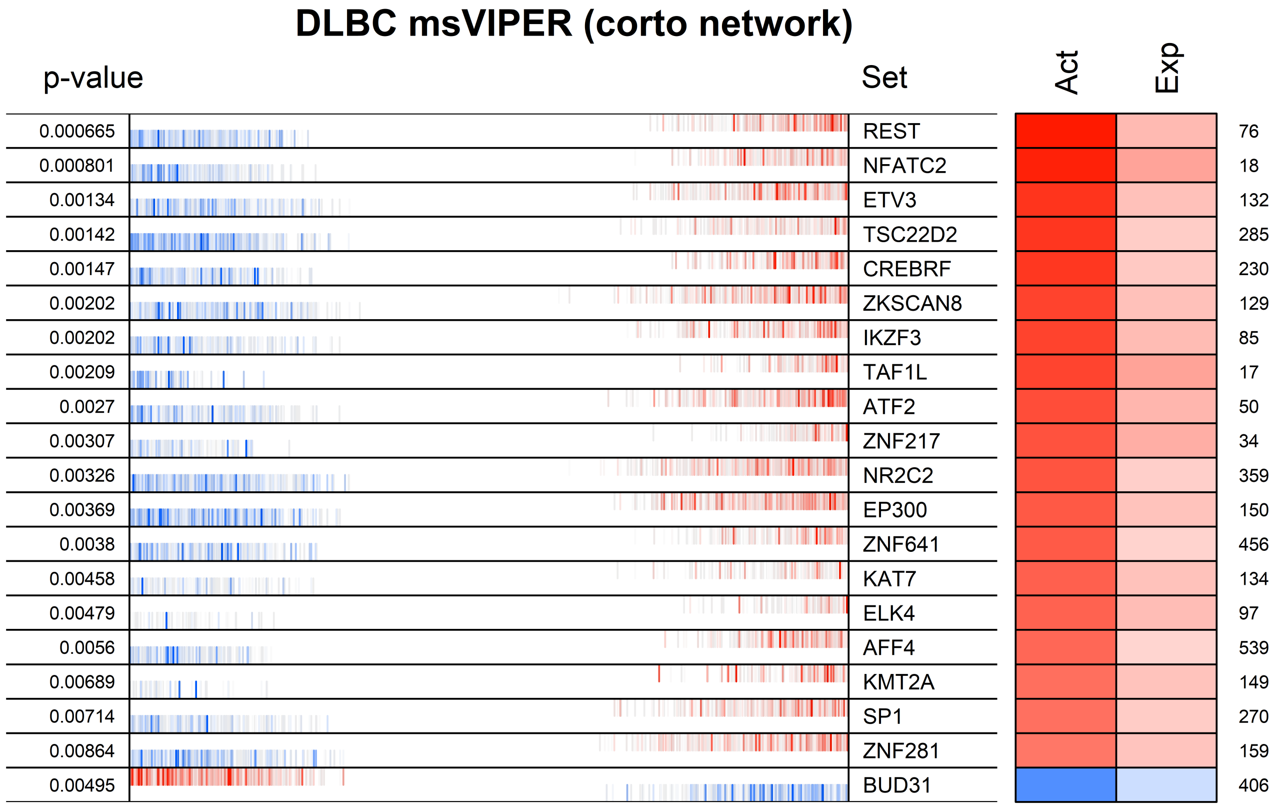
DLBC msVIPER (corto network)
p-value
Set
Act
Exp
0.000665 REST 76
0.000801 NFATC2 18
0.00134 ETV3 132
0.00142 TSC22D2 285
0.00147 CREBRF 230
0.00202 ZKSCAN8 129
0.00202 IKZF3 85
0.00209 TAF1L 17
0.0027 ATF2 50
0.00307 ZNF217 34
0.00326 NR2C2 359
0.00369 EP300 150
0.0038 ZNF641 456
0.00458 KAT7 134
0.00479 ELK4 97
0.0056 AFF4 539
0.00689 KMT2A 149
0.00714 SP1 270
0.00864 ZNF281 159
0.00495 BUD31 406

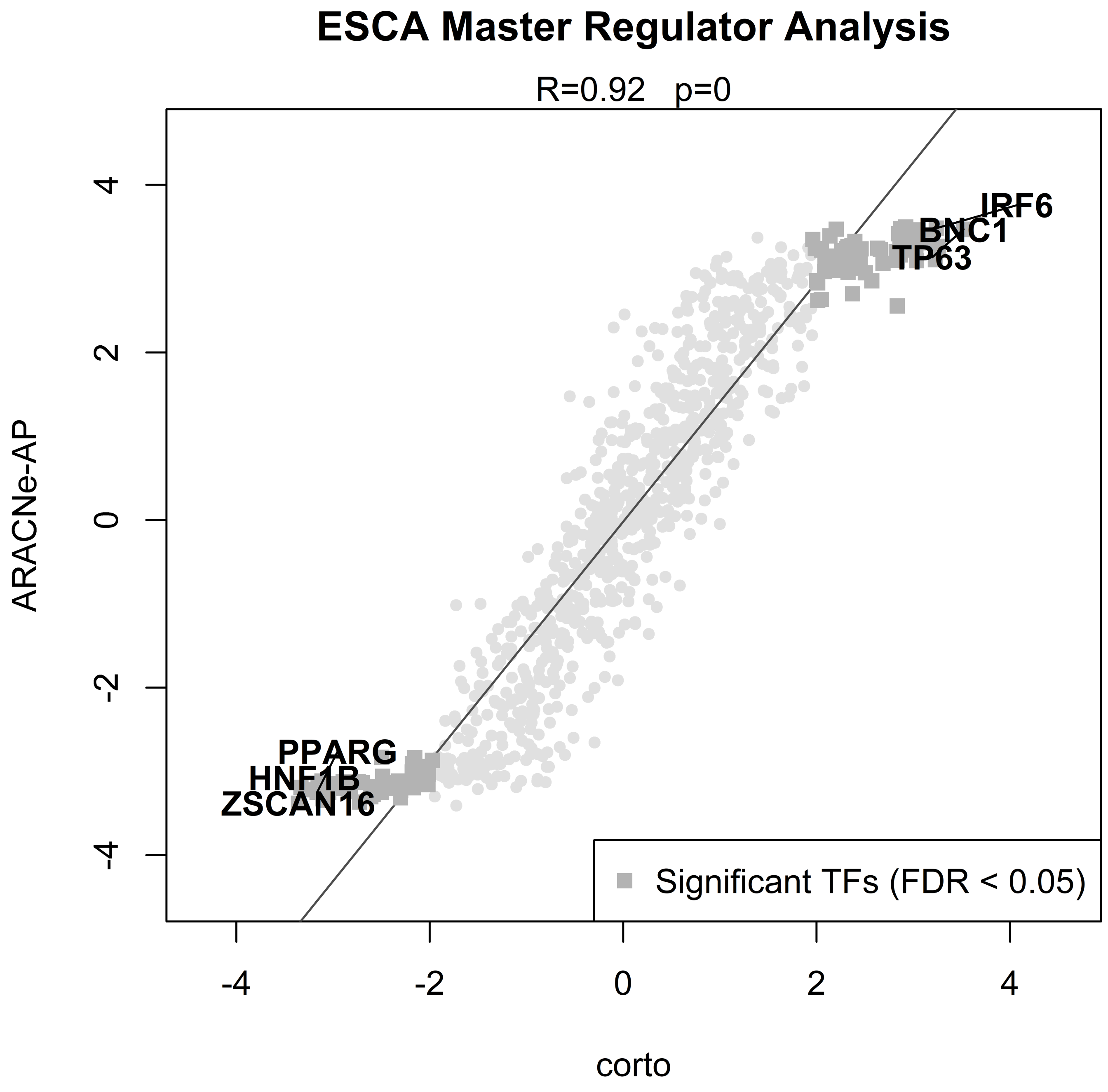

ESCA Master Regulator Analysis
R=0.92 p=0
ARACNe-AP
corto
IRF6
BNC1
TP63
PPARG
HNF1B
ZSCAN16
Significant TFs (FDR < 0.05)
4
2
0
-2
-4
-4
-2
0
2
4

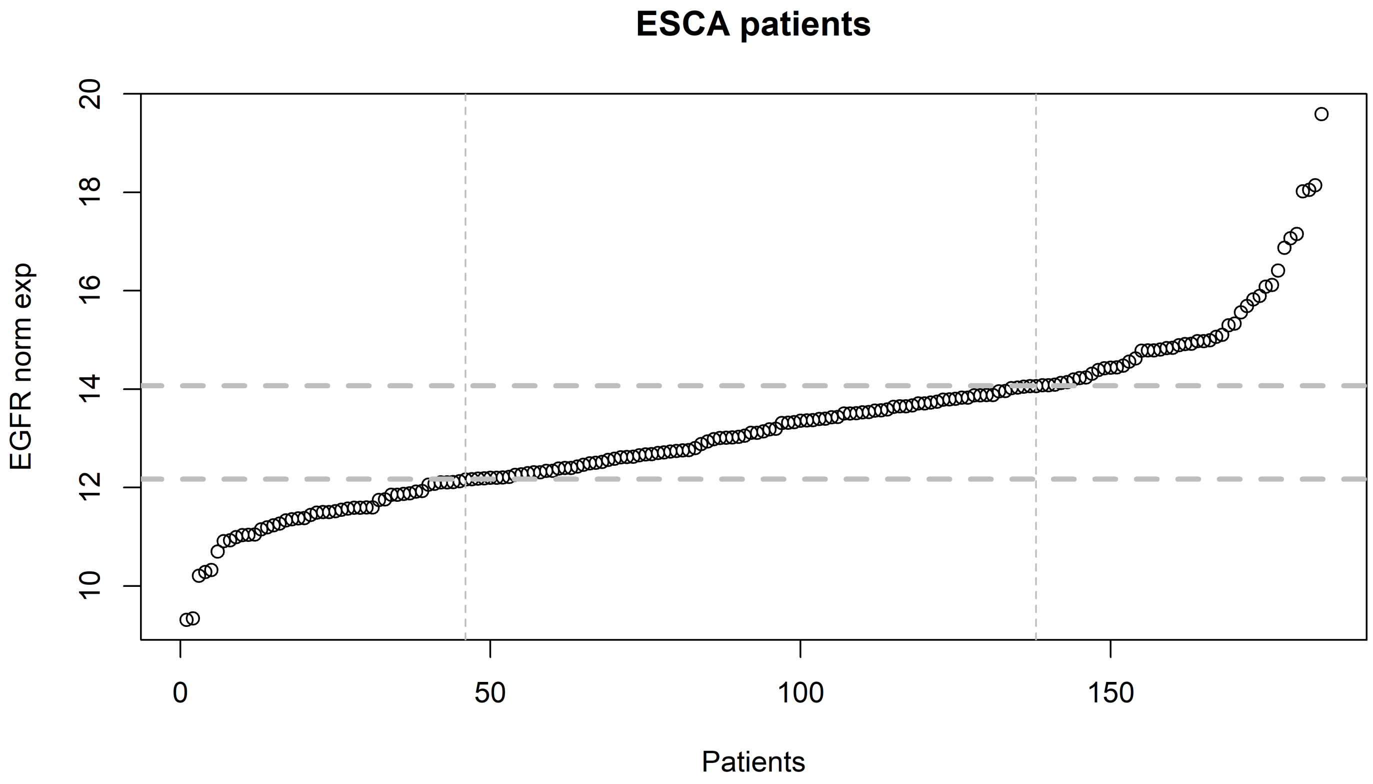
ESCA patients
EGFR norm exp
Patients
10
12
14
16
18
20
0
50
100
150

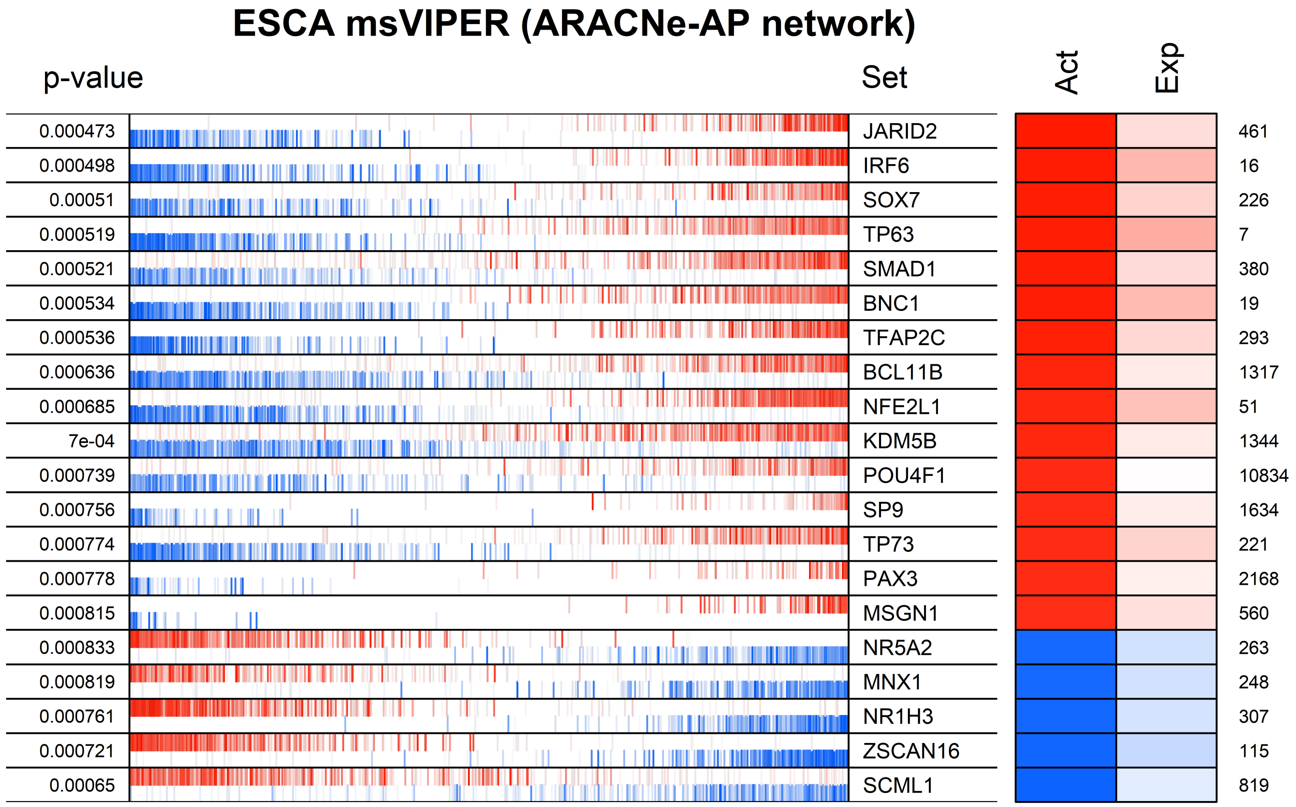
ESCA msVIPER (ARACNe-AP network)
p-value
Set
Act
Exp
0.000473 JARID2 461
0.000498 IRF6 16
0.00051 SOX7 226
0.000519 TP63 7
0.000521 SMAD1 380
0.000534 BNC1 19
0.000536 TFAP2C 293
0.000636 BCL11B 1317
0.000685 NFE2L1 51
7e-04 KDM5B 1344
0.000739 POU4F1 10834
0.000756 SP9 1634
0.000774 TP73 221
0.000778 PAX3 2168
0.000815 MSGN1 560
0.000833 NR5A2 263
0.000819 MNX1 248
0.000761 NR1H3 307
0.000721 ZSCAN16 115
0.00065 SCML1 819

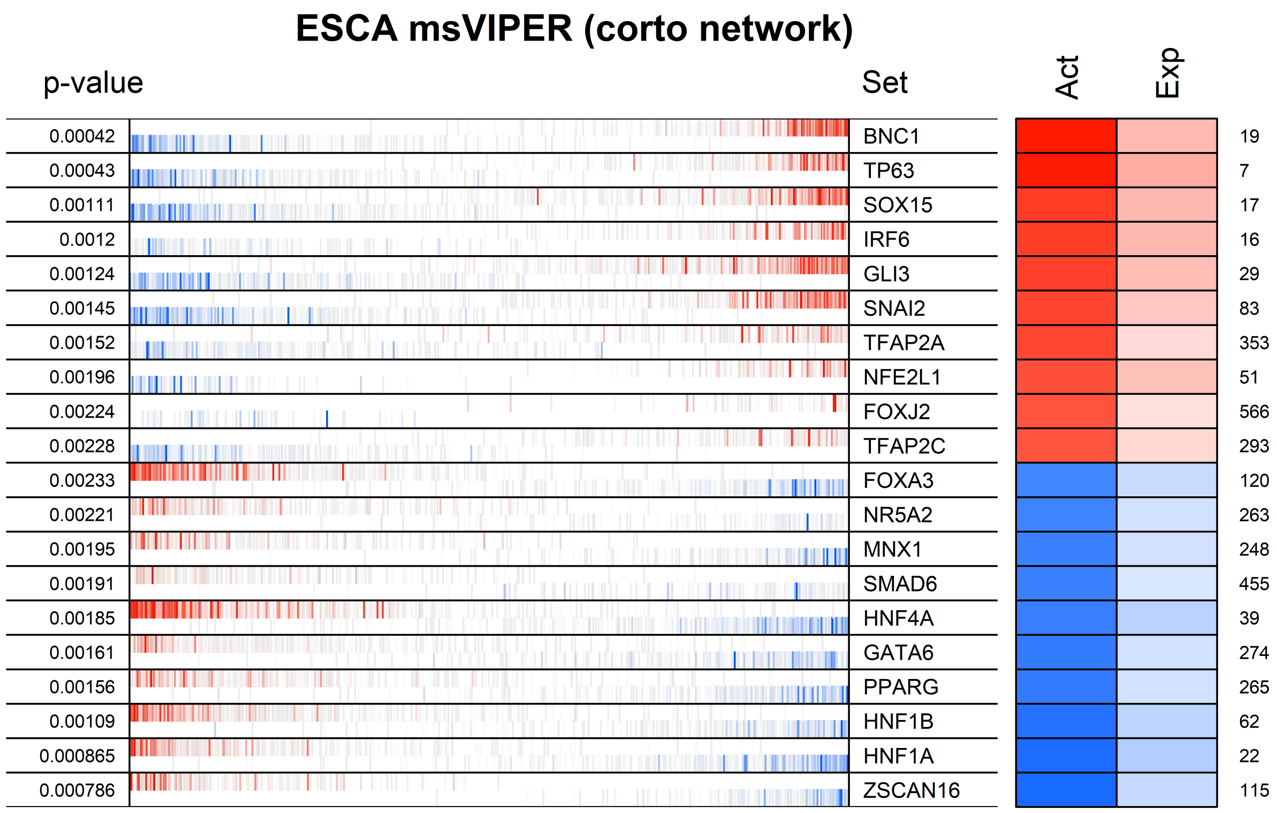

ESCA msVIPER (corto network)
p-value
Set
Act
Exp
0.00042
BNC1
19
0.00043
TP63
7
0.00111
SOX15
17
0.0012
IRF6
16
0.00124
GLI3
29
0.00145
SNAI2
83
0.00152
TFAP2A
353
0.00196
NFE2L1
51
0.00224
FOXJ2
566
0.00228
TFAP2C
293
0.00233
FOXA3
120
0.00221
NR5A2
263
0.00195
MNX1
248
0.00191
SMAD6
455
0.00185
HNF4A
39
0.00161
GATA6
274
0.00156
PPARG
265
0.00109
HNF1B
62
0.000865
HNF1A
22
0.000786
ZSCAN16
115

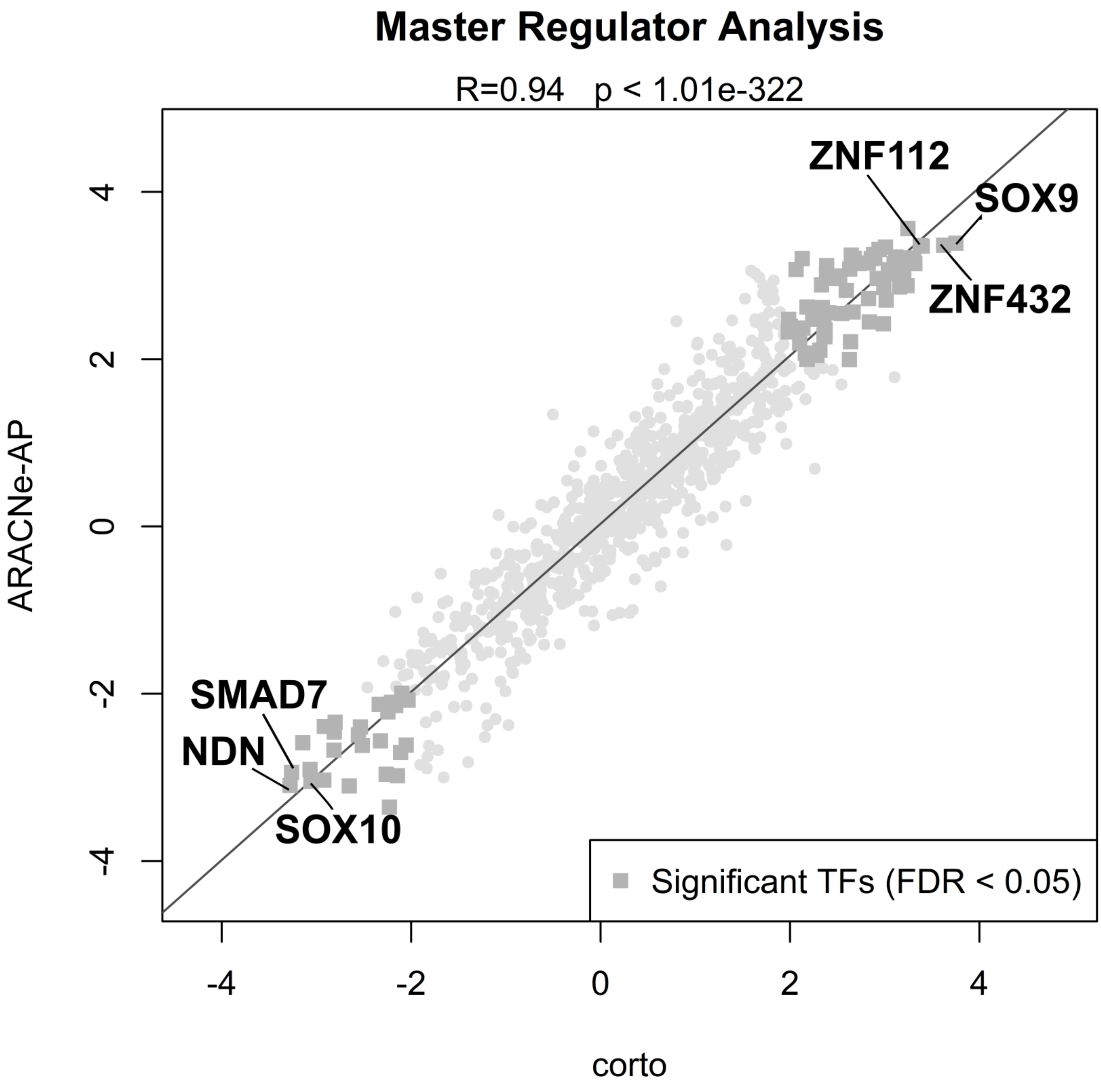

Master Regulator Analysis
R=0.94 p < 1.01e-322
ZNF112
SOX9
ZNF432
SMAD7
NDN
SOX10
Significant TFs (FDR < 0.05)
ARACNe-AP
corto
4
2
0
-2
-4
-4
-2
0
2
4

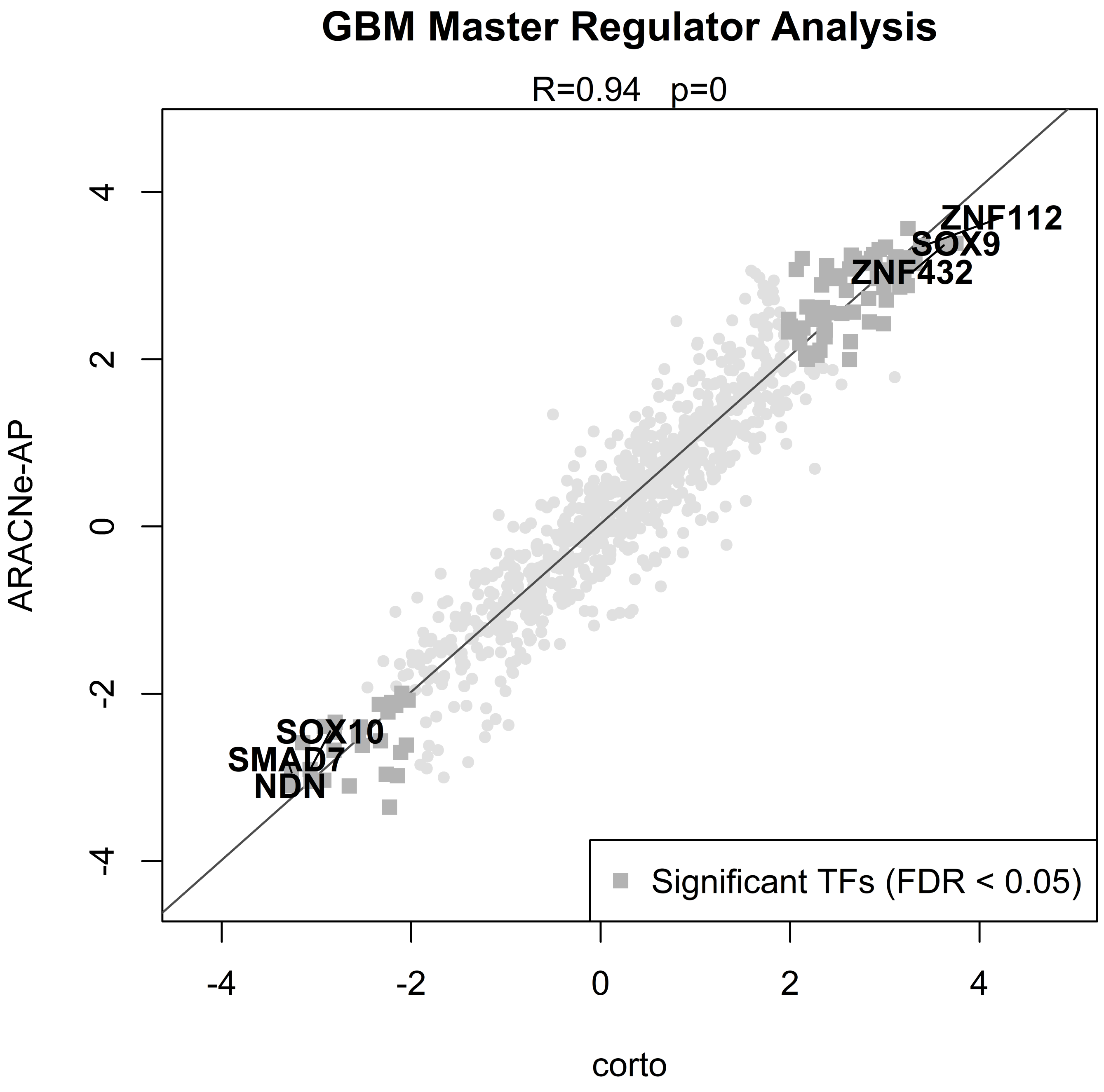
GBM Master Regulator Analysis
R=0.94 p=0
ZNF112
SOX9
ZNF432
SOX10
SMAD7
NDN
Significant TFs (FDR < 0.05)
ARACNe-AP
corto
4
2
0
-2
-4
-4
-2
0
2
4

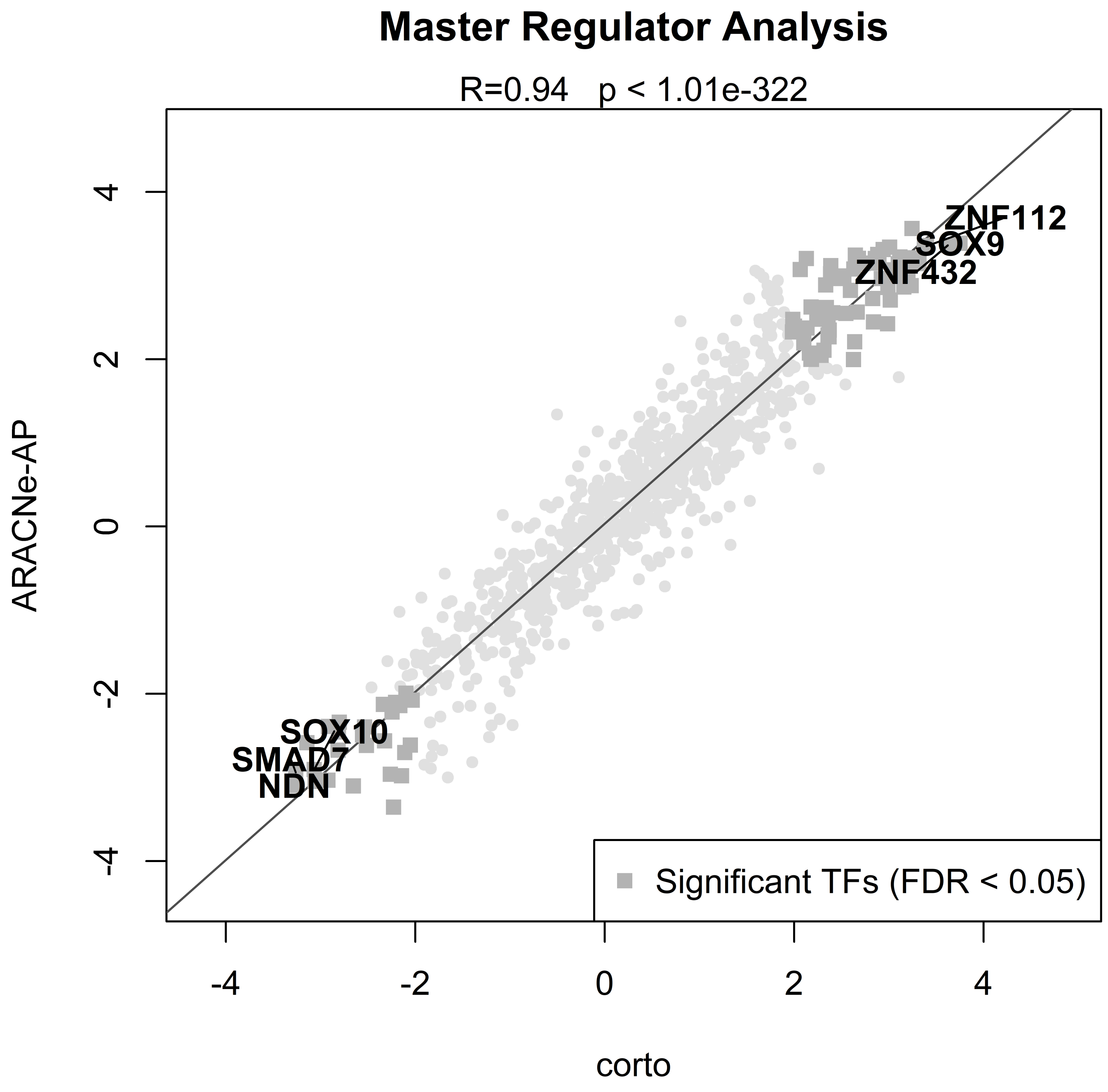

Master Regulator Analysis
R=0.94 p < 1.01e-322
ARACNe-AP
corto
-4
-2
0
2
4
ZNF112
SOX9
ZNF432
SOX10
SMAD7
NDN
Significant TFs (FDR < 0.05)

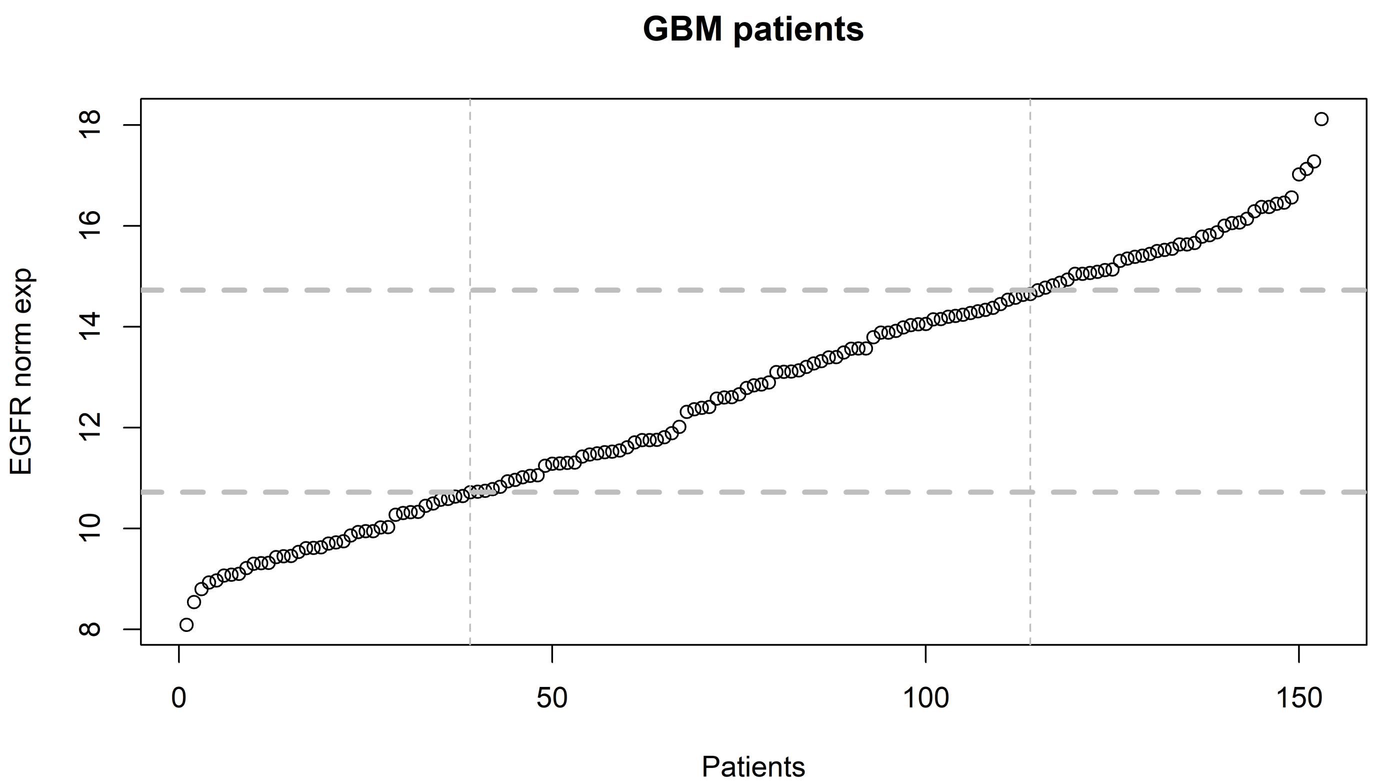
GBM patients
EGFR norm exp
8
10
12
14
16
18
0
50
100
150
Patients

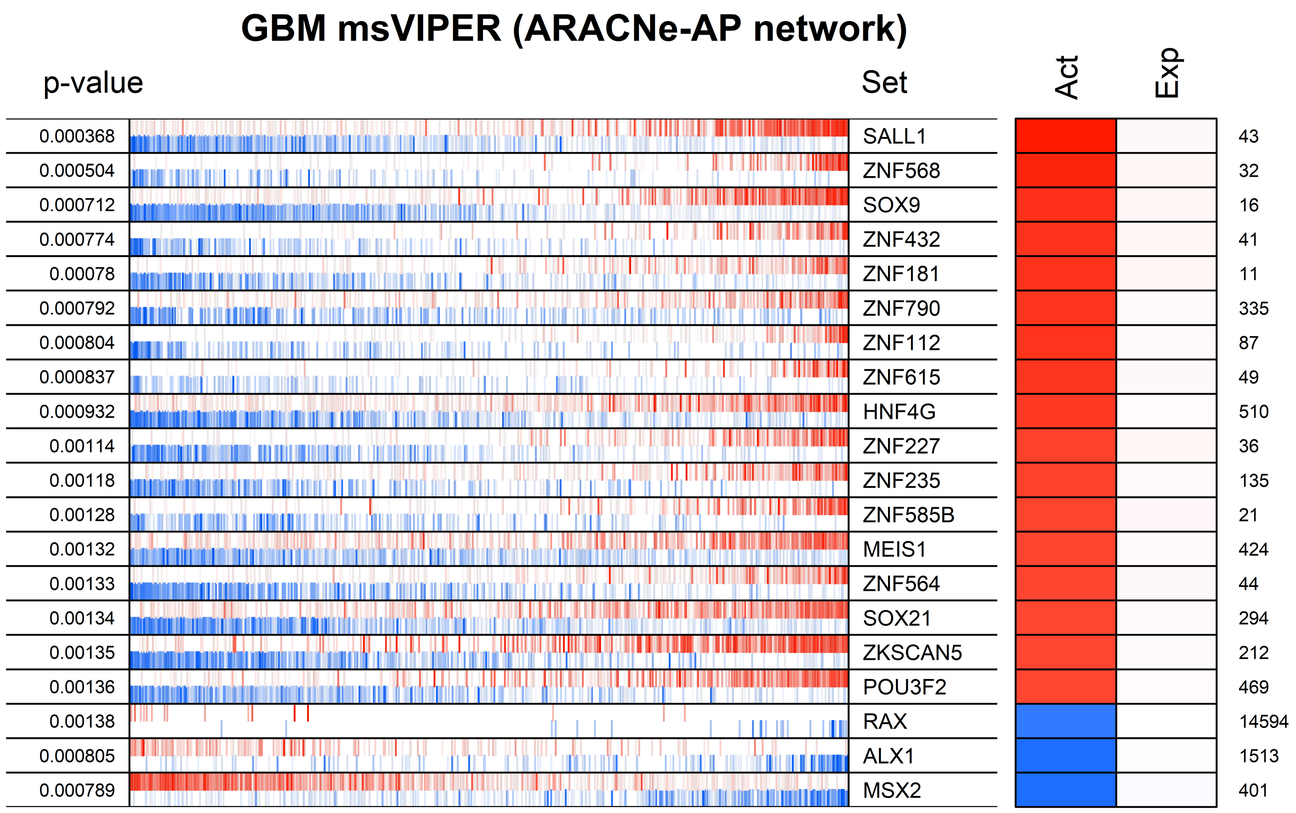
GBM msVIPER (ARACNe-AP network)
p-value
Set
Act
Exp
0.000368 SALL1 43
0.000504 ZNF568 32
0.000712 SOX9 16
0.000774 ZNF432 41
0.00078 ZNF181 11
0.000792 ZNF790 335
0.000804 ZNF112 87
0.000837 ZNF615 49
0.000932 HNF4G 510
0.00114 ZNF227 36
0.00118 ZNF235 135
0.00128 ZNF585B 21
0.00132 MEIS1 424
0.00133 ZNF564 44
0.00134 SOX21 294
0.00135 ZKSCAN5 212
0.00136 POU3F2 469
0.00138 RAX 14594
0.000805 ALX1 1513
0.000789 MSX2 401

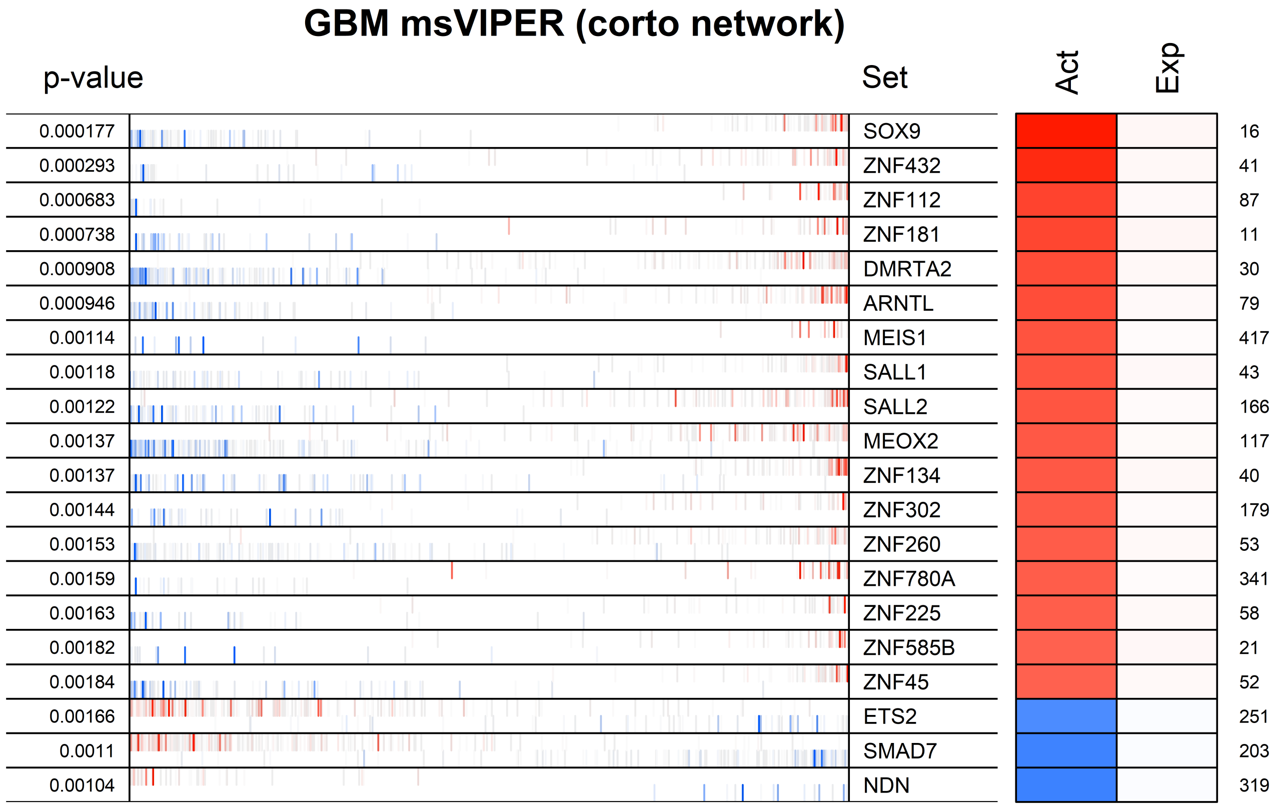
GBM msVIPER (corto network)
p-value
Set
Act
Exp
0.000177 SOX9 16
0.000293 ZNF432 41
0.000683 ZNF112 87
0.000738 ZNF181 11
0.000908 DMRTA2 30
0.000946 ARNTL 79
0.00114 MEIS1 417
0.00118 SALL1 43
0.00122 SALL2 166
0.00137 MEOX2 117
0.00137 ZNF134 40
0.00144 ZNF302 179
0.00153 ZNF260 53
0.00159 ZNF780A 341
0.00163 ZNF225 58
0.00182 ZNF585B 21
0.00184 ZNF45 52
0.00166 ETS2 251
0.0011 SMAD7 203
0.00104 NDN 319

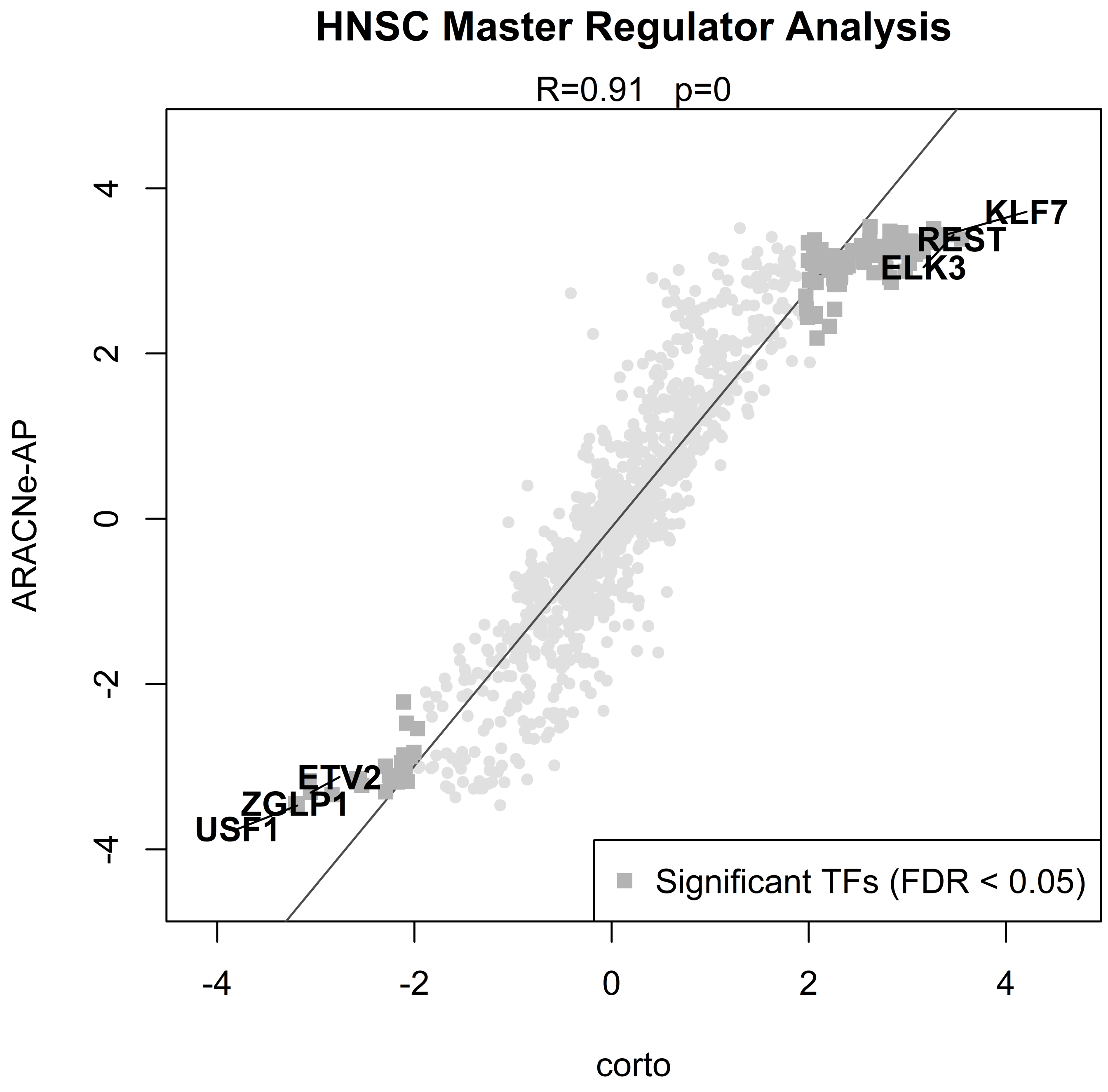

HNSC Master Regulator Analysis
R=0.91 p=0
ARACNe-AP
corto
4
2
0
-2
-4
KLF7
REST
ELK3
ETV2
ZGLP1
USF1
Significant TFs (FDR < 0.05)

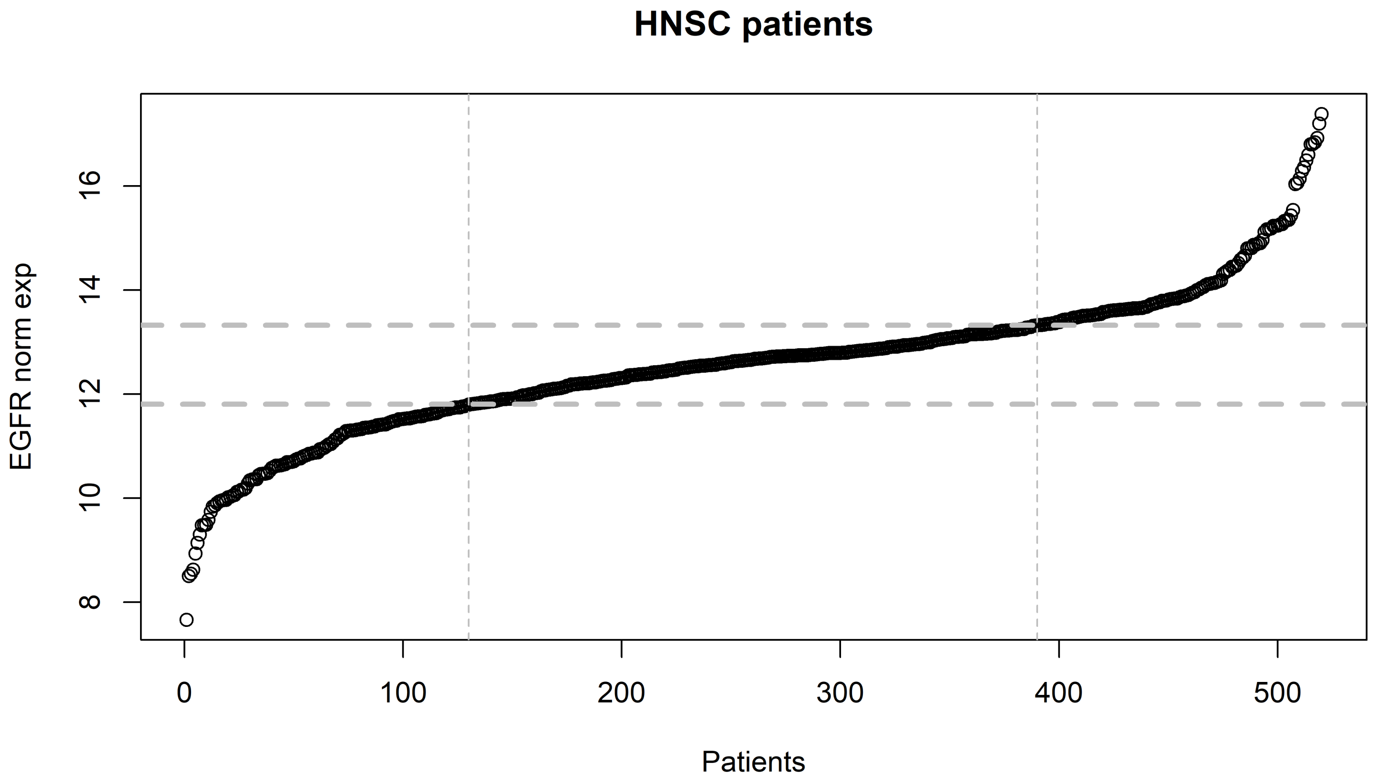
HNSC patients
EGFR norm exp
8
10
12
14
16
0
100
200
300
400
500
Patients

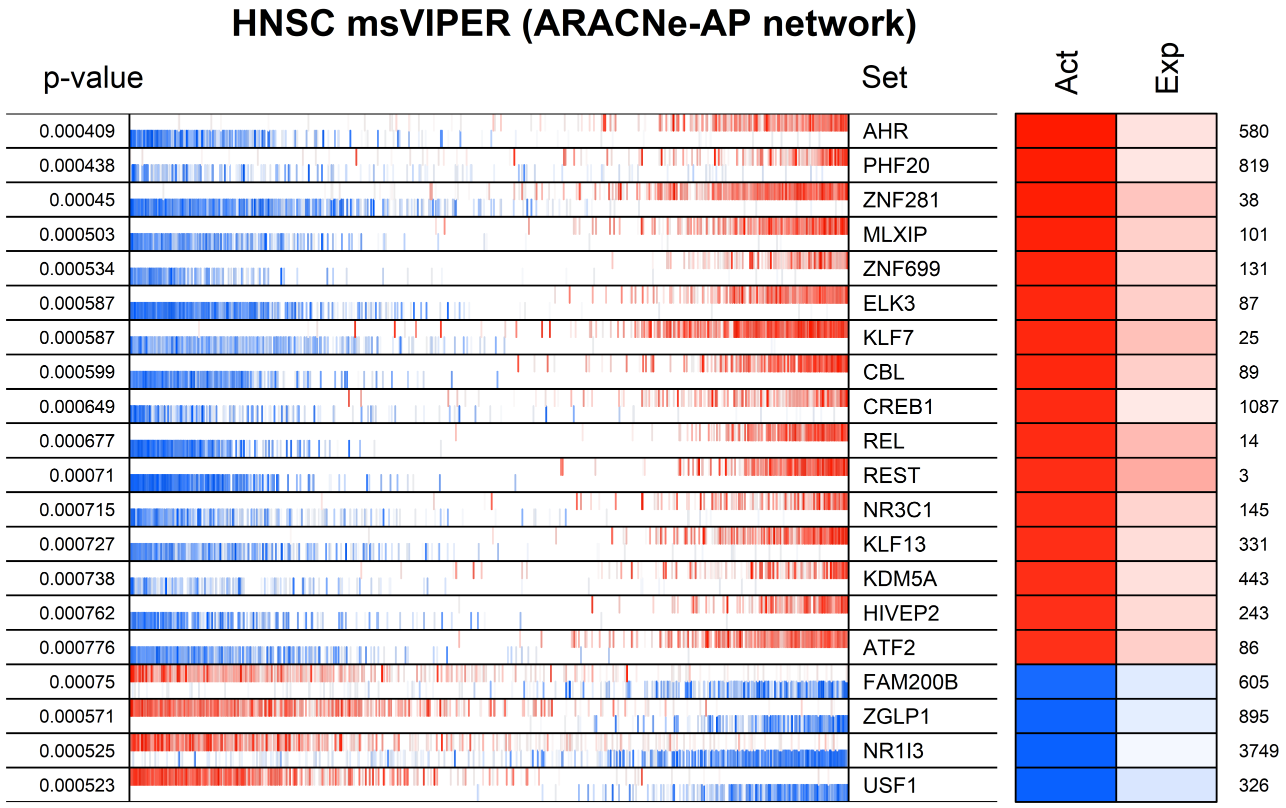
HNSC msVIPER (ARACNe-AP network)
p-value
Set
Act
Exp
0.000409
0.000438
0.00045
0.000503
0.000534
0.000587
0.000587
0.000599
0.000649
0.000677
0.00071
0.000715
0.000727
0.000738
0.000762
0.000776
0.00075
0.000571
0.000525
0.000523
AHR
PHF20
ZNF281
MLXIP
ZNF699
ELK3
KLF7
CBL
CREB1
REL
REST
NR3C1
KLF13
KDM5A
HIVEP2
ATF2
FAM200B
ZGLP1
NR1I3
USF1
580
819
38
101
131
87
25
89
1087
14
3
145
331
443
243
86
605
895
3749
326

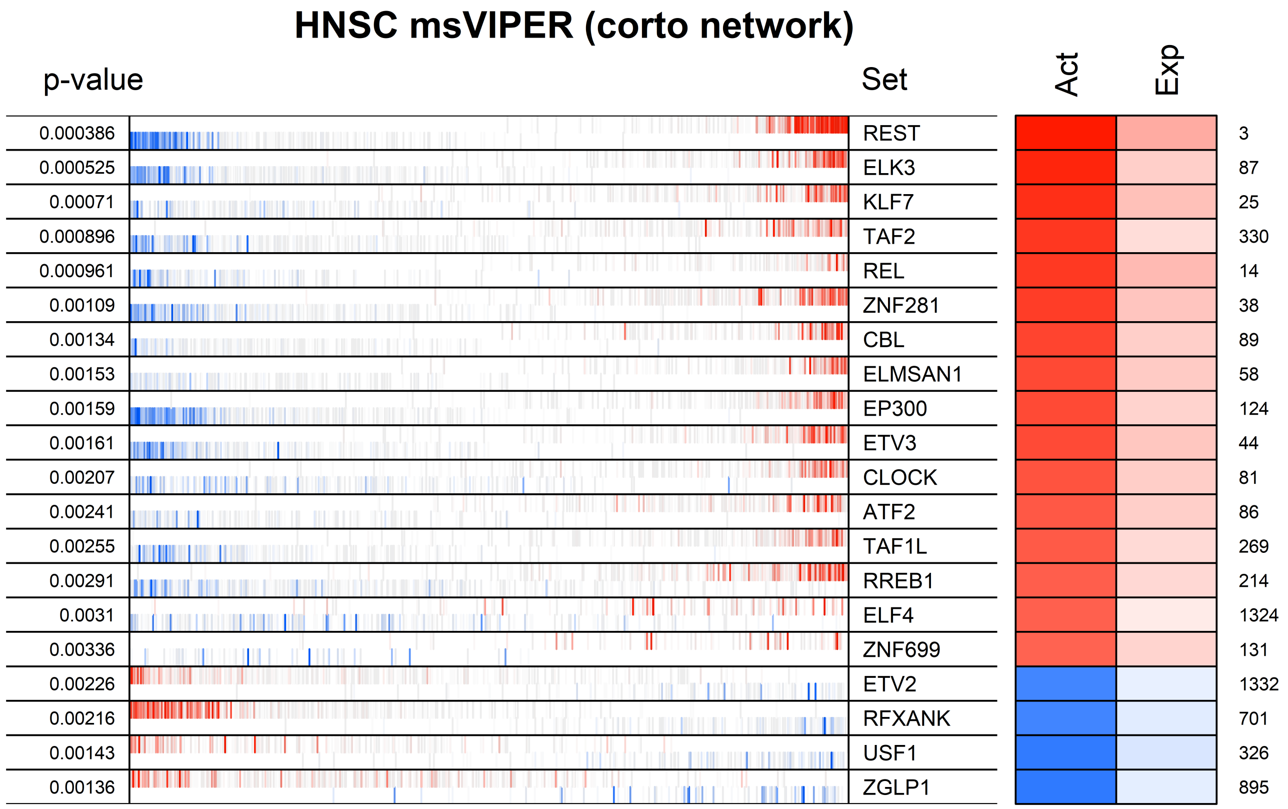
HNSC msVIPER (corto network)
p-value
Set
Act
Exp
0.000386 REST 3
0.000525 ELK3 87
0.00071 KLF7 25
0.000896 TAF2 330
0.000961 REL 14
0.00109 ZNF281 38
0.00134 CBL 89
0.00153 ELMSAN1 58
0.00159 EP300 124
0.00161 ETV3 44
0.00207 CLOCK 81
0.00241 ATF2 86
0.00255 TAF1L 269
0.00291 RREB1 214
0.0031 ELF4 1324
0.00336 ZNF699 131
0.00226 ETV2 1332
0.00216 RFXANK 701
0.00143 USF1 326
0.00136 ZGLP1 895

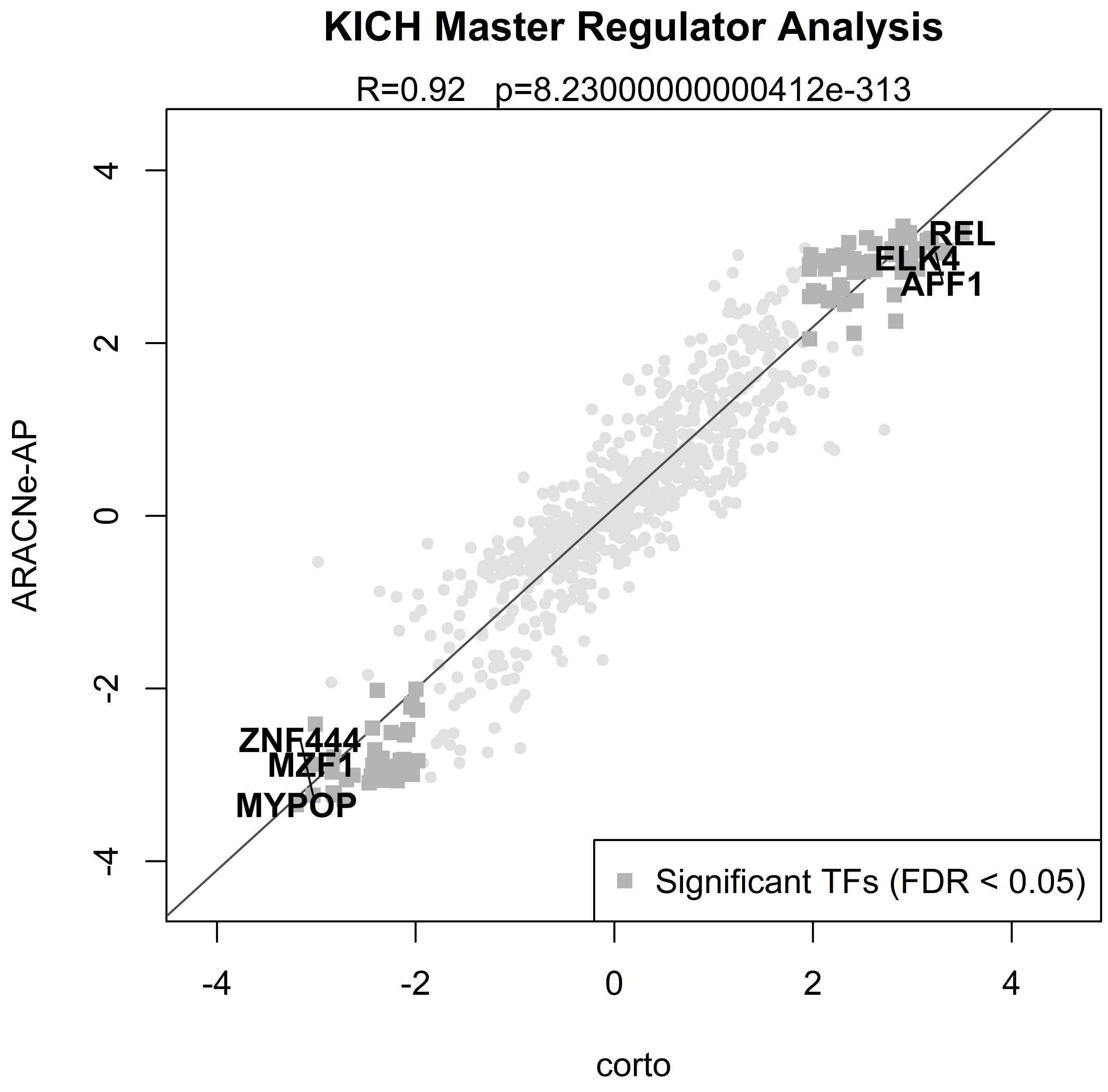
KICH Master Regulator Analysis
R=0.92   p=8.23000000000412e-313
ARACNe-AP
corto
4
2
0
-2
-4
REL
ELK4
AFF1
ZNF444
MZF1
MYPOP
Significant TFs (FDR < 0.05)

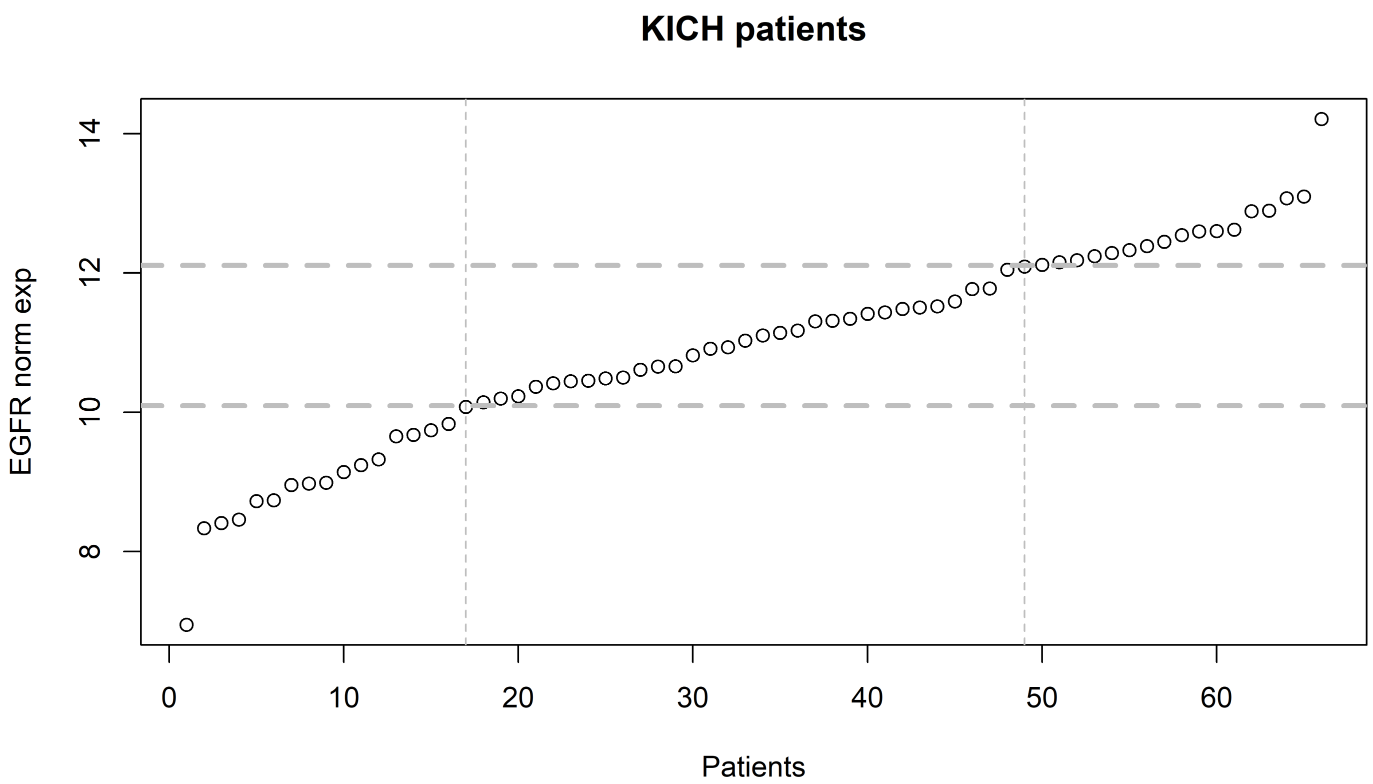
KICH patients
EGFR norm exp
8
10
12
14
0
10
20
30
40
50
60
Patients

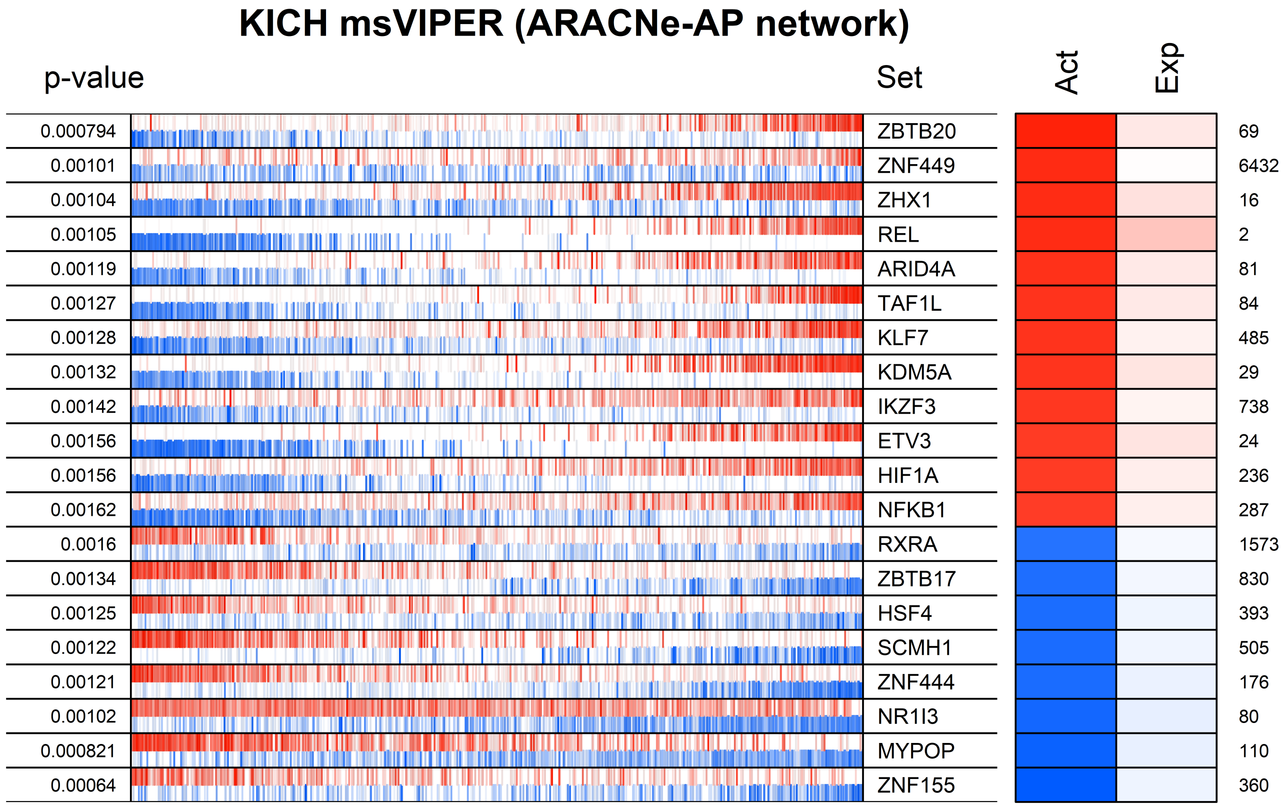

KICH msVIPER (ARACNe-AP network)
p-value
Set
Act
Exp
0.000794
0.00101
0.00104
0.00105
0.00119
0.00127
0.00128
0.00132
0.00142
0.00156
0.00156
0.00162
0.0016
0.00134
0.00125
0.00122
0.00121
0.00102
0.000821
0.00064
ZBTB20
ZNF449
ZHX1
REL
ARID4A
TAF1L
KLF7
KDM5A
IKZF3
ETV3
HIF1A
NFKB1
RXRA
ZBTB17
HSF4
SCMH1
ZNF444
NR1I3
MYPOP
ZNF155
69
6432
16
2
81
84
485
29
738
24
236
287
1573
830
393
505
176
80
110
360

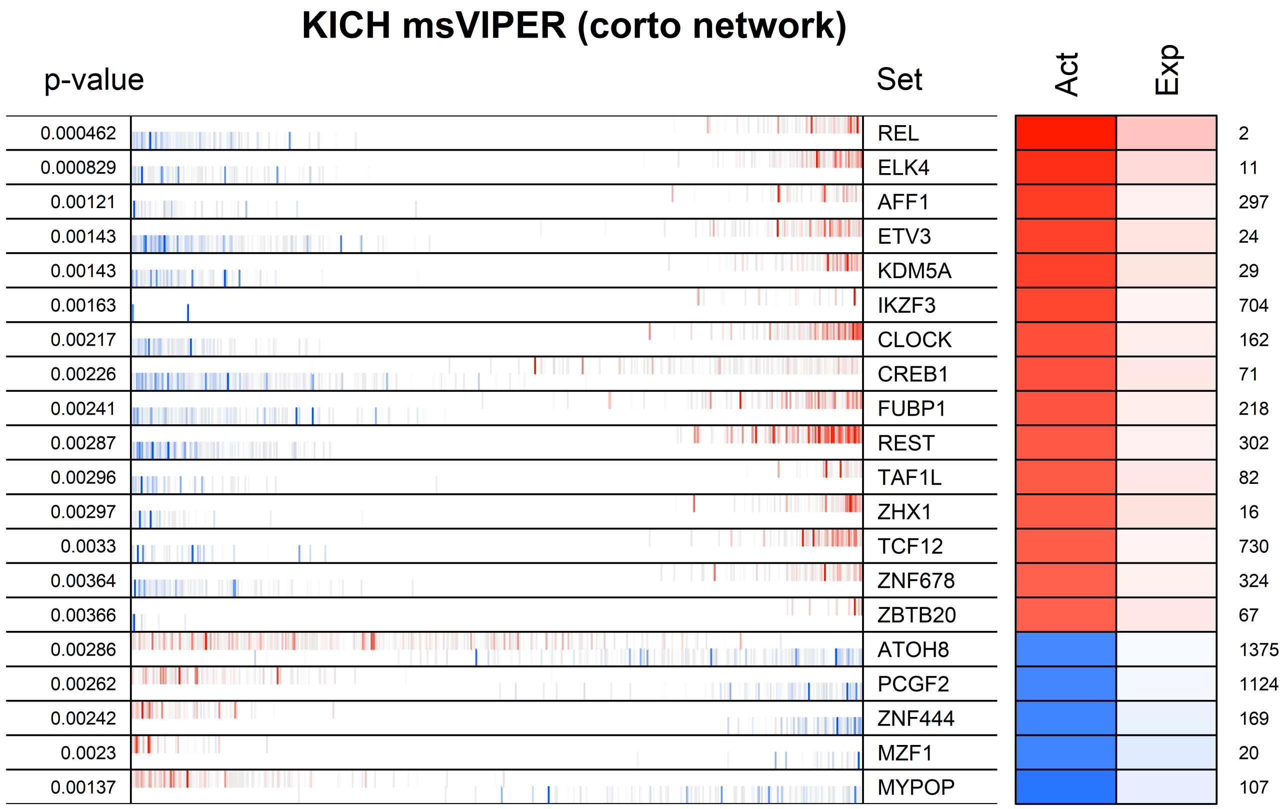
KICH msVIPER (corto network)
p-value
Set
Act
Exp
0.000462 REL 2
0.000829 ELK4 11
0.00121 AFF1 297
0.00143 ETV3 24
0.00143 KDM5A 29
0.00163 IKZF3 704
0.00217 CLOCK 162
0.00226 CREB1 71
0.00241 FUBP1 218
0.00287 REST 302
0.00296 TAF1L 82
0.00297 ZHX1 16
0.0033 TCF12 730
0.00364 ZNF678 324
0.00366 ZBTB20 67
0.00286 ATOH8 1375
0.00262 PCGF2 1124
0.00242 ZNF444 169
0.0023 MZF1 20
0.00137 MYPOP 107

### KIRC Master Regulator Analysis

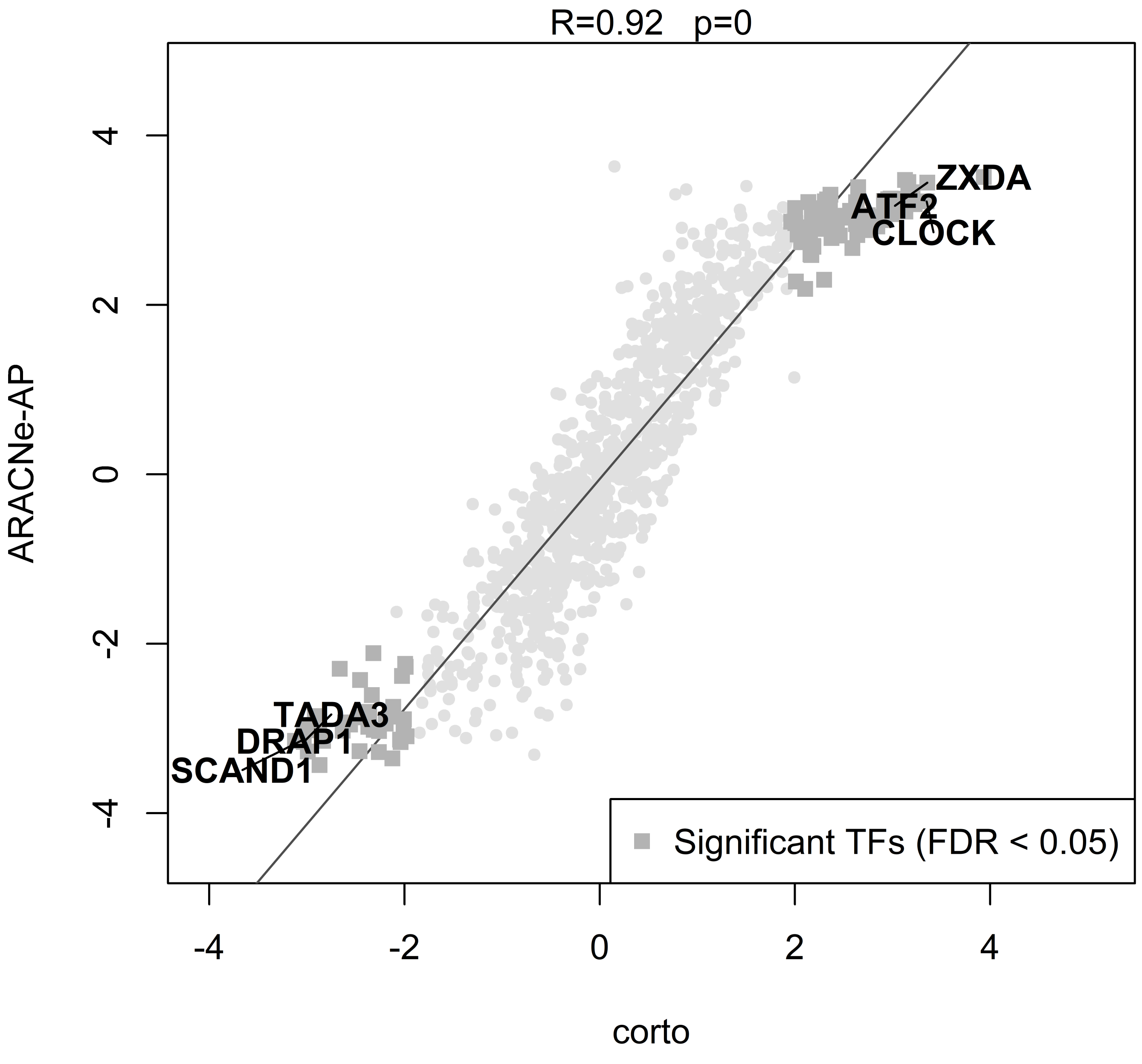

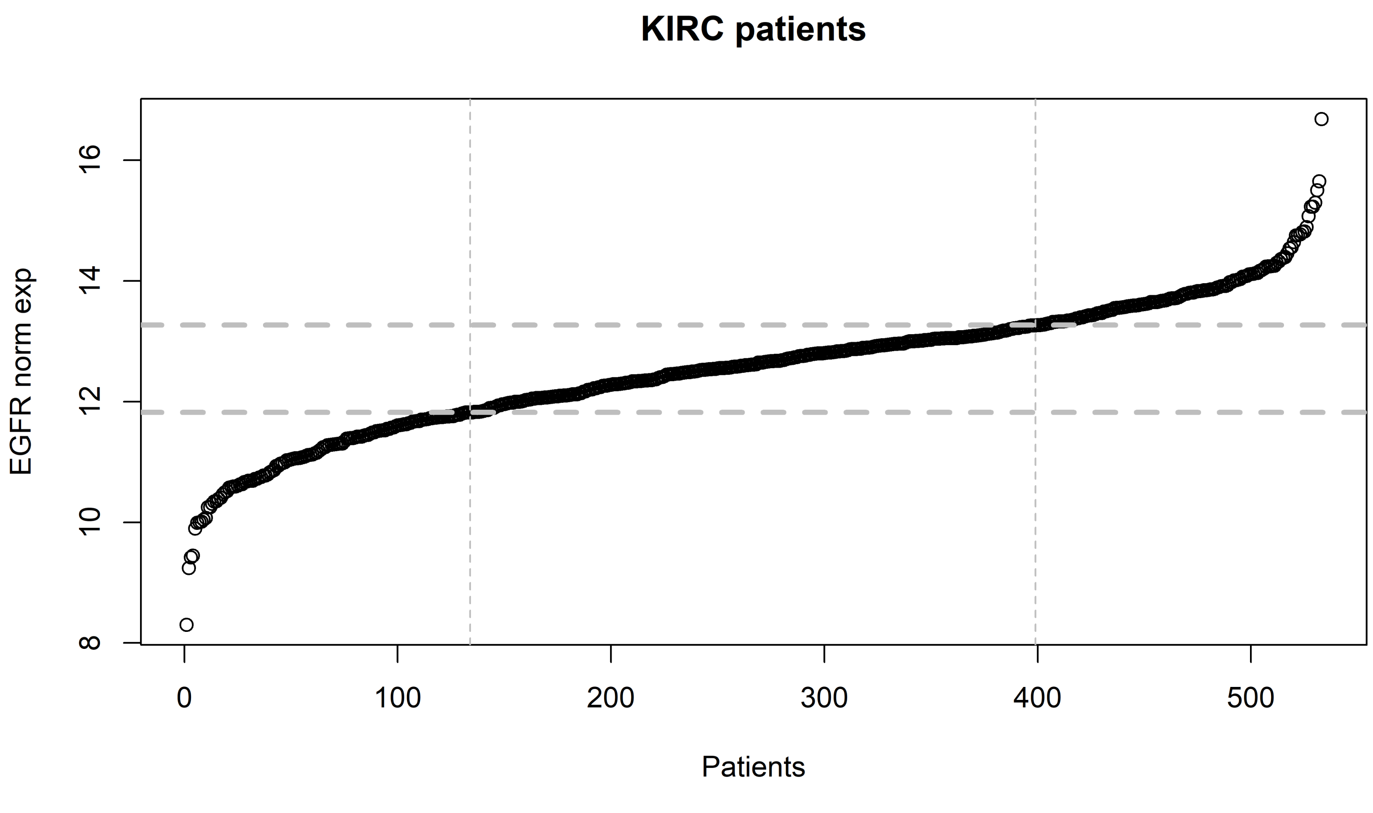

KIRC patients
EGFR norm exp
8
10
12
14
16
0
100
200
300
400
500
Patients

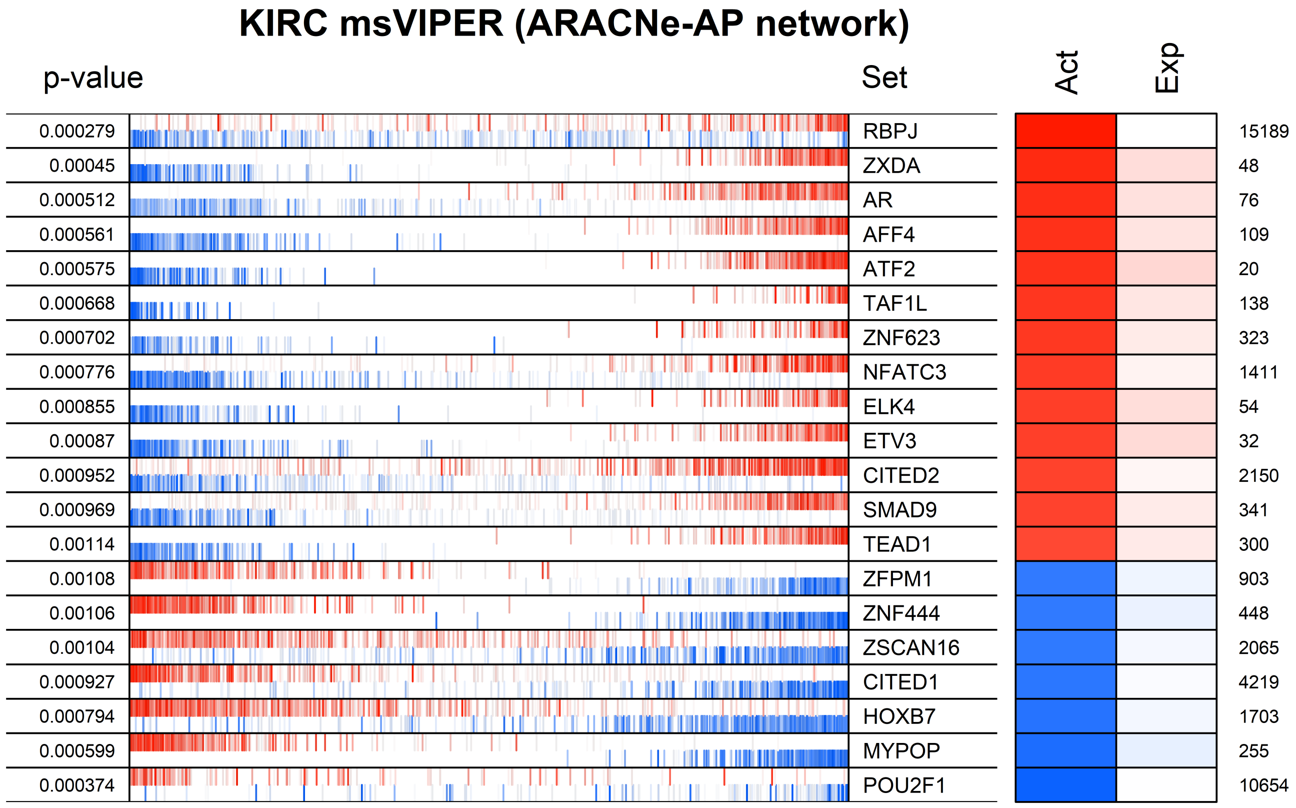

KIRC msVIPER (ARACNe-AP network)
p-value
Set
Act
Exp
0.000279 RBPJ 15189
0.00045 ZXDA 48
0.000512 AR 76
0.000561 AFF4 109
0.000575 ATF2 20
0.000668 TAF1L 138
0.000702 ZNF623 323
0.000776 NFATC3 1411
0.000855 ELK4 54
0.00087 ETV3 32
0.000952 CITED2 2150
0.000969 SMAD9 341
0.00114 TEAD1 300
0.00108 ZFPM1 903
0.00106 ZNF444 448
0.00104 ZSCAN16 2065
0.000927 CITED1 4219
0.000794 HOXB7 1703
0.000599 MYPOP 255
0.000374 POU2F1 10654

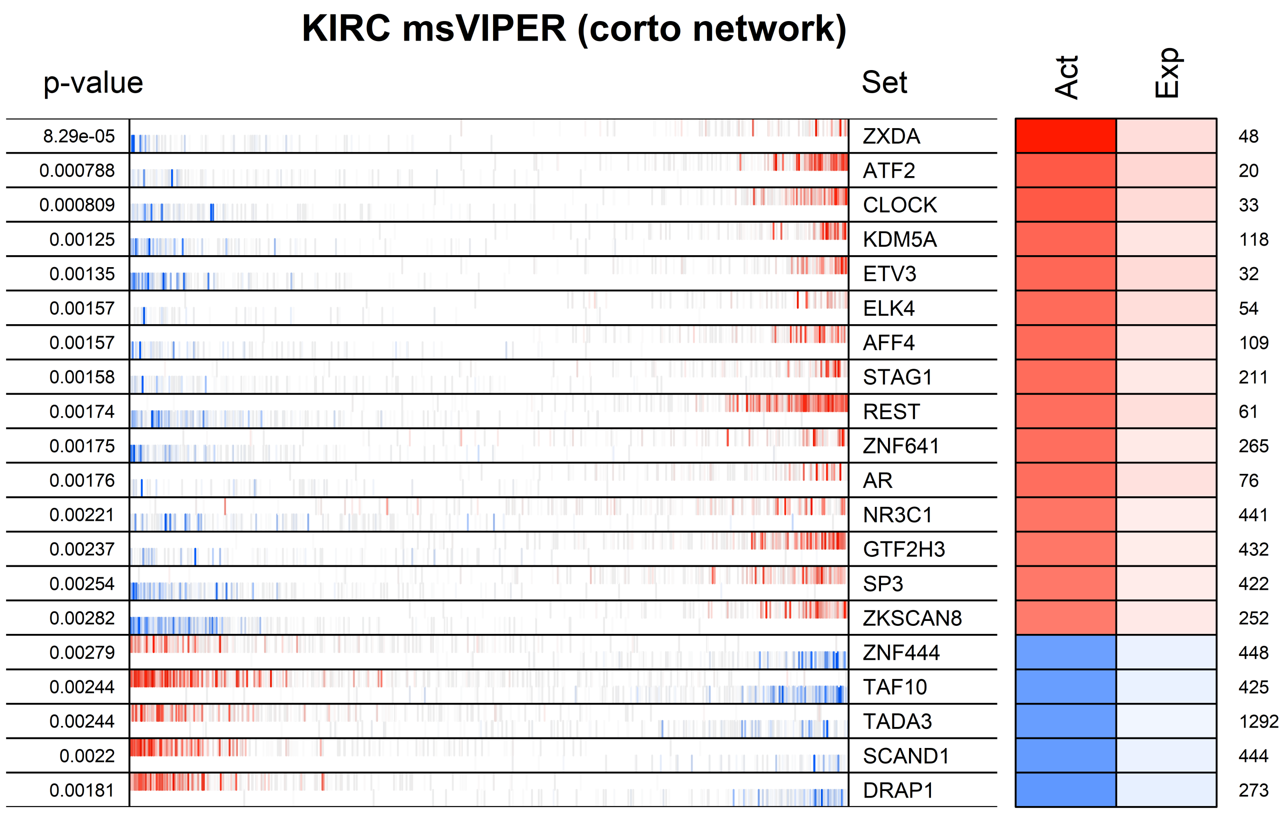
KIRC msVIPER (corto network)
p-value
Set
Act
Exp
8.29e-05
ZXDA
48
0.000788
ATF2
20
0.000809
CLOCK
33
0.00125
KDM5A
118
0.00135
ETV3
32
0.00157
ELK4
54
0.00157
AFF4
109
0.00158
STAG1
211
0.00174
REST
61
0.00175
ZNF641
265
0.00176
AR
76
0.00221
NR3C1
441
0.00237
GTF2H3
432
0.00254
SP3
422
0.00282
ZKSCAN8
252
0.00279
ZNF444
448
0.00244
TAF10
425
0.00244
TADA3
1292
0.0022
SCAND1
444
0.00181
DRAP1
273

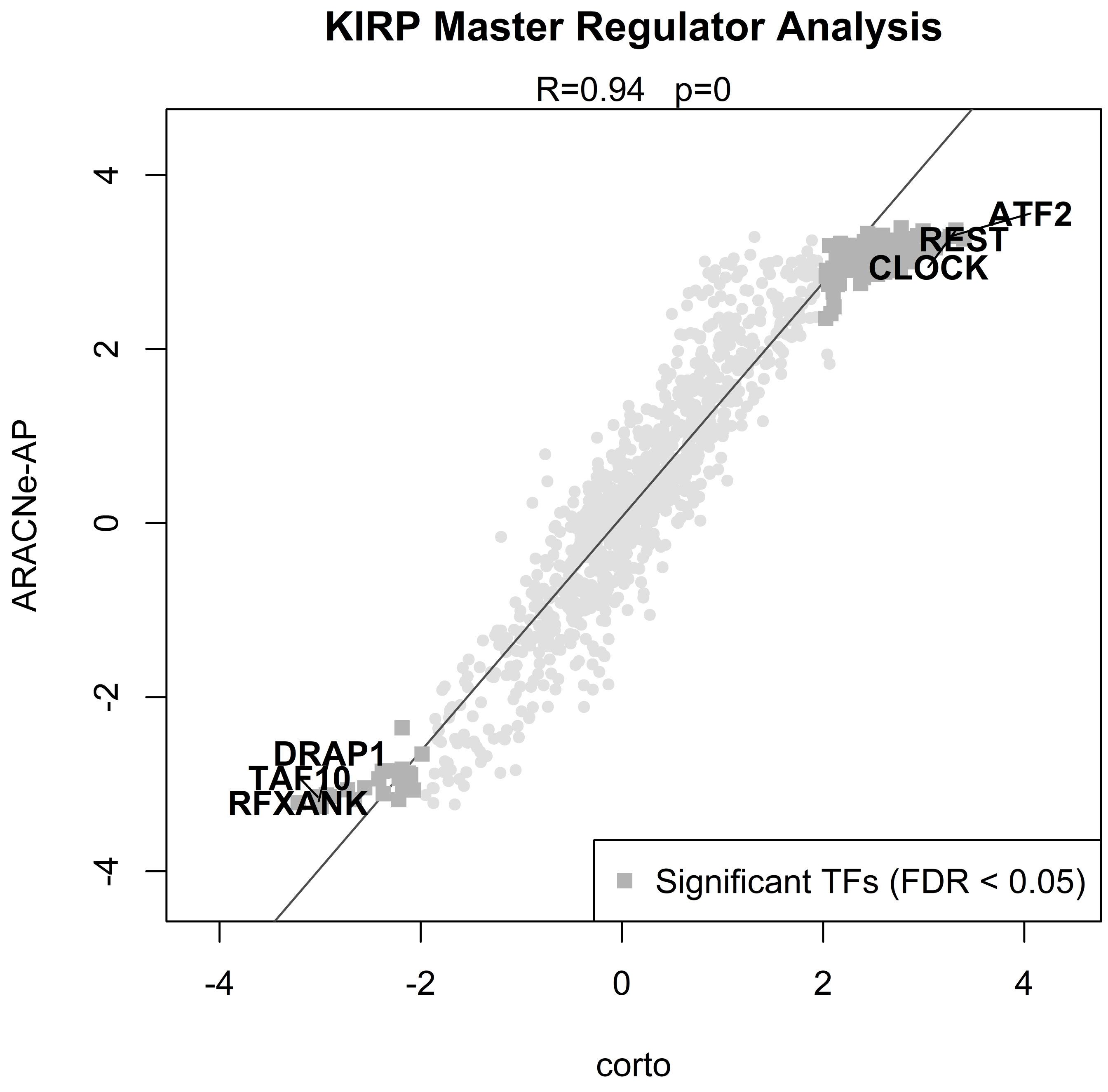
KIRP Master Regulator Analysis
R=0.94 p=0
ATF2
REST
CLOCK
DRAP1
TAF10
RFXANK
Significant TFs (FDR < 0.05)
ARACNe-AP
corto
4
2
0
-2
-4

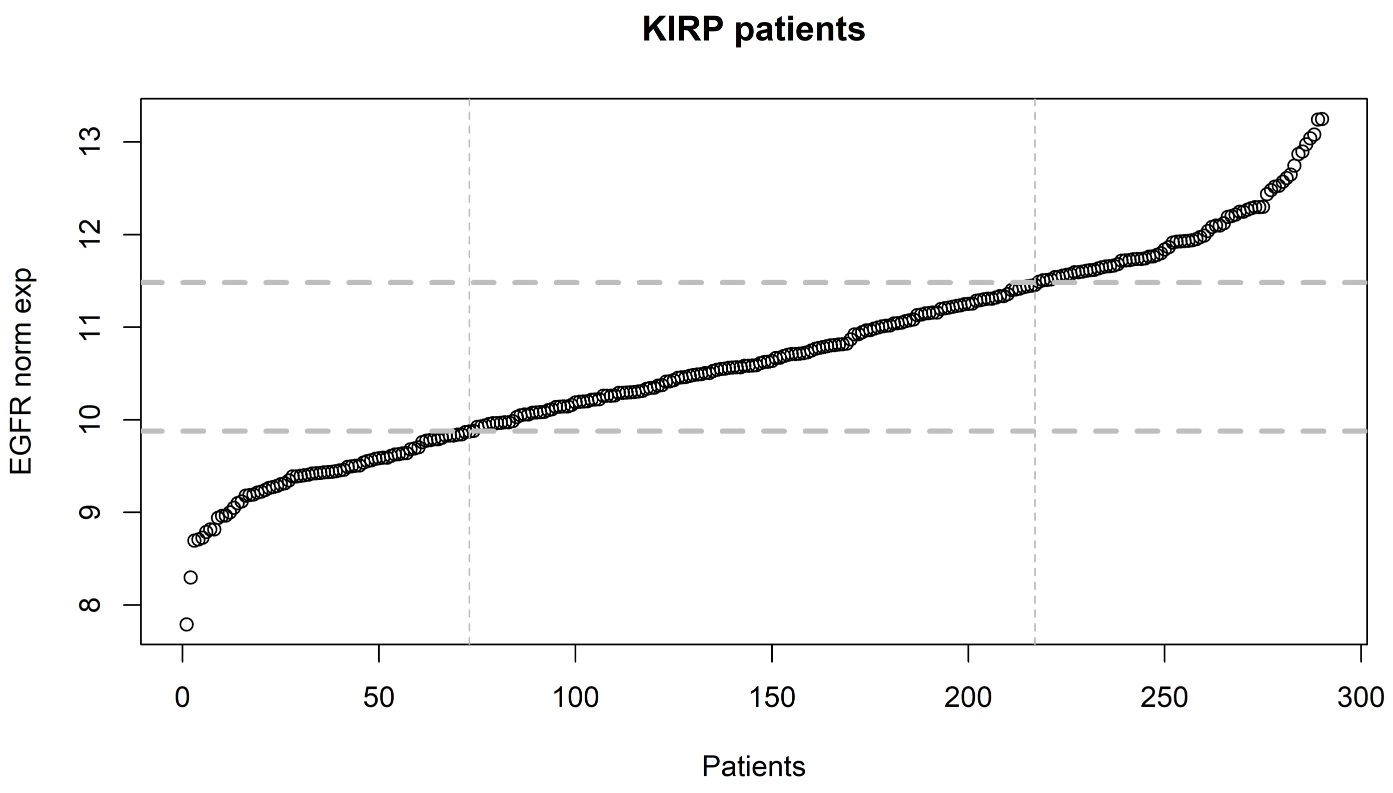

KIRP patients
EGFR norm exp
Patients
8
9
10
11
12
13
0
50
100
150
200
250
300

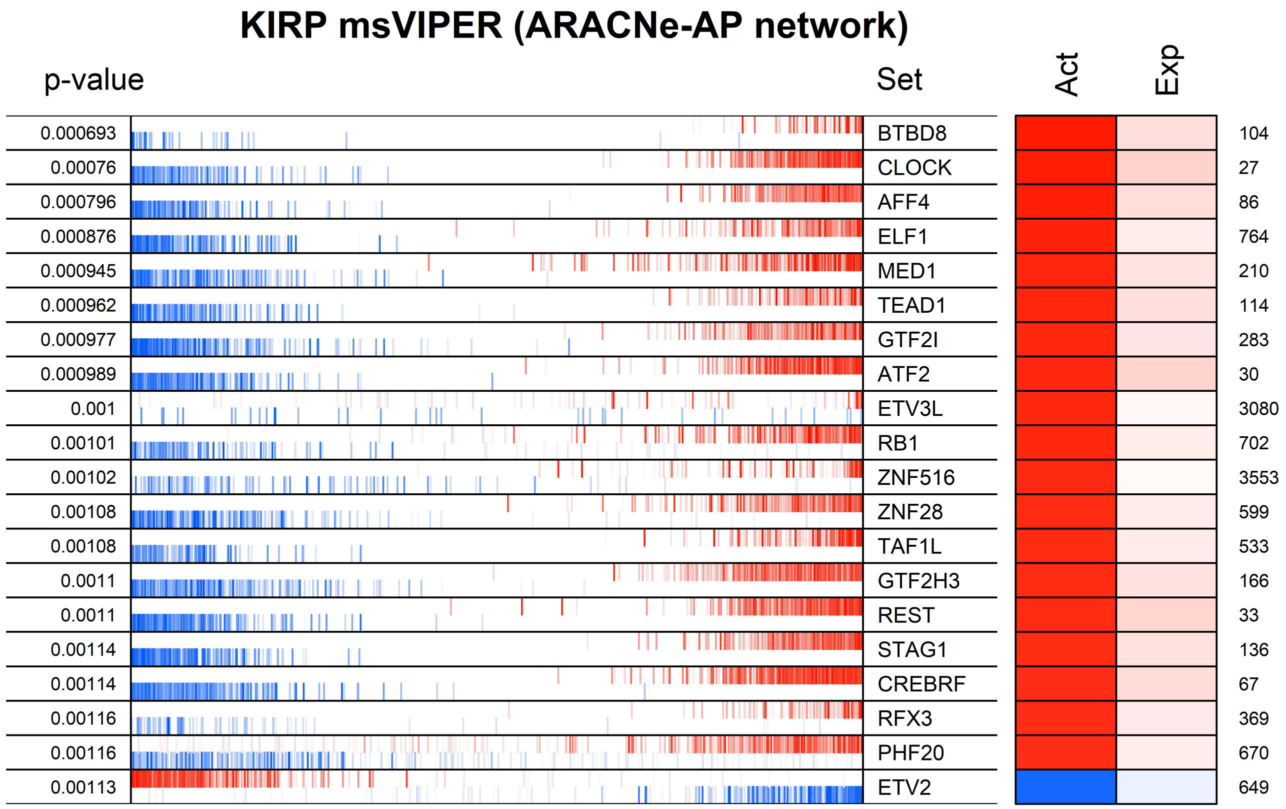

KIRP msVIPER (ARACNe-AP network)
p-value
Set
Act
Exp
0.000693 BTBD8 104
0.00076 CLOCK 27
0.000796 AFF4 86
0.000876 ELF1 764
0.000945 MED1 210
0.000962 TEAD1 114
0.000977 GTF2I 283
0.000989 ATF2 30
0.001 ETV3L 3080
0.00101 RB1 702
0.00102 ZNF516 3553
0.00108 ZNF28 599
0.00108 TAF1L 533
0.0011 GTF2H3 166
0.0011 REST 33
0.00114 STAG1 136
0.00114 CREBRF 67
0.00116 RFX3 369
0.00116 PHF20 670
0.00113 ETV2 649

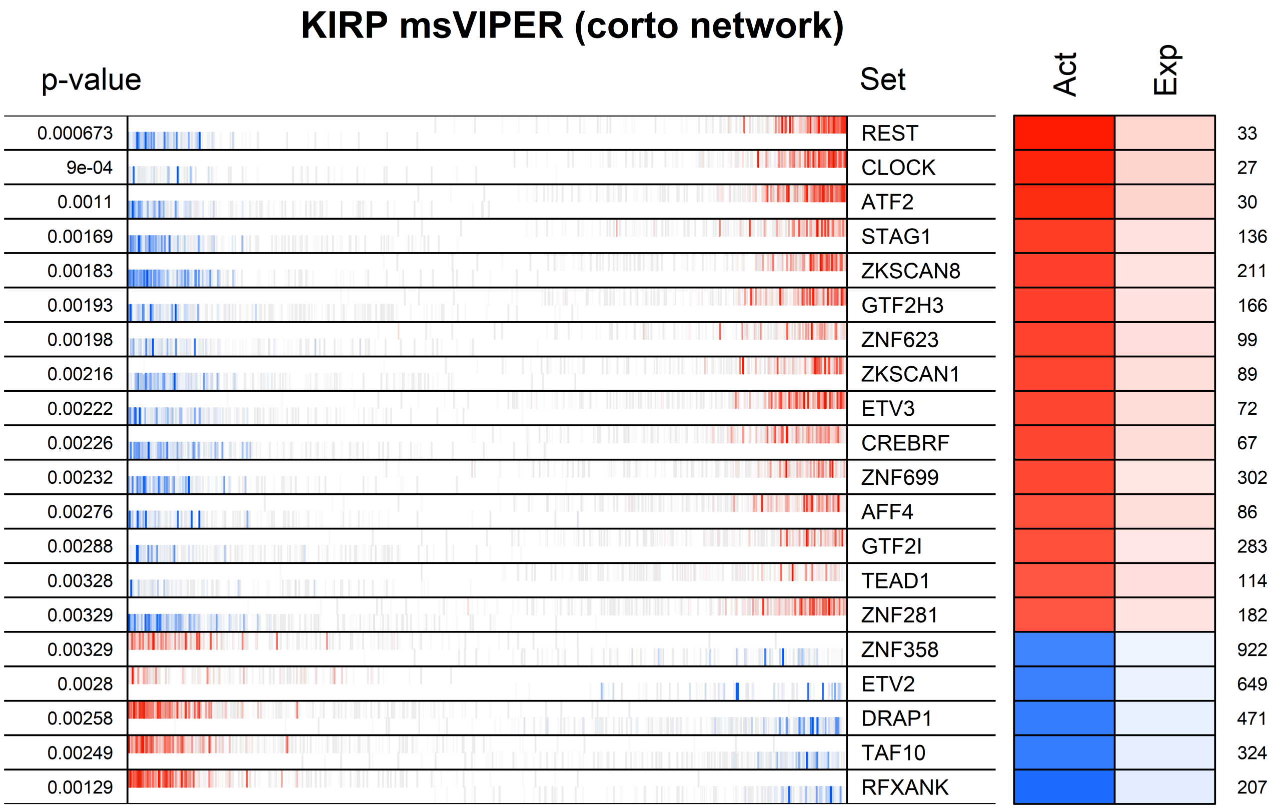
KIRP msVIPER (corto network)
p-value
Set
Act
Exp
0.000673
9e-04
0.0011
0.00169
0.00183
0.00193
0.00198
0.00216
0.00222
0.00226
0.00232
0.00276
0.00288
0.00328
0.00329
0.00329
0.0028
0.00258
0.00249
0.00129
REST
CLOCK
ATF2
STAG1
ZKSCAN8
GTF2H3
ZNF623
ZKSCAN1
ETV3
CREBRF
ZNF699
AFF4
GTF2I
TEAD1
ZNF281
ZNF358
ETV2
DRAP1
TAF10
RFXANK
33
27
30
136
211
166
99
89
72
67
302
86
283
114
182
922
649
471
324
207

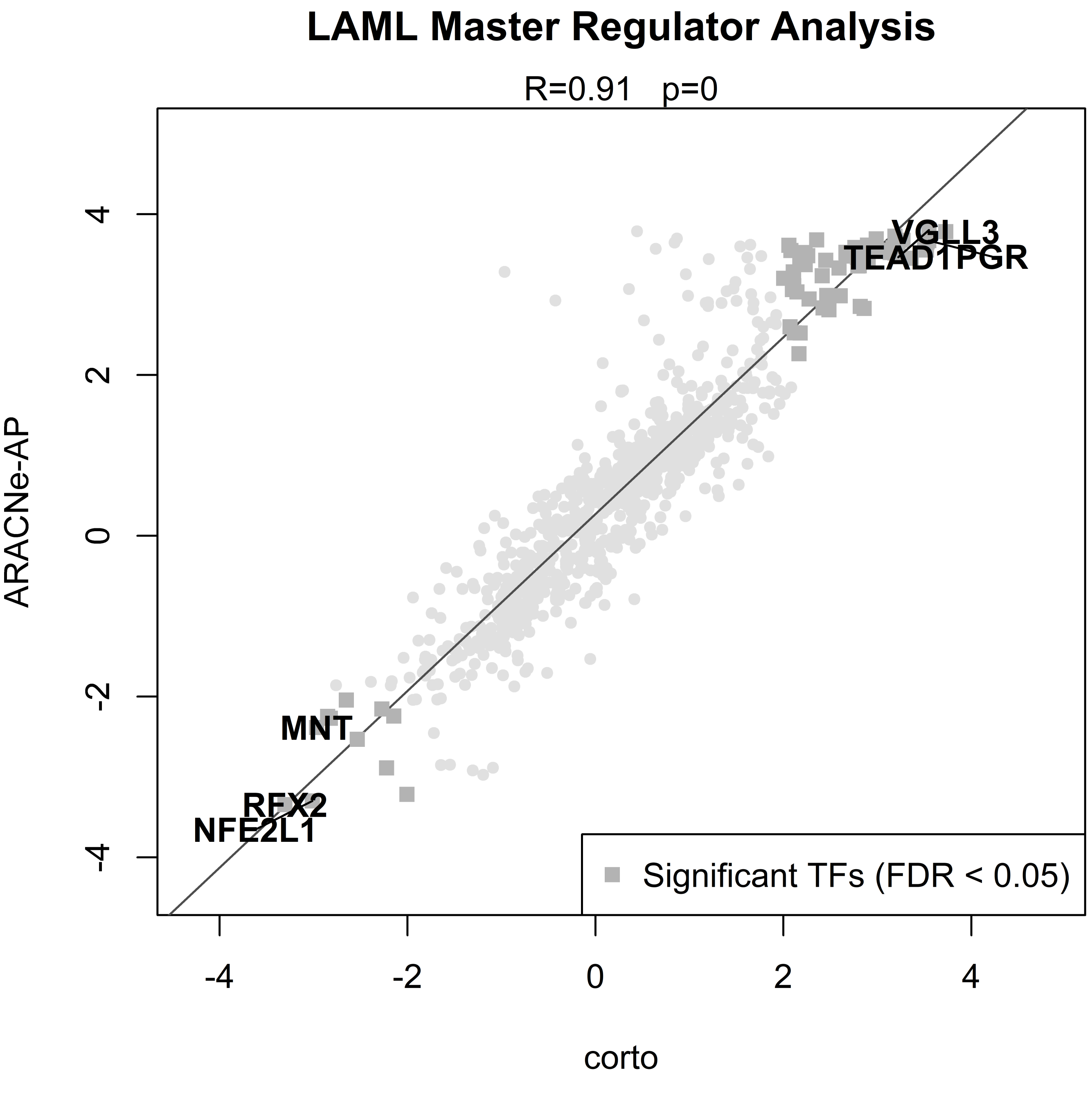
LAML Master Regulator Analysis
R=0.91 p=0
ARACNe-AP
corto
VGLL3
TEAD1PGR
MNT
RFX2
NFE2L1
Significant TFs (FDR < 0.05)
-4
-2
0
2
4

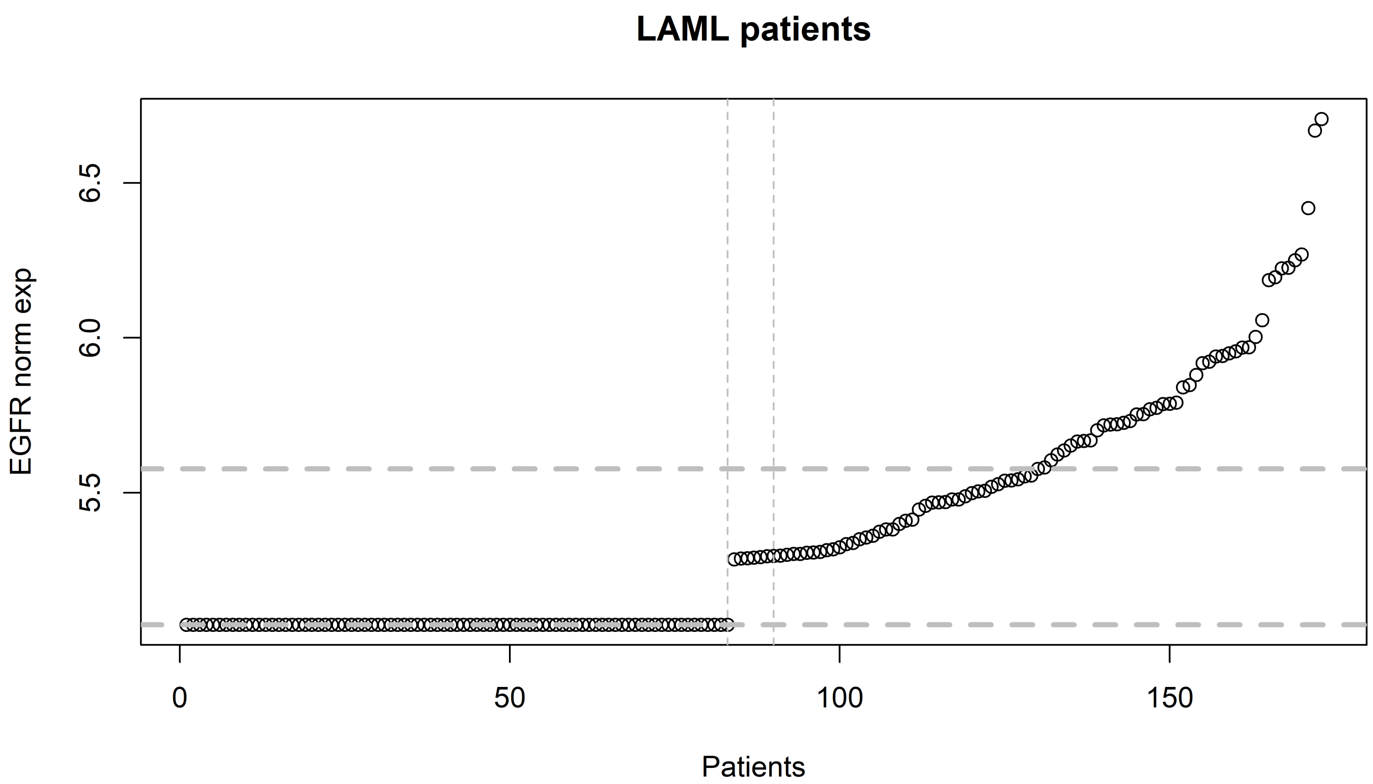

LAML patients
EGFR norm exp
6.5
6.0
5.5
0
50
100
150
Patients

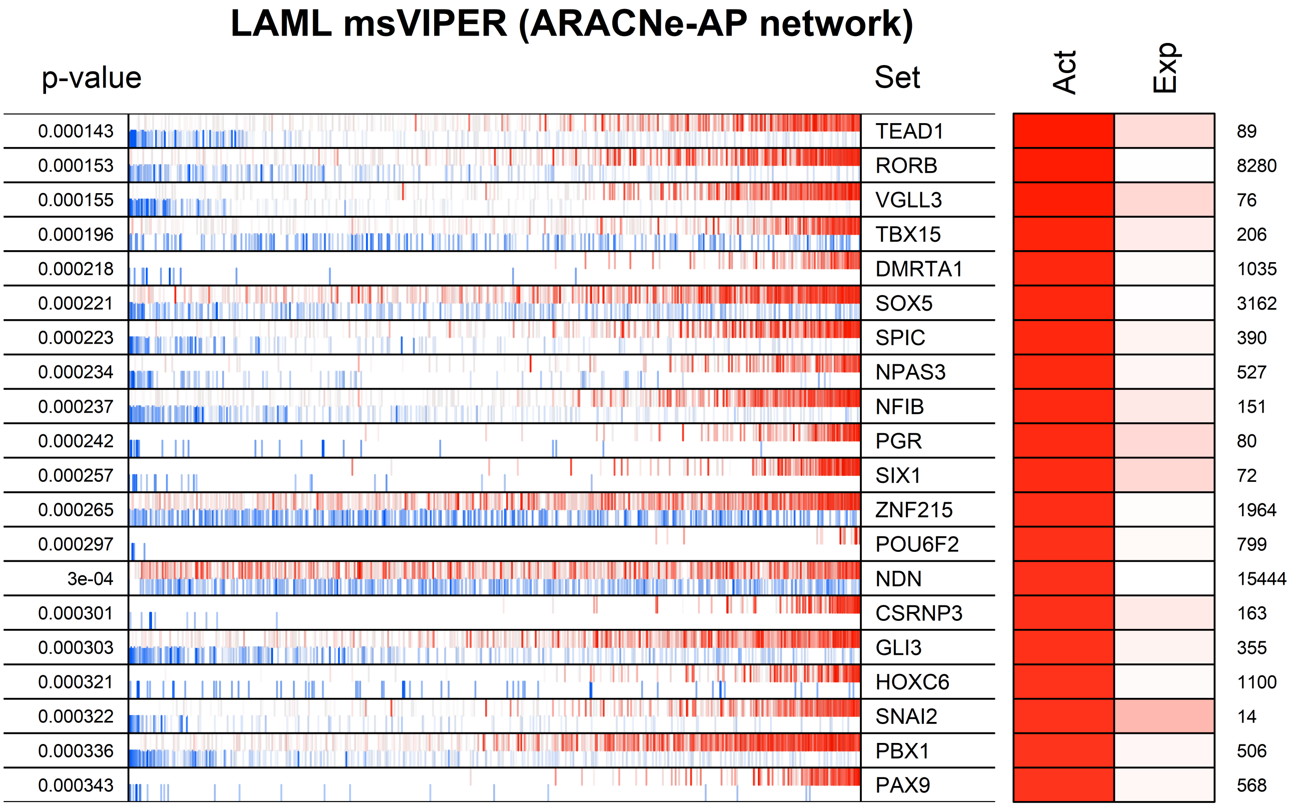
LAML msVIPER (ARACNe-AP network)
p-value
Set
Act
Exp
0.000143 TEAD1 89
0.000153 RORB 8280
0.000155 VGLL3 76
0.000196 TBX15 206
0.000218 DMRTA1 1035
0.000221 SOX5 3162
0.000223 SPIC 390
0.000234 NPAS3 527
0.000237 NFIB 151
0.000242 PGR 80
0.000257 SIX1 72
0.000265 ZNF215 1964
0.000297 POU6F2 799
3e-04 NDN 15444
0.000301 CSRNP3 163
0.000303 GLI3 355
0.000321 HOXC6 1100
0.000322 SNAI2 14
0.000336 PBX1 506
0.000343 PAX9 568

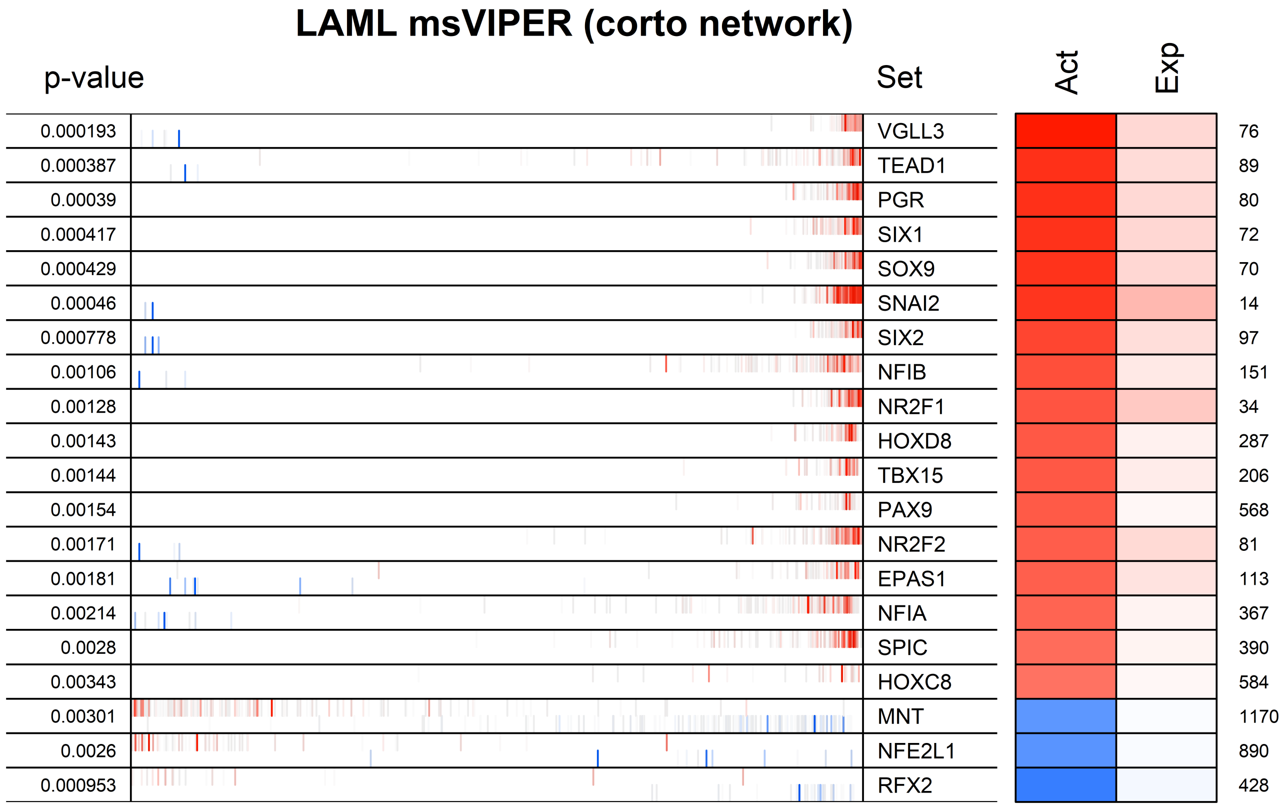

LAML msVIPER (corto network)
p-value
Set
Act
Exp
0.000193
0.000387
0.00039
0.000417
0.000429
0.00046
0.000778
0.00106
0.00128
0.00143
0.00144
0.00154
0.00171
0.00181
0.00214
0.0028
0.00343
0.00301
0.0026
0.000953
VGLL3
TEAD1
PGR
SIX1
SOX9
SNAI2
SIX2
NFIB
NR2F1
HOXD8
TBX15
PAX9
NR2F2
EPAS1
NFIA
SPIC
HOXC8
MNT
NFE2L1
RFX2
76
89
80
72
70
14
97
151
34
287
206
568
81
113
367
390
584
1170
890
428

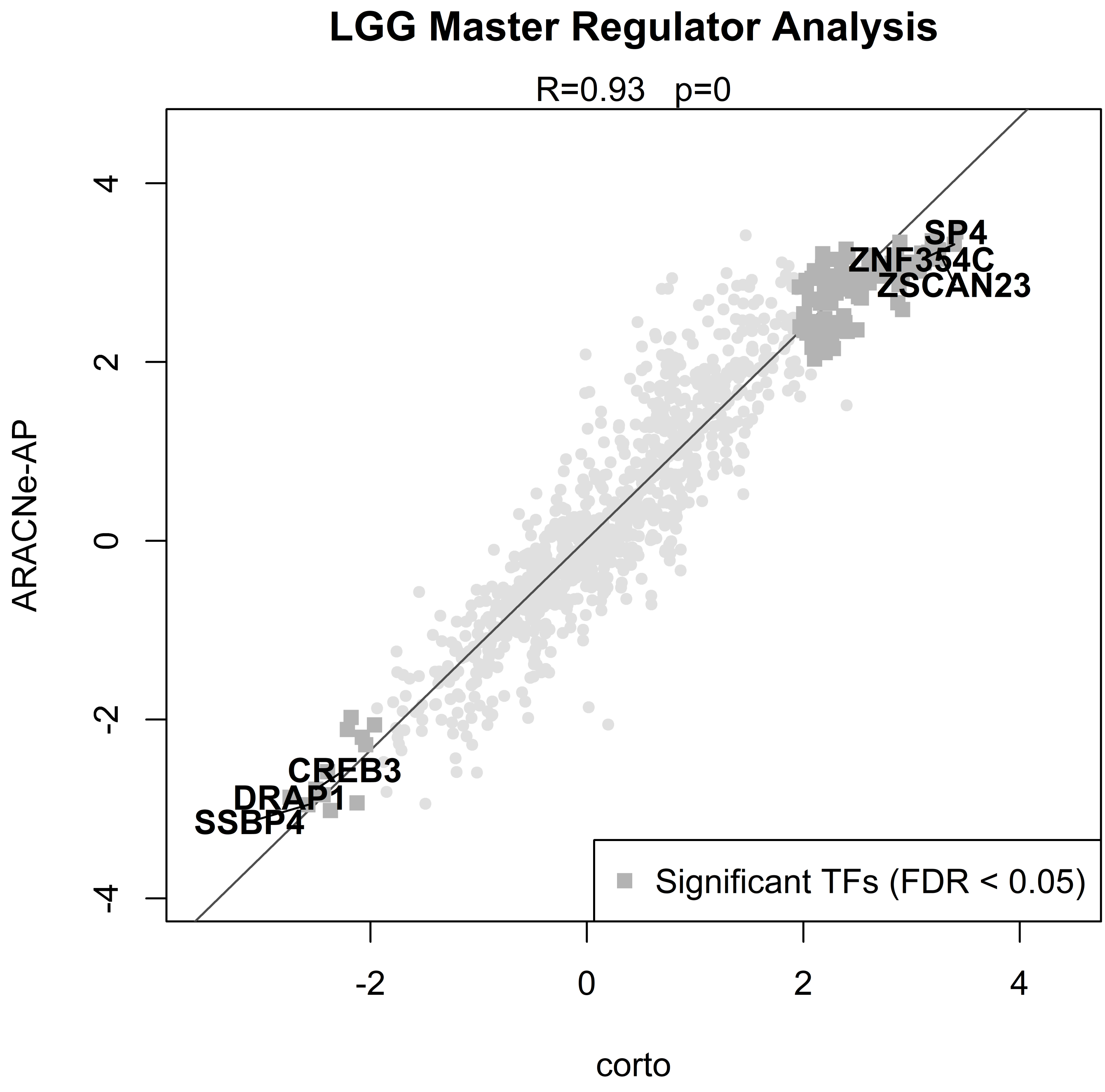

LGG Master Regulator Analysis
R=0.93 p=0
SP4
ZNF354C
ZSCAN23
CREB3
DRAP1
SSBP4
Significant TFs (FDR < 0.05)
ARACNe-AP
corto
4
2
0
-2
-4
-2
0
2
4

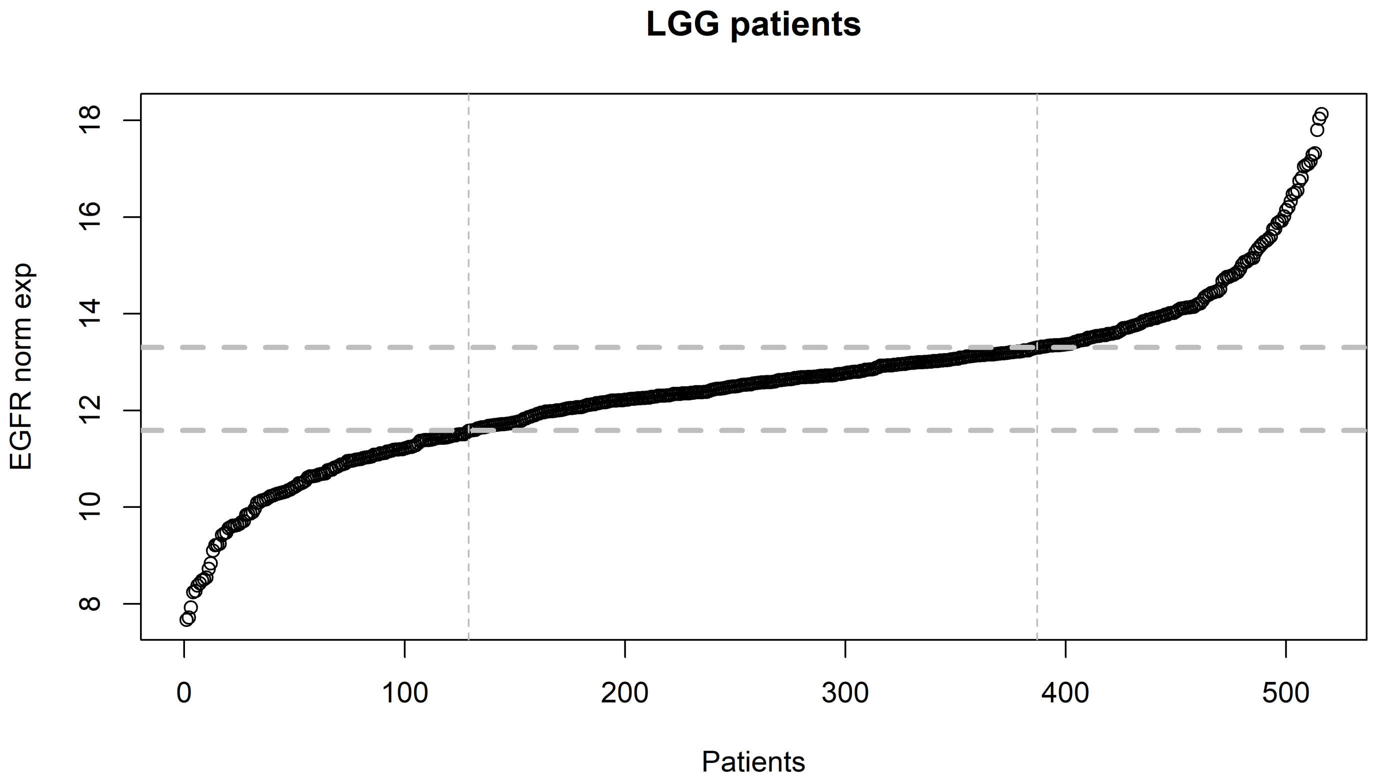

LGG patients
EGFR norm exp
Patients
8
10
12
14
16
18
0
100
200
300
400
500

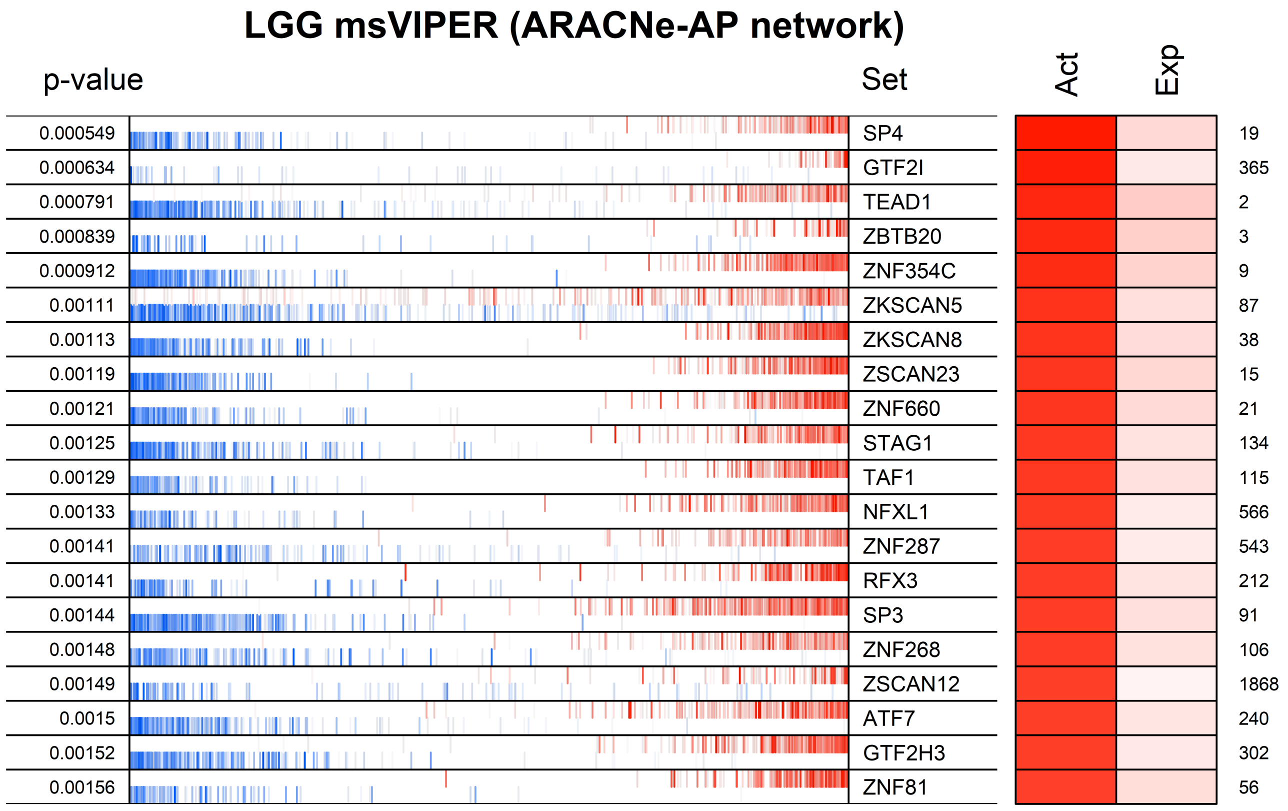
LGG msVIPER (ARACNe-AP network)
p-value
Set
Act
Exp
0.000549
SP4
19
0.000634
GTF2I
365
0.000791
TEAD1
2
0.000839
ZBTB20
3
0.000912
ZNF354C
9
0.00111
ZKSCAN5
87
0.00113
ZKSCAN8
38
0.00119
ZSCAN23
15
0.00121
ZNF660
21
0.00125
STAG1
134
0.00129
TAF1
115
0.00133
NFXL1
566
0.00141
ZNF287
543
0.00141
RFX3
212
0.00144
SP3
91
0.00148
ZNF268
106
0.00149
ZSCAN12
1868
0.0015
ATF7
240
0.00152
GTF2H3
302
0.00156
ZNF81
56

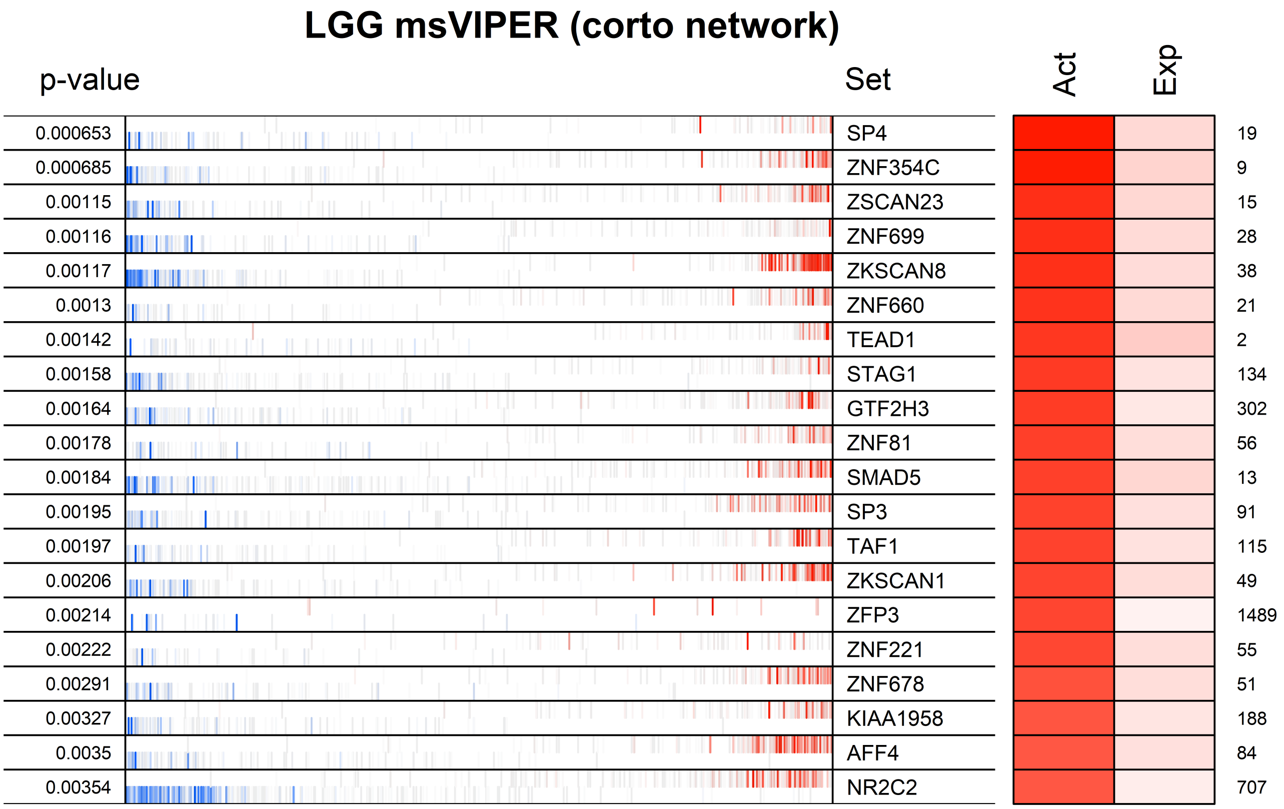
LGG msVIPER (corto network)
p-value
Set
Act
Exp
0.000653 SP4 19
0.000685 ZNF354C 9
0.00115 ZSCAN23 15
0.00116 ZNF699 28
0.00117 ZKSCAN8 38
0.0013 ZNF660 21
0.00142 TEAD1 2
0.00158 STAG1 134
0.00164 GTF2H3 302
0.00178 ZNF81 56
0.00184 SMAD5 13
0.00195 SP3 91
0.00197 TAF1 115
0.00206 ZKSCAN1 49
0.00214 ZFP3 1489
0.00222 ZNF221 55
0.00291 ZNF678 51
0.00327 KIAA1958 188
0.0035 AFF4 84
0.00354 NR2C2 707

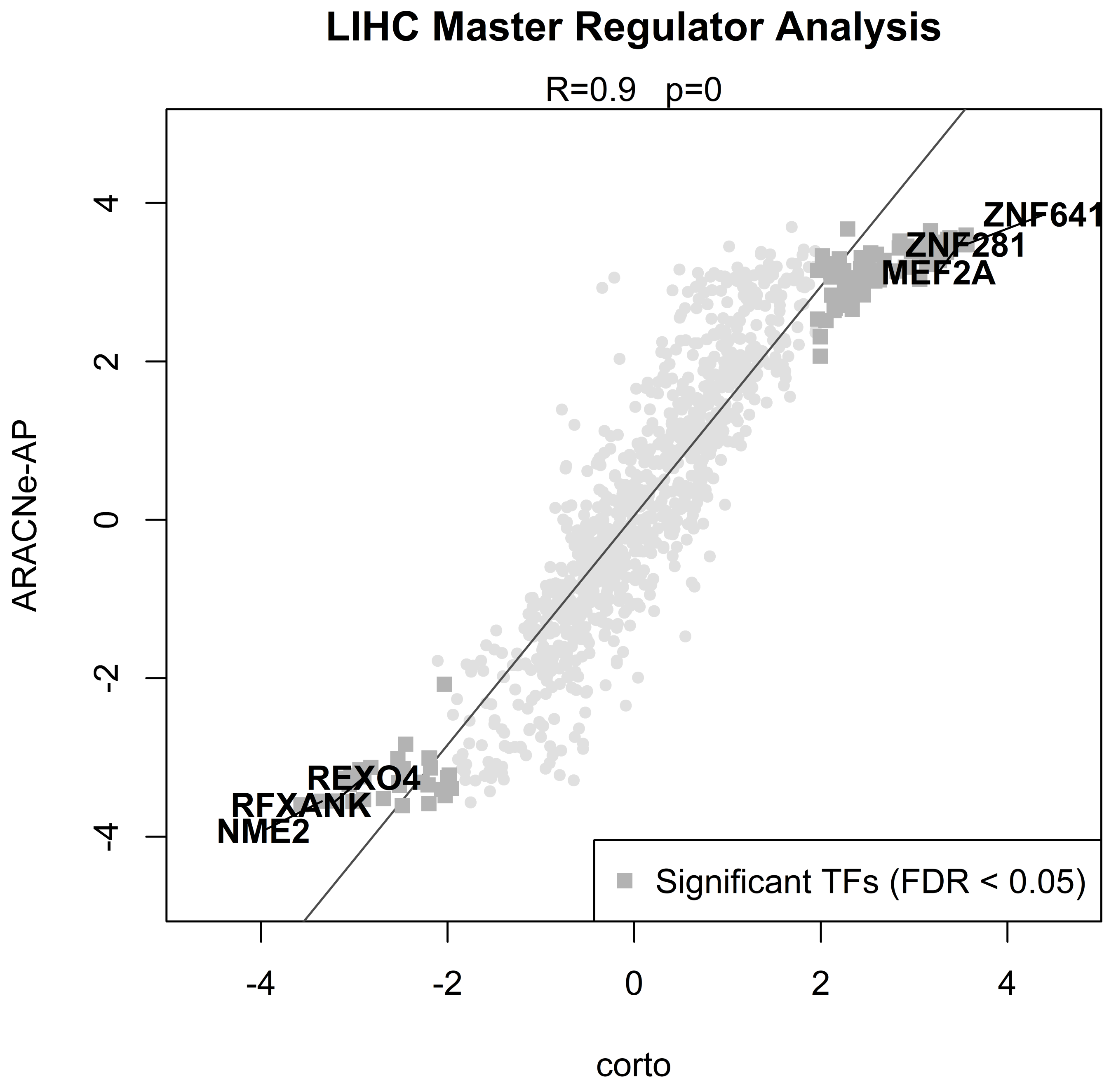

LIHC Master Regulator Analysis
R=0.9 p=0
ARACNe-AP
corto
4
2
0
-2
-4
ZNF641
ZNF281
MEF2A
REXO4
RFXANK
NME2
Significant TFs (FDR < 0.05)

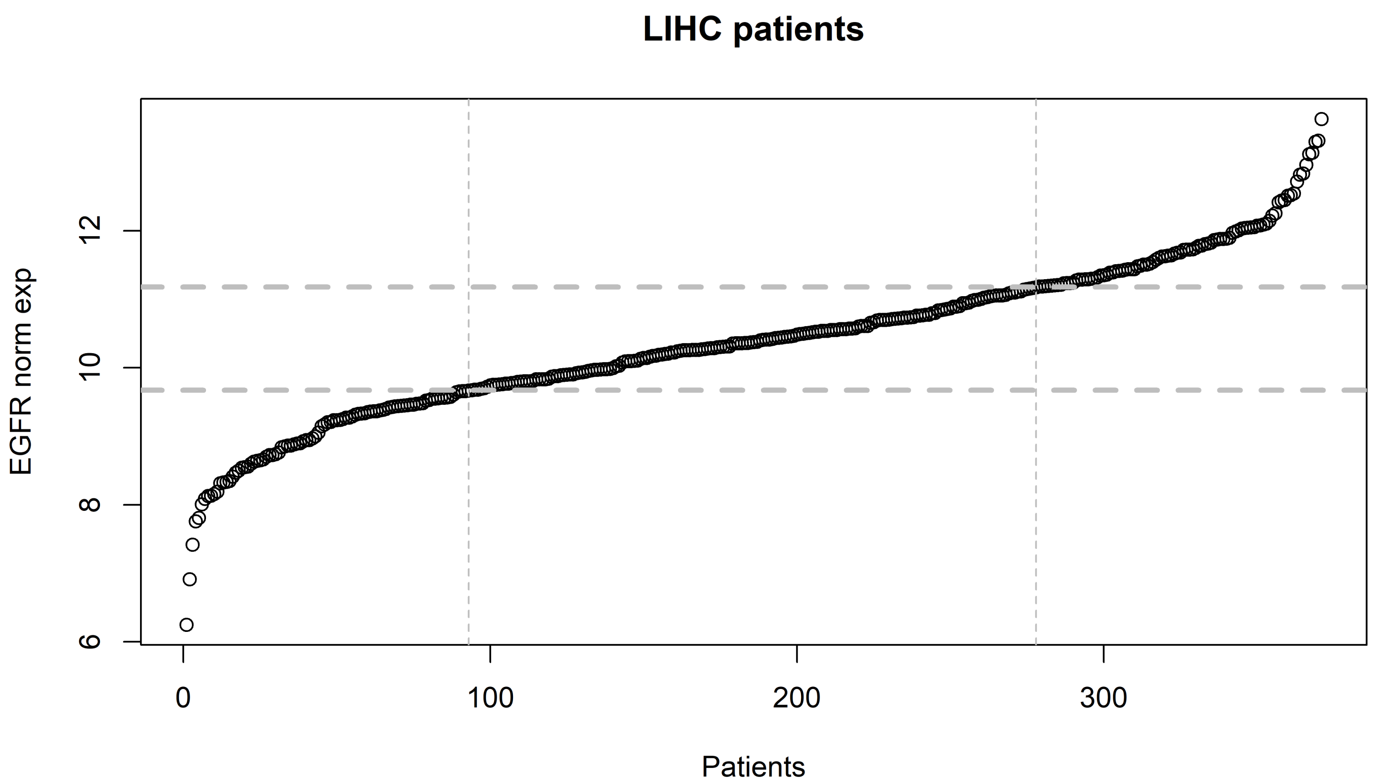

LIHC patients
EGFR norm exp
6
8
10
12
0
100
200
300
Patients

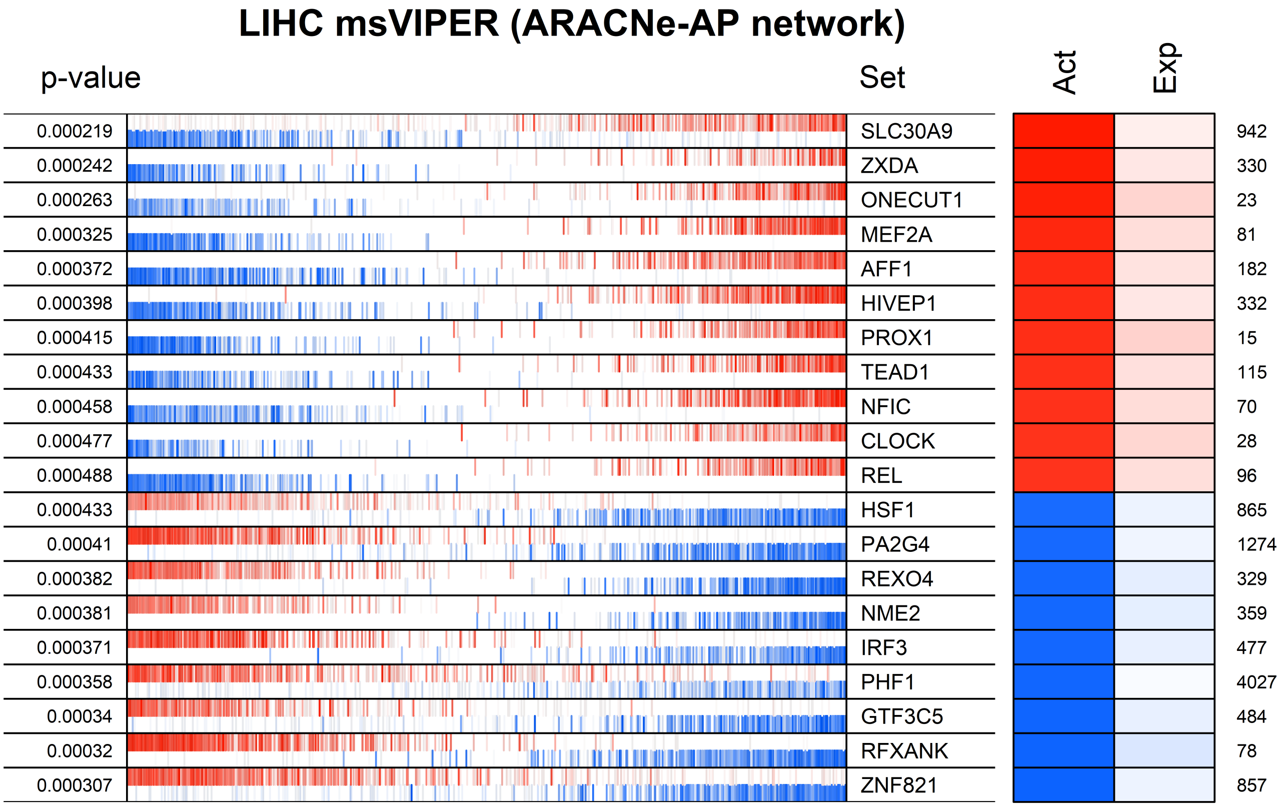

LIHC msVIPER (ARACNe-AP network)
p-value
Set
Act
Exp
0.000219 SLC30A9 942
0.000242 ZXDA 330
0.000263 ONECUT1 23
0.000325 MEF2A 81
0.000372 AFF1 182
0.000398 HIVEP1 332
0.000415 PROX1 15
0.000433 TEAD1 115
0.000458 NFIC 70
0.000477 CLOCK 28
0.000488 REL 96
0.000433 HSF1 865
0.00041 PA2G4 1274
0.000382 REXO4 329
0.000381 NME2 359
0.000371 IRF3 477
0.000358 PHF1 4027
0.00034 GTF3C5 484
0.00032 RFXANK 78
0.000307 ZNF821 857

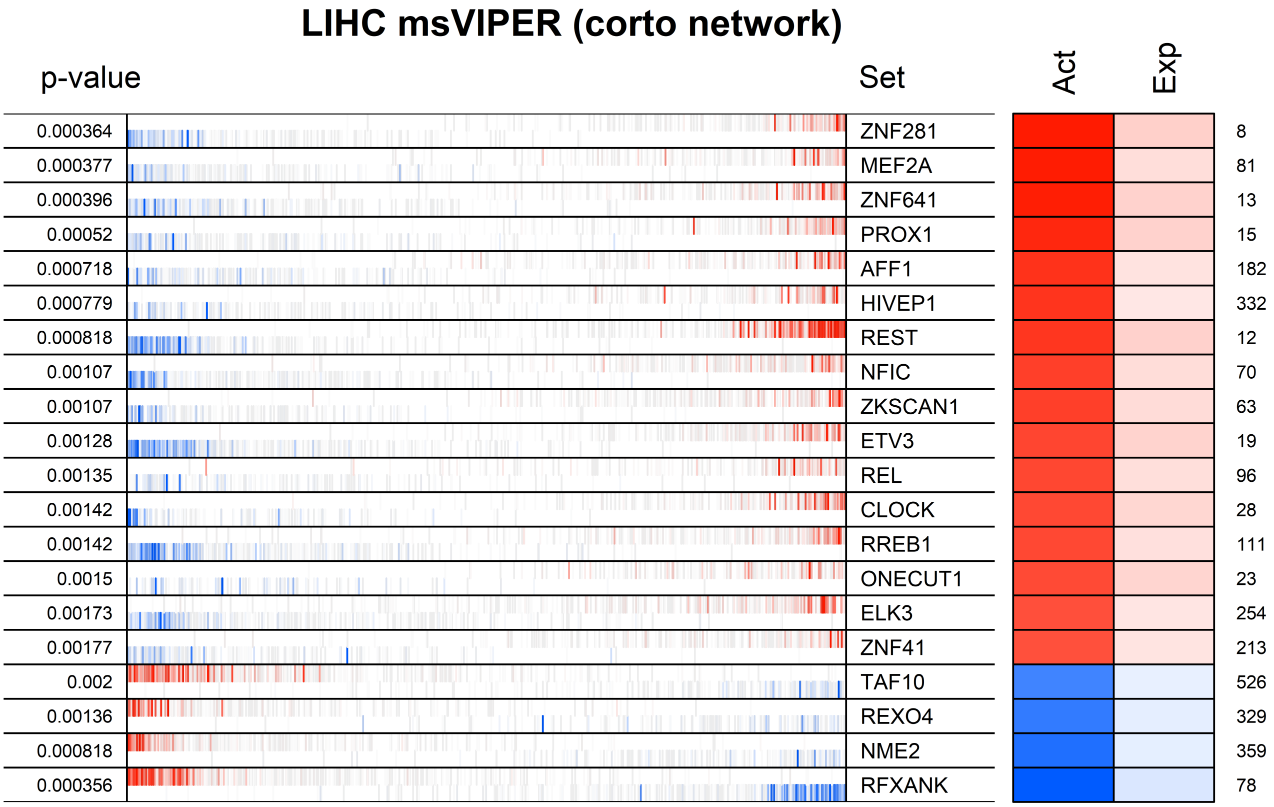
LIHC msVIPER (corto network)
p-value
Set
Act
Exp
0.000364 ZNF281 8
0.000377 MEF2A 81
0.000396 ZNF641 13
0.00052 PROX1 15
0.000718 AFF1 182
0.000779 HIVEP1 332
0.000818 REST 12
0.00107 NFIC 70
0.00107 ZKSCAN1 63
0.00128 ETV3 19
0.00135 REL 96
0.00142 CLOCK 28
0.00142 RREB1 111
0.0015 ONECUT1 23
0.00173 ELK3 254
0.00177 ZNF41 213
0.002 TAF10 526
0.00136 REXO4 329
0.000818 NME2 359
0.000356 RFXANK 78

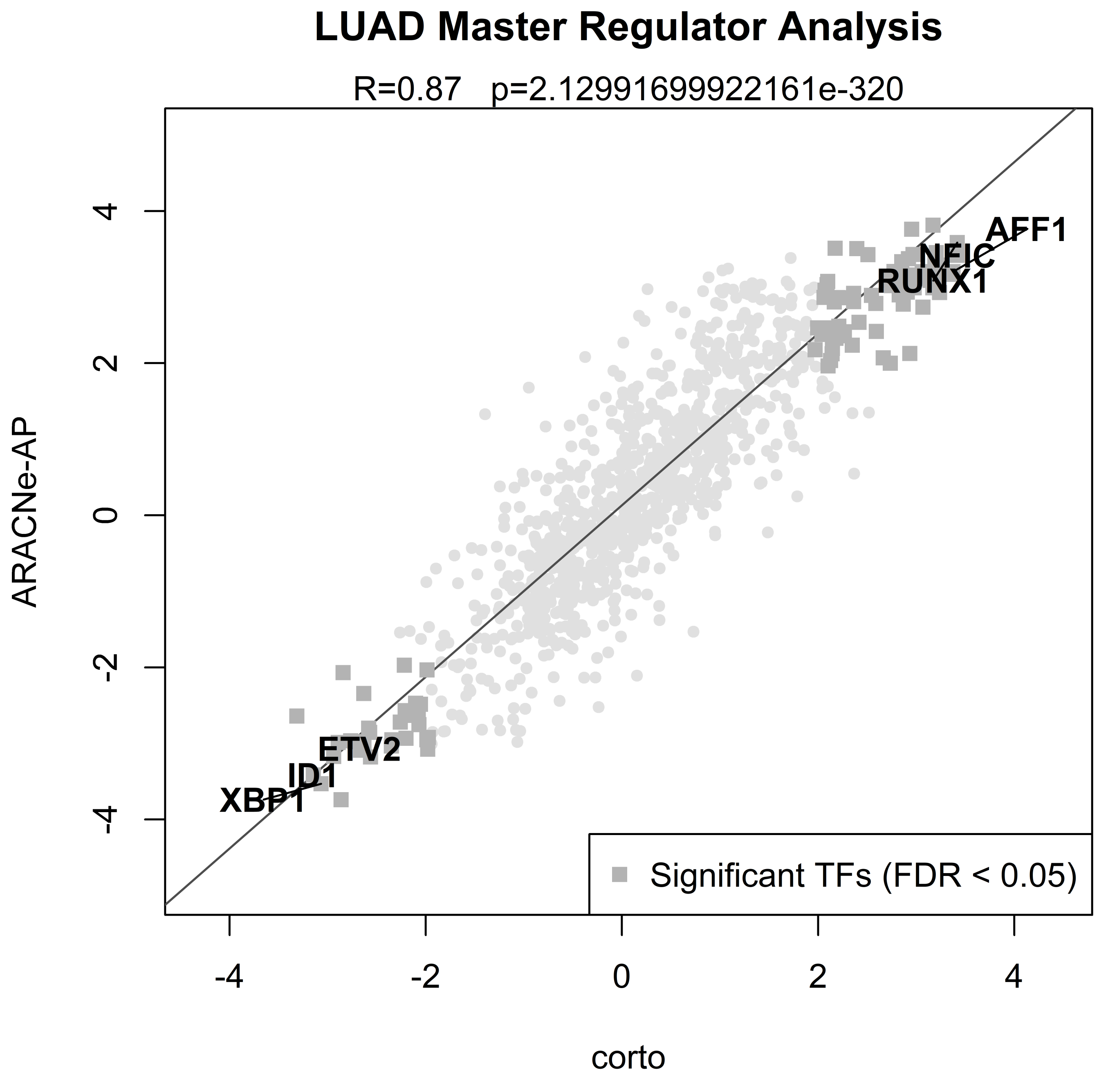

LUAD Master Regulator Analysis
R=0.87   p=2.12991699922161e-320
ARACNe-AP
corto
AFF1
NFIC
RUNX1
ETV2
ID1
XBP1
Significant TFs (FDR < 0.05)
4
2
0
-2
-4
-4
-2
0
2
4

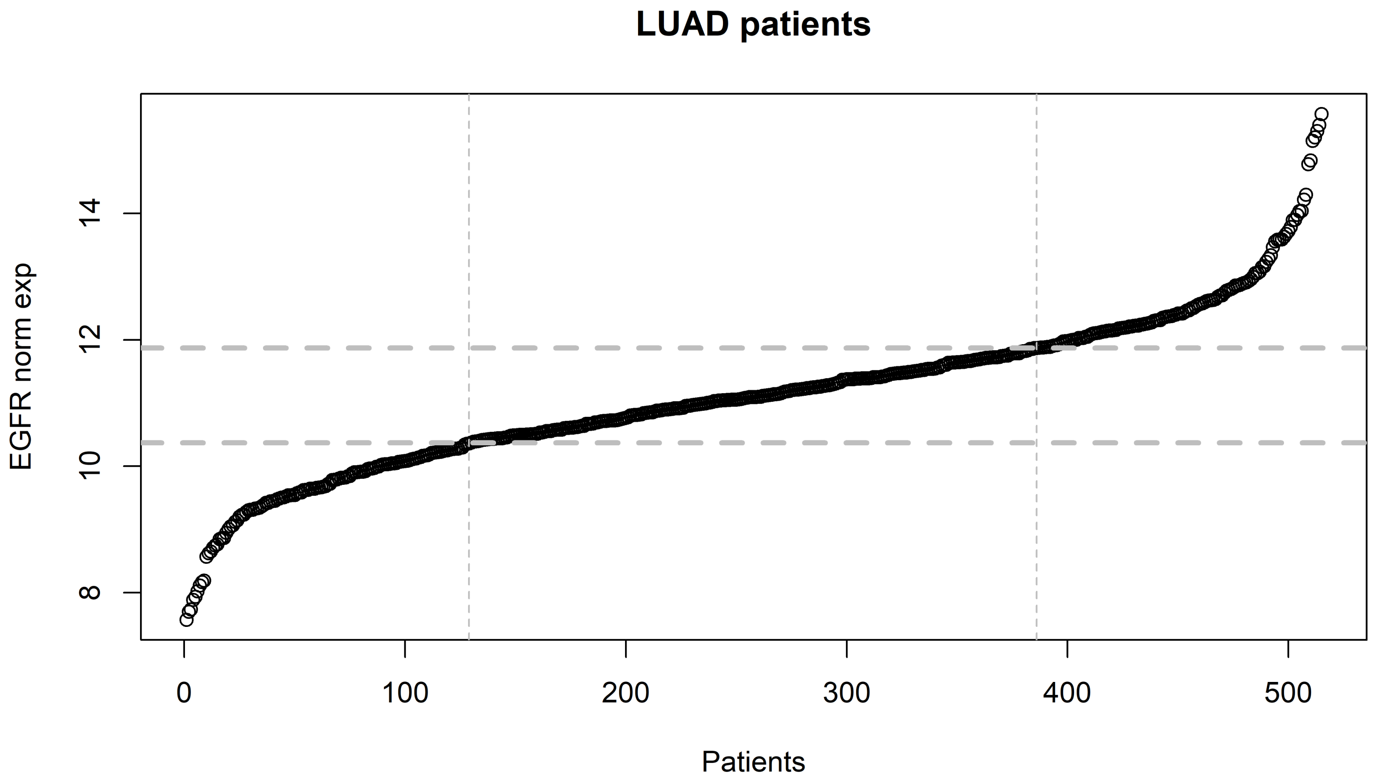
LUAD patients
EGFR norm exp
8
10
12
14
0
100
200
300
400
500
Patients

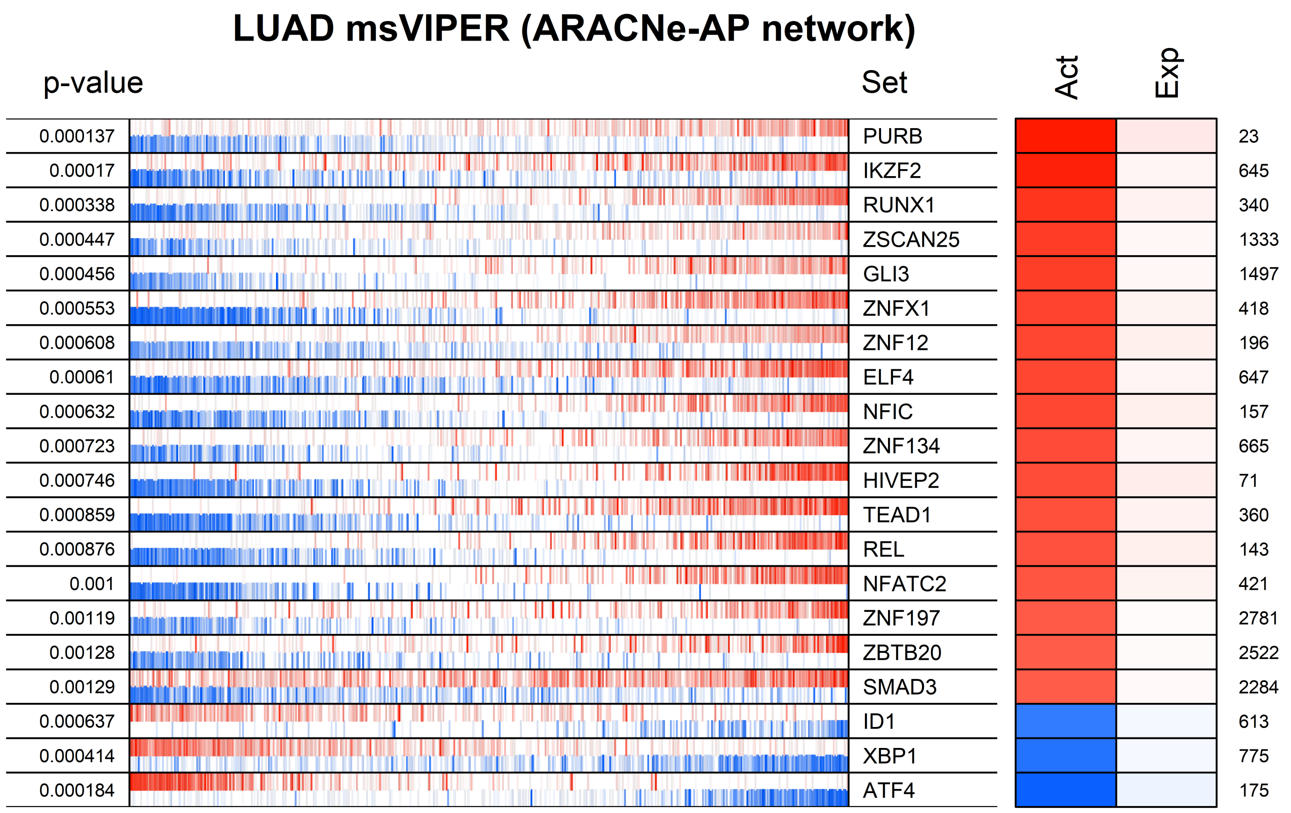

LUAD msVIPER (ARACNe-AP network)
p-value
Set
Act
Exp
0.000137
PURB
23
0.00017
IKZF2
645
0.000338
RUNX1
340
0.000447
ZSCAN25
1333
0.000456
GLI3
1497
0.000553
ZNFX1
418
0.000608
ZNF12
196
0.00061
ELF4
647
0.000632
NFIC
157
0.000723
ZNF134
665
0.000746
HIVEP2
71
0.000859
TEAD1
360
0.000876
REL
143
0.001
NFATC2
421
0.00119
ZNF197
2781
0.00128
ZBTB20
2522
0.00129
SMAD3
2284
0.000637
ID1
613
0.000414
XBP1
775
0.000184
ATF4
175

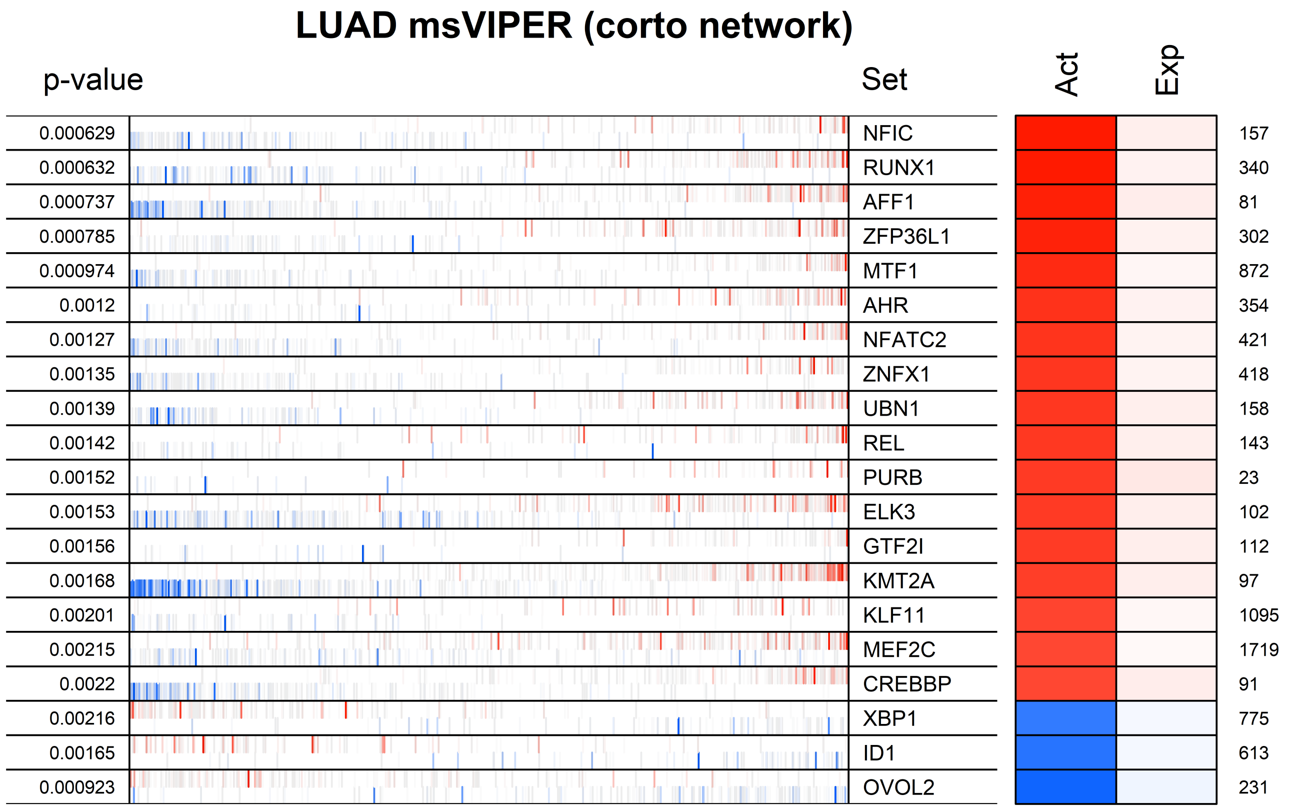
LUAD msVIPER (corto network)
p-value
Set
Act
Exp
0.000629 NFIC 157
0.000632 RUNX1 340
0.000737 AFF1 81
0.000785 ZFP36L1 302
0.000974 MTF1 872
0.0012 AHR 354
0.00127 NFATC2 421
0.00135 ZNFX1 418
0.00139 UBN1 158
0.00142 REL 143
0.00152 PURB 23
0.00153 ELK3 102
0.00156 GTF2I 112
0.00168 KMT2A 97
0.00201 KLF11 1095
0.00215 MEF2C 1719
0.0022 CREBBP 91
0.00216 XBP1 775
0.00165 ID1 613
0.000923 OVOL2 231

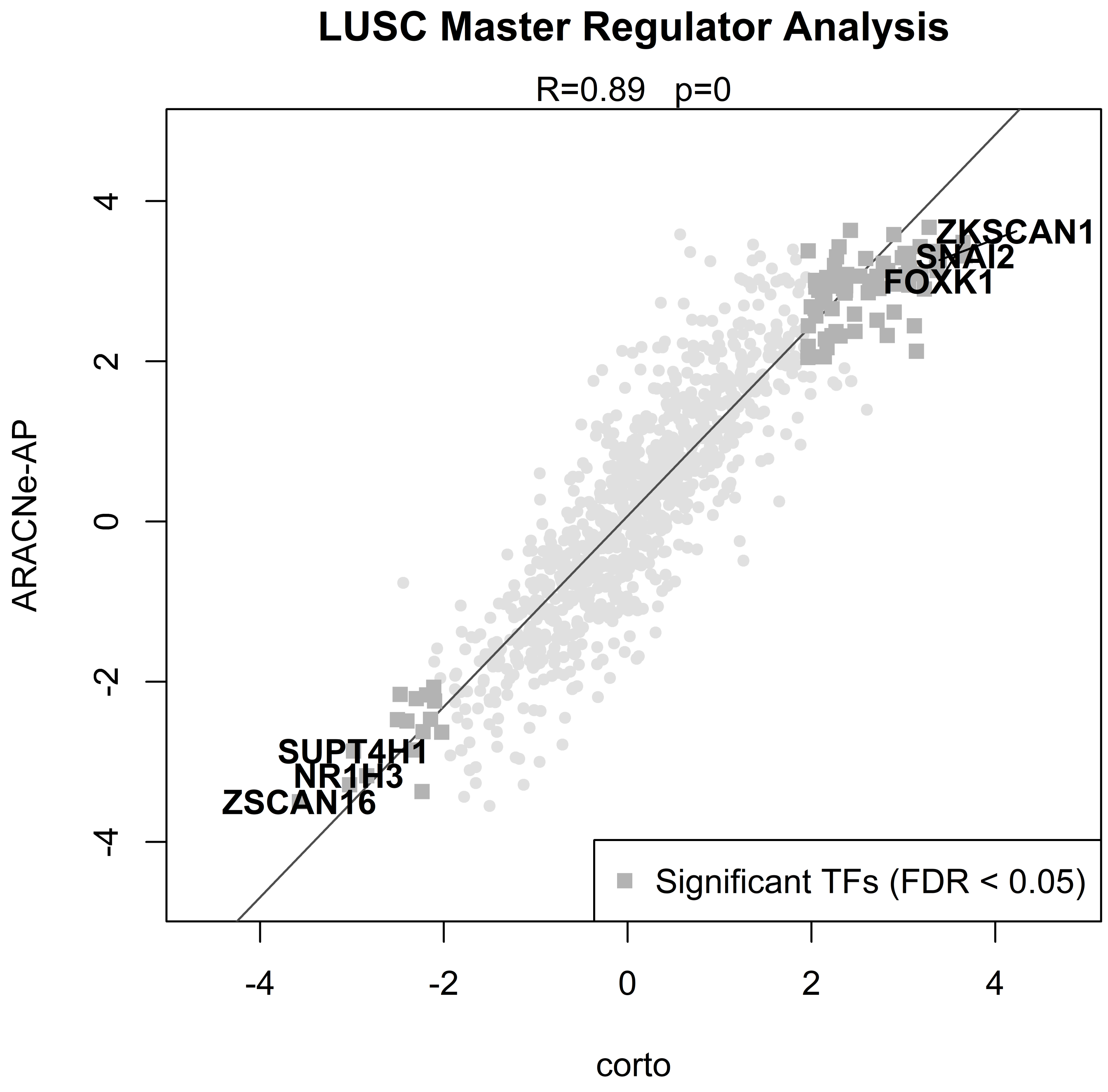

LUSC Master Regulator Analysis
R=0.89 p=0
ARACNe-AP
corto
4
2
0
-2
-4
-4
-2
0
2
4
ZKSCAN1
SNAI2
FOXK1
SUPT4H1
NR1H3
ZSCAN16
Significant TFs (FDR < 0.05)

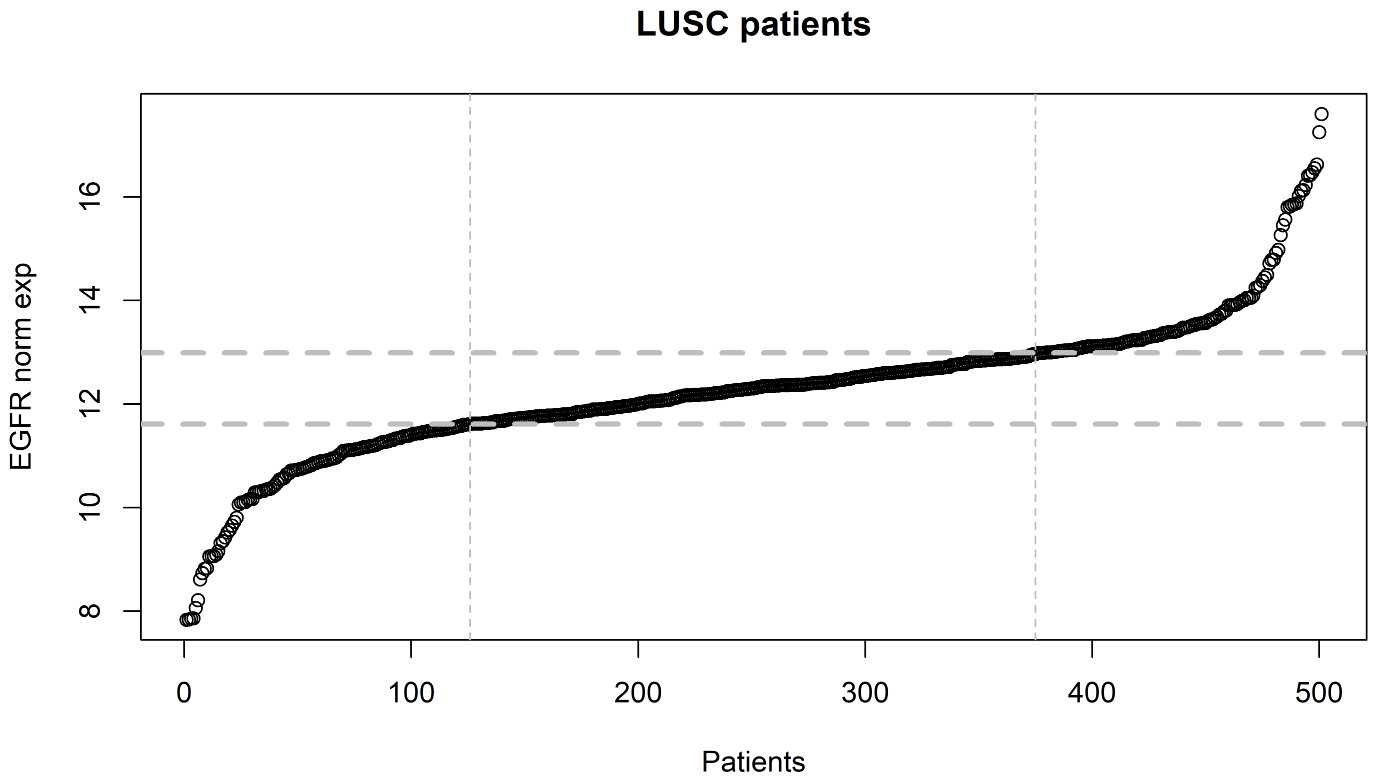

LUSC patients
EGFR norm exp
Patients
8
10
12
14
16
0
100
200
300
400
500

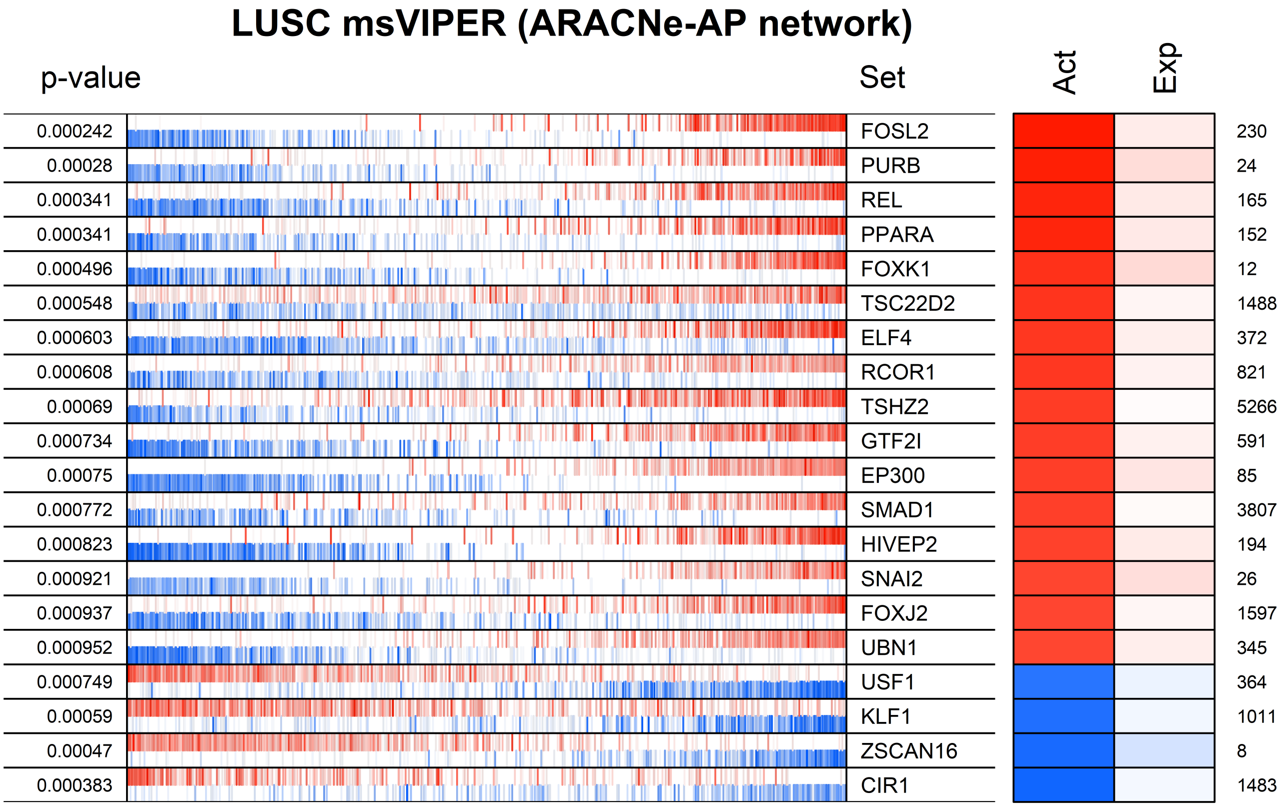

LUSC msVIPER (ARACNe-AP network)
p-value
Set
Act
Exp
0.000242
FOSL2
230
0.00028
PURB
24
0.000341
REL
165
0.000341
PPARA
152
0.000496
FOXK1
12
0.000548
TSC22D2
1488
0.000603
ELF4
372
0.000608
RCOR1
821
0.00069
TSHZ2
5266
0.000734
GTF2I
591
0.00075
EP300
85
0.000772
SMAD1
3807
0.000823
HIVEP2
194
0.000921
SNAI2
26
0.000937
FOXJ2
1597
0.000952
UBN1
345
0.000749
USF1
364
0.00059
KLF1
1011
0.00047
ZSCAN16
8
0.000383
CIR1
1483

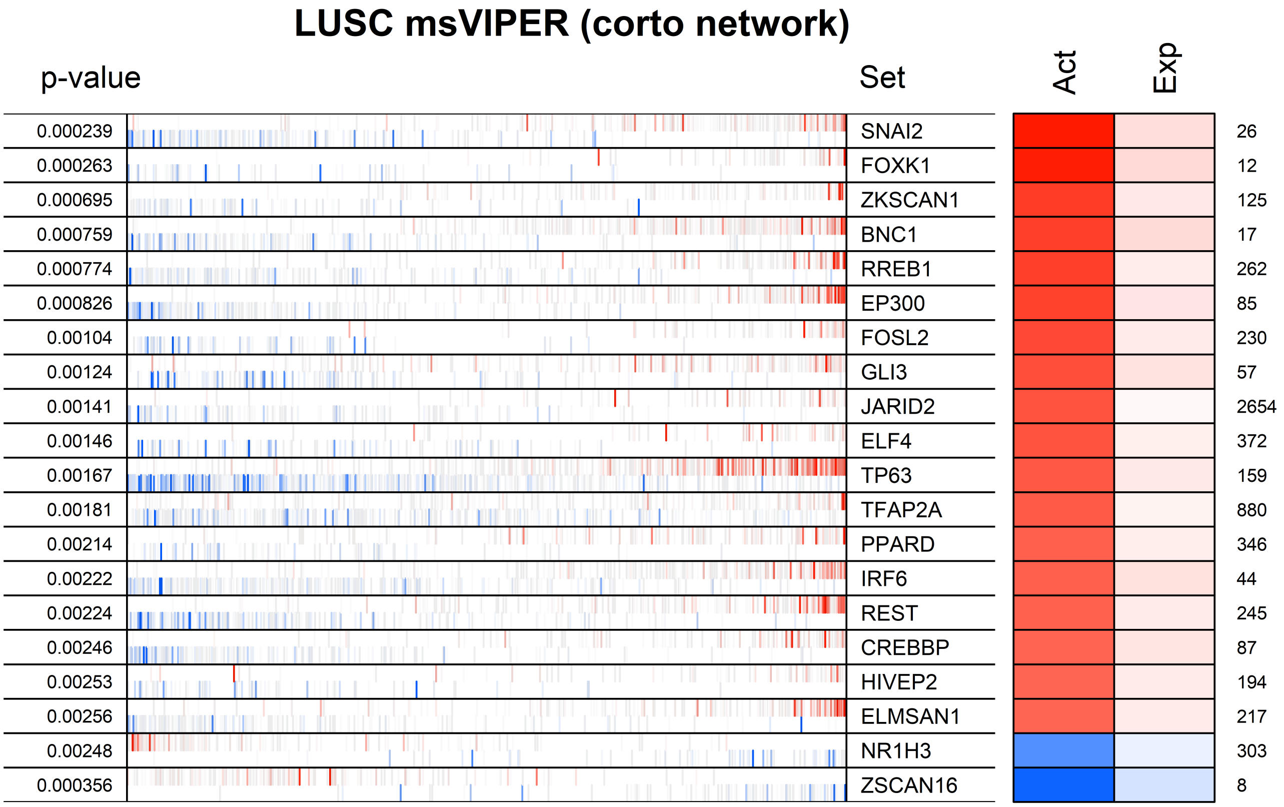
LUSC msVIPER (corto network)
p-value
Set
Act
Exp
0.000239 SNAI2 26
0.000263 FOXK1 12
0.000695 ZKSCAN1 125
0.000759 BNC1 17
0.000774 RREB1 262
0.000826 EP300 85
0.00104 FOSL2 230
0.00124 GLI3 57
0.00141 JARID2 2654
0.00146 ELF4 372
0.00167 TP63 159
0.00181 TFAP2A 880
0.00214 PPARD 346
0.00222 IRF6 44
0.00224 REST 245
0.00246 CREBBP 87
0.00253 HIVEP2 194
0.00256 ELMSAN1 217
0.00248 NR1H3 303
0.000356 ZSCAN16 8

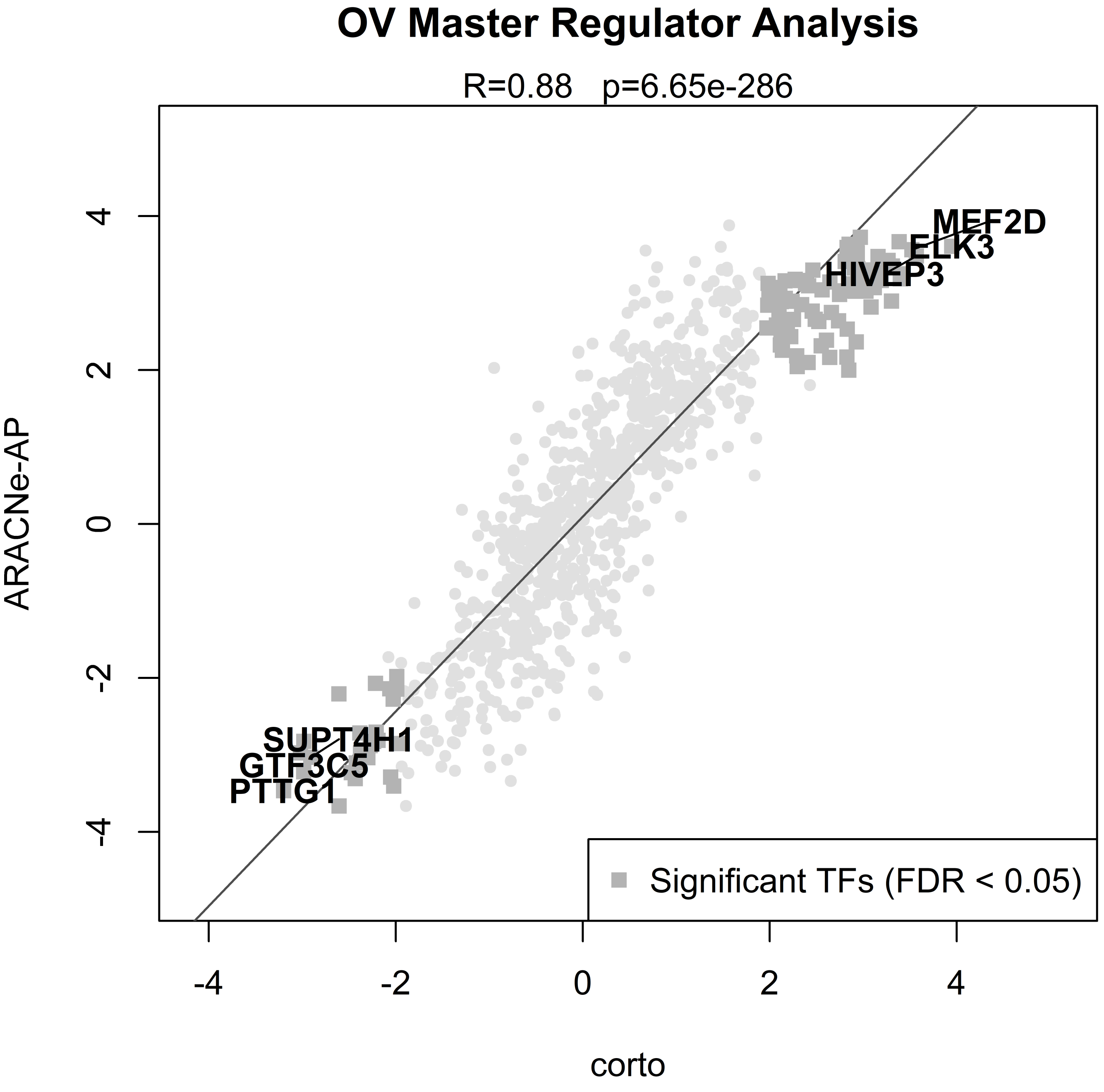

OV Master Regulator Analysis
R=0.88   p=6.65e-286
MEF2D
ELK3
HIVEP3
SUPT4H1
GTF3C5
PTTG1
Significant TFs (FDR < 0.05)
ARACNe-AP
corto
4
2
0
-2
-4
-4
-2
0
2
4

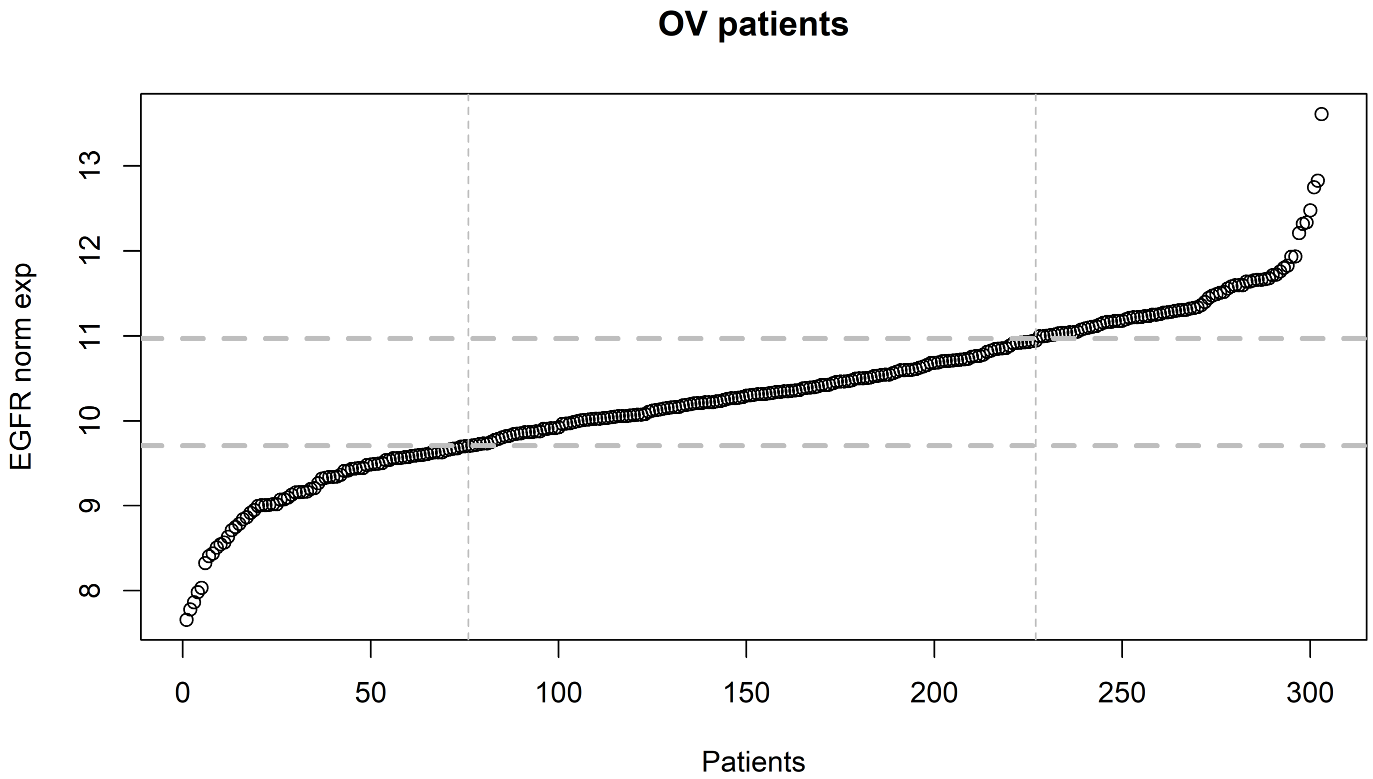
OV patients
EGFR norm exp
8
9
10
11
12
13
0
50
100
150
200
250
300
Patients

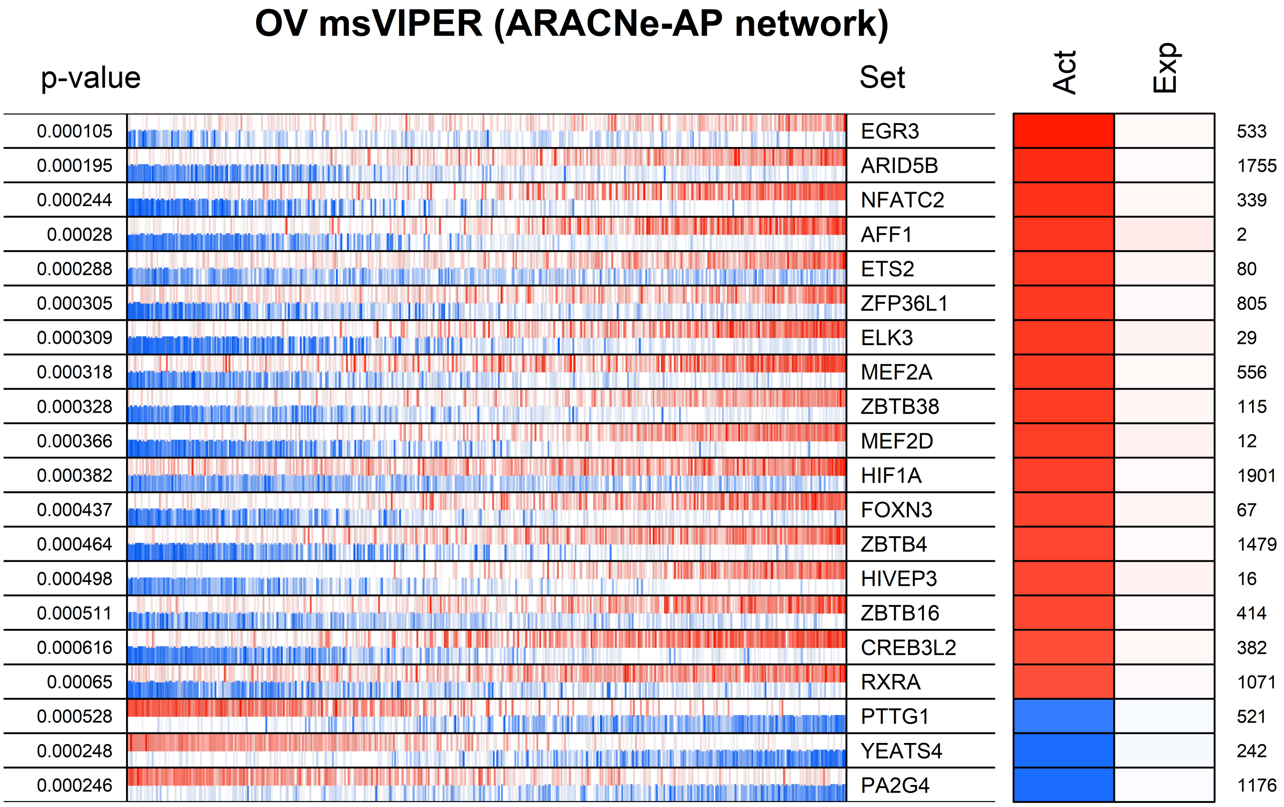
OV msVIPER (ARACNe-AP network)
p-value
Set
Act
Exp
0.000105 EGR3 533
0.000195 ARID5B 1755
0.000244 NFATC2 339
0.00028 AFF1 2
0.000288 ETS2 80
0.000305 ZFP36L1 805
0.000309 ELK3 29
0.000318 MEF2A 556
0.000328 ZBTB38 115
0.000366 MEF2D 12
0.000382 HIF1A 1901
0.000437 FOXN3 67
0.000464 ZBTB4 1479
0.000498 HIVEP3 16
0.000511 ZBTB16 414
0.000616 CREB3L2 382
0.00065 RXRA 1071
0.000528 PTTG1 521
0.000248 YEATS4 242
0.000246 PA2G4 1176

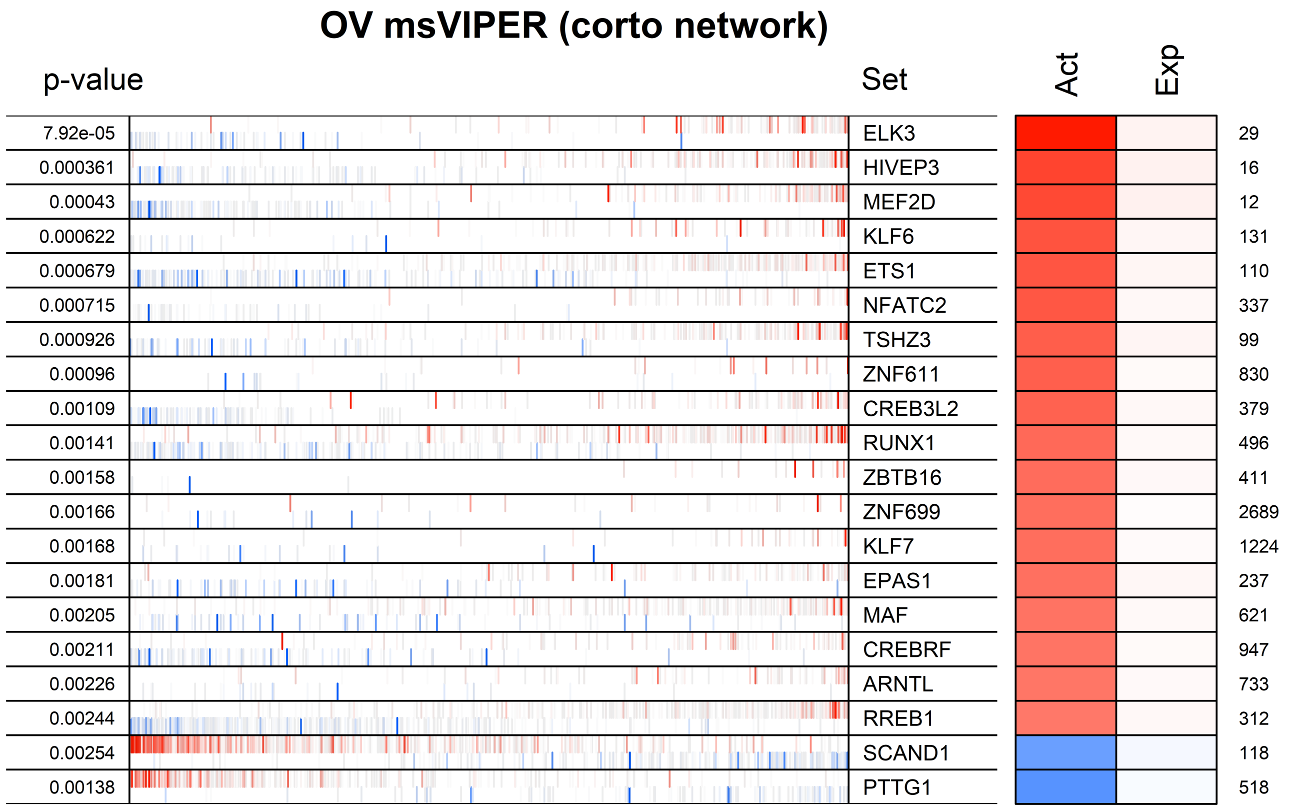
OV msVIPER (corto network)
p-value
Set
Act
Exp
7.92e-05 ELK3 29
0.000361 HIVEP3 16
0.00043 MEF2D 12
0.000622 KLF6 131
0.000679 ETS1 110
0.000715 NFATC2 337
0.000926 TSHZ3 99
0.00096 ZNF611 830
0.00109 CREB3L2 379
0.00141 RUNX1 496
0.00158 ZBTB16 411
0.00166 ZNF699 2689
0.00168 KLF7 1224
0.00181 EPAS1 237
0.00205 MAF 621
0.00211 CREBRF 947
0.00226 ARNTL 733
0.00244 RREB1 312
0.00254 SCAND1 118
0.00138 PTTG1 518

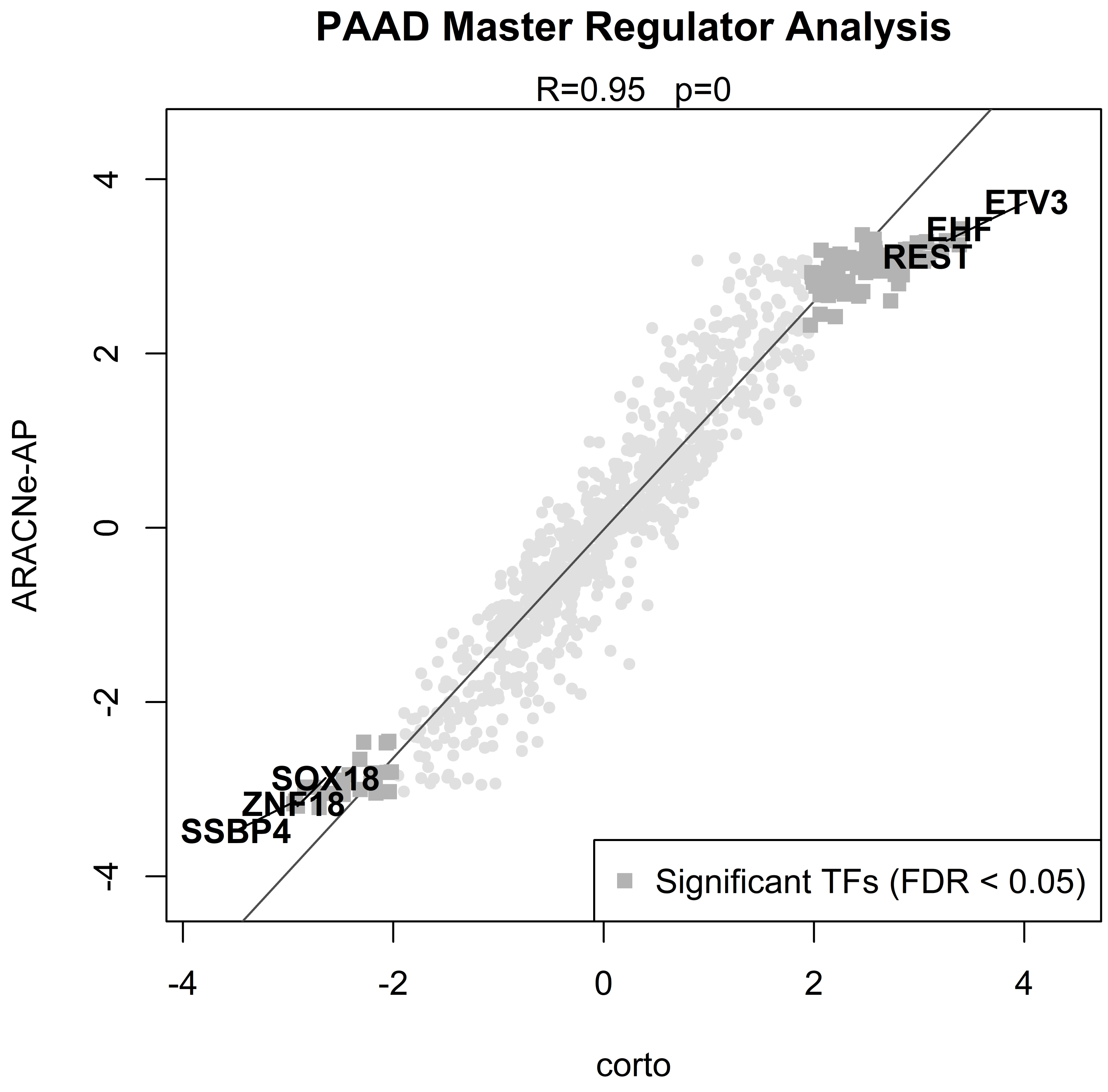
PAAD Master Regulator Analysis
R=0.95 p=0
ARACNe-AP
corto
-4
-2
0
2
4
ETV3
EHF
REST
SOX18
ZNF18
SSBP4
Significant TFs (FDR < 0.05)

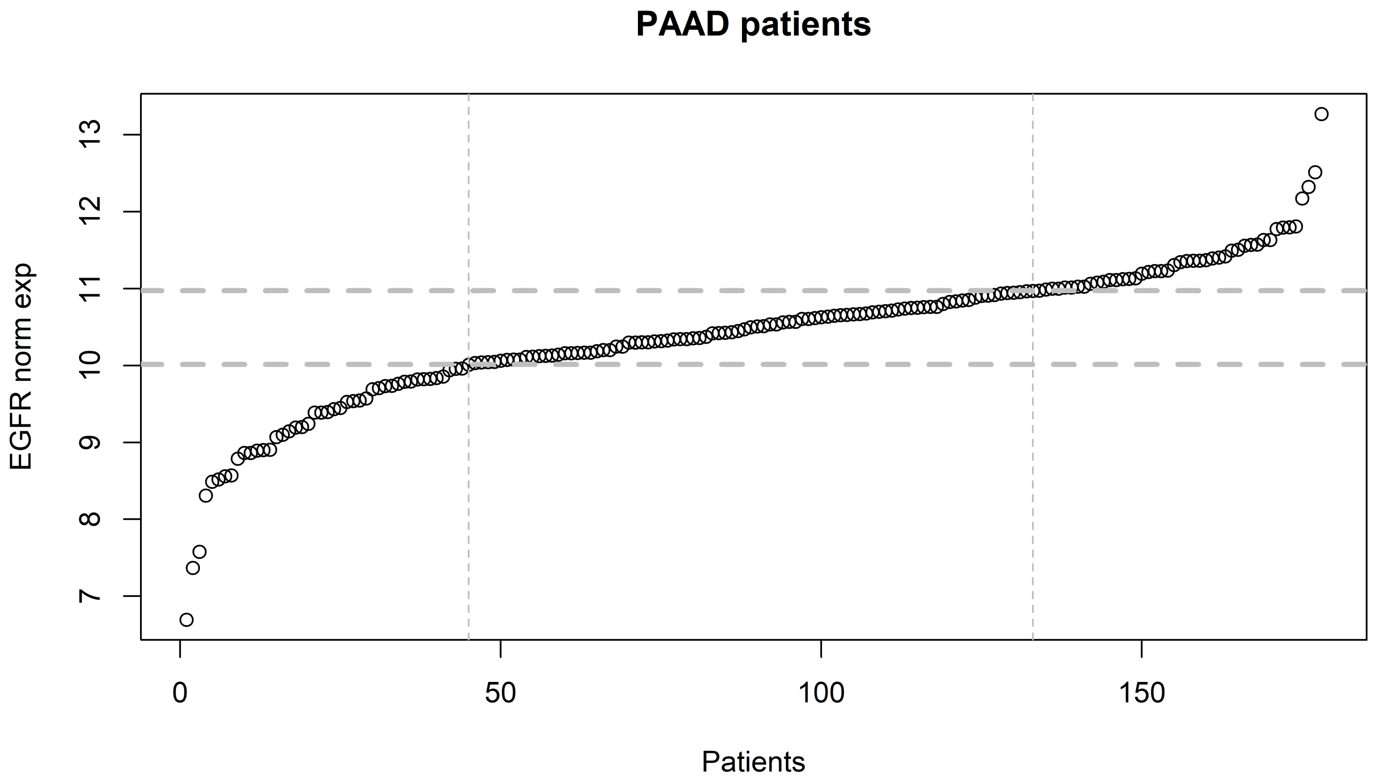
PAAD patients
EGFR norm exp
7
8
9
10
11
12
13
0
50
100
150
Patients

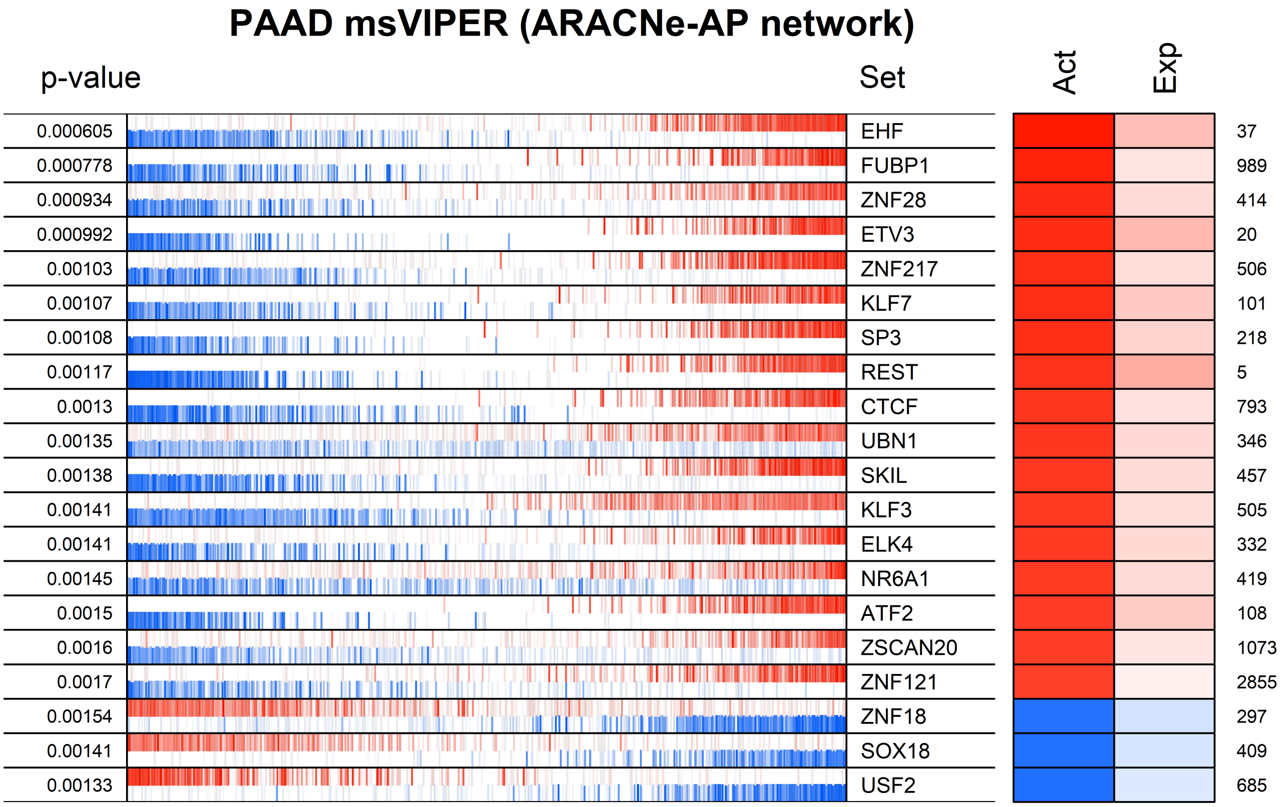
PAAD msVIPER (ARACNe-AP network)
p-value
Set
Act
Exp
0.000605 EHF 37
0.000778 FUBP1 989
0.000934 ZNF28 414
0.000992 ETV3 20
0.00103 ZNF217 506
0.00107 KLF7 101
0.00108 SP3 218
0.00117 REST 5
0.0013 CTCF 793
0.00135 UBN1 346
0.00138 SKIL 457
0.00141 KLF3 505
0.00141 ELK4 332
0.00145 NR6A1 419
0.0015 ATF2 108
0.0016 ZSCAN20 1073
0.0017 ZNF121 2855
0.00154 ZNF18 297
0.00141 SOX18 409
0.00133 USF2 685

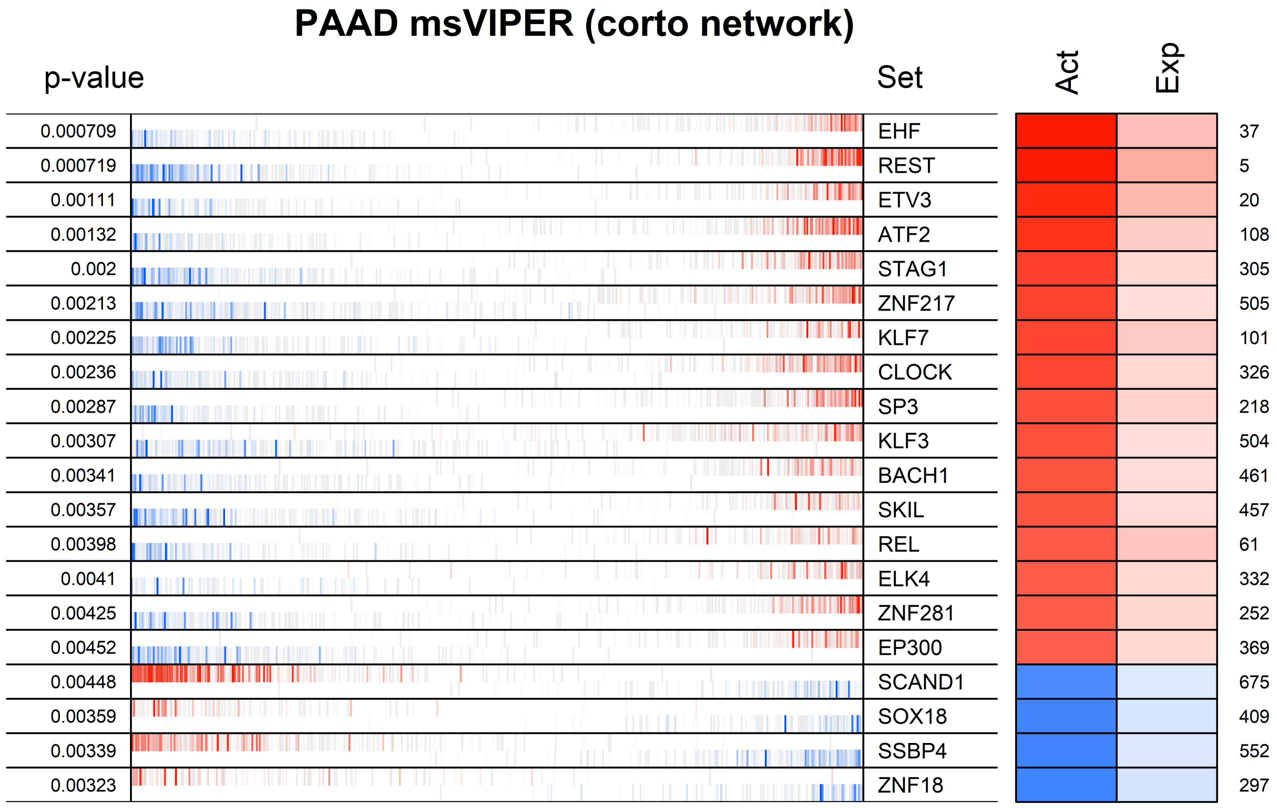
PAAD msVIPER (corto network)
p-value
Set
Act
Exp
0.000709 EHF 37
0.000719 REST 5
0.00111 ETV3 20
0.00132 ATF2 108
0.002 STAG1 305
0.00213 ZNF217 505
0.00225 KLF7 101
0.00236 CLOCK 326
0.00287 SP3 218
0.00307 KLF3 504
0.00341 BACH1 461
0.00357 SKIL 457
0.00398 REL 61
0.0041 ELK4 332
0.00425 ZNF281 252
0.00452 EP300 369
0.00448 SCAND1 675
0.00359 SOX18 409
0.00339 SSBP4 552
0.00323 ZNF18 297

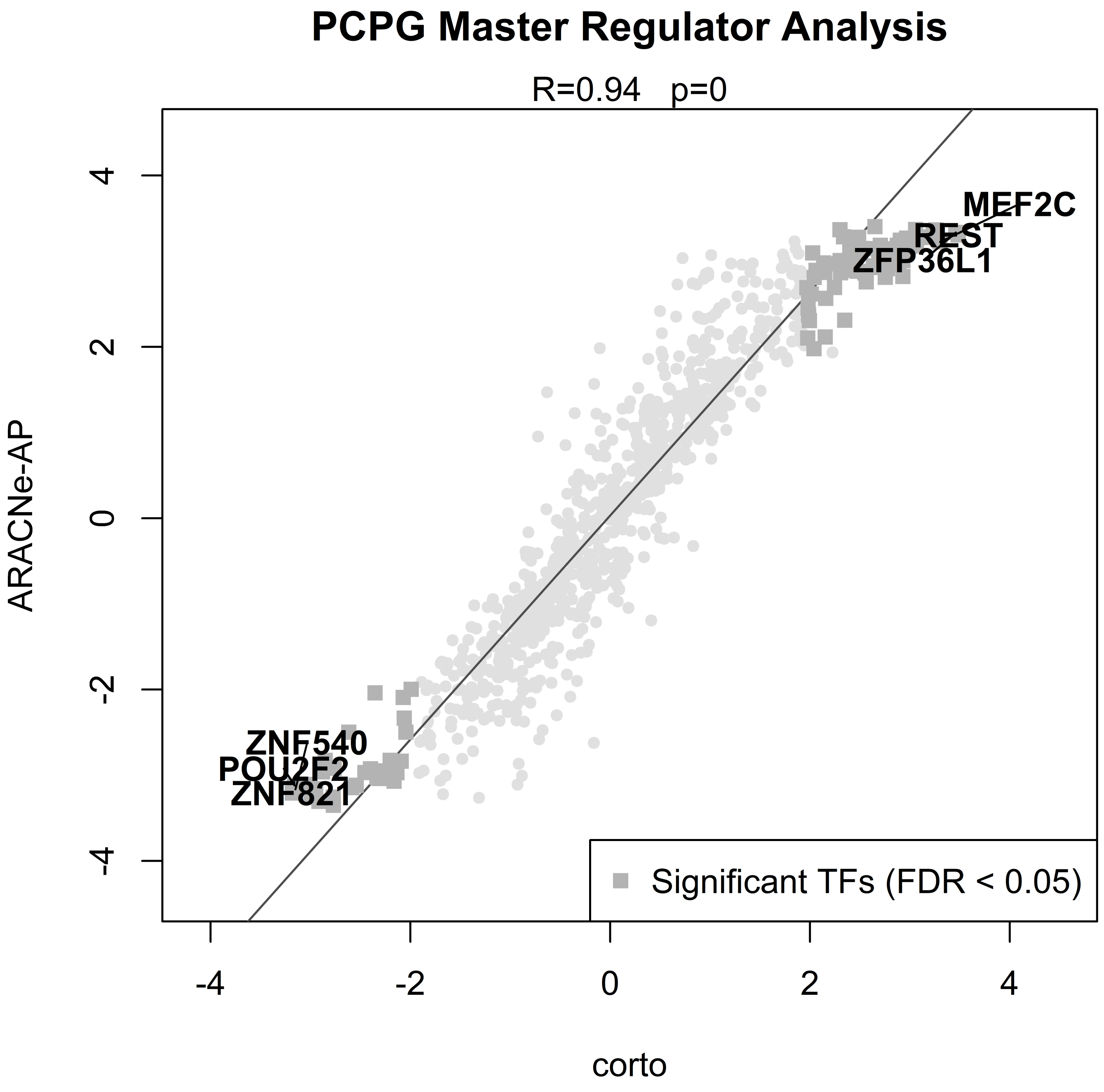
PCPG Master Regulator Analysis
R=0.94 p=0
ARACNe-AP
corto
4
2
0
-2
-4
-4
-2
0
2
4
MEF2C
REST
ZFP36L1
ZNF540
POU2F2
ZNF821
Significant TFs (FDR < 0.05)

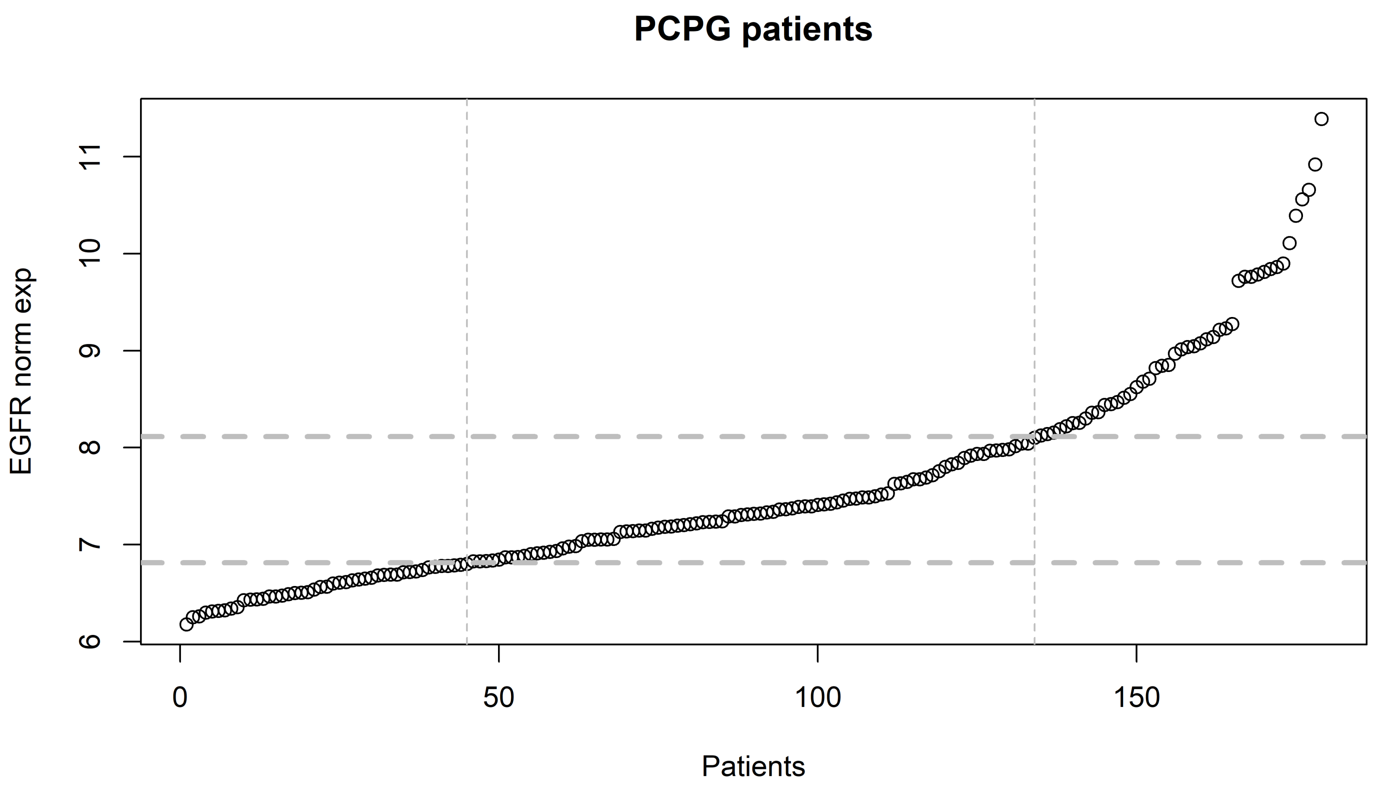
PCPG patients
EGFR norm exp
6
7
8
9
10
11
0
50
100
150
Patients

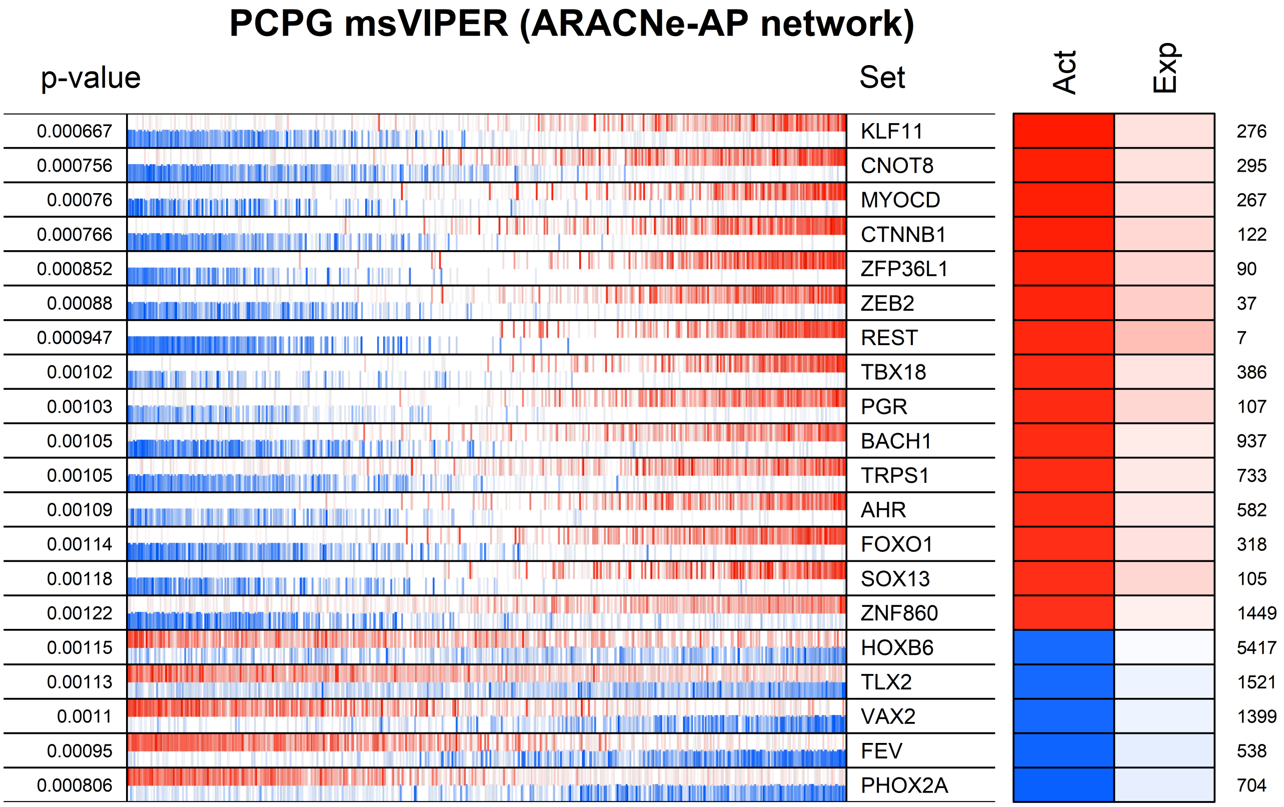

PCPG msVIPER (ARACNe-AP network)
p-value
Set
Act
Exp
0.000667 KLF11 276
0.000756 CNOT8 295
0.00076 MYOCD 267
0.000766 CTNNB1 122
0.000852 ZFP36L1 90
0.00088 ZEB2 37
0.000947 REST 7
0.00102 TBX18 386
0.00103 PGR 107
0.00105 BACH1 937
0.00105 TRPS1 733
0.00109 AHR 582
0.00114 FOXO1 318
0.00118 SOX13 105
0.00122 ZNF860 1449
0.00115 HOXB6 5417
0.00113 TLX2 1521
0.0011 VAX2 1399
0.00095 FEV 538
0.000806 PHOX2A 704

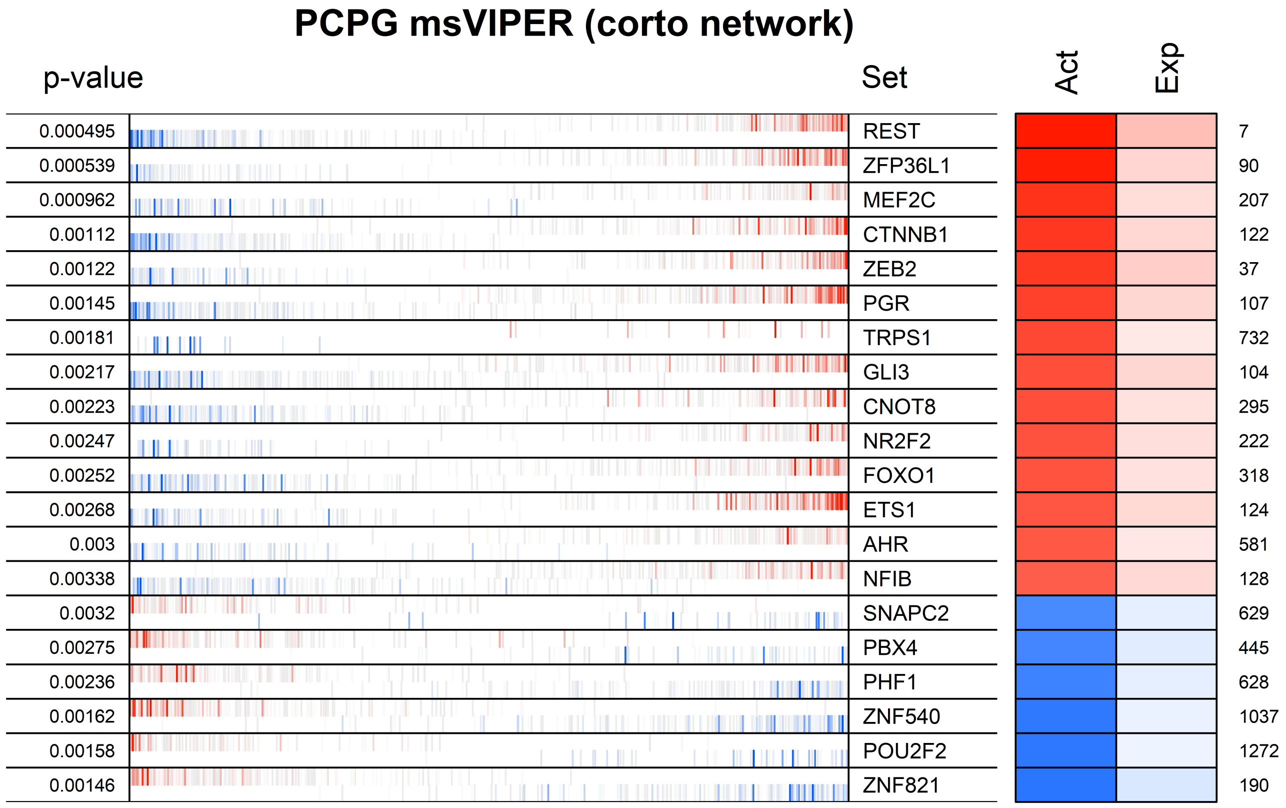
PCPG msVIPER (corto network)
p-value
Set
Act
Exp
0.000495 REST 7
0.000539 ZFP36L1 90
0.000962 MEF2C 207
0.00112 CTNNB1 122
0.00122 ZEB2 37
0.00145 PGR 107
0.00181 TRPS1 732
0.00217 GLI3 104
0.00223 CNOT8 295
0.00247 NR2F2 222
0.00252 FOXO1 318
0.00268 ETS1 124
0.003 AHR 581
0.00338 NFIB 128
0.0032 SNAPC2 629
0.00275 PBX4 445
0.00236 PHF1 628
0.00162 ZNF540 1037
0.00158 POU2F2 1272
0.00146 ZNF821 190

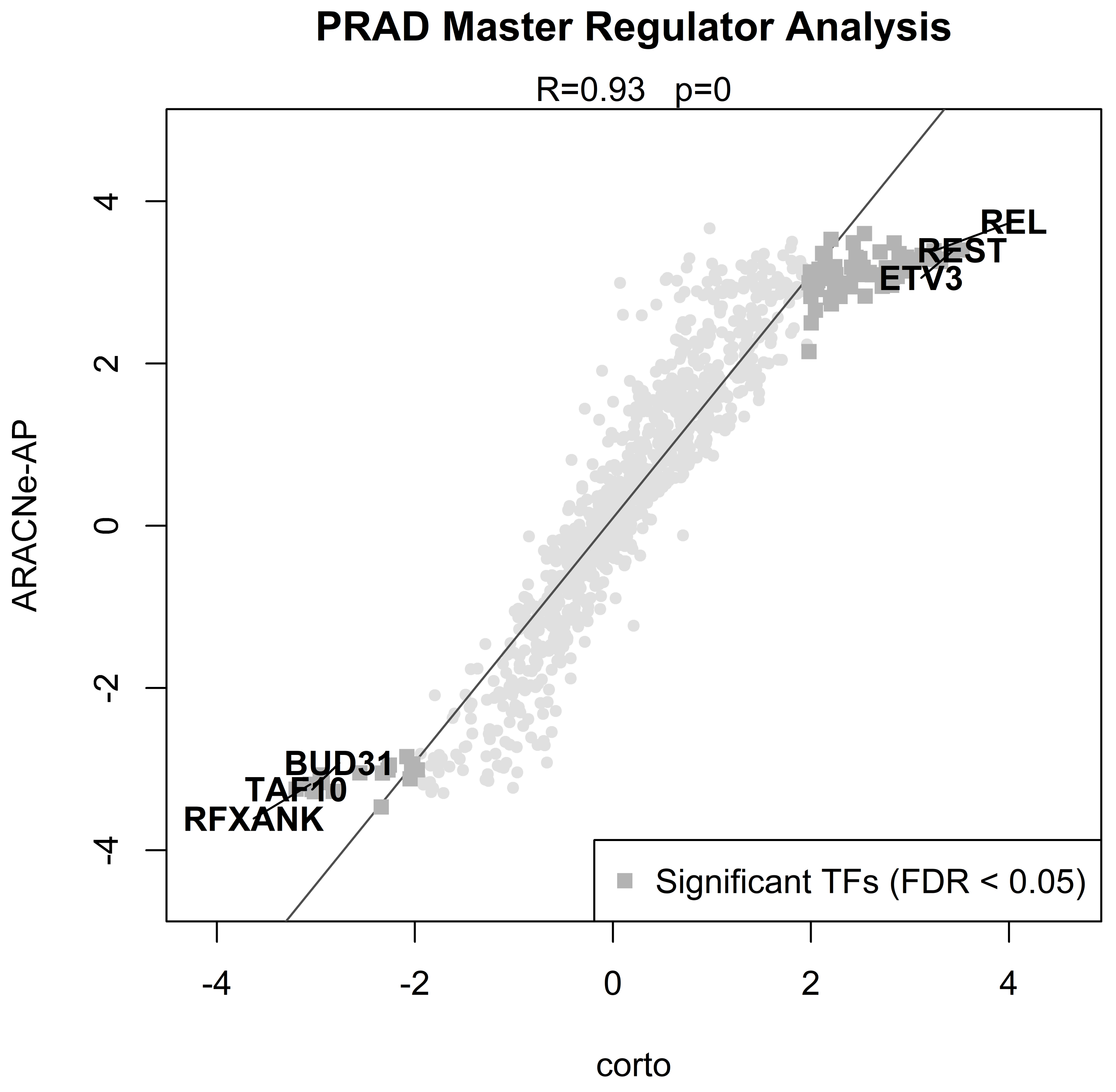
PRAD Master Regulator Analysis
R=0.93 p=0
ARACNe-AP
corto
4
2
0
-2
-4
-4
-2
0
2
4
REL
REST
ETV3
BUD31
TAF10
RFXANK
Significant TFs (FDR < 0.05)

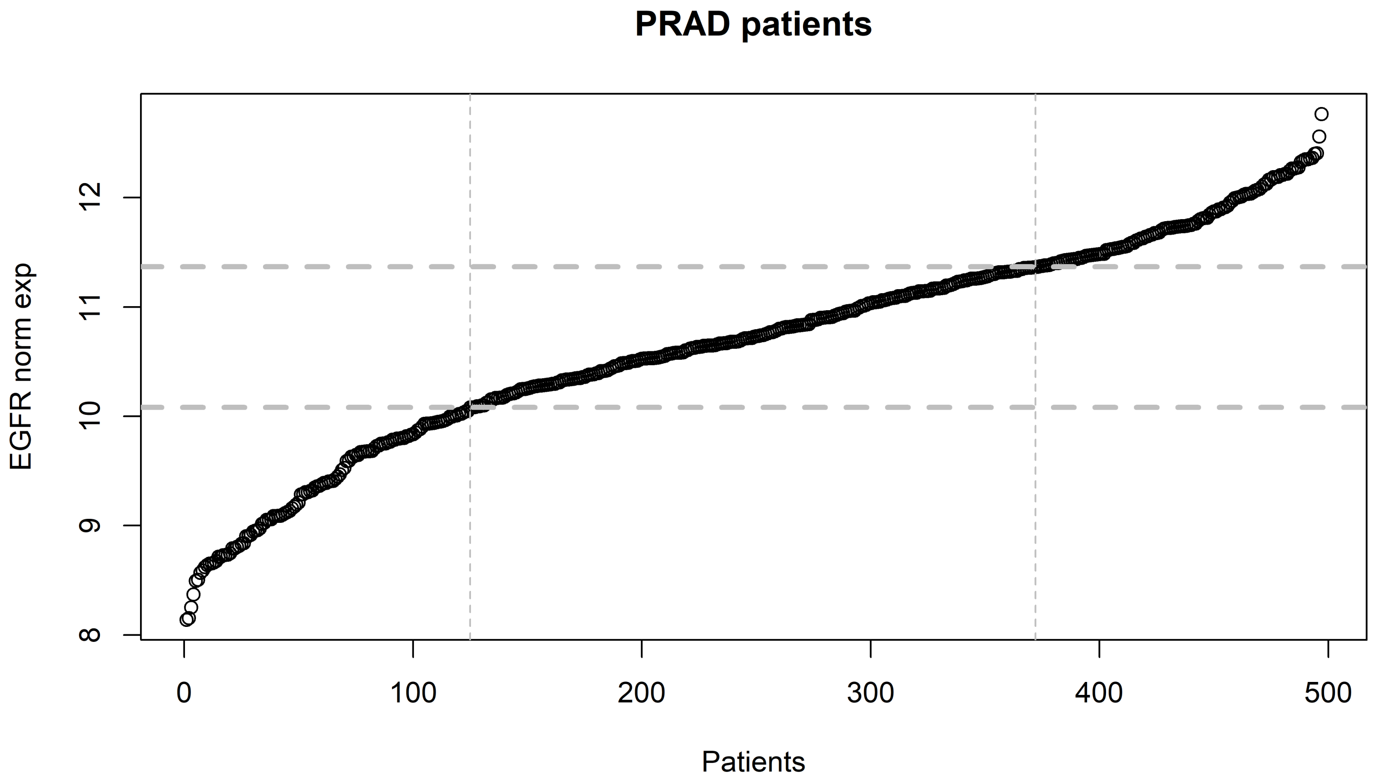

PRAD patients
EGFR norm exp
Patients
8
9
10
11
12
0
100
200
300
400
500

PRAD msVIPER (ARACNe-AP network)

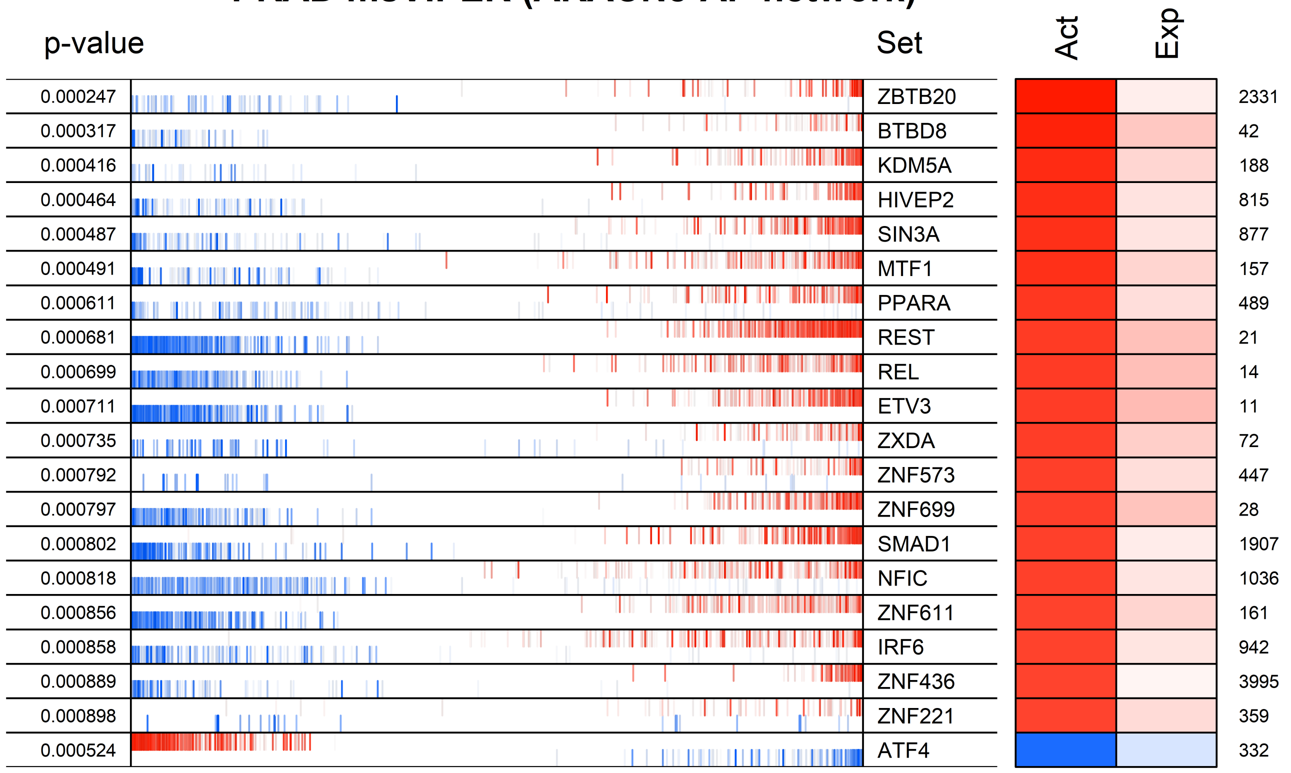

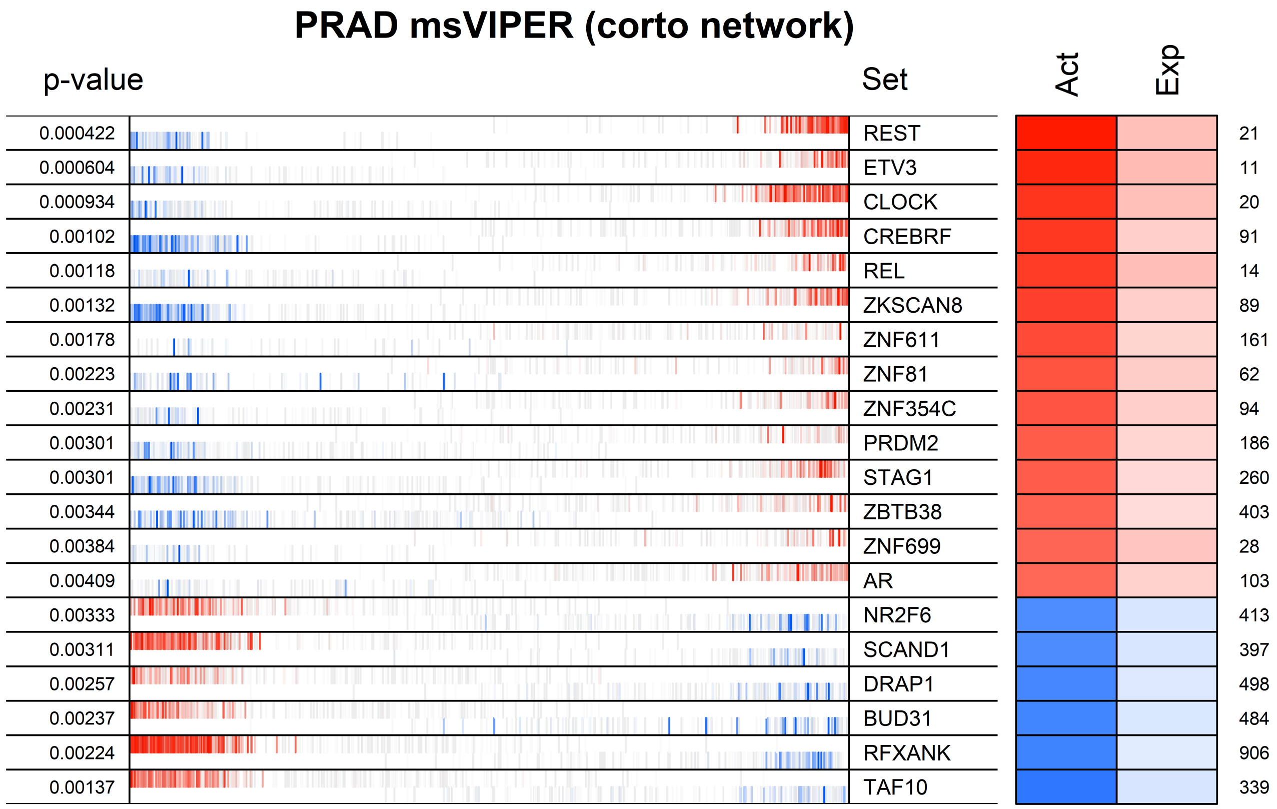
PRAD msVIPER (corto network)
p-value
Set
Act
Exp
0.000422
0.000604
0.000934
0.00102
0.00118
0.00132
0.00178
0.00223
0.00231
0.00301
0.00301
0.00344
0.00384
0.00409
0.00333
0.00311
0.00257
0.00237
0.00224
0.00137
REST
ETV3
CLOCK
CREBRF
REL
ZKSCAN8
ZNF611
ZNF81
ZNF354C
PRDM2
STAG1
ZBTB38
ZNF699
AR
NR2F6
SCAND1
DRAP1
BUD31
RFXANK
TAF10
21
11
20
91
14
89
161
62
94
186
260
403
28
103
413
397
498
484
906
339

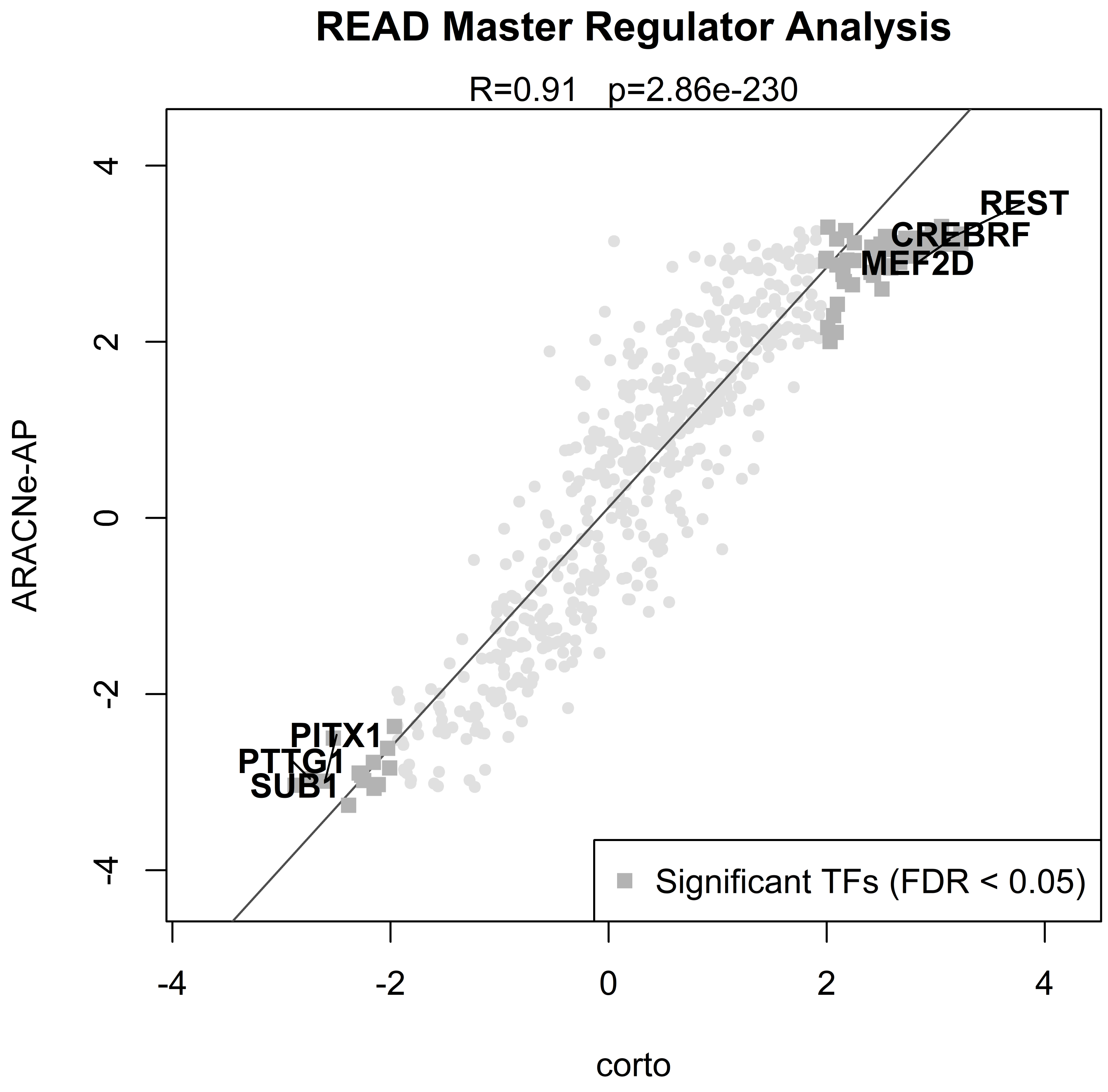
READ Master Regulator Analysis
R=0.91 p=2.86e-230
ARACNe-AP
corto
4
2
0
-2
-4
-4
-2
0
2
4
REST
CREBRF
MEF2D
PITX1
PTTG1
SUB1
Significant TFs (FDR < 0.05)

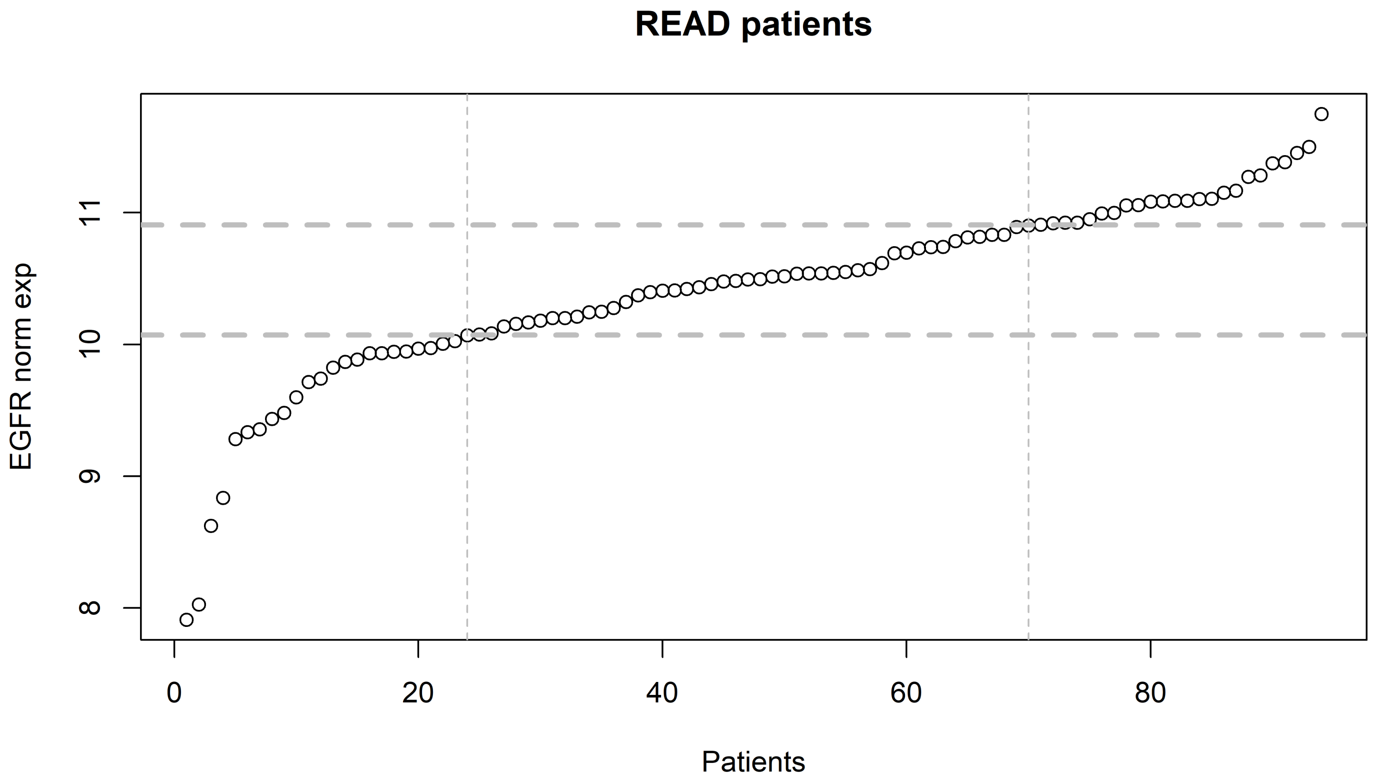

READ patients
EGFR norm exp
8
9
10
11
0
20
40
60
80
Patients

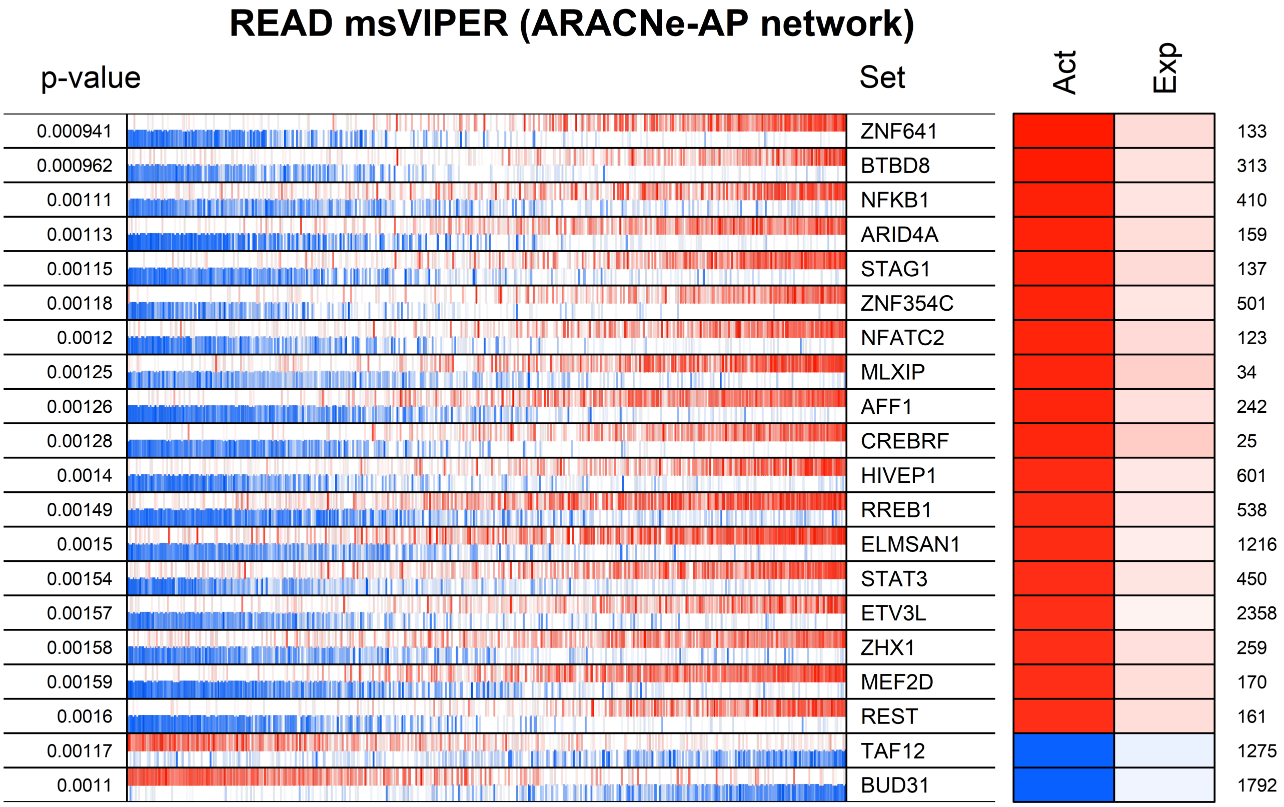
READ msVIPER (ARACNe-AP network)
p-value
Set
Act
Exp
0.000941
ZNF641
133
0.000962
BTBD8
313
0.00111
NFKB1
410
0.00113
ARID4A
159
0.00115
STAG1
137
0.00118
ZNF354C
501
0.0012
NFATC2
123
0.00125
MLXIP
34
0.00126
AFF1
242
0.00128
CREBRF
25
0.0014
HIVEP1
601
0.00149
RREB1
538
0.0015
ELMSAN1
1216
0.00154
STAT3
450
0.00157
ETV3L
2358
0.00158
ZHX1
259
0.00159
MEF2D
170
0.0016
REST
161
0.00117
TAF12
1275
0.0011
BUD31
1792

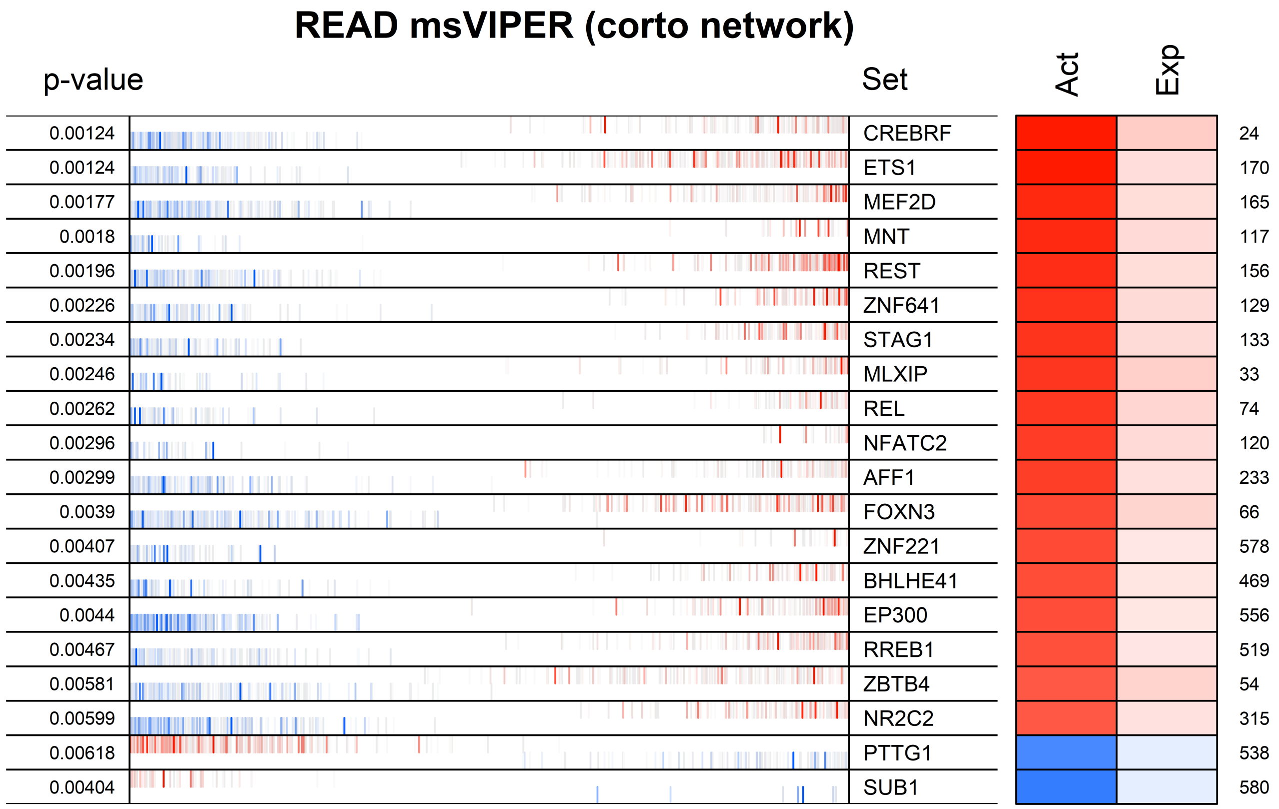
READ msVIPER (corto network)
p-value
Set
Act
Exp
0.00124 CREBRF 24
0.00124 ETS1 170
0.00177 MEF2D 165
0.0018 MNT 117
0.00196 REST 156
0.00226 ZNF641 129
0.00234 STAG1 133
0.00246 MLXIP 33
0.00262 REL 74
0.00296 NFATC2 120
0.00299 AFF1 233
0.0039 FOXN3 66
0.00407 ZNF221 578
0.00435 BHLHE41 469
0.0044 EP300 556
0.00467 RREB1 519
0.00581 ZBTB4 54
0.00599 NR2C2 315
0.00618 PTTG1 538
0.00404 SUB1 580

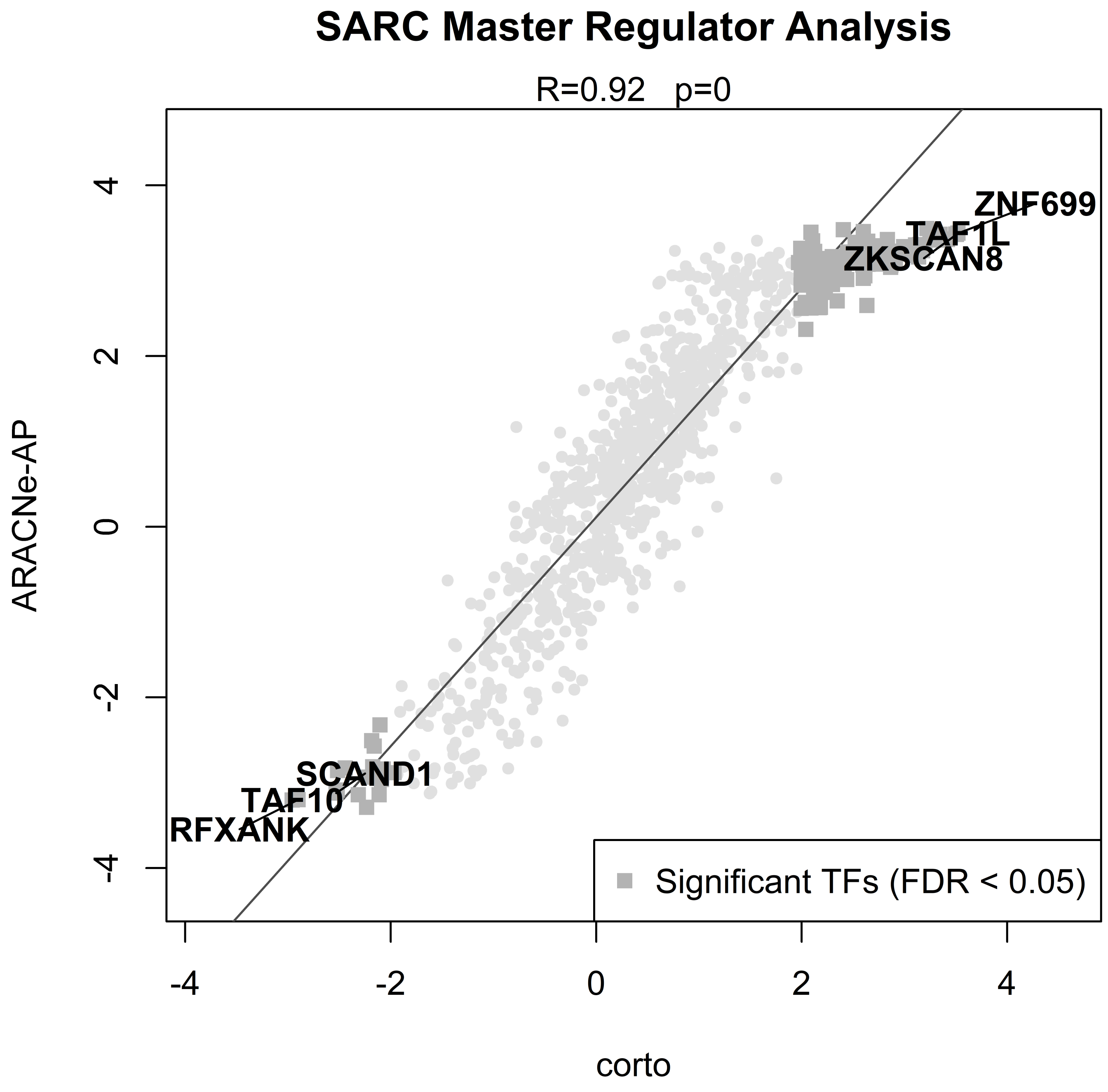
SARC Master Regulator Analysis
R=0.92 p=0
ZNF699
TAF1L
ZKSCAN8
SCAND1
TAF10
RFXANK
Significant TFs (FDR < 0.05)
ARACNe-AP
corto
4
2
0
-2
-4
-4
-2
0
2
4

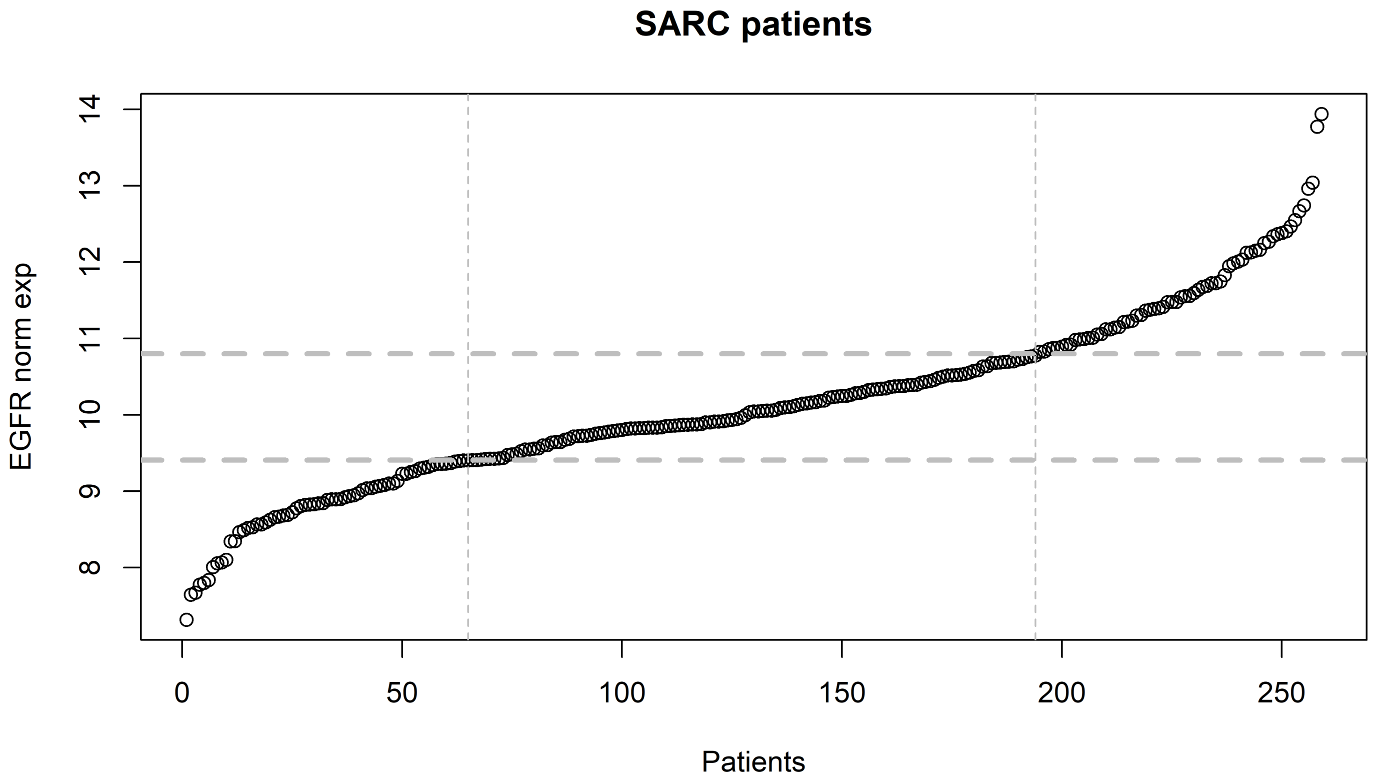

SARC patients
EGFR norm exp
14
13
12
11
10
9
8
0
50
100
150
200
250
Patients

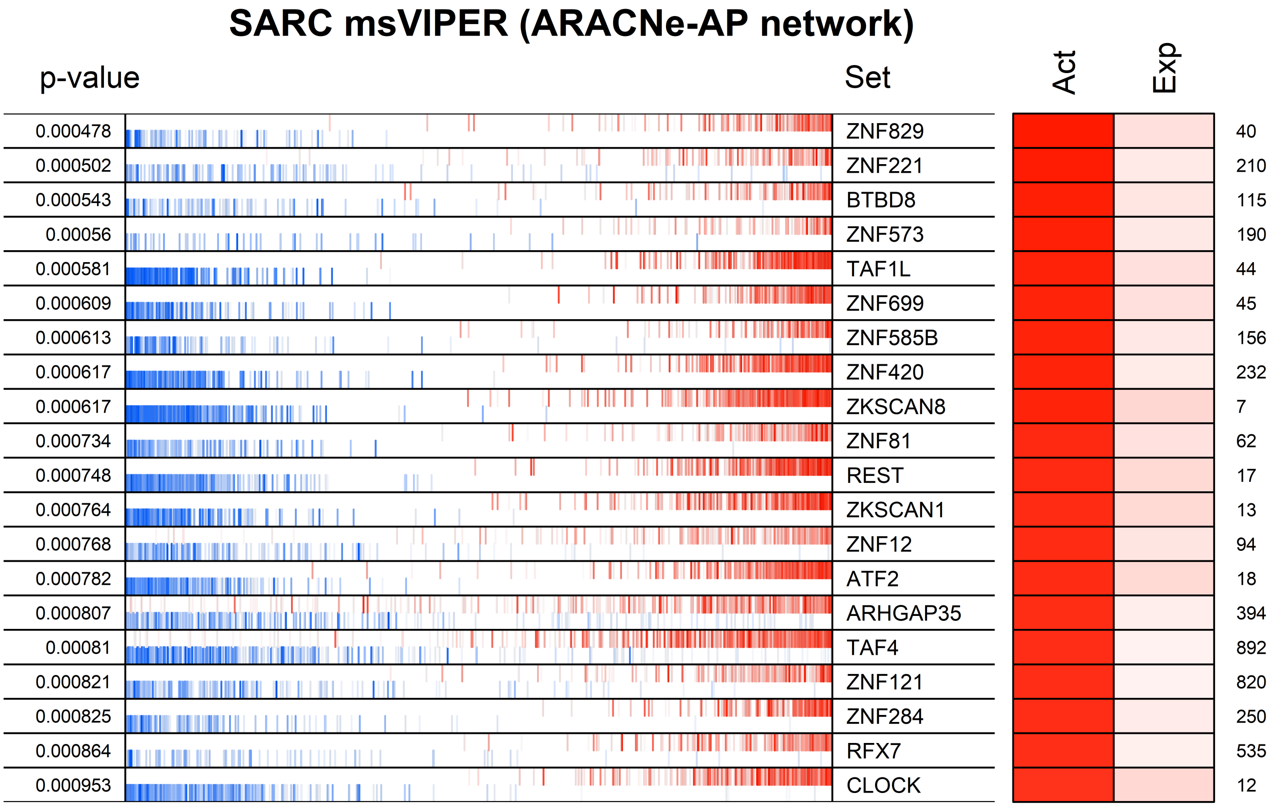
SARC msVIPER (ARACNe-AP network)
p-value
Set
Act
Exp
0.000478
ZNF829
40
0.000502
ZNF221
210
0.000543
BTBD8
115
0.00056
ZNF573
190
0.000581
TAF1L
44
0.000609
ZNF699
45
0.000613
ZNF585B
156
0.000617
ZNF420
232
0.000617
ZKSCAN8
7
0.000734
ZNF81
62
0.000748
REST
17
0.000764
ZKSCAN1
13
0.000768
ZNF12
94
0.000782
ATF2
18
0.000807
ARHGAP35
394
0.00081
TAF4
892
0.000821
ZNF121
820
0.000825
ZNF284
250
0.000864
RFX7
535
0.000953
CLOCK
12

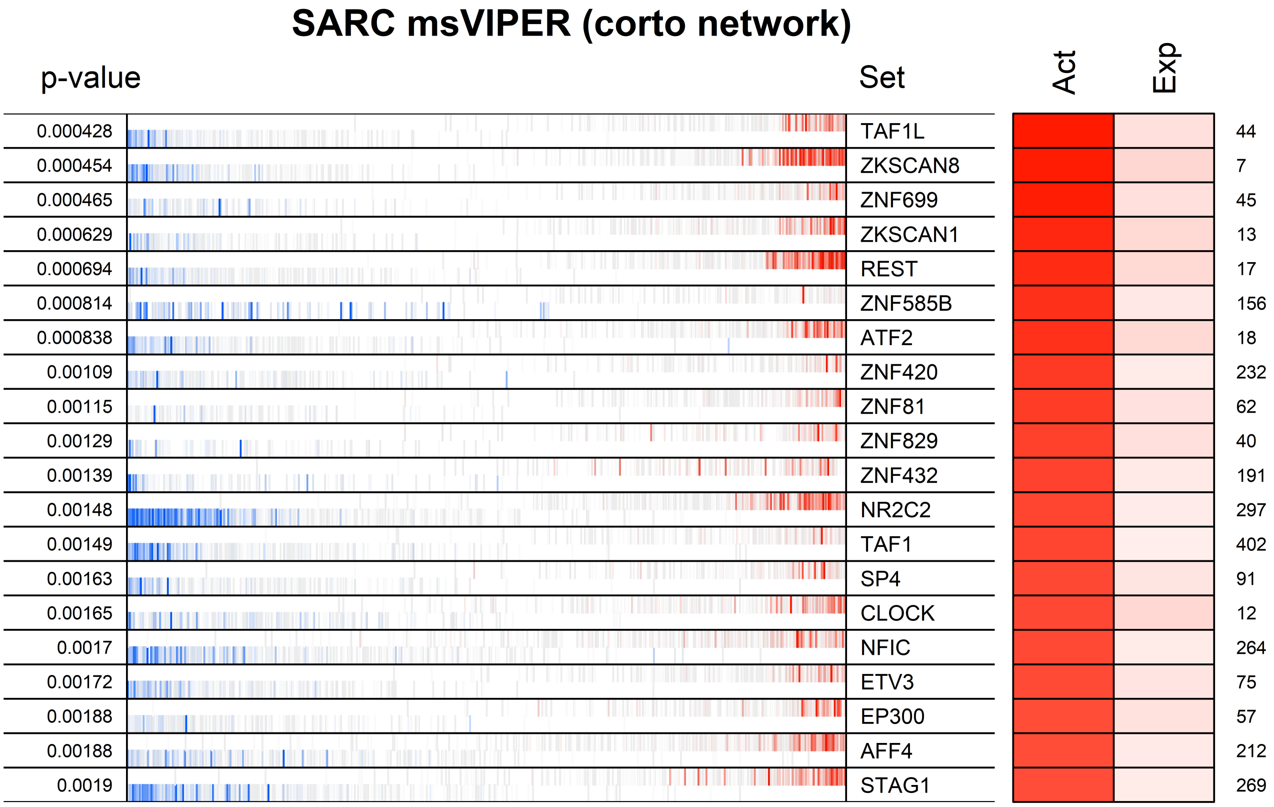
SARC msVIPER (corto network)
p-value
Set
Act
Exp
0.000428 TAF1L 44
0.000454 ZKSCAN8 7
0.000465 ZNF699 45
0.000629 ZKSCAN1 13
0.000694 REST 17
0.000814 ZNF585B 156
0.000838 ATF2 18
0.00109 ZNF420 232
0.00115 ZNF81 62
0.00129 ZNF829 40
0.00139 ZNF432 191
0.00148 NR2C2 297
0.00149 TAF1 402
0.00163 SP4 91
0.00165 CLOCK 12
0.0017 NFIC 264
0.00172 ETV3 75
0.00188 EP300 57
0.00188 AFF4 212
0.0019 STAG1 269

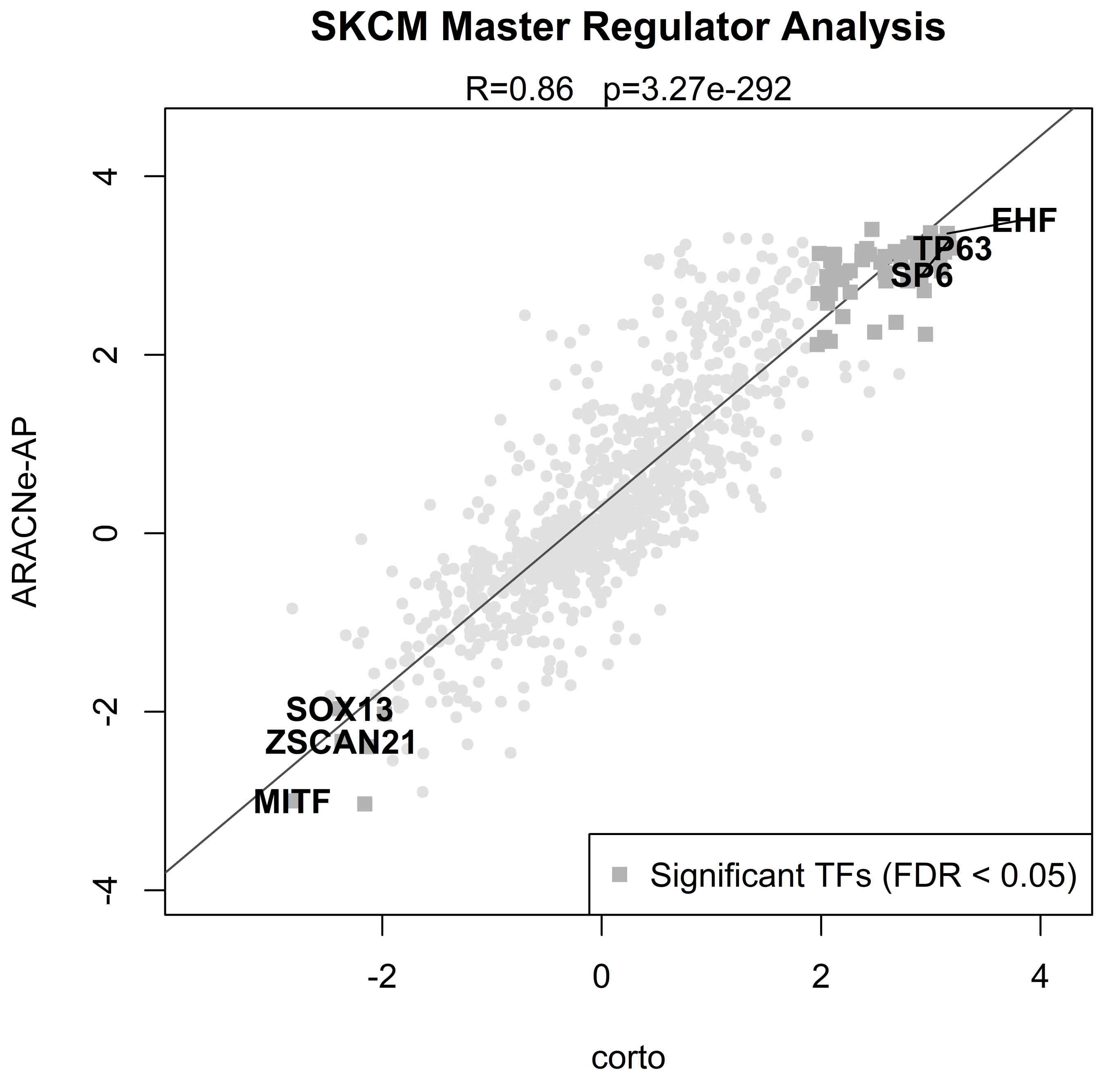
SKCM Master Regulator Analysis
R=0.86 p=3.27e-292
ARACNe-AP
corto
EHF
TP63
SP6
SOX13
ZSCAN21
MITF
Significant TFs (FDR < 0.05)
4
2
0
-2
-4
-2
0
2
4

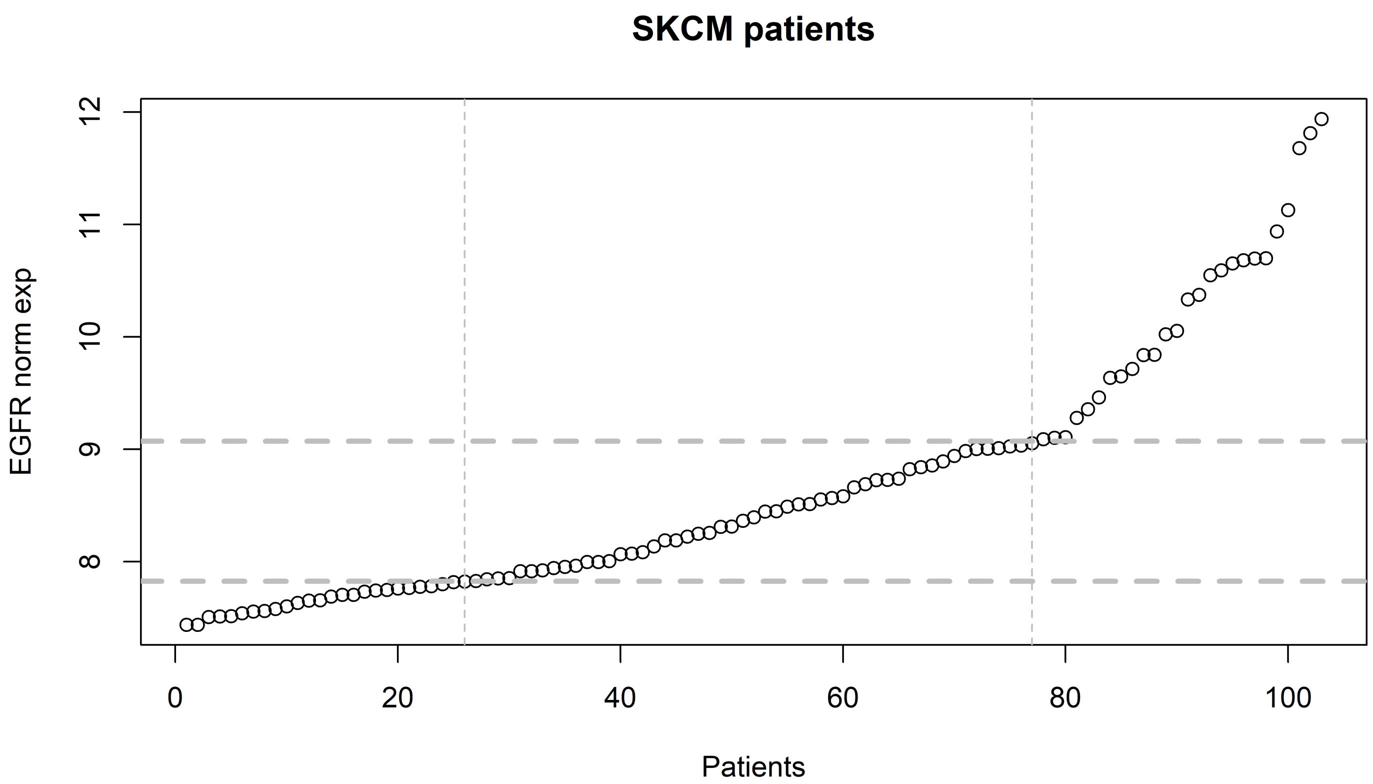
SKCM patients
EGFR norm exp
Patients
8
9
10
11
12
0
20
40
60
80
100

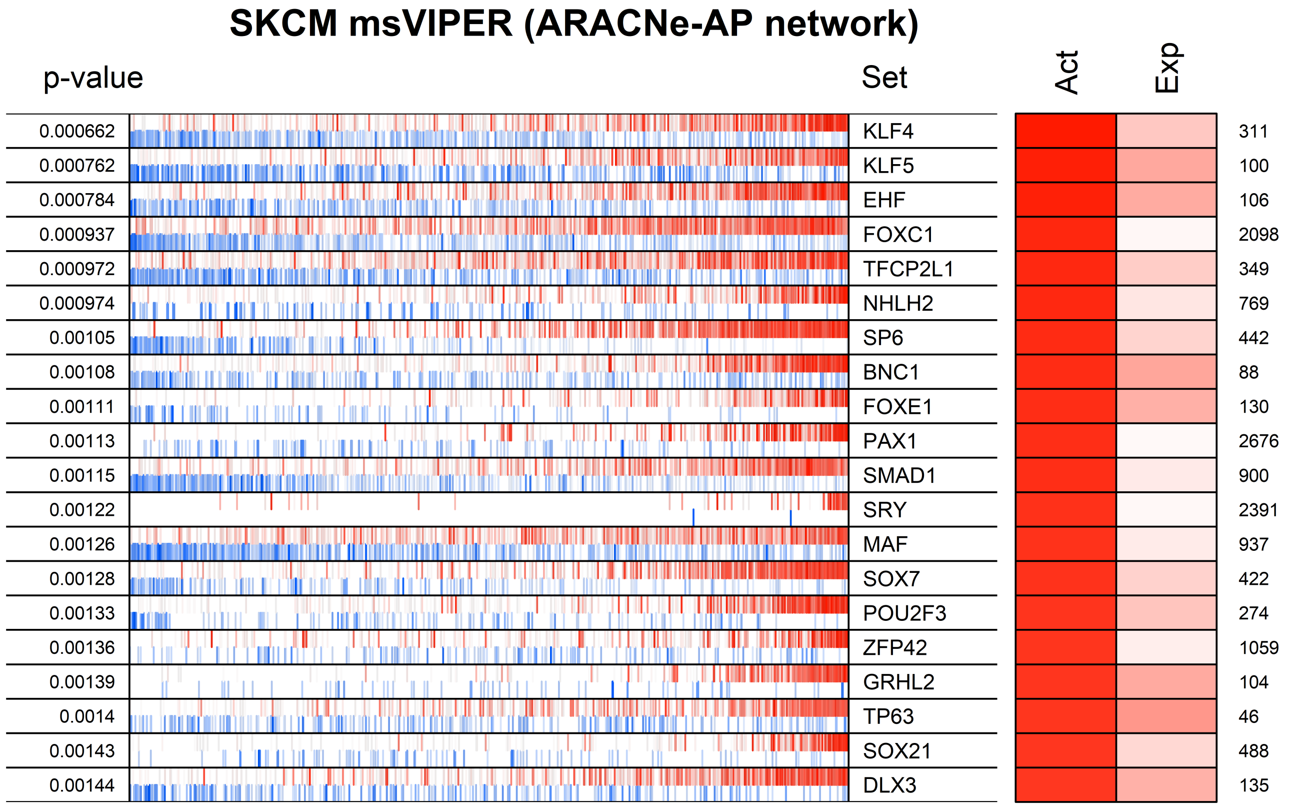
SKCM msVIPER (ARACNe-AP network)
p-value
Set
Act
Exp
0.000662 KLF4 311
0.000762 KLF5 100
0.000784 EHF 106
0.000937 FOXC1 2098
0.000972 TFCP2L1 349
0.000974 NHLH2 769
0.00105 SP6 442
0.00108 BNC1 88
0.00111 FOXE1 130
0.00113 PAX1 2676
0.00115 SMAD1 900
0.00122 SRY 2391
0.00126 MAF 937
0.00128 SOX7 422
0.00133 POU2F3 274
0.00136 ZFP42 1059
0.00139 GRHL2 104
0.0014 TP63 46
0.00143 SOX21 488
0.00144 DLX3 135

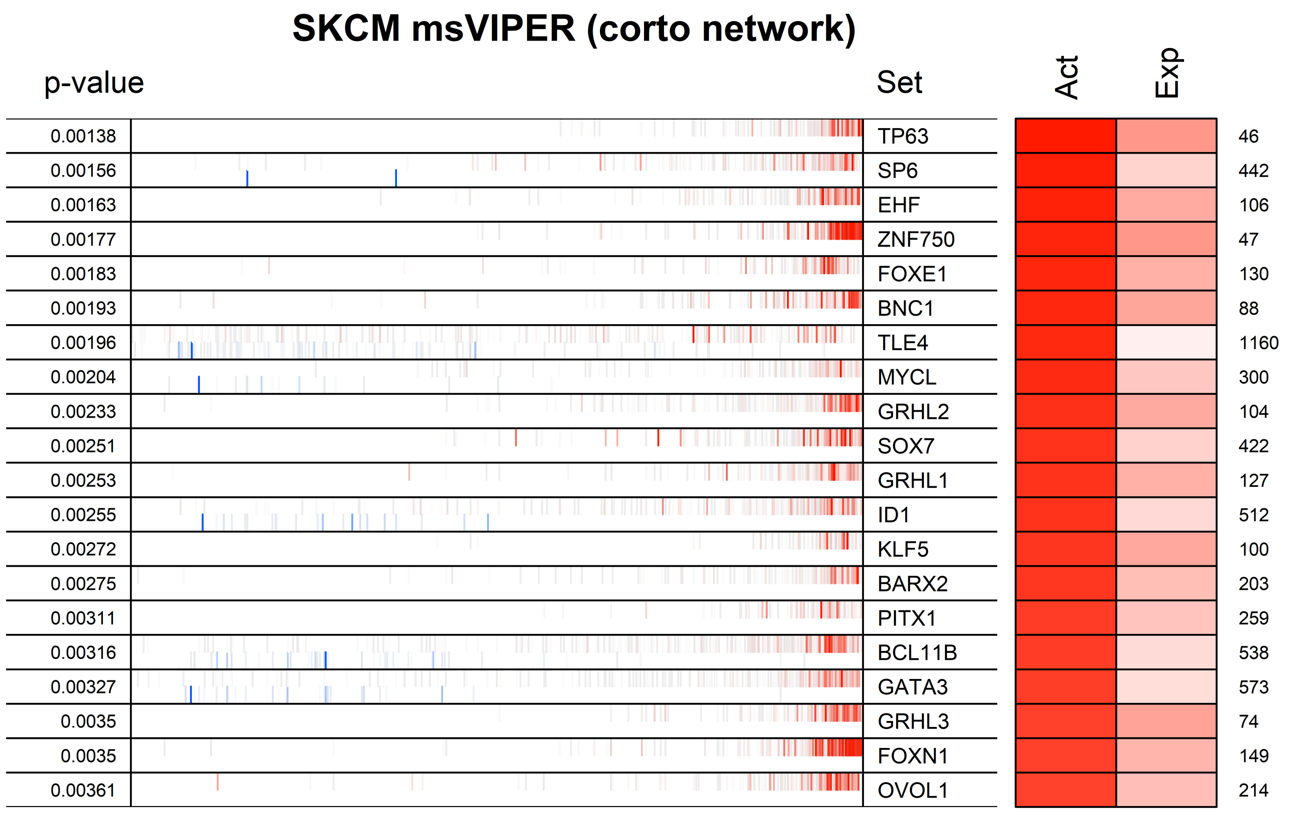
SKCM msVIPER (corto network)
p-value
Set
Act
Exp
0.00138 TP63 46
0.00156 SP6 442
0.00163 EHF 106
0.00177 ZNF750 47
0.00183 FOXE1 130
0.00193 BNC1 88
0.00196 TLE4 1160
0.00204 MYCL 300
0.00233 GRHL2 104
0.00251 SOX7 422
0.00253 GRHL1 127
0.00255 ID1 512
0.00272 KLF5 100
0.00275 BARX2 203
0.00311 PITX1 259
0.00316 BCL11B 538
0.00327 GATA3 573
0.0035 GRHL3 74
0.0035 FOXN1 149
0.00361 OVOL1 214

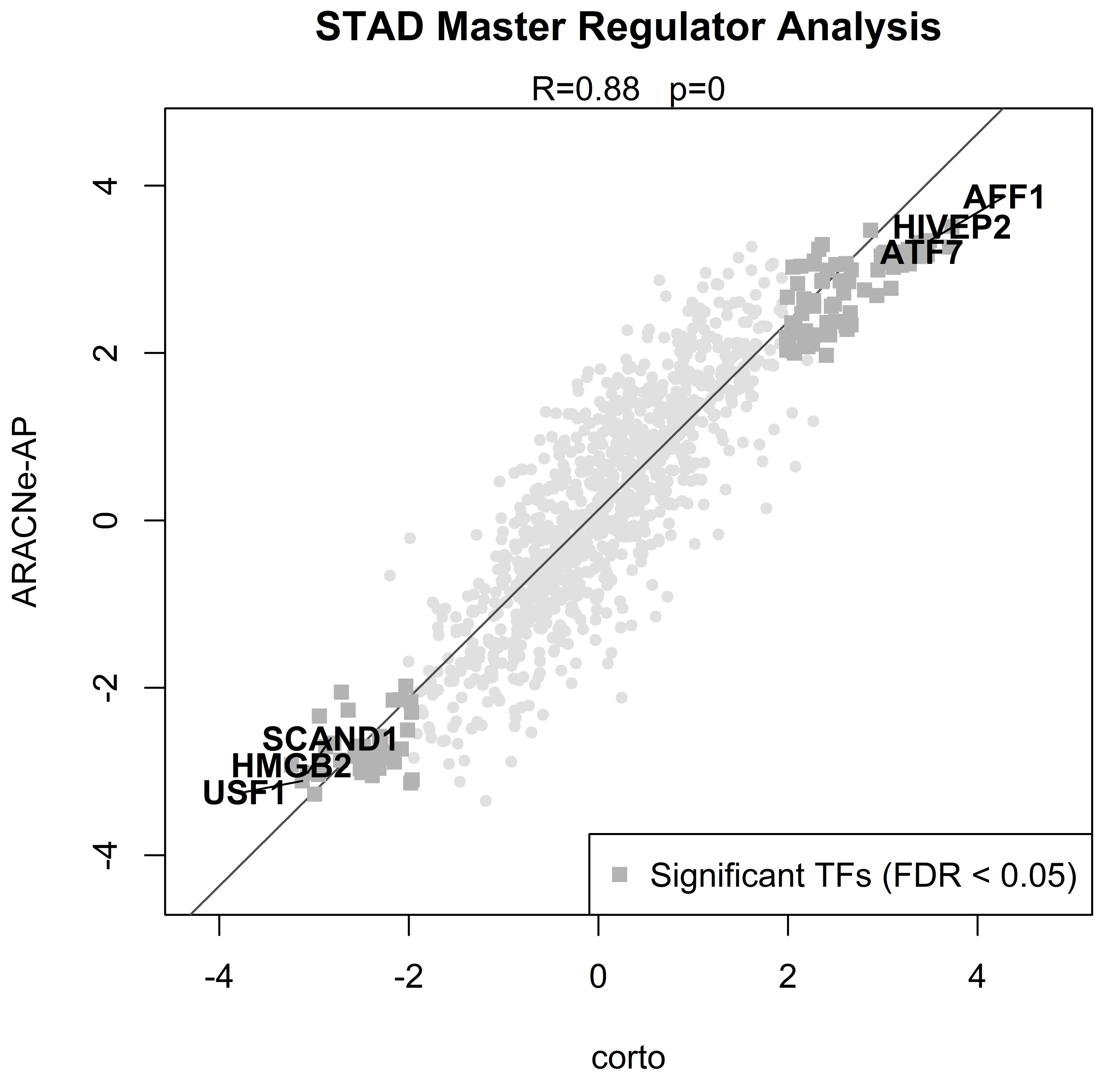
STAD Master Regulator Analysis
R=0.88 p=0
ARACNe-AP
corto
AFF1
HIVEP2
ATF7
SCAND1
HMGB2
USF1
Significant TFs (FDR < 0.05)
4
2
0
-2
-4
-4
-2
0
2
4

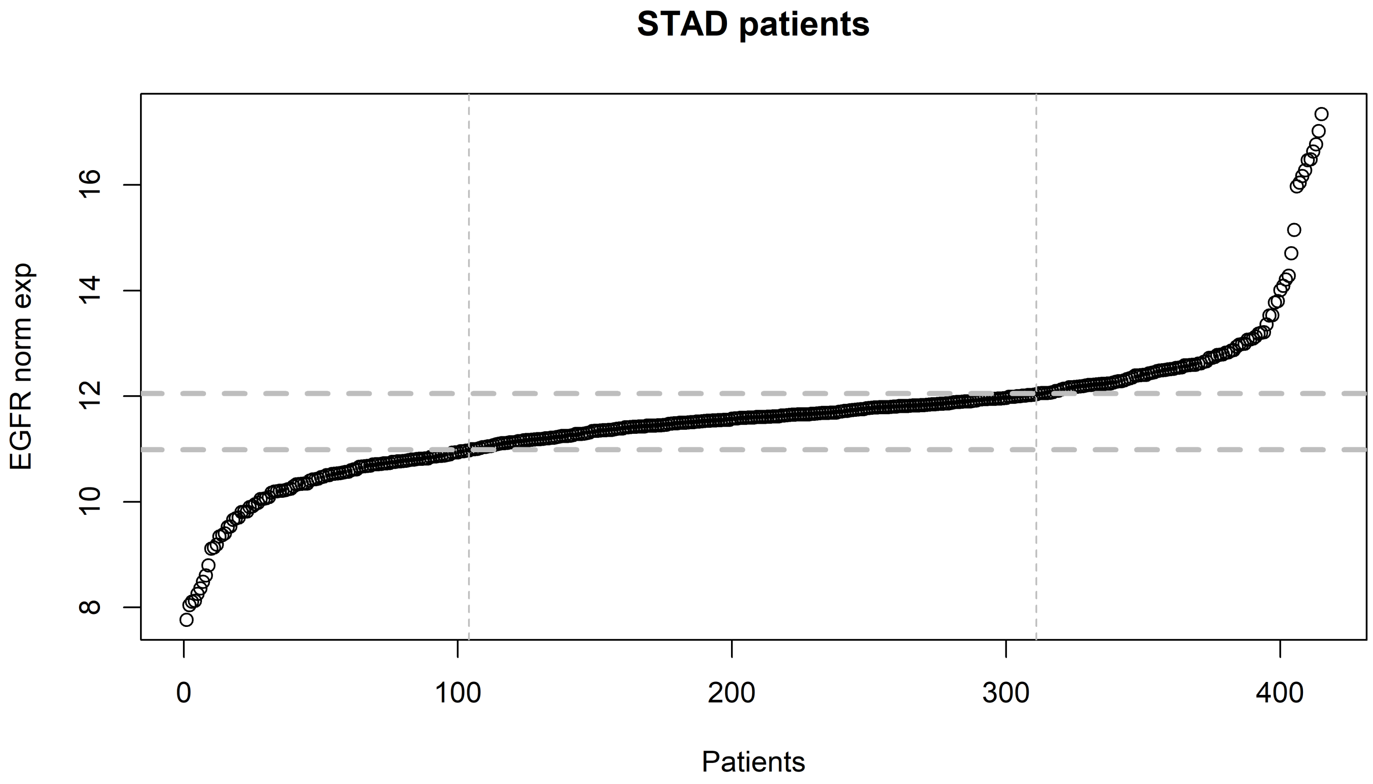
STAD patients
EGFR norm exp
8
10
12
14
16
0
100
200
300
400
Patients

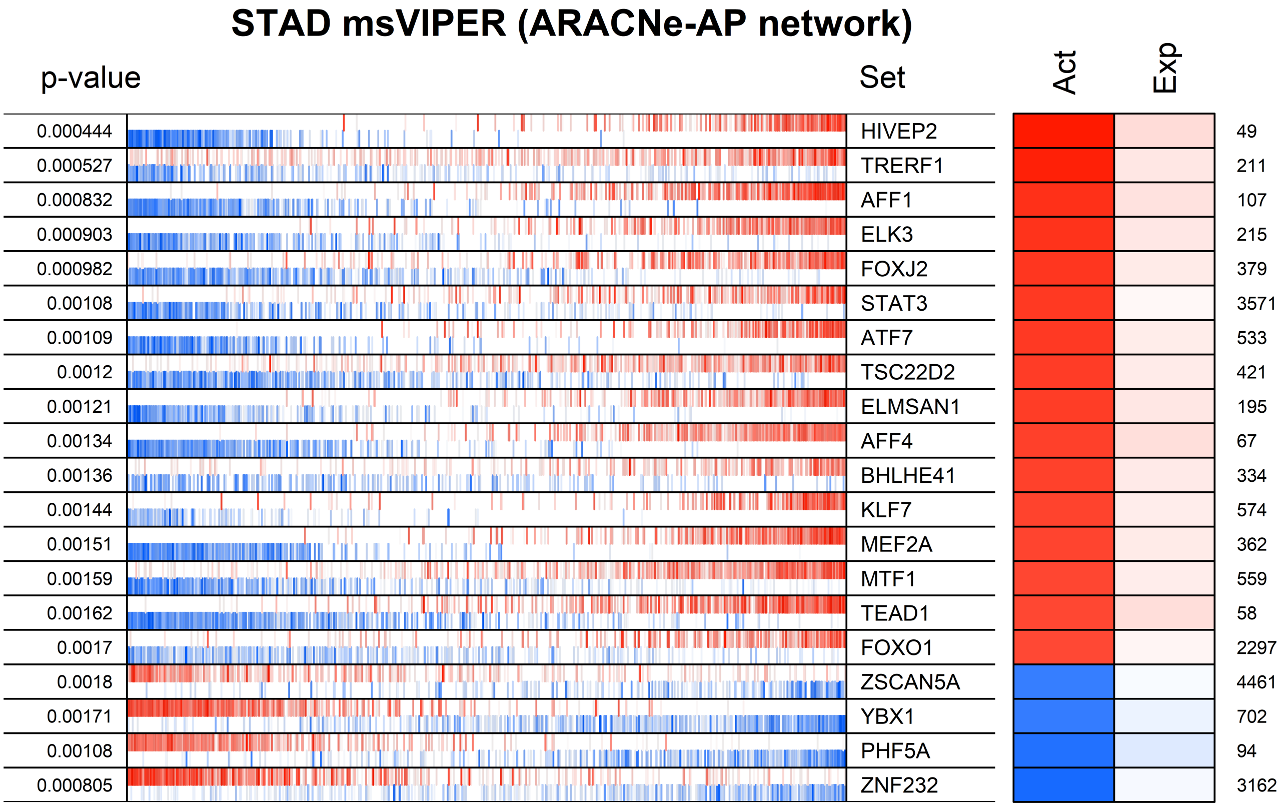
STAD msVIPER (ARACNe-AP network)
p-value
Set
Act
Exp
0.000444 HIVEP2 49
0.000527 TRERF1 211
0.000832 AFF1 107
0.000903 ELK3 215
0.000982 FOXJ2 379
0.00108 STAT3 3571
0.00109 ATF7 533
0.0012 TSC22D2 421
0.00121 ELMSAN1 195
0.00134 AFF4 67
0.00136 BHLHE41 334
0.00144 KLF7 574
0.00151 MEF2A 362
0.00159 MTF1 559
0.00162 TEAD1 58
0.0017 FOXO1 2297
0.0018 ZSCAN5A 4461
0.00171 YBX1 702
0.00108 PHF5A 94
0.000805 ZNF232 3162

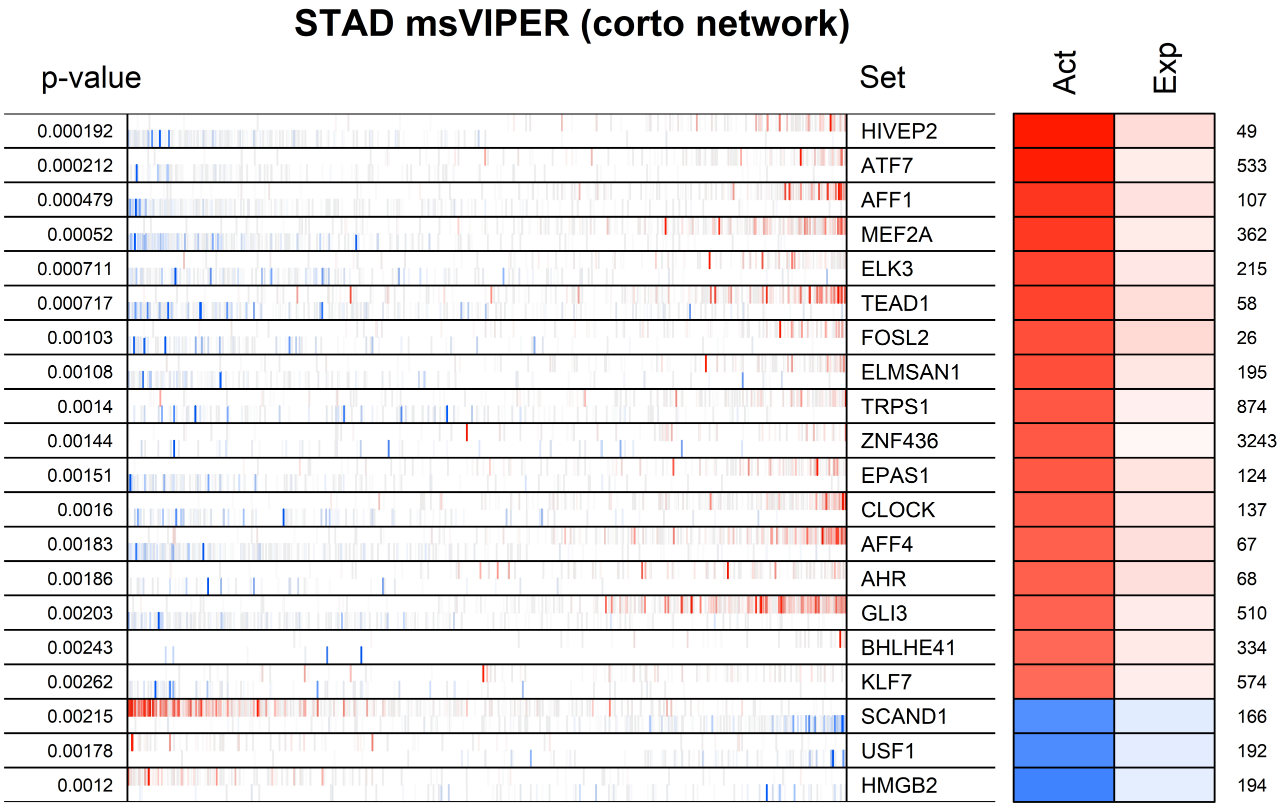
STAD msVIPER (corto network)
p-value
Set
Act
Exp
0.000192
HIVEP2
49
0.000212
ATF7
533
0.000479
AFF1
107
0.00052
MEF2A
362
0.000711
ELK3
215
0.000717
TEAD1
58
0.00103
FOSL2
26
0.00108
ELMSAN1
195
0.0014
TRPS1
874
0.00144
ZNF436
3243
0.00151
EPAS1
124
0.0016
CLOCK
137
0.00183
AFF4
67
0.00186
AHR
68
0.00203
GLI3
510
0.00243
BHLHE41
334
0.00262
KLF7
574
0.00215
SCAND1
166
0.00178
USF1
192
0.0012
HMGB2
194

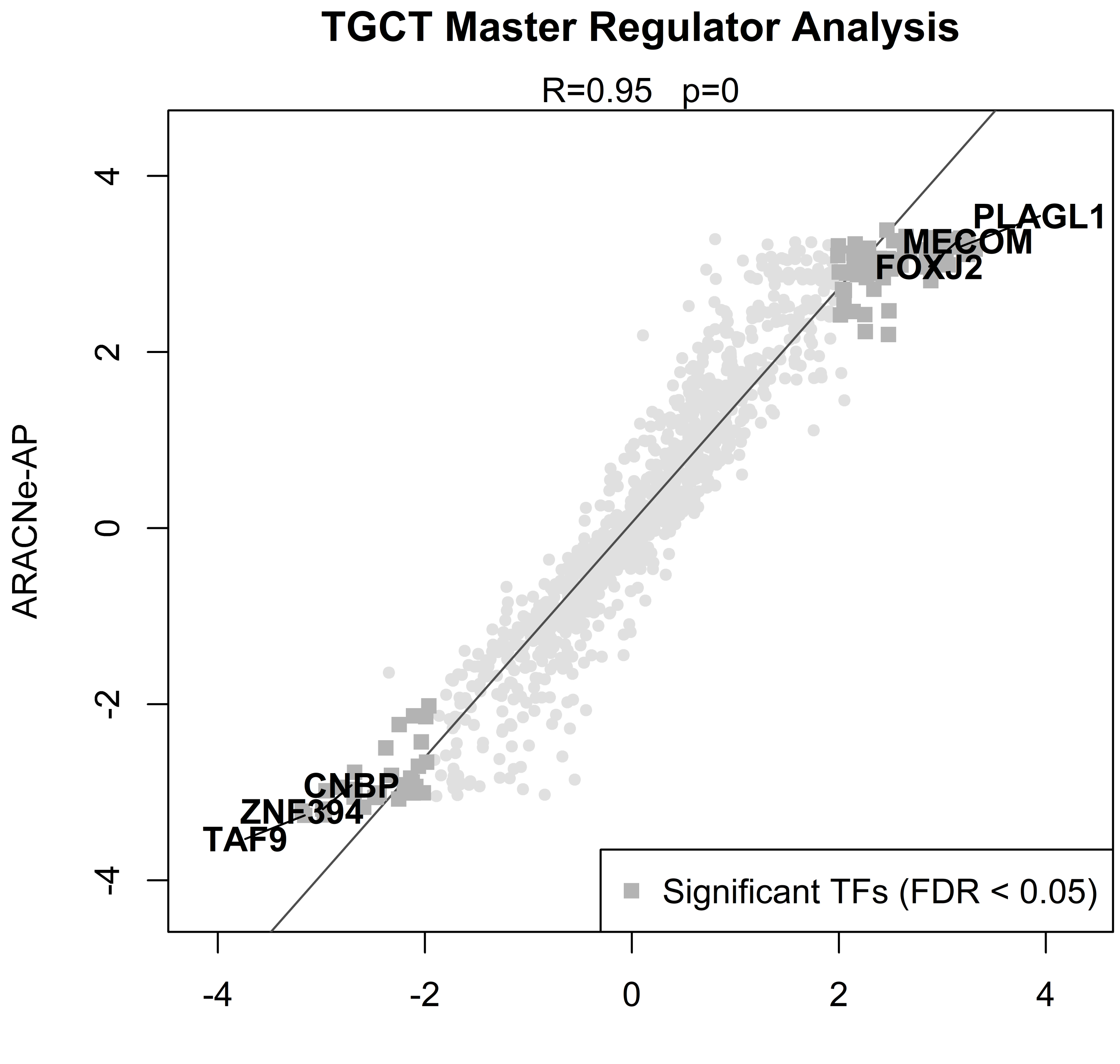
TGCT Master Regulator Analysis
R=0.95 p=0
ARACNe-AP
corto
-4
-2
0
2
4
PLAGL1
MECOM
FOXJ2
CNBP
ZNF394
TAF9
Significant TFs (FDR < 0.05)

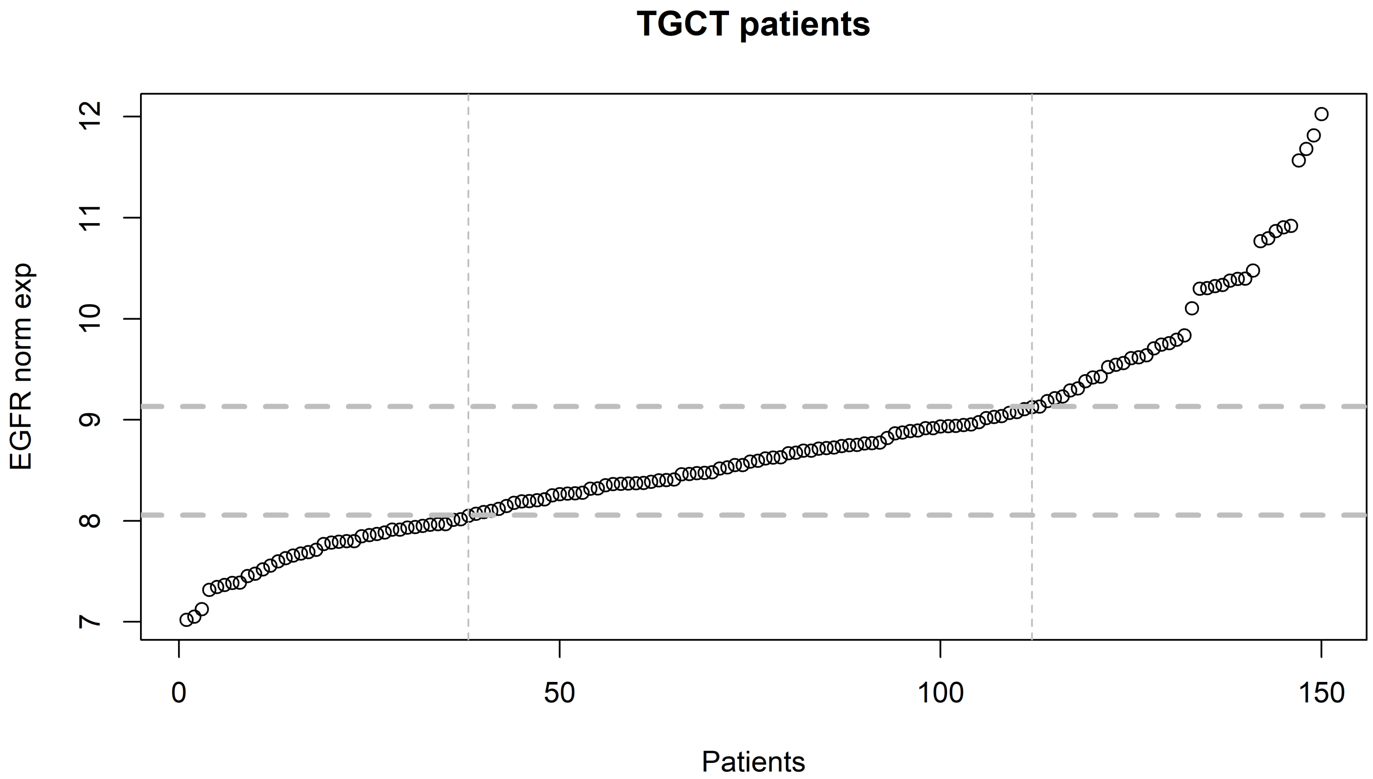

TGCT patients
EGFR norm exp
Patients
7
8
9
10
11
12
0
50
100
150

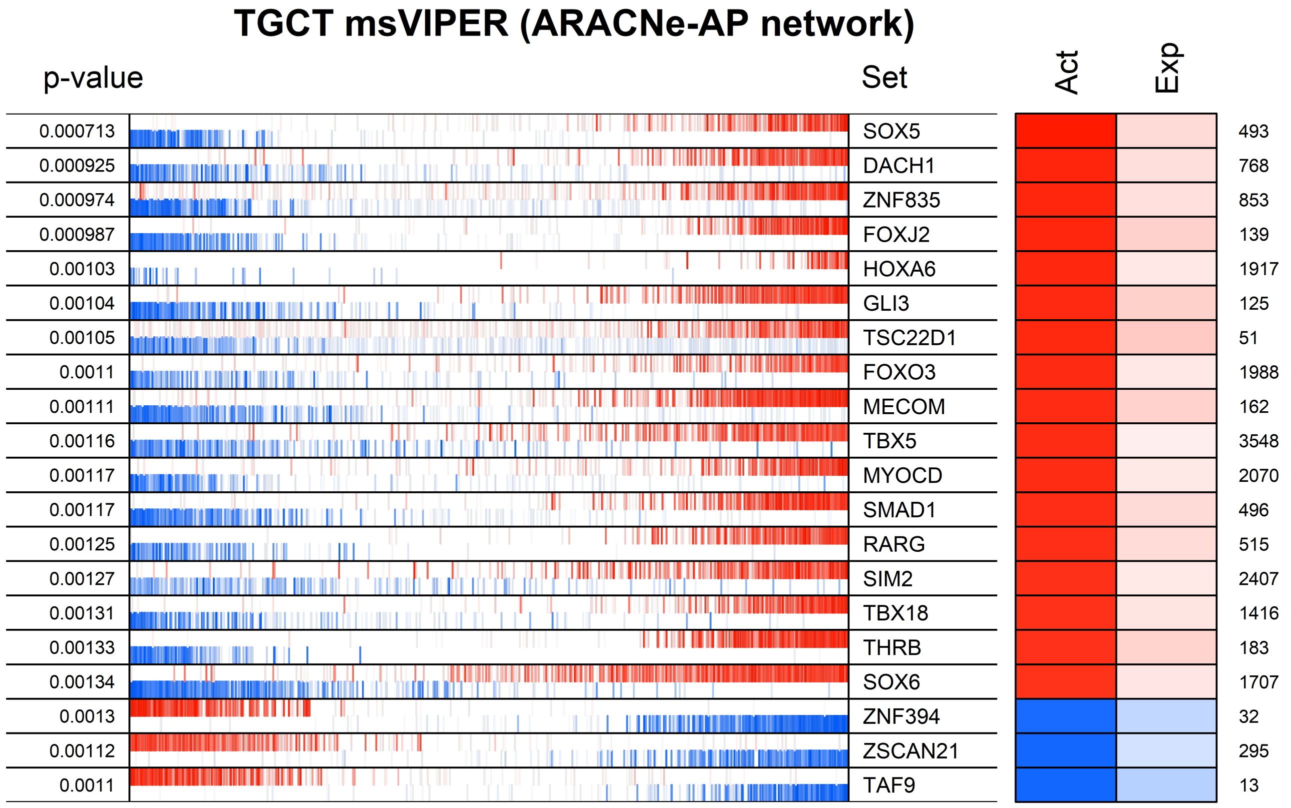

TGCT msVIPER (ARACNe-AP network)
p-value
Set
Act
Exp
0.000713 SOX5 493
0.000925 DACH1 768
0.000974 ZNF835 853
0.000987 FOXJ2 139
0.00103 HOXA6 1917
0.00104 GLI3 125
0.00105 TSC22D1 51
0.0011 FOXO3 1988
0.00111 MECOM 162
0.00116 TBX5 3548
0.00117 MYOCD 2070
0.00117 SMAD1 496
0.00125 RARG 515
0.00127 SIM2 2407
0.00131 TBX18 1416
0.00133 THRB 183
0.00134 SOX6 1707
0.0013 ZNF394 32
0.00112 ZSCAN21 295
0.0011 TAF9 13

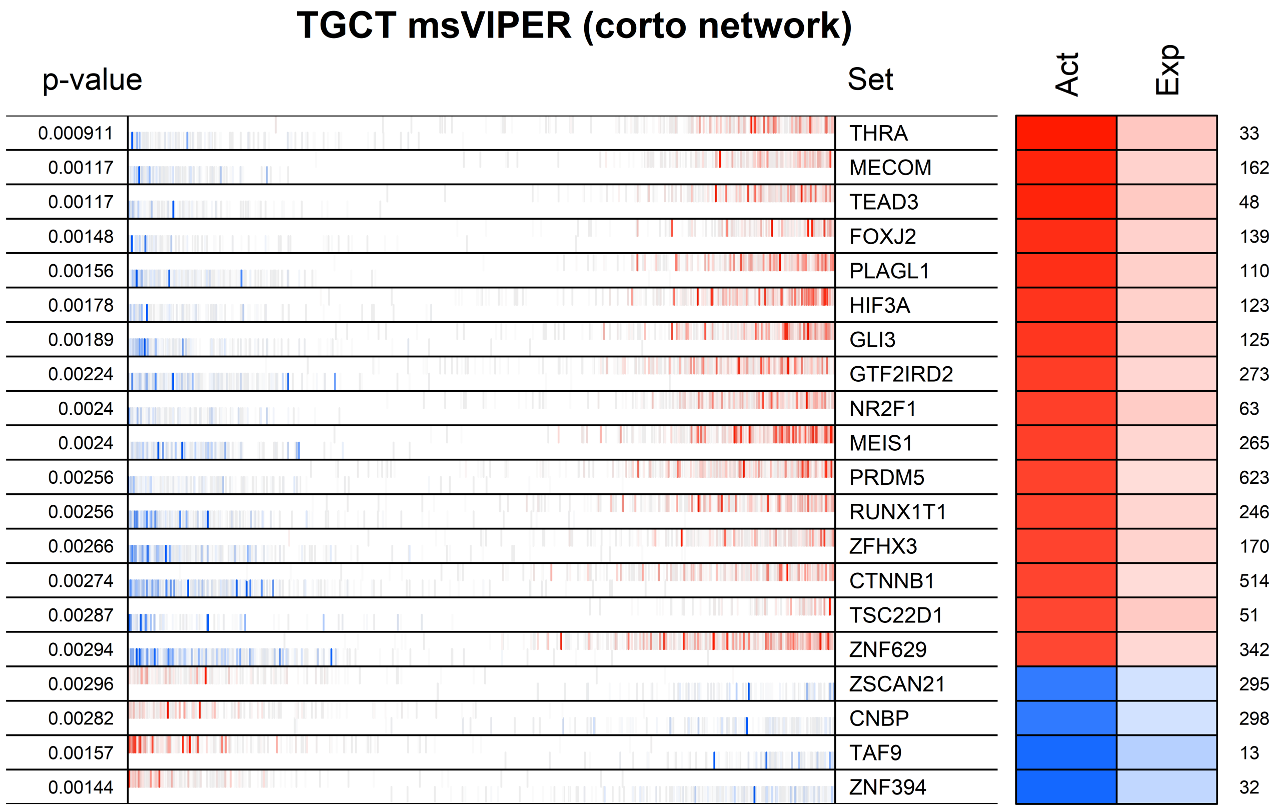
TGCT msVIPER (corto network)
p-value
Set
Act
Exp
0.000911 THRA 33
0.00117 MECOM 162
0.00117 TEAD3 48
0.00148 FOXJ2 139
0.00156 PLAGL1 110
0.00178 HIF3A 123
0.00189 GLI3 125
0.00224 GTF2IRD2 273
0.0024 NR2F1 63
0.0024 MEIS1 265
0.00256 PRDM5 623
0.00256 RUNX1T1 246
0.00266 ZFHX3 170
0.00274 CTNNB1 514
0.00287 TSC22D1 51
0.00294 ZNF629 342
0.00296 ZSCAN21 295
0.00282 CNBP 298
0.00157 TAF9 13
0.00144 ZNF394 32

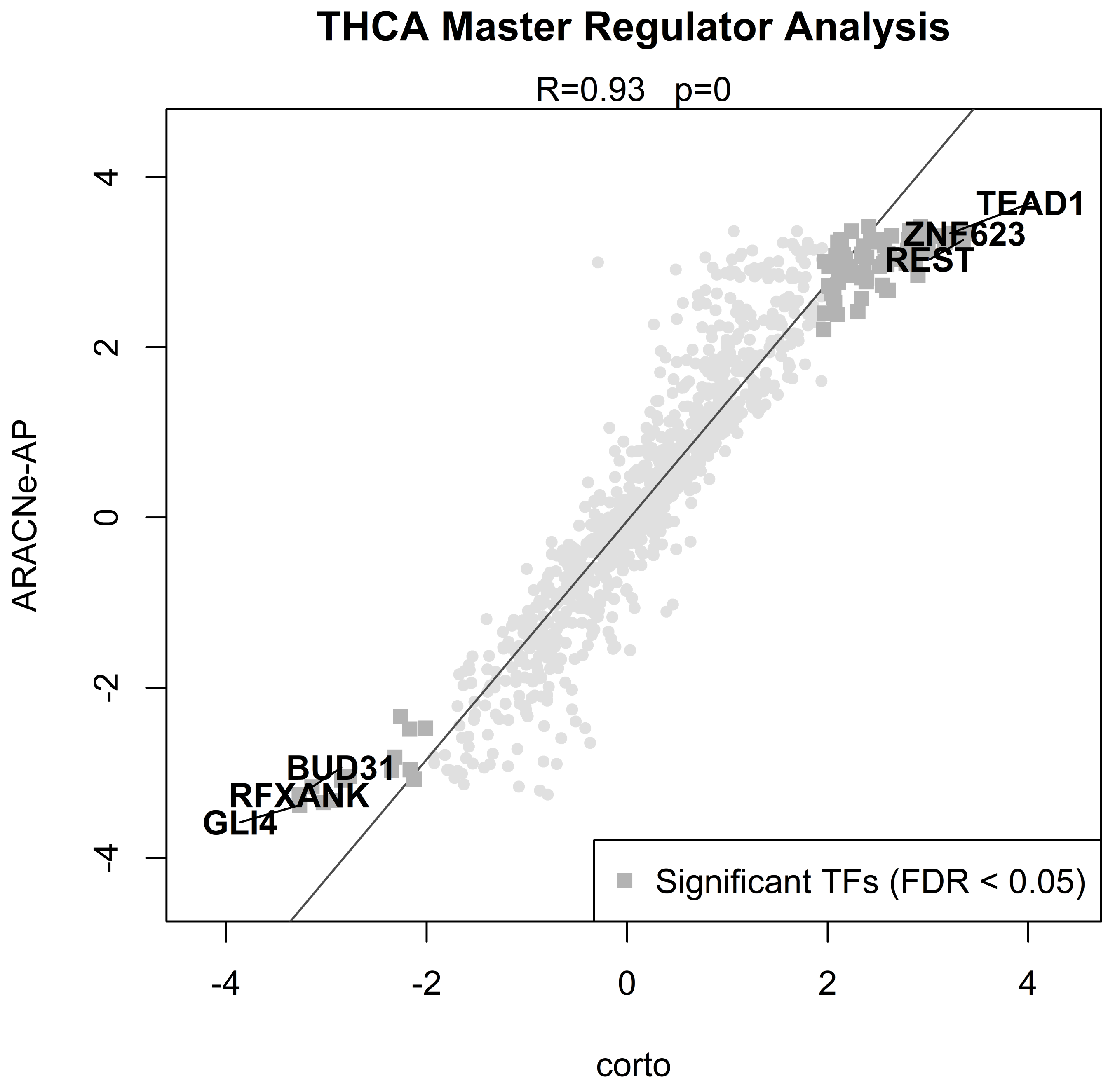
THCA Master Regulator Analysis
R=0.93 p=0
ARACNe-AP
corto
4
2
0
-2
-4
TEAD1
ZNF623
REST
BUD31
RFXANK
GLI4
Significant TFs (FDR < 0.05)

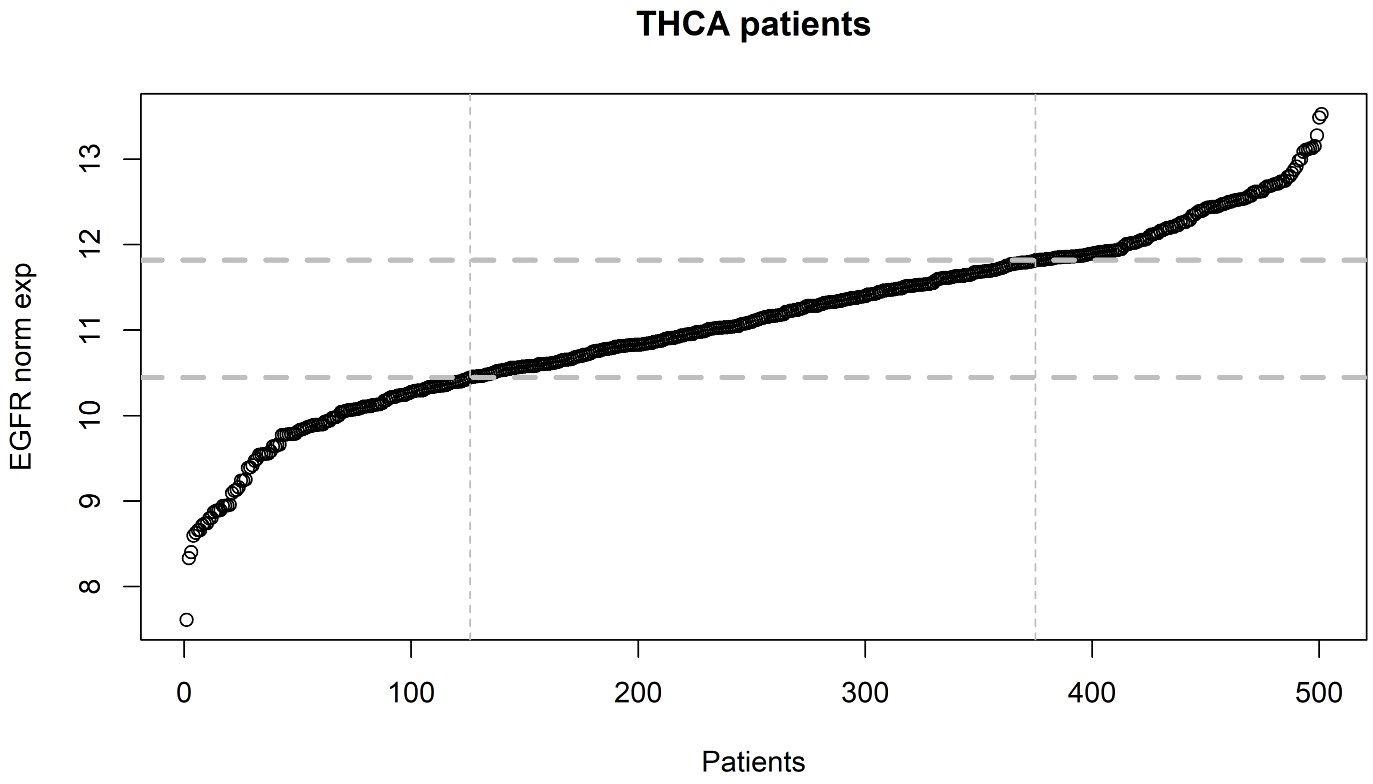
THCA patients
EGFR norm exp
Patients
8
9
10
11
12
13
0
100
200
300
400
500

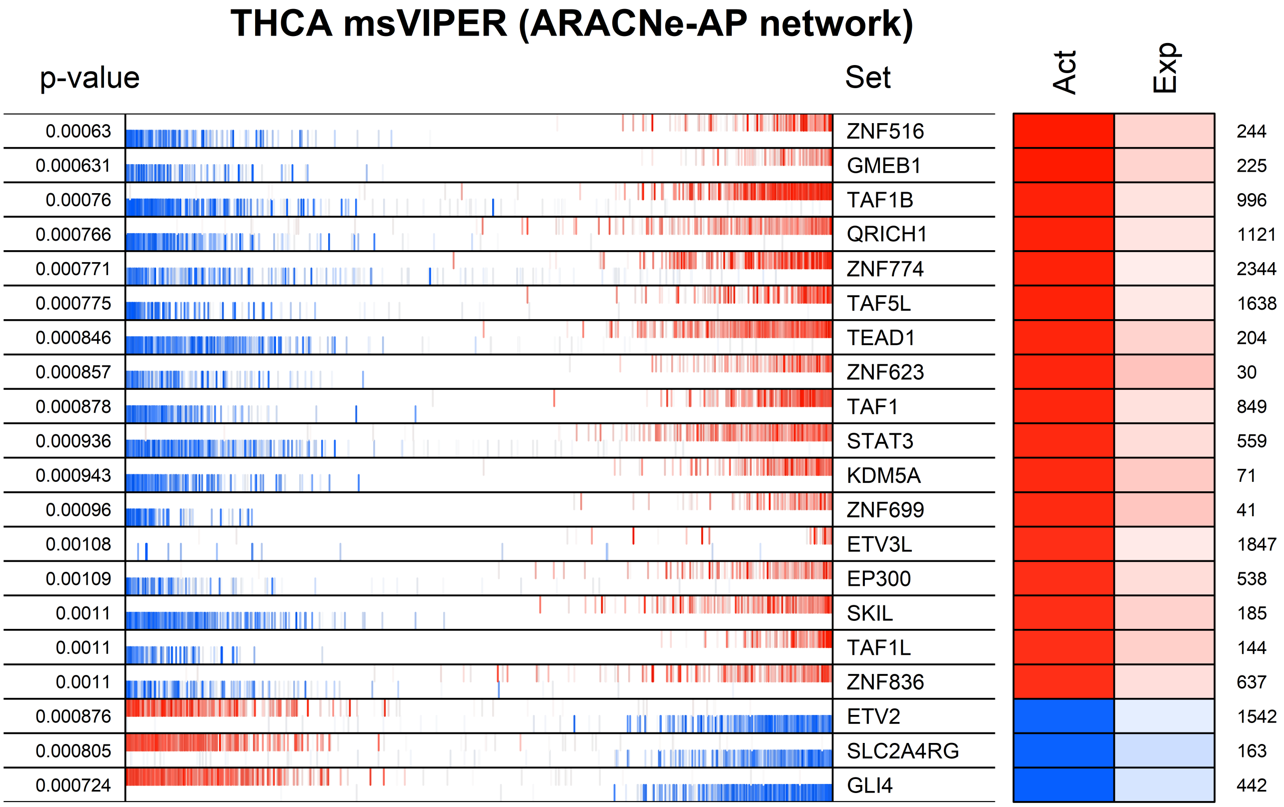
THCA msVIPER (ARACNe-AP network)
p-value
Set
Act
Exp
0.00063
ZNF516
244
0.000631
GMEB1
225
0.00076
TAF1B
996
0.000766
QRICH1
1121
0.000771
ZNF774
2344
0.000775
TAF5L
1638
0.000846
TEAD1
204
0.000857
ZNF623
30
0.000878
TAF1
849
0.000936
STAT3
559
0.000943
KDM5A
71
0.00096
ZNF699
41
0.00108
ETV3L
1847
0.00109
EP300
538
0.0011
SKIL
185
0.0011
TAF1L
144
0.0011
ZNF836
637
0.000876
ETV2
1542
0.000805
SLC2A4RG
163
0.000724
GLI4
442

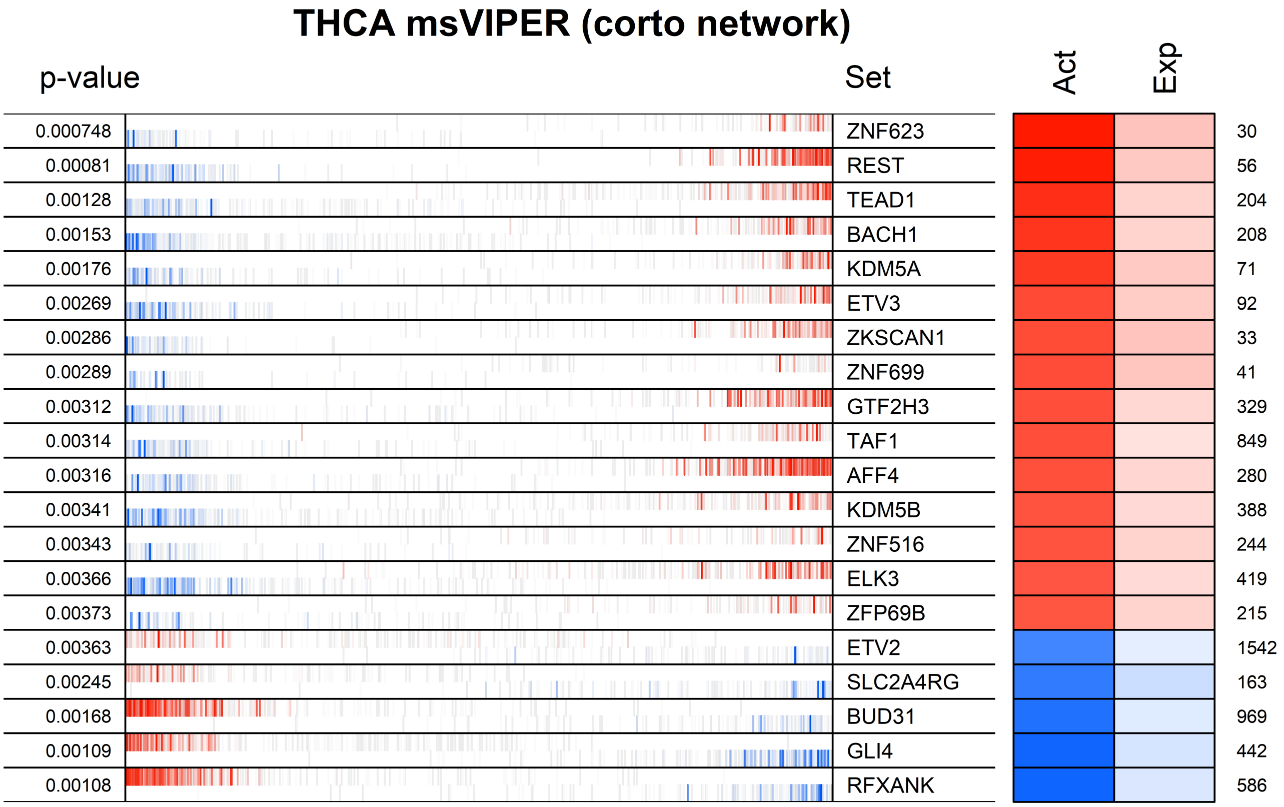
THCA msVIPER (corto network)
p-value
Set
Act
Exp
0.000748 ZNF623 30
0.00081 REST 56
0.00128 TEAD1 204
0.00153 BACH1 208
0.00176 KDM5A 71
0.00269 ETV3 92
0.00286 ZKSCAN1 33
0.00289 ZNF699 41
0.00312 GTF2H3 329
0.00314 TAF1 849
0.00316 AFF4 280
0.00341 KDM5B 388
0.00343 ZNF516 244
0.00366 ELK3 419
0.00373 ZFP69B 215
0.00363 ETV2 1542
0.00245 SLC2A4RG 163
0.00168 BUD31 969
0.00109 GLI4 442
0.00108 RFXANK 586

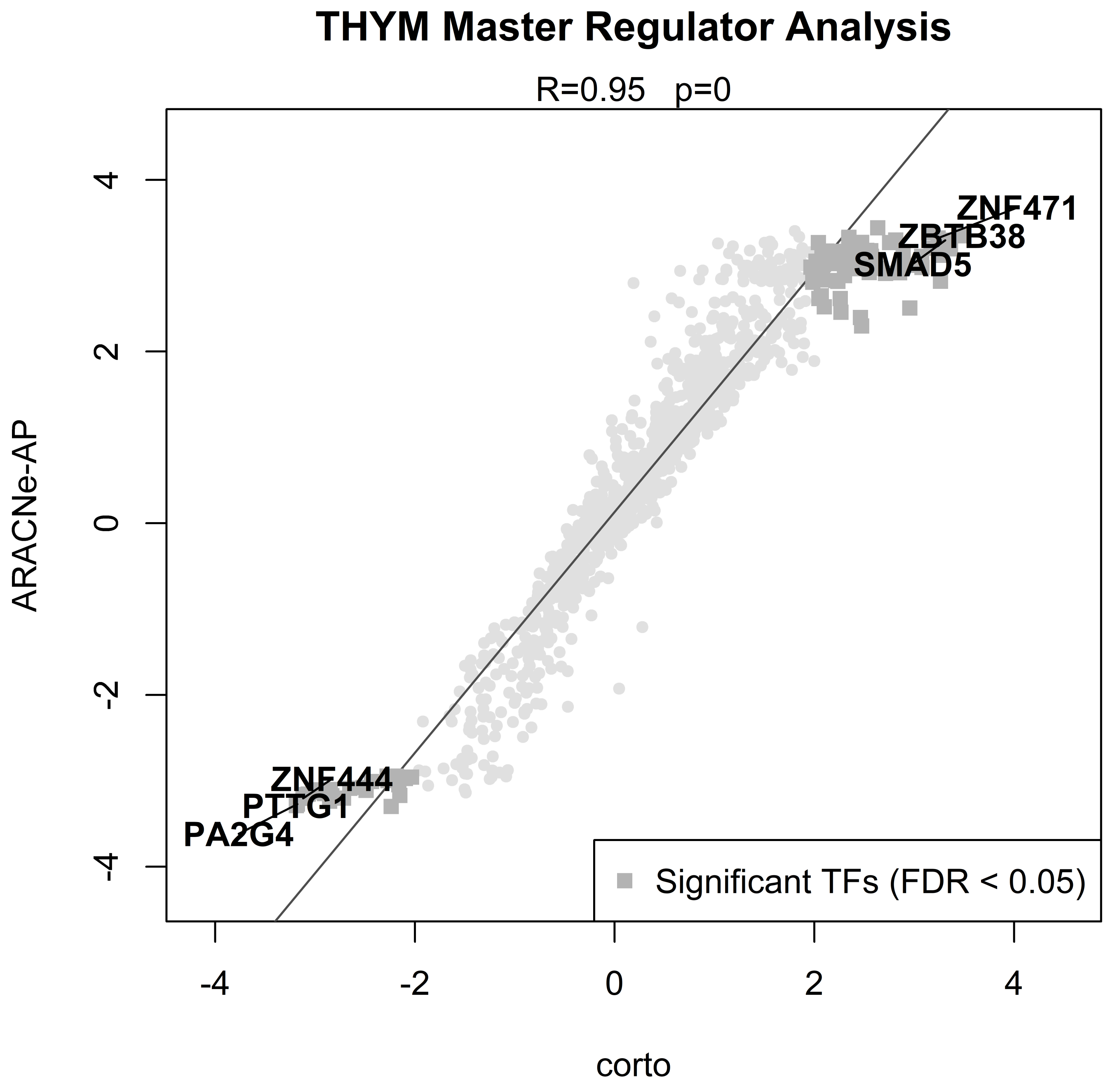

THYM Master Regulator Analysis
R=0.95 p=0
ARACNe-AP
corto
4
2
0
-2
-4
-4
-2
0
2
4
ZNF471
ZBTB38
SMAD5
ZNF444
PTTG1
PA2G4
Significant TFs (FDR < 0.05)

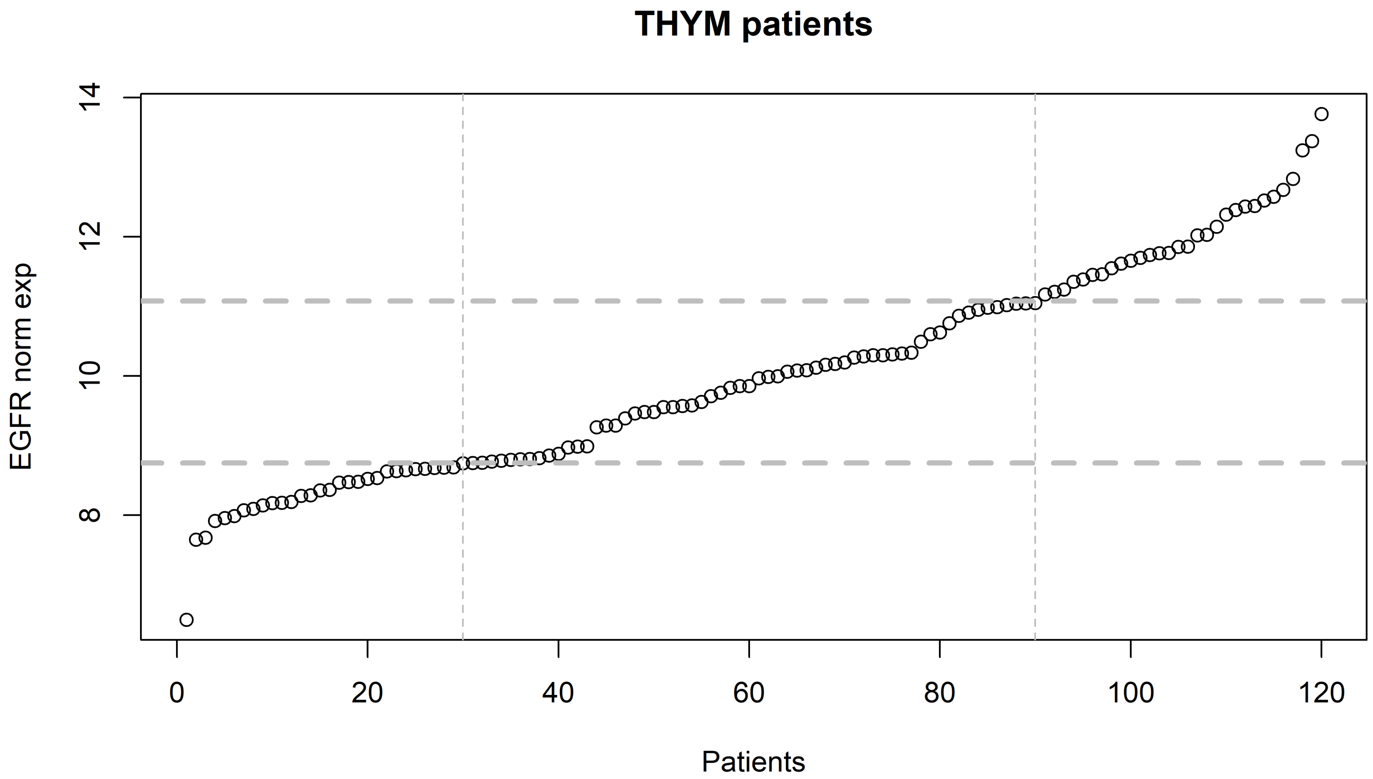

THYM patients
EGFR norm exp
Patients
0
20
40
60
80
100
120
8
10
12
14

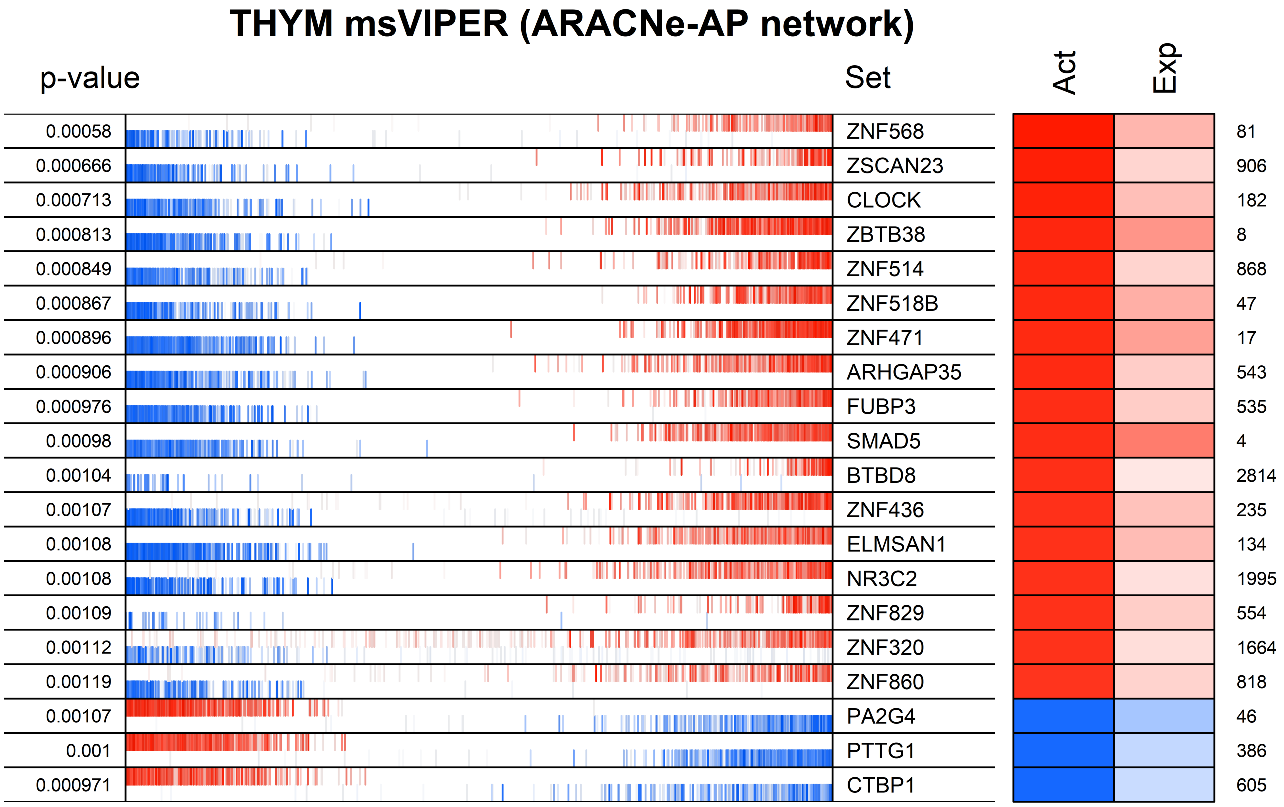
THYM msVIPER (ARACNe-AP network)
p-value
Set
Act
Exp
0.00058 ZNF568 81
0.000666 ZSCAN23 906
0.000713 CLOCK 182
0.000813 ZBTB38 8
0.000849 ZNF514 868
0.000867 ZNF518B 47
0.000896 ZNF471 17
0.000906 ARHGAP35 543
0.000976 FUBP3 535
0.00098 SMAD5 4
0.00104 BTBD8 2814
0.00107 ZNF436 235
0.00108 ELMSAN1 134
0.00108 NR3C2 1995
0.00109 ZNF829 554
0.00112 ZNF320 1664
0.00119 ZNF860 818
0.00107 PA2G4 46
0.001 PTTG1 386
0.000971 CTBP1 605

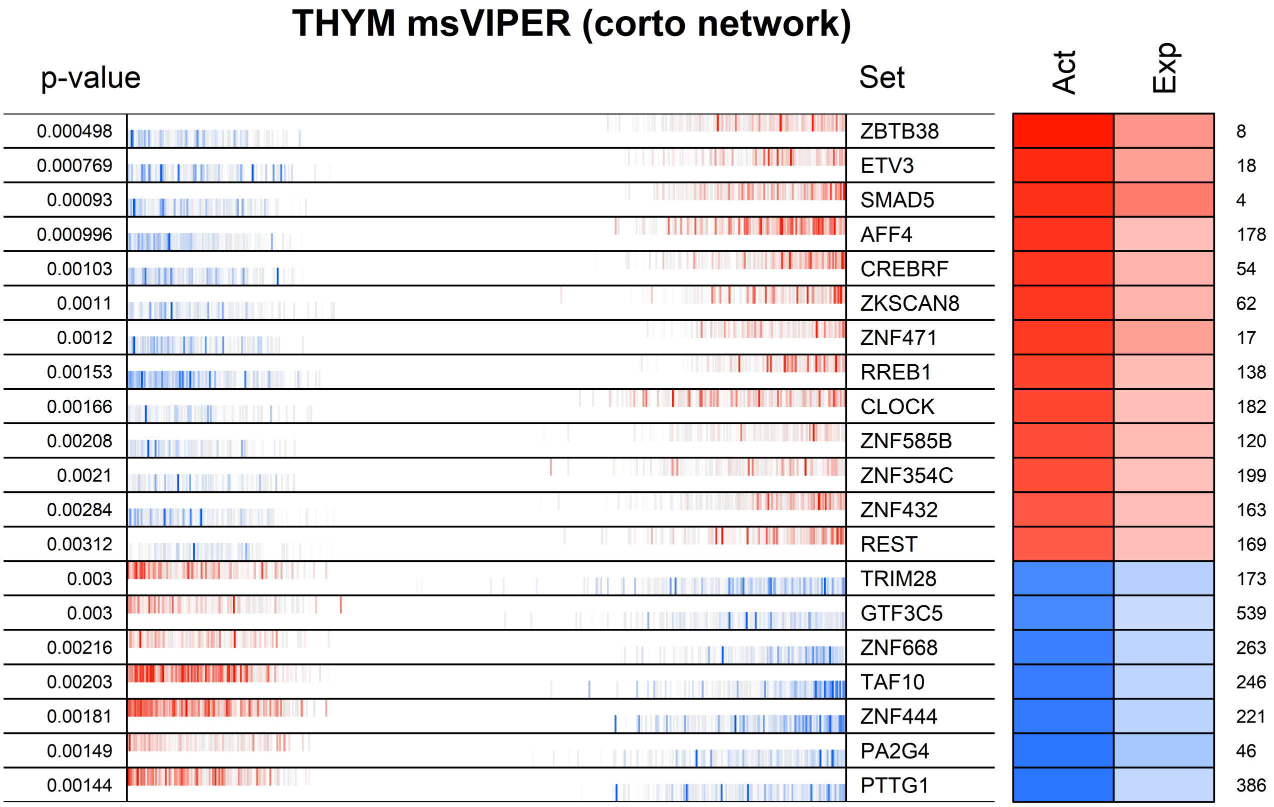
THYM msVIPER (corto network)
p-value
Set
Act
Exp
0.000498
ZBTB38
8
0.000769
ETV3
18
0.00093
SMAD5
4
0.000996
AFF4
178
0.00103
CREBRF
54
0.0011
ZKSCAN8
62
0.0012
ZNF471
17
0.00153
RREB1
138
0.00166
CLOCK
182
0.00208
ZNF585B
120
0.0021
ZNF354C
199
0.00284
ZNF432
163
0.00312
REST
169
0.003
TRIM28
173
0.003
GTF3C5
539
0.00216
ZNF668
263
0.00203
TAF10
246
0.00181
ZNF444
221
0.00149
PA2G4
46
0.00144
PTTG1
386

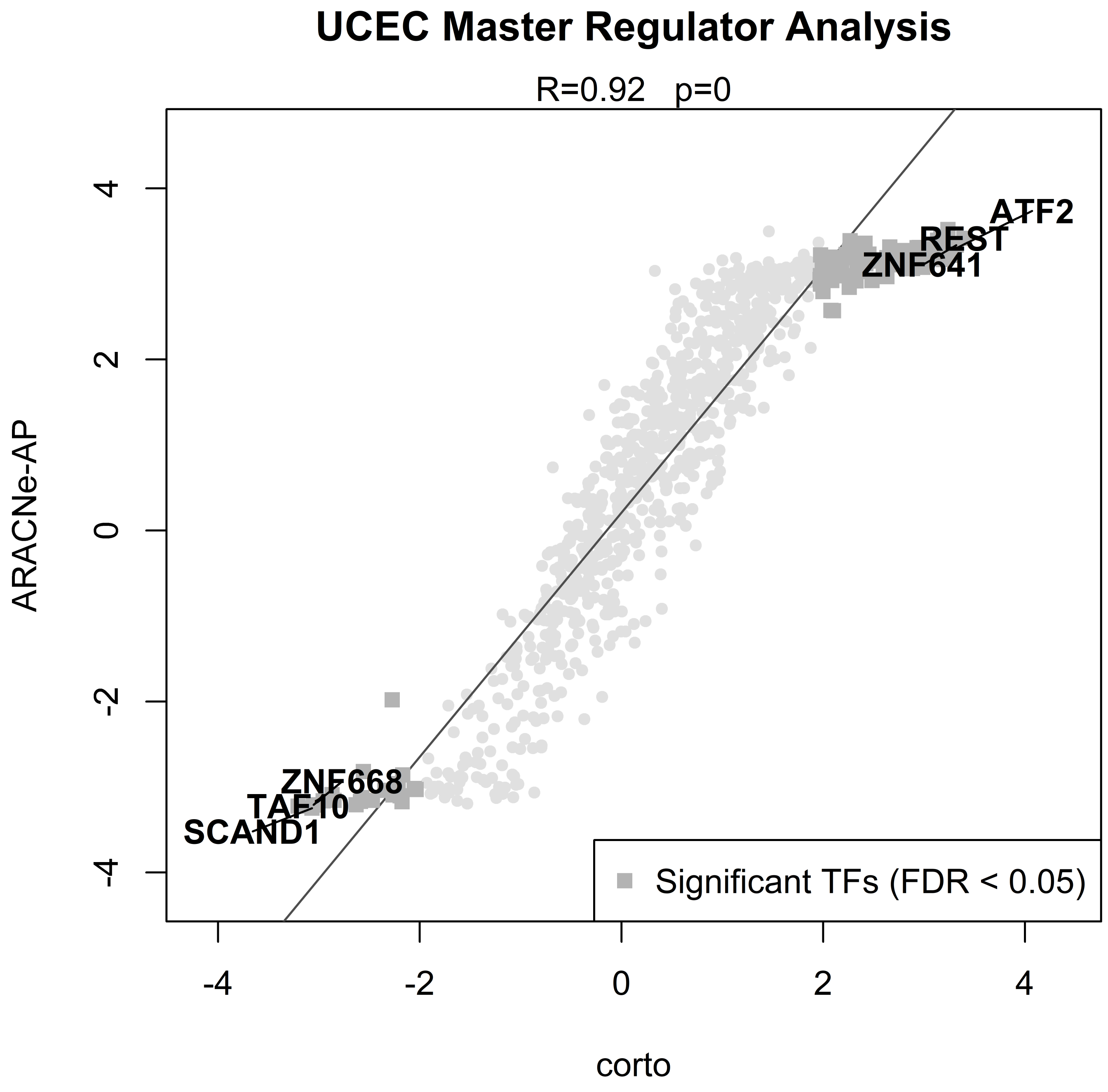
UCEC Master Regulator Analysis
R=0.92 p=0
ARACNe-AP
corto
-4
-2
0
2
4
ATF2
REST
ZNF641
ZNF668
TAF10
SCAND1
Significant TFs (FDR < 0.05)

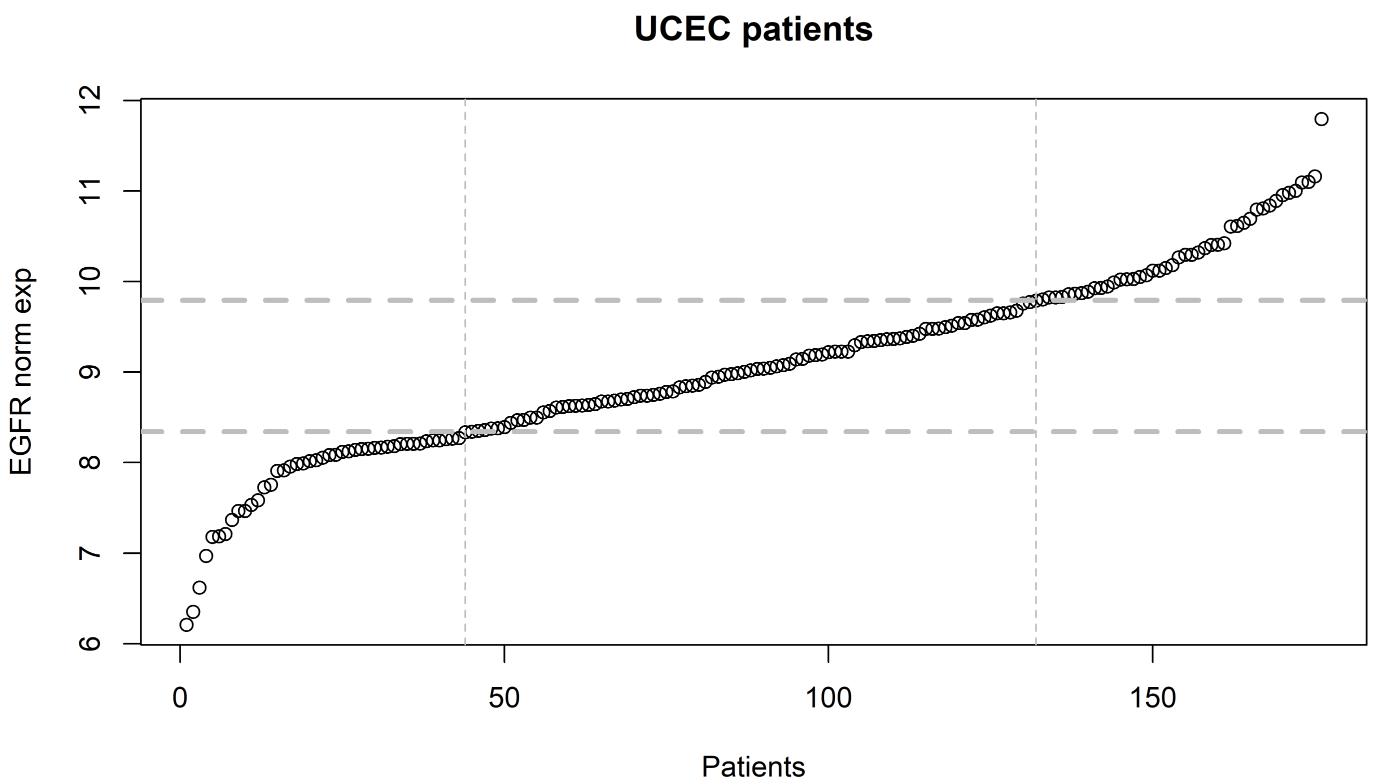
UCEC patients
EGFR norm exp
6
7
8
9
10
11
12
0
50
100
150
Patients

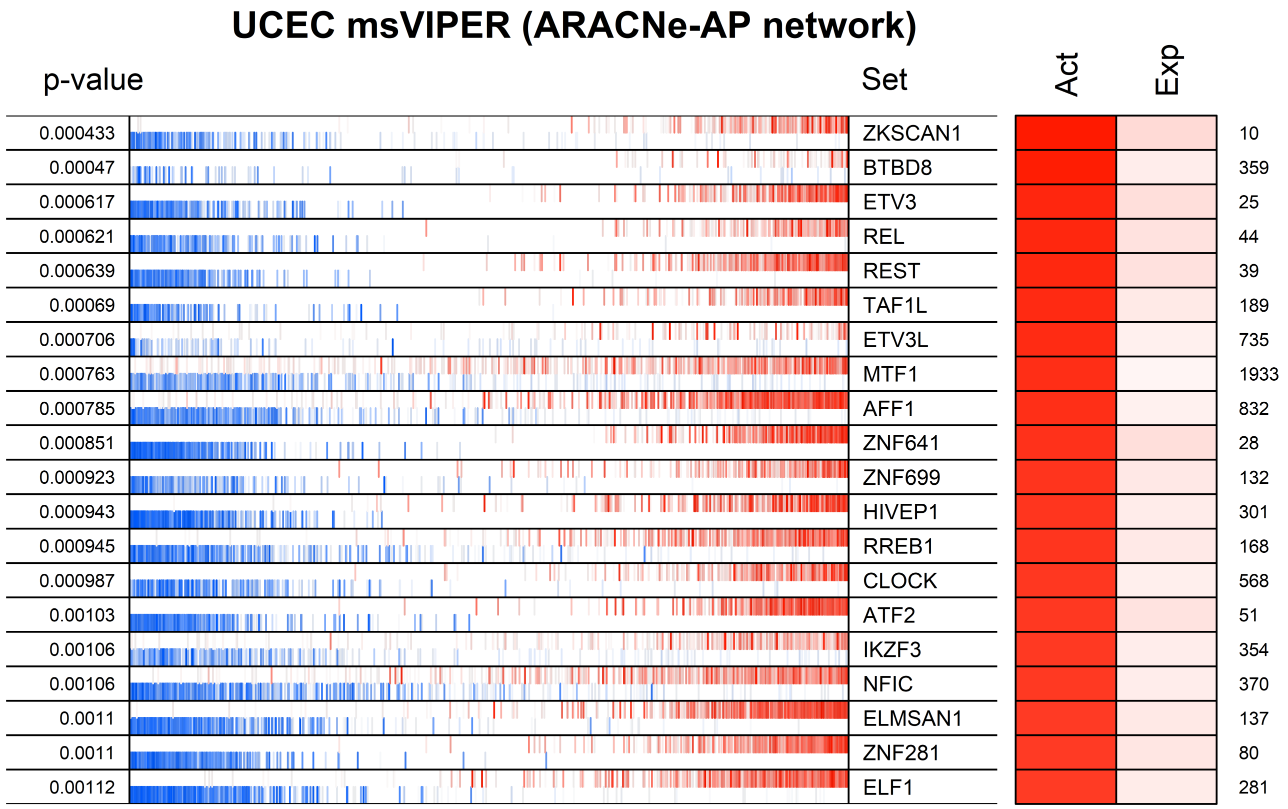
UCEC msVIPER (ARACNe-AP network)
p-value
Set
Act
Exp
0.000433 ZKSCAN1 10
0.00047 BTBD8 359
0.000617 ETV3 25
0.000621 REL 44
0.000639 REST 39
0.00069 TAF1L 189
0.000706 ETV3L 735
0.000763 MTF1 1933
0.000785 AFF1 832
0.000851 ZNF641 28
0.000923 ZNF699 132
0.000943 HIVEP1 301
0.000945 RREB1 168
0.000987 CLOCK 568
0.00103 ATF2 51
0.00106 IKZF3 354
0.00106 NFIC 370
0.0011 ELMSAN1 137
0.0011 ZNF281 80
0.00112 ELF1 281

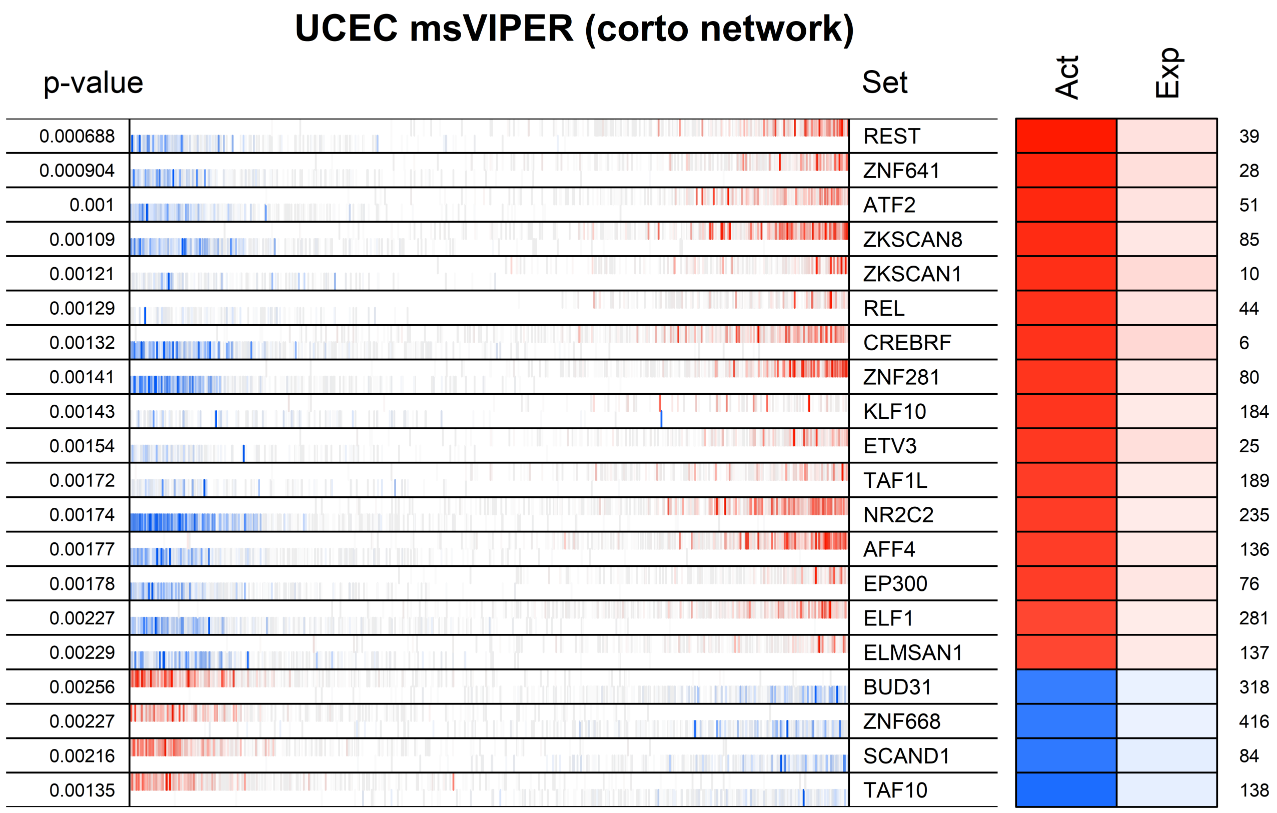
UCEC msVIPER (corto network)
p-value
Set
Act
Exp
0.000688
0.000904
0.001
0.00109
0.00121
0.00129
0.00132
0.00141
0.00143
0.00154
0.00172
0.00174
0.00177
0.00178
0.00227
0.00229
0.00256
0.00227
0.00216
0.00135
REST
ZNF641
ATF2
ZKSCAN8
ZKSCAN1
REL
CREBRF
ZNF281
KLF10
ETV3
TAF1L
NR2C2
AFF4
EP300
ELF1
ELMSAN1
BUD31
ZNF668
SCAND1
TAF10
39
28
51
85
10
44
6
80
184
25
189
235
136
76
281
137
318
416
84
138

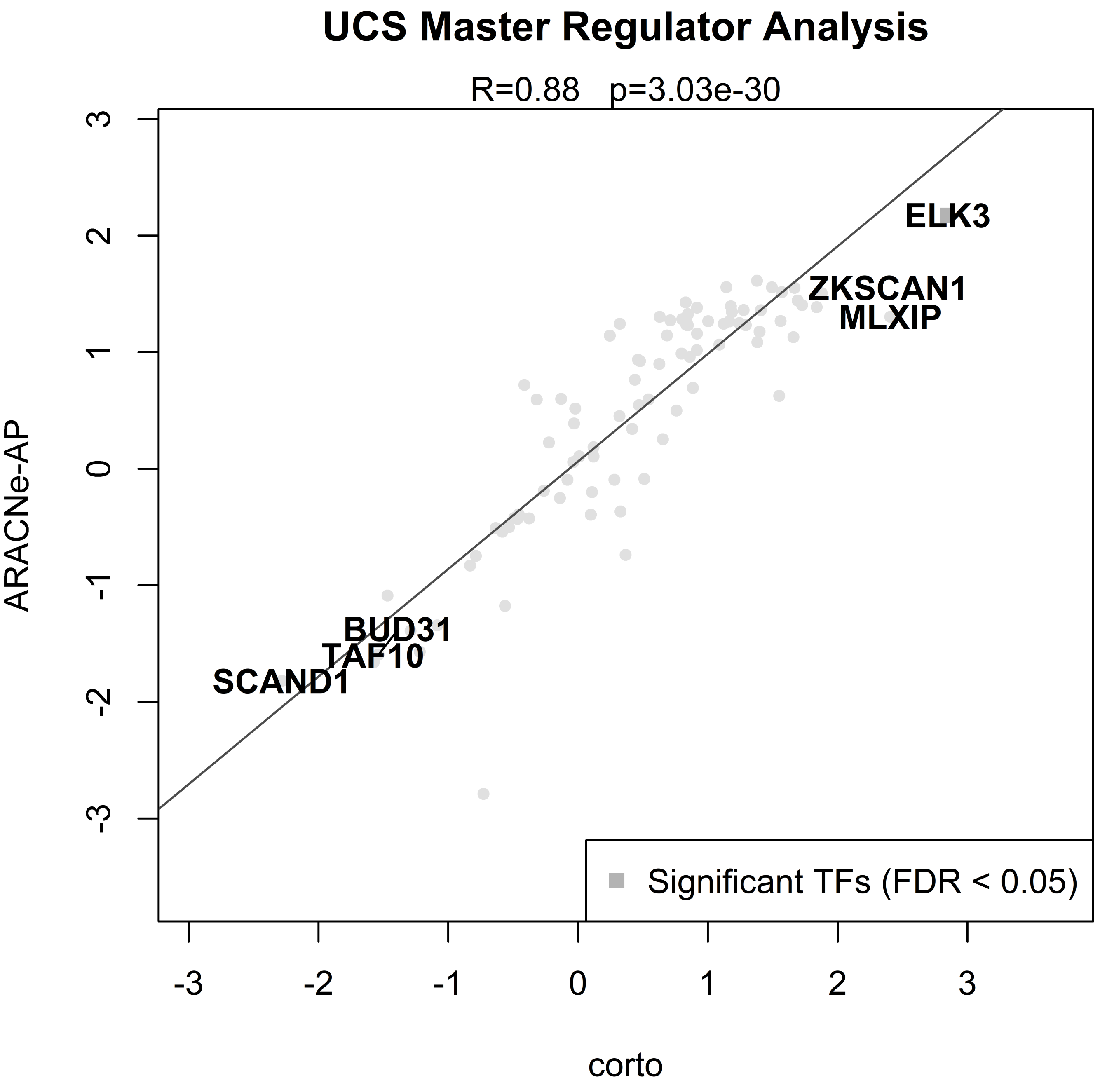

UCS Master Regulator Analysis
R=0.88 p=3.03e-30
ARACNe-AP
corto
ELK3
ZKSCAN1
MLXIP
BUD31
TAF10
SCAND1
Significant TFs (FDR < 0.05)
3
2
1
0
-1
-2
-3

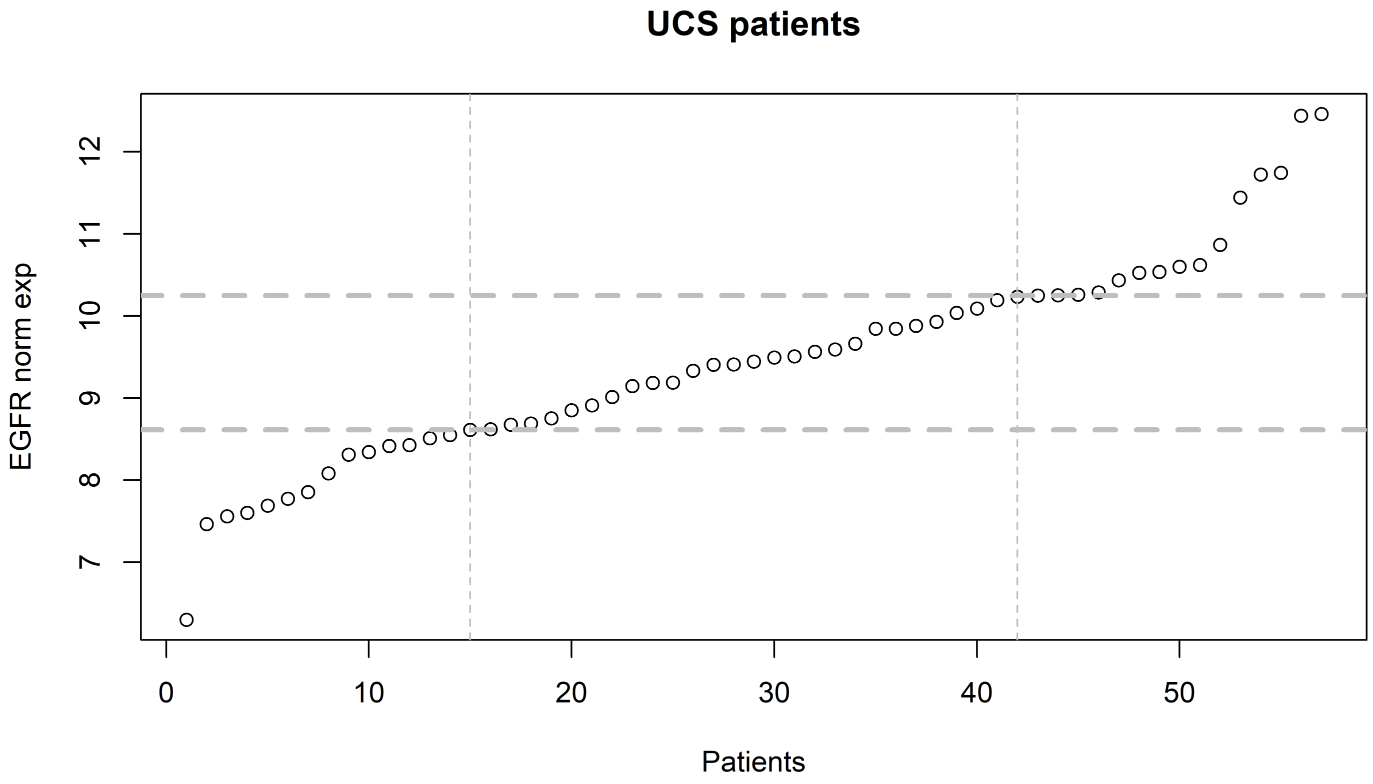
UCS patients
EGFR norm exp
Patients
7
8
9
10
11
12
0
10
20
30
40
50

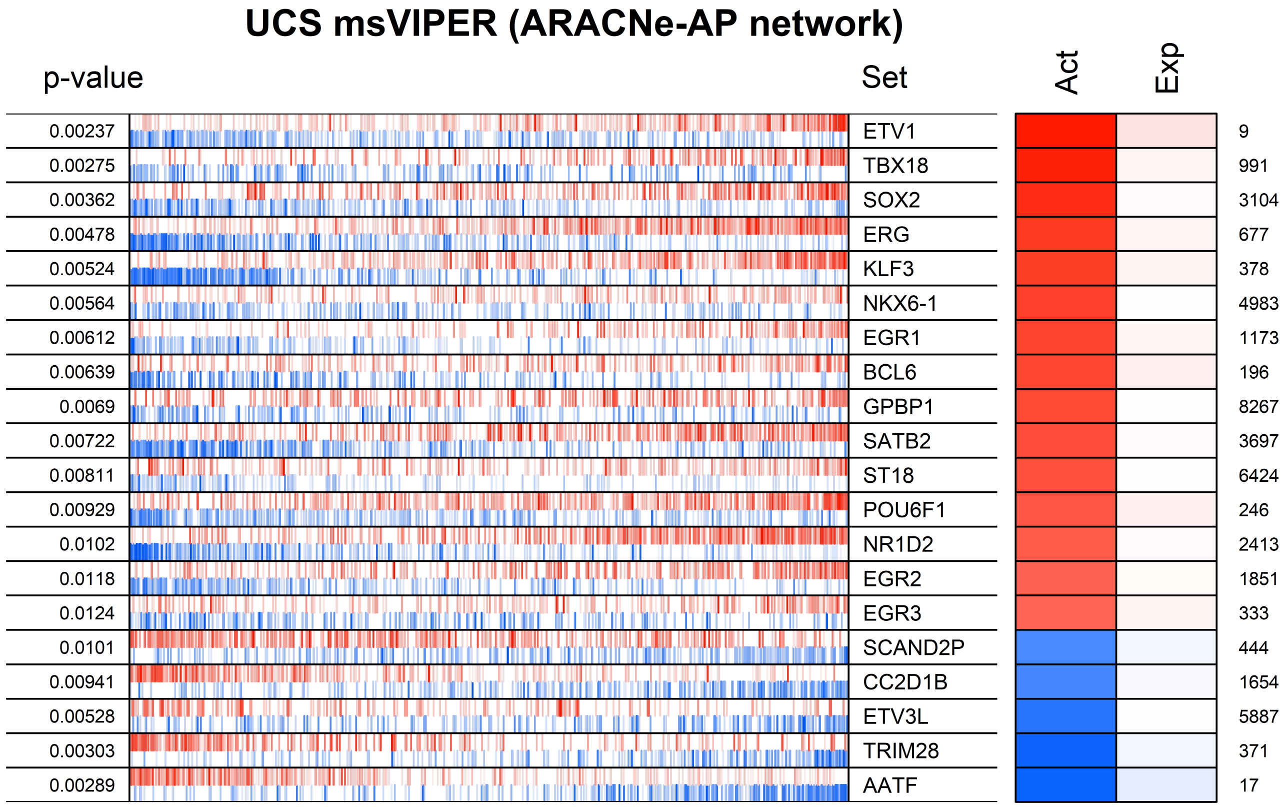
UCS msVIPER (ARACNe-AP network)
p-value
Set
Act
Exp
0.00237 ETV1 9
0.00275 TBX18 991
0.00362 SOX2 3104
0.00478 ERG 677
0.00524 KLF3 378
0.00564 NKX6-1 4983
0.00612 EGR1 1173
0.00639 BCL6 196
0.0069 GPBP1 8267
0.00722 SATB2 3697
0.00811 ST18 6424
0.00929 POU6F1 246
0.0102 NR1D2 2413
0.0118 EGR2 1851
0.0124 EGR3 333
0.0101 SCAND2P 444
0.00941 CC2D1B 1654
0.00528 ETV3L 5887
0.00303 TRIM28 371
0.00289 AATF 17

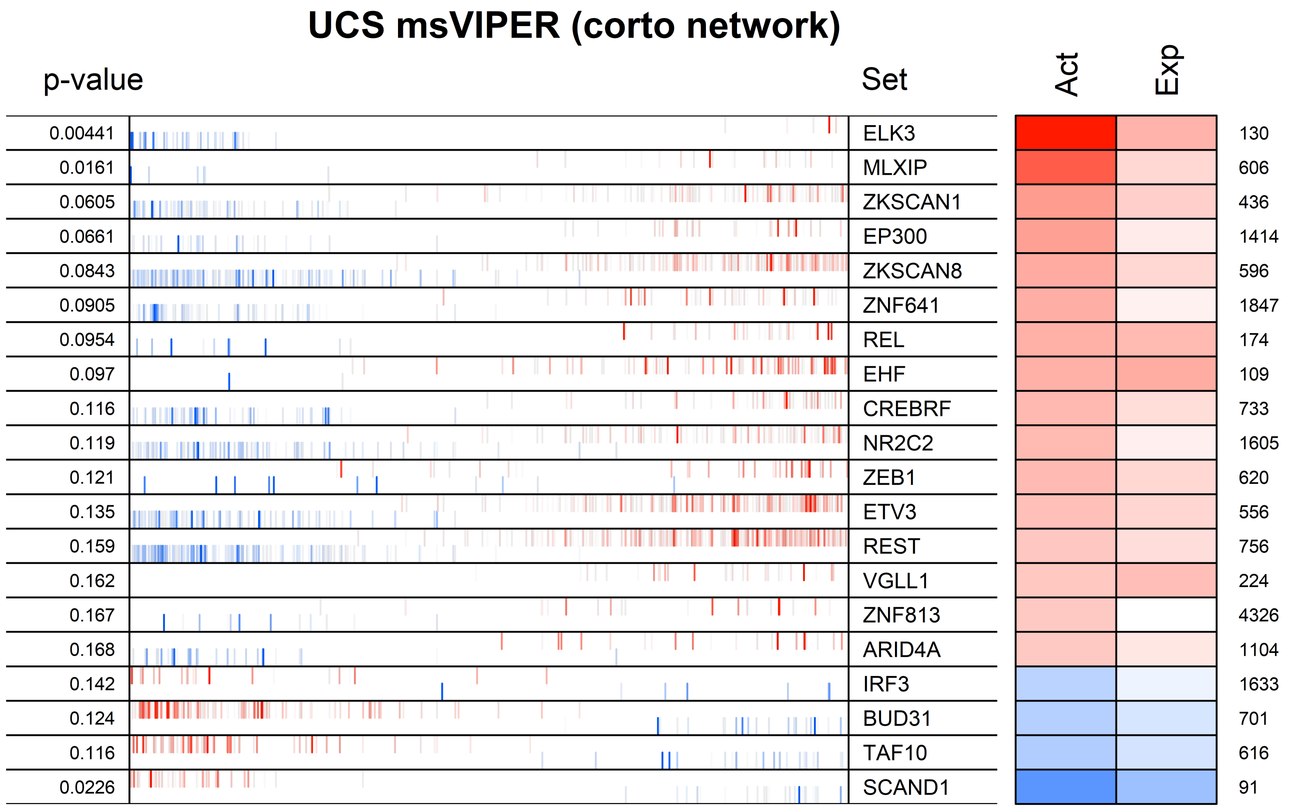
UCS msVIPER (corto network)
p-value
Set
Act
Exp
0.00441 ELK3 130
0.0161 MLXIP 606
0.0605 ZKSCAN1 436
0.0661 EP300 1414
0.0843 ZKSCAN8 596
0.0905 ZNF641 1847
0.0954 REL 174
0.097 EHF 109
0.116 CREBRF 733
0.119 NR2C2 1605
0.121 ZEB1 620
0.135 ETV3 556
0.159 REST 756
0.162 VGLL1 224
0.167 ZNF813 4326
0.168 ARID4A 1104
0.142 IRF3 1633
0.124 BUD31 701
0.116 TAF10 616
0.0226 SCAND1 91

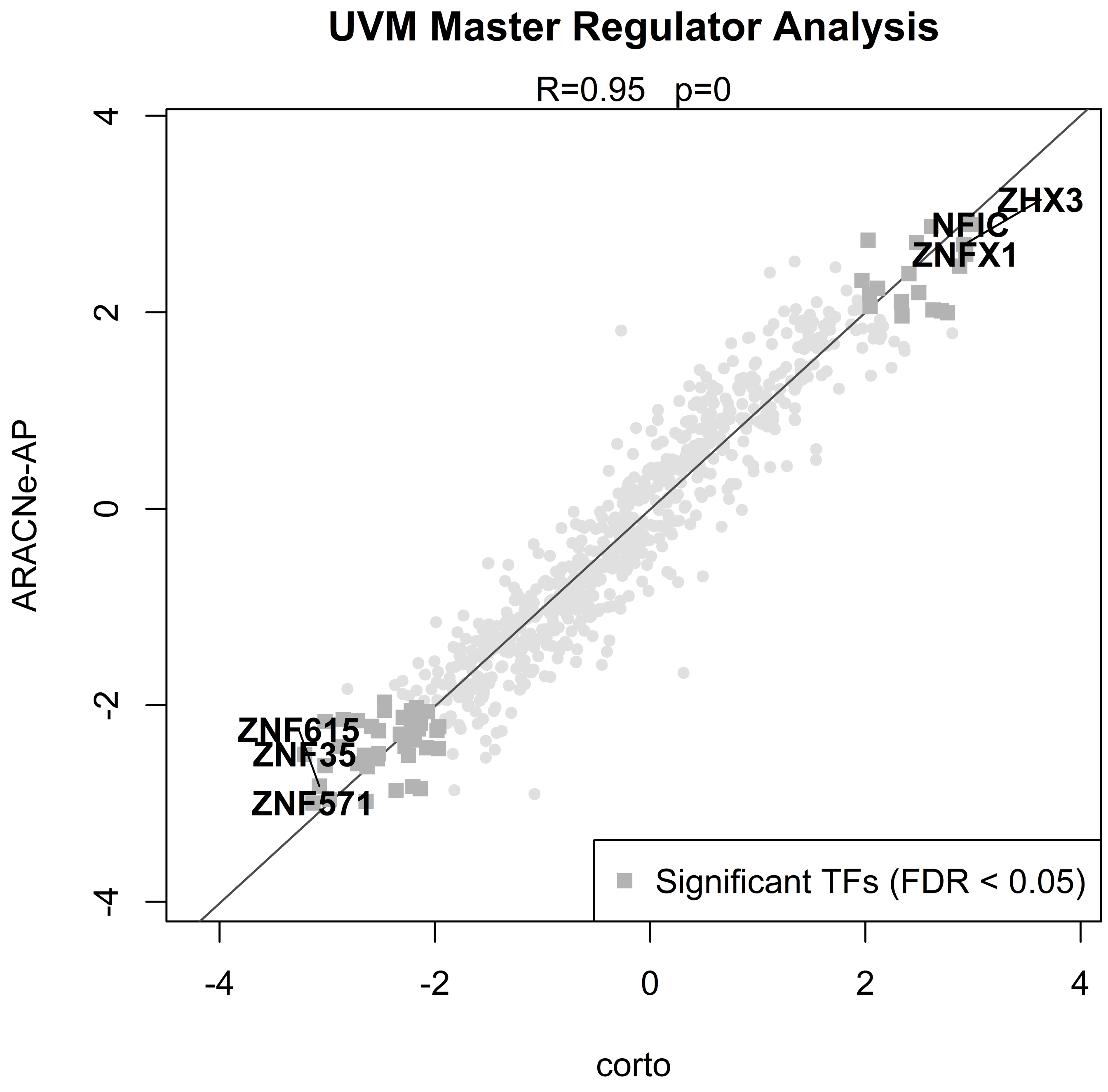

UVM Master Regulator Analysis
R=0.95 p=0
ZHX3
NFIC
ZNFX1
ZNF615
ZNF35
ZNF571
Significant TFs (FDR < 0.05)
ARACNe-AP
corto
4
2
0
-2
-4
-4
-2
0
2
4

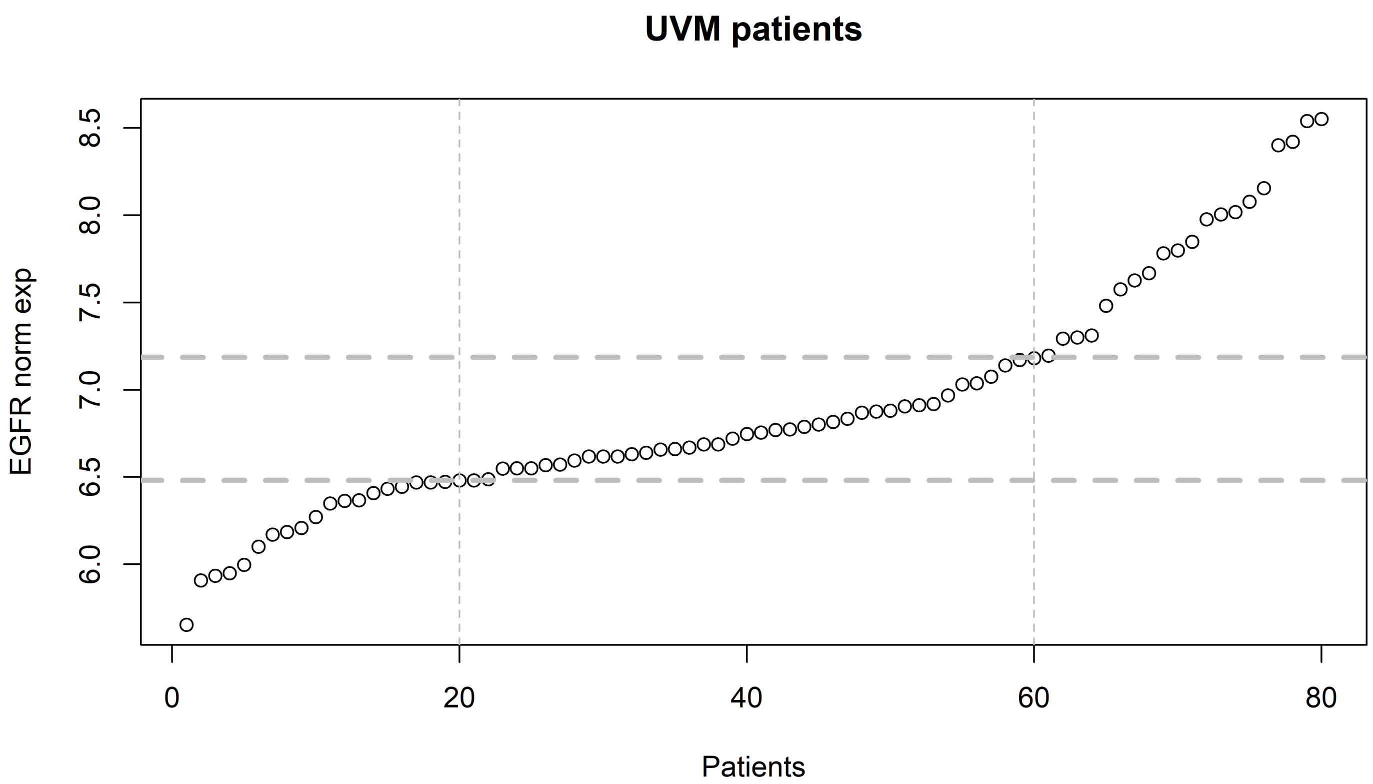
UVM patients
EGFR norm exp
8.5
8.0
7.5
7.0
6.5
6.0
0
20
40
60
80
Patients

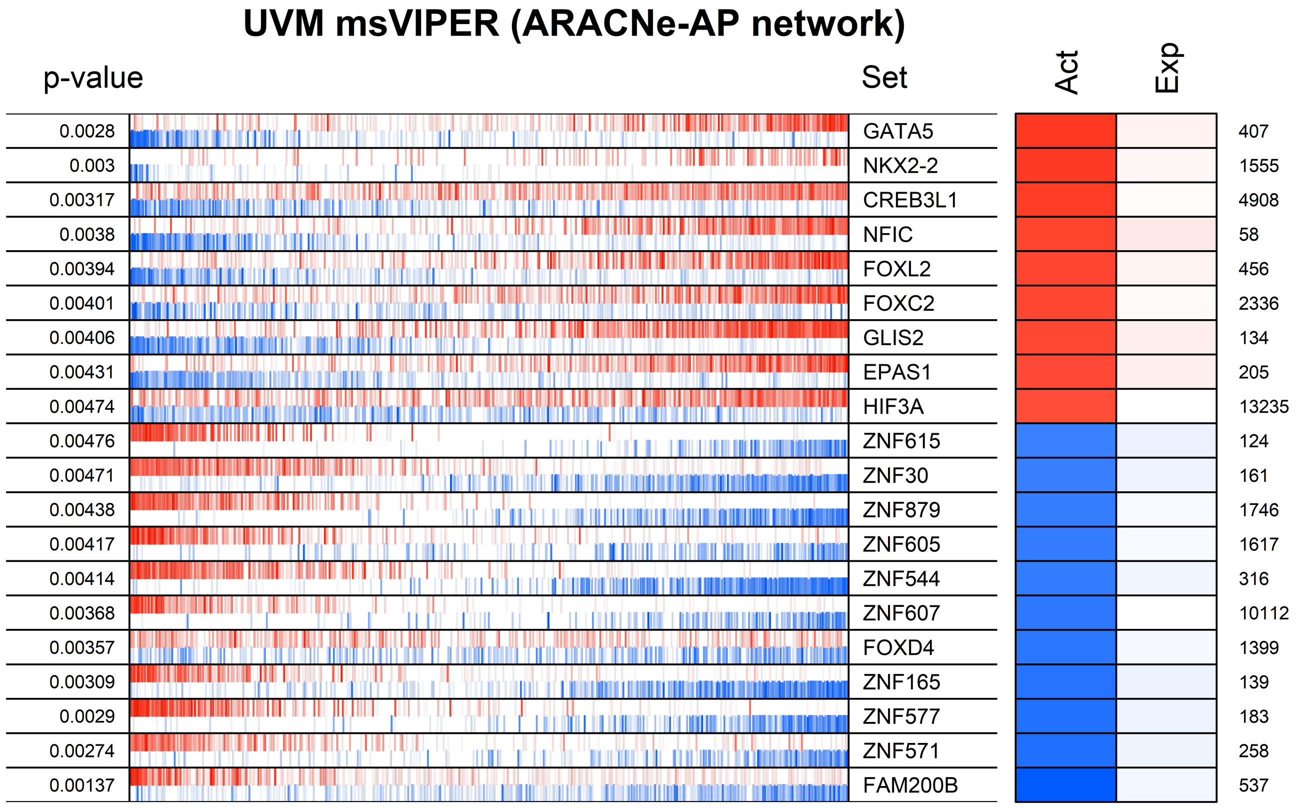
UVM msVIPER (ARACNe-AP network)
p-value
Set
Act
Exp
0.0028
GATA5
407
0.003
NKX2-2
1555
0.00317
CREB3L1
4908
0.0038
NFIC
58
0.00394
FOXL2
456
0.00401
FOXC2
2336
0.00406
GLIS2
134
0.00431
EPAS1
205
0.00474
HIF3A
13235
0.00476
ZNF615
124
0.00471
ZNF30
161
0.00438
ZNF879
1746
0.00417
ZNF605
1617
0.00414
ZNF544
316
0.00368
ZNF607
10112
0.00357
FOXD4
1399
0.00309
ZNF165
139
0.0029
ZNF577
183
0.00274
ZNF571
258
0.00137
FAM200B
537

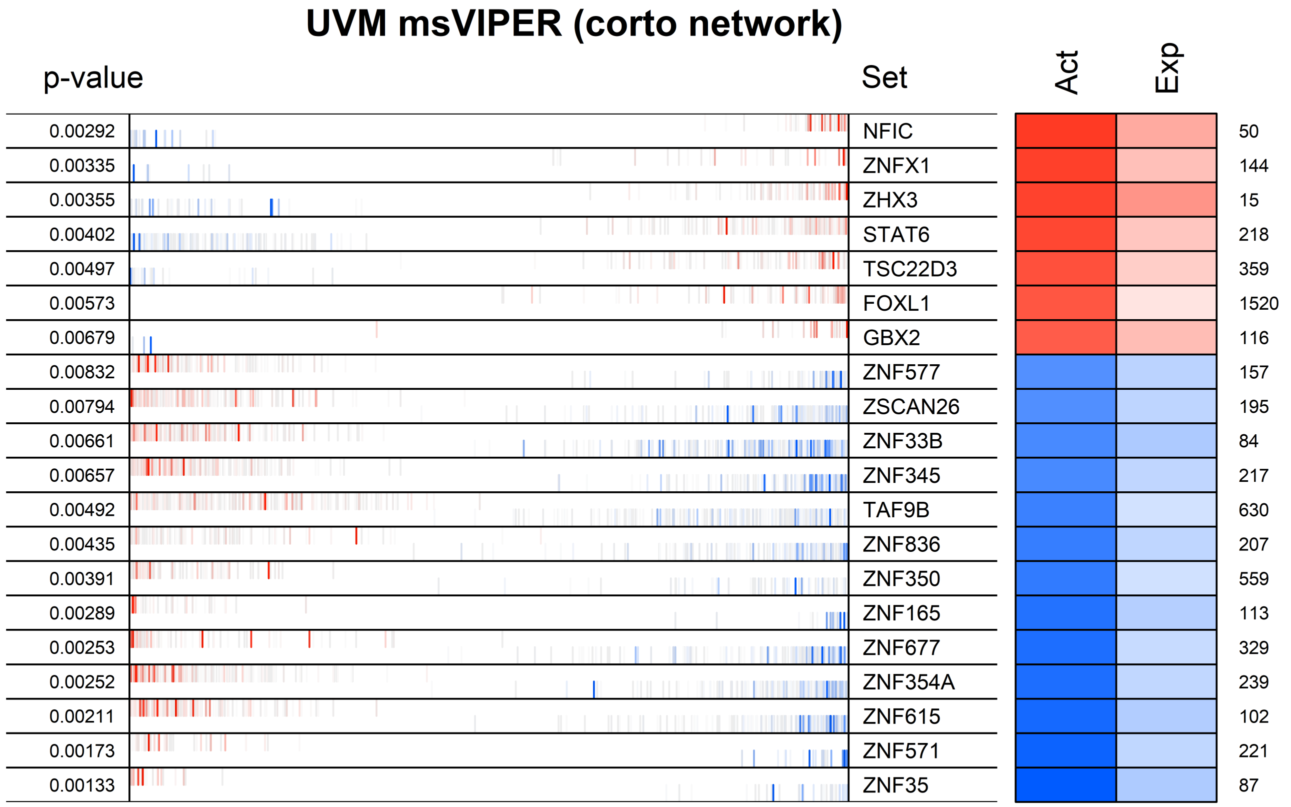
UVM msVIPER (corto network)
p-value
Set
Act
Exp
0.00292 NFIC 50
0.00335 ZNFX1 144
0.00355 ZHX3 15
0.00402 STAT6 218
0.00497 TSC22D3 359
0.00573 FOXL1 1520
0.00679 GBX2 116
0.00832 ZNF577 157
0.00794 ZSCAN26 195
0.00661 ZNF33B 84
0.00657 ZNF345 217
0.00492 TAF9B 630
0.00435 ZNF836 207
0.00391 ZNF350 559
0.00289 ZNF165 113
0.00253 ZNF677 329
0.00252 ZNF354A 239
0.00211 ZNF615 102
0.00173 ZNF571 221
0.00133 ZNF35 87
