## Supplementary material for "*corto*: a lightweight R package for Gene Network Inference and Master Regulator Analysis": Supp Vignette

- Supplementary Vignette 1, on Master Regulator Analysis on a MYCN amplification signature in Neuroblastoma: [link here](https://www.dropbox.com/s/3ltiftpz6nx0wfu/SuppVignette1_mycnamp.zip?dl=0)
- Supplementary Vignette 2, about *corto* operating on parallel RNA-Seq and ATAC-Seq signatures in Fulvestrant-treated cells: [link here](https://www.dropbox.com/s/da289ku3ugq55wl/SuppVignette2_atac.zip?dl=0)
- Supplementary Vignette 3, about running *corto* and other tools to reconstruct gene networks in a pure R environment: [link here](https://www.dropbox.com/s/42ejtl6b0alx0d3/SuppVignette3_moretools.zip?dl=0)
- Supplementary Vignette 4, about comparing *corto* and *ARACNe-AP* in Master Regulator Analysis: [link here](https://www.dropbox.com/s/wcyjmwc3j18lmg0/SuppVignette4_mracomparison.zip?dl=0)
