## Supplementary figures and images for "*corto*: a lightweight R package for Gene Network Inference and Master Regulator Analysis"

### Supp Figure S1

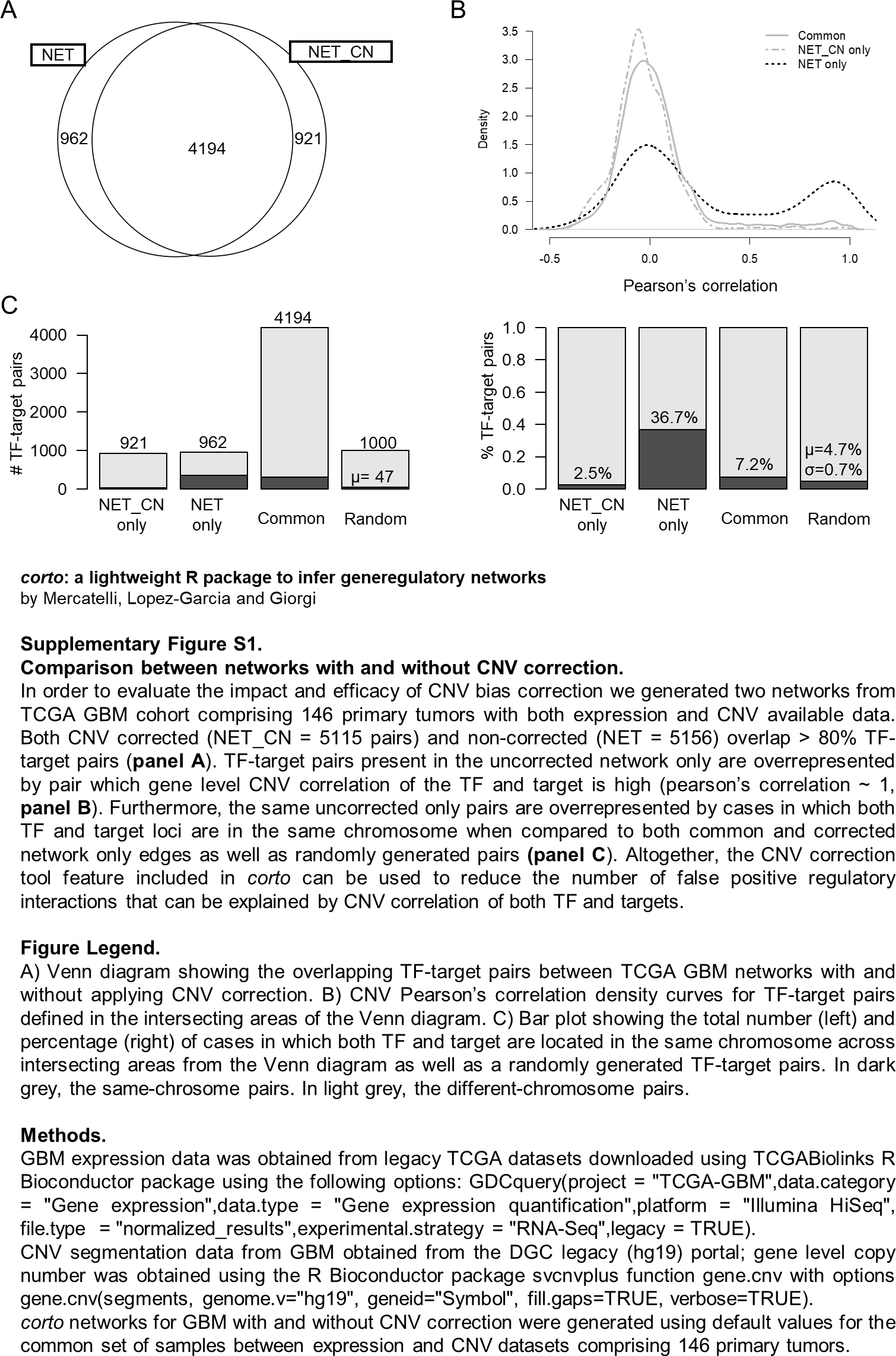
